# GW182 silencing domain engages the tristetraprolin-binding pocket of CNOT1 to recruit CNOT1 into multiprotein condensates

**DOI:** 10.64898/2026.07.16.738963

**Authors:** Michał K. Białobrzewski, Maja K. Cieplak-Rotowska, Agnieszka Michaś, Zuzanna Staszałek, Nahum Sonenberg, Michal Dadlez, Anna Niedzwiecka

## Abstract

GW182 recruits the CCR4-NOT deadenylase complex through its CIM1 region, yet the CNOT1 surface mediating this interaction has remained unknown. Using hydrogen– deuterium exchange mass spectrometry, we map CIM1 binding to a hydrophobic groove within CNOT1 residues 800-999. Unexpectedly, this groove coincides with the tristetraprolin (TTP)-binding pocket. CIM1 and the C-terminal TTP peptide engage this surface through a shared RL(P/ζ)XΩ sequence pattern despite lacking overall homology, suggesting a convergently evolved short linear motif (SLiM). We further show that the GW182 silencing domain (GW182 SD) and TTP compete for CNOT1 binding using orthogonal *in vitro* assays, including a reconstituted liquid-liquid phase separation system in which GW182 SD acts as a scaffold and recruits CNOT1 as a client through specific binding. These findings define a shared CNOT1 recognition site and reveal SLiM-mediated competition as a molecular principle governing CCR4-NOT engagement, with consequences for GW182-driven silencing condensate assembly *in vitro*.

**Significance:** GW182 silencing domain binds the tristetraprolin pocket of CNOT1 and recruits it into multiprotein condensates, providing a biophysical framework for coordinated miRNA- and ARE-mediated gene expression regulation.

## Introduction

Post-transcriptional regulation *via* mRNA decay is a central layer of eukaryotic gene expression and is orchestrated by multiple cellular pathways that dynamically remodel the composition of multiprotein complexes. Specific interactions between *cis*-acting elements located in the mRNA 3’ untranslated regions (UTRs), including microRNA (miRNA) binding sites and AU-rich elements (AREs), and their cognate effectors, miRNAs or ARE-binding proteins (ARE-BPs), impact mRNA translation, subcellular localization and stability through distinct yet interconnected mechanisms^1–3^. Moreover, recent transcriptome-wide mapping shows that discrete protein-and miRNA-associated RNA elements can recruit CCR4–NOT and regulate gene expression, underscoring the diversity of sequence-defined routes to CCR4–NOT engagement^4^.

AREs present in mRNAs encoding nuclear transcription factors, proto-oncogenes, and cytokines are specifically recognized by tristetraprolin (TTP)^5^, a member of the tandem zinc finger proteins family^6^. TTP promotes rapid mRNA decay and modulates signaling pathways by directly interacting with the multisubunit CCR4-NOT deadenylase complex^6–8^, a master regulator of mRNA stability and translation^9^. While the molecular architecture of CCR4-NOT was resolved only for *Schizosaccharomyces pombe* using cryo-EM^10^, studies in mammals have also established that its central subunit, CNOT1, serves as a scaffold to which the other components are anchored^11–18^. A fragment of the N-terminal region of CNOT1 (residues 800-999; Fig. 1a) interacts with the conserved C-terminal sequence of TTP (residues 315-323) with an affinity of ∼2 μM and is required for ARE-mediated deadenylation^7^.

**Figure 1.**
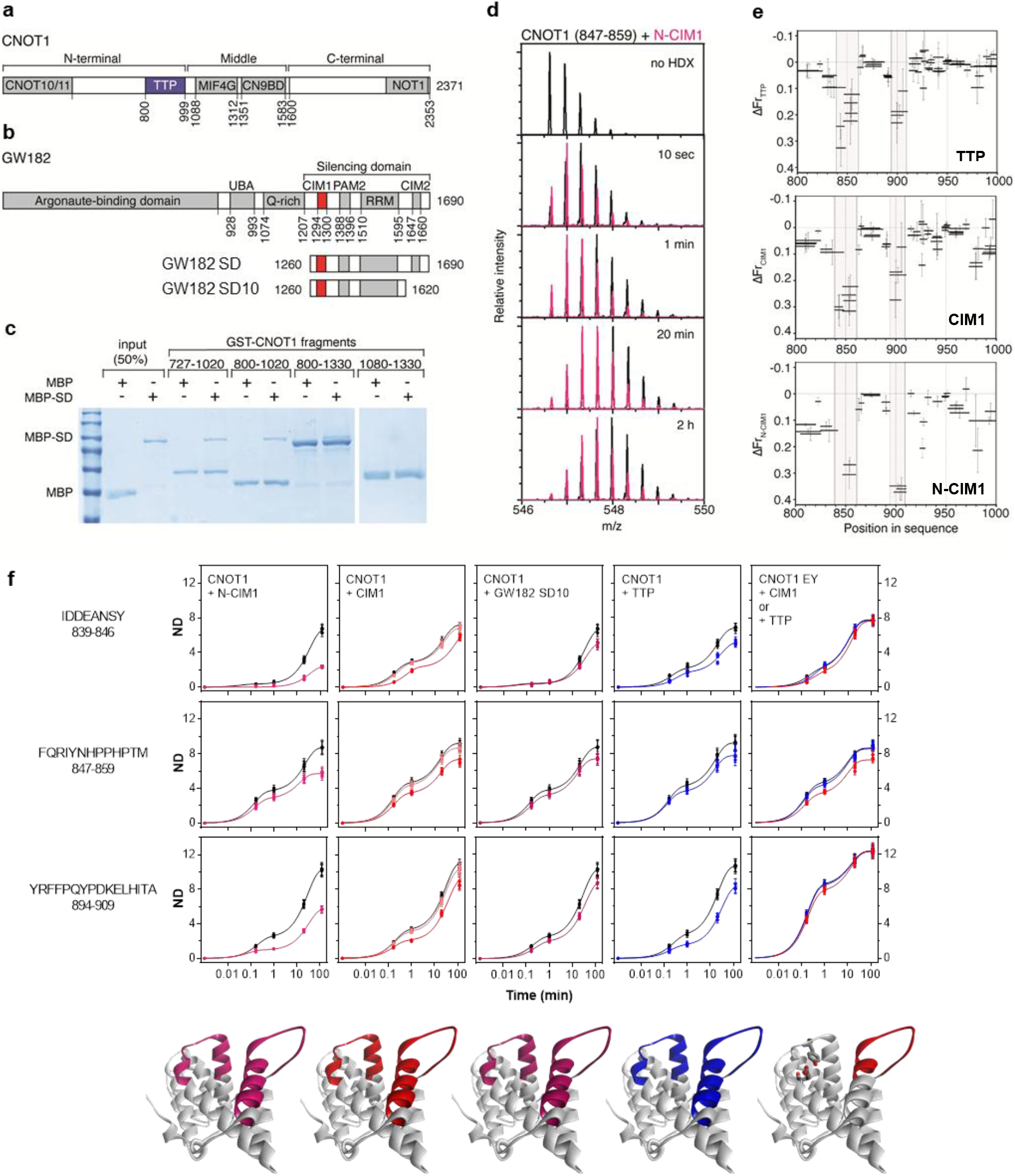
GW182 SD interacts with the tristetraprolin-binding domain of CNOT1. Schematic diagrams of domain composition of **a**, human CNOT1 (isoform 2), and **b,** GW182 (TNRC6C isoform 1), its silencing domain (GW182 SD), and a C-terminally truncated SD fragment (GW182 SD10). **c**, SDS PAGE of GST pulldown assay showing that MBP-tagged GW182 SD interacts with CNOT1(800-1020). MBP alone was used as a negative control. **d**, Mass spectra of 847-859 peptide of CNOT1(800-999) alone (black) and in the presence of N-CIM1 peptide at 140 µM (red) after different deuteration times. **e**, HDX differences (ΔFr) for pepsin-generated CNOT1 peptides in the presence of TTP (3.3 µM, upper panel), CIM1 (126 µM middle panel), and N-CIM1 (140 µM bottom panel) peptides *vs*. the *apo* state after 2 h of deuteration. Vertical thin lines show the standard deviation, n=3. Two regions of the highest protection against HDX in the presence of TTP-or GW182SD-derived peptides are marked grey. **f**, HDX kinetics for selected CNOT1 peptides showing the largest changes in in deuterium uptake (ND) between the *apo* state (black) and in the presence of the following binding partners, from left to right: N-CIM1 peptide (140 µM, magenta); CIM1 peptide (12.6 µM, orange; 126 µM, red); GW182 SD10 protein (6 µM, dark magenta); and TTP peptide (3.3 µM, blue). Right panel, HDX kinetics of CNOT1 E893A/Y900A in the presence of CIM1 peptide (126 µM, red) or TTP peptide (3.3 µM, blue). Interaction with TTP peptide is completely abolished, while a residual evidence of CIM1 binding is visible for the 847-859 CNOT1 fragment. (Bottom) Structure of CNOT1(800-999) (pdb 4J8S), with colored segments marking peptic peptides protected from HDX in the presence of each ligand.

In parallel, miRNA-dependent gene silencing also involves CCR4-NOT-mediated deadenylation^19,20^, in addition to inhibition of translation initiation^13,21–23^. In this pathway, CCR4-NOT is recruited to target mRNAs through the successive actions of the miRNA^24^, an Argonaute (Ago) protein, and the Ago-binding GW-rich proteins of 182 kDa (GW182). In mammals, GW182 is encoded as three paralogs (TNRC6A, TNRC6B and TNRC6C)^25^. GW182 directly interacts with CCR4-NOT^14,15,21,26,27^ and is an essential component of the miRNA-induced silencing complex (miRISC)^28^ that organizes higher-order assemblies containing Ago and CCR4-NOT^15,21,26,29,30^. Although largely functionally redundant in cell culture systems^31^, the paralogs are not interchangeable *in vivo*, since knockout of TNRC6A, - B or-C results in distinct phenotypes, and loss of individual paralogs is lethal^32^. Consistently, haploinsufficient variants in TNRC6B have been linked to a human genetic disorder characterized by neurocognitive and behavioral abnormalities^33^. The intrinsically disordered C-terminal domain of GW182^34^ is critical for silencing due to its involvement in both deadenylation^26,35^ and translation repression^35^. Within the silencing domain (SD) of GW182 (TNRC6 C, Fig. 1b), two CCR4-NOT-interacting motifs (CIM1, CIM2) were identified^26^, and CIM1 (^1294^QSRLPQW^1300^) was shown to interact directly with CNOT1^26^. However, the corresponding GW182-binding site of CNOT1 has not been delineated due to the large size of both protein and the intrinsically disordered nature of the GW182 SD^34^, which make structural studies challenging.

GW182 and Argonaute proteins are constitutive components of cytoplasmic processing bodies (P-bodies, PBs)^28,36,37^ that form upon activation of miRNA-mediated gene silencing^38^ and serve as sites of mRNA storage of translationally repressed transcripts^39–41^. PBs are membrane-less organelles (MLOs) with a distinct molecular composition^42^ that are spatially, compositionally and functionally linked with other MLOs, *i.e*., stress granules^35,42–44^. Like other MLOs, PBs form as biomolecular condensates *via* liquid-liquid phase separation (LLPS)^46^, driven primarily by the condensation of multivalent intrinsically disordered proteins (IDPs)^47,48^. Condensate components are dynamic, exchanging with the surrounding soluble pool, and either retain liquid-like properties or mature into gel-like and solid-like state^48,49^. Dysregulation of such IDP-driven condensation has been implicated in cancer, neurodegeneration, inflammation, and autoimmune disorders^50,51^.

Many proteins are incorporated into MLOs not through intrinsic phase-separating capacity, but *via* interactions with scaffold proteins that drive the phase transition^52^. These client proteins are recruited through networks of homotypic and heterotypic interactions^53^. Recently, the N-terminal Ago-binding domain (ABD) of GW182 (TNRC6B) was shown to function as a condensate scaffold that recruits Ago2 as a client, promoting miRISC phase separation both *in vitro* and in cells^54^ leading to deadenylase recruitment and accelerated deadenylation of targeted mRNAs^54^. In contrast, the contribution of the C-terminal GW182 silencing domain to homotypic and heterotypic condensate formation remained unexplored. Since quantitative analysis of P-body components through *in vitro* reconstitution of multicomponent condensates is a major focus of current research^55^, we address the mechanistic basis of multiprotein condensate formation involving the human GW182 SD and a fragment of CNOT1.

It is increasingly evident that 3’ UTR-mediated silencing pathways converge at multiple levels. First, both GW182^14,15^ and TTP^56^ interact with the CNOT9 subunit of CCR4-NOT *via* two conserved tryptophan-binding pockets, enabling deadenylation. Second, GW182 and TTP participate in transcript-specific 4EHP/GIGYF2/DDX6-and 4EHP/4E-T/DDX6-dependent pathways that couple 5’ translational repression with 3’ mRNA decay^56–59^. This coupling between translational control and mRNA stability extends beyond GW182 and TTP, as illustrated by ZC3H7A/B RNA-binding proteins, which coordinate CCR4–NOT-dependent decay with GIGYF2/4EHP-mediated translational repression^61^. Additional convergence has recently been demonstrated for GW182, where the TRIM71 protein silences hairpin-containing 3’ UTR targets through an RNA-dependent interaction with GW182, competing with Ago and UPF1 in a transcript-selective manner^62^. Furthermore, a recent reconstitution study has indicated that RNA adaptors can engage CCR4–NOT through multivalent IDR contacts rather than single isolated motifs, and that phosphorylation can tune these interactions to modulate deadenylation rates, with related evidence for human TTP^63^.

Here, we combine hydrogen-deuterium exchange mass spectrometry (HDX-MS), fluorescence correlation spectroscopy (FCS), affinity chromatography, and liquid-liquid phase separation-based assays to reveal an additional point of convergence between GW182-and TTP-dependent silencing pathways. This work aims to elucidate how distinct silencing factors access CCR4-NOT through convergent minimal interaction motifs and to define the role of the GW182 silencing domain and CNOT1 in multiprotein condensate organization. The findings provide new insight into the regulation of miRNA-and ARE-mediated RNA silencing and reveal biophysical principles underlying protein-protein interactions that contribute to the formation of human P-bodies.

## Results

### GW182 silencing domain directly binds the TTP-binding region of CNOT1

To identify the GW182 SD binding site within CNOT1, we focused on the previously characterized TTP-binding region of CNOT1, since deletion of residues 727-1068 or 840-1068 was shown to strongly impair mRNA repression even in the presence of other conserved regions^14^. GST pulldown assays revealed that MBP-tagged GW182 SD (TNRC6C) is retained by GST-CNOT1 fragments encompassing residues 800-1020, 727-1020, and 800-1330, whereas no interaction is detected with CNOT1(1080-1330) (Fig. 1c). These results identify CNOT1(800-1020) as sufficient for GW182 SD binding. Subsequent analyses were therefore performed using the minimal CNOT1(800-999) fragment (referred to as CNOT1), whose structure in the complex with the C-terminal TTP peptide was previously resolved^7^.

### CIM1 of GW182 and TTP engage the same hydrophobic groove of CNOT1

Given the intrinsically disordered nature of the GW182 SD^34^ and the limited solubility of both interaction partners, we used hydrogen-deuterium exchange mass spectrometry (HDX-MS) as the method of choice to map the interaction surfaces^64–66^. HDX-MS experiments were performed on CNOT1 in the presence of peptides corresponding to GW182 CIM1 (QSRLPQWTHP, denoted to as CIM1)^26^, an N-terminally extended CIM1 peptide (N-CIM1; DPSQSQSRLPQWTHPN), the GW182 SD10 fragment (residues 1260-1620), or the TTP peptide (APRRLPIFNRISVSE, denoted as TTP)^7^ (Fig. 1d, e).

Both CIM1 and N-CIM1 peptides induced HDX protection patterns in CNOT1 that closely resembled those observed with TTP, with strongest protection localized to regions spanning residues 839-859 and 894-909 (Fig. 1e,f), in contrast to the unprotected regions (Fig. S2-S5).Specifically, each of the three ligands reduced HDX over identical peptic peptides spanning helices α1, α3, and α4, consistent with a shared binding surface or the same allosteric stabilization. These regions form the hydrophobic groove that accommodates TTP in the crystal structure^7^, suggesting that CIM1 of GW182 engages the same CNOT1 surface. In contrast, the CIM2 peptide^26^ had no detectable effect on CNOT1 HDX (Fig. S1a). HDX-MS further revealed similar bi-exponential exchange kinetics for CNOT1 bound to TTP, CIM1, N-CIM1, or GW182 SD10 across most peptic peptides (Fig. 1f, S2-S6, and Suppl. Tables T1-T5), supporting a shared binding mode for CIM1 and TTP on CNOT1.

The HDX protection of the CNOT1 binding groove by N-CIM1 was substantially stronger than that observed for the originally proposed CIM1 sequence^26^. This was evident from a consistent decrease in the number of deuterons taken up (ND, Fig. 1f) across CNOT1 peptides spanning residues 839-846, 847-859, and 894-909, with values of 2.44 ± 0.11, 5.72 ± 0.09, and 5.79 ± 0.06 for N-CIM1 compared with 6.74 ± 0.06, 7.34 ± 0.11, and 9.4 ± 0.2 for CIM1, respectively (mean ± s.d., Supp. Tables T1-T2), These differences correspond to an approximately 20-60% reduction in deuterium incorporation relative to CIM1 under comparable conditions and exceed experimental uncertainty in all three regions. Together, these results indicate stronger CNOT1 engagement by N-CIM1 and show that the functional CIM1 element extends beyond the previously defined sequence. However, none of the GW182 SD-derived fragments tested bound as strongly as TTP.

### Distinct binding modes of CIM1 and TTP within the shared groove

Despite engaging the same hydrophobic groove, the HDX patterns revealed subtle ligand-specific differences that argue against an identical binding mode (Fig. 1f, S2-S5). In particular, the protection effect was stronger at the IDDEANSY(839-846) peptide relative to the residues 847-859 for N-CNOT1 than that for TTP. Accordingly, the coarse-grained docking suggests that the N-CIM1 chain has the same N-to-C orientation as TTP in the complex^7^, but its alignment with the binding groove is less parallel than that of TTP (Fig. S1 b).

To further probe the overlap of the TTP-and CIM1-binding sites and their binding mode differences, we introduced E893A/Y900A point mutations into CNOT1 (referred to as CNOT1 EY), which disrupt key interactions with TTP^7^. HDX-MS confirmed that the mutations do not alter the wild-type (wt) HEAT repeat structure of CNOT1^67^. As expected, these mutations abolished protection induced by TTP across the hydrophobic groove (Fig. 1f, right panel). In contrast, CIM1-induced protection was only partially lost: while the protection in the regions 839-846 and 894-909 was eliminated, residual protection persisted in the residues 847-859. This indicates that CIM1 binding is weakened but not completely abolished, consistent with a distinct engagement of the same hydrophobic groove. Thus, while both ligands rely on the same core residues for docking, CIM1 retains alternative stabilizing contacts that partially compensate for loss of the canonical TTP interaction hotspot.

Coarse-grained docking of N-CIM1 to CNOT1 using CABS-Dock^68^ yielded models consistent with these observations (average RSMD 3.8 Å, Fig. S1b). In the best-scoring model, the aromatic side chain of GW182 Trp1300 occupies the same position as TTP Phe319 in the crystal structure^7^, while the peptide backbone adopts a slightly altered orientation relative to the groove (Fig. S1b).

### GW182 CIM1 binds CNOT1 with SLiM-like affinity

To quantify the interaction, we performed HDX-MS titrations of CNOT1 with increasing concentrations of N-CIM1 (Fig. 2a, S7). Analysis of mass shifts allowed estimation of apparent local dissociation constants (*K_d_^local^*) for individual CNOT1 peptides. The strongest effects, engaging ∼40 to 80% of the exchangeable amide hydrogens, were observed for peptides spanning residues 839-847, 847-859, and 894-909, with *K_d_^local^* values in the low-to-mid micromolar range (Suppl. Table T6), consistent with transient interactions mediated by short linear motifs (SLiMs)^69,70^. Protection of residues 800-819 exhibited the lowest apparent *K_d_^local^* but involved only a small fraction of exchangeable amides, indicating an auxiliary contribution.

**Figure 2.**
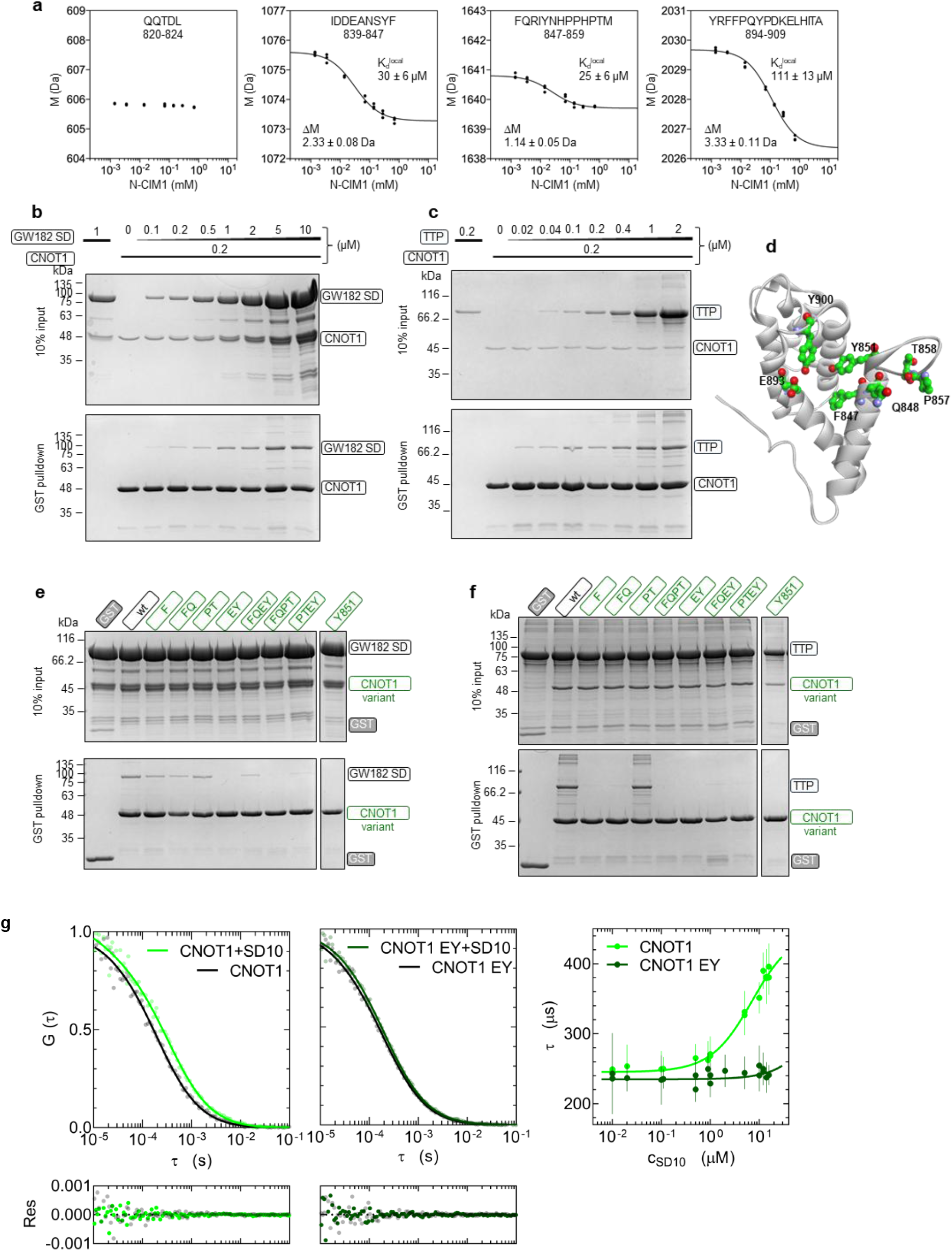
GW182 SD interacts with the tristetraprolin-binding site of CNOT1 with a micromolar affinity. **a**, Binding isotherms obtained as changes in mass (M in Da) of selected CNOT1 peptides after 2 hours of HDX in the presence of increasing concentrations of the N-CIM1 peptide, reflecting the local involvement in N-CIM1 binding; *K_d_^local^*, apparent dissociation constant for N-CIM1 binding to the CNOT1 peptide fragment; *ΔM*, maximal protection against HDX of the CNOT1 peptide in the complex with N-CIM1. CNOT1 concentration was 2 µM. **b,** GST pulldown assay for GST-CNOT1(800-999) titration with MBP-GW182 SD (see also Fig. S9) or, **c**, with MBP-TTP. **d**, Selected point mutations around the TTP-binding site shown in the crystal structure of CNOT1(800-999) (pdb 4J8S^7^). See Fig. S8 for all mutations tested. **e**, Results of GST pulldown assays for binding of MBP-GW182 SD (see also Fig. S9) or, **f**, MBP-TTP to GST-CNOT1(800-999) with the point mutations. **g**, Normalized FCS autocorrelation curve for fluorescently labeled (Cy5) GST-CNOT1(800-999) at 1 μM (left panel) and its E893A/Y900A mutant (middle panel) in the absence (black line) and presence of 15.6 μM GW182 SD10 (green line) fitted to experimental data (translucent points), together with their raw fitting residuals (bottom panels). (Right panel) quantitative determination of the equilibrium dissociation constant, *K_d_*, for GW182 SD10 binding to CNOT1(800-999) (light green) and CNOT1(800-999) E893A/Y900A (dark green) by FCS titration. The diffusion times, *τ*, of GST-CNOT1 become longer at increasing concentrations of GW182 SD10 due to their complex formation. The slight increase in τ observed for CNOT1 EY is indicative of residual interactions with GW182 SD10. For negative control with use of α chymotrypsinogen, see Fig. S10.

Pulldown titrations using GST-CNOT1 and MBP-tagged GW182 SD (residues 1260-1690) showed that several-fold higher concentrations of GW182 SD are required to achieve retention comparable to that of full-length TTP (Fig. 2b, c), whose affinity for CNOT1 is ∼2 μM^7^. These data suggest an overall affinity for GW182 SD in the ∼10 μM range, consistent with the upper limiting *K_d_^local^* values for N-CIM1 (Fig. 2a). Such micromolar affinities are characteristic of SLiMs^69,70^ operating in dynamic multiprotein assemblies, where reversibility rather than stability is required for regulatory switching.

Fluorescence correlation spectroscopy (FCS) measurements of the diffusion of fluorescently labeled GST-CNOT1 directly quantified its interaction with GW182 SD10 (Fig. 2g). Increasing concentrations of GW182 SD10 induced a twofold increase in the CNOT1 diffusion time, yielding a dissociation constant of 6.6 ± 2.1 μM and a corresponding ΔG⁰ of - 29.0 ± 0.8 kJ/mol. This affinity is weaker than that of TTP^7^, with a ΔΔG⁰ of ∼3 kJ/mol, consistent with the energetic contribution of a single non-covalent contact and supporting the notion of a shared site with slightly distinct binding modes.

Mutational analysis using GST-CNOT1 variants further corroborated these conclusions (Fig. 2e, f, S8). The E893A/Y900A or Y851A mutations abolished binding to both TTP and GW182 SD under pulldown conditions, whereas mutations affecting Phe847A/Q848A or Pro857A/Thr858A (Fig. 2d) differentially impaired interactions with TTP or GW182 SD, respectively. FCS titrations using the CNOT1 EY mutant showed only minimal changes in diffusion time, indicating severely reduced affinity to GW182 SD10 (Fig. 2g; negative control FCS experiments with α-chymotrypsinogen are shown in Fig. S10).

Collectively, HDX-MS, pulldown, and FCS titration analyses demonstrate that the N-CIM1 peptide of GW182 SD binds the same hydrophobic groove of CNOT1 as TTP, albeit through a partially distinct set of contacts (Fig. 2d, S1b).

### GW182 SD and TTP compete for CNOT1 binding *in vitro*

Given the shared interacting site, direct *in vitro* competition was further examined using FCS-based displacement assays with rhodamine B-labeled CIM1 peptide (RhB-CIM1_c_) at 20 nM (Fig. 3a, S12). Binding of GST-CNOT1 increased the diffusion time of RhB-CIM1_c_ by ∼2.5-fold, whereas GST-CNOT1 EY induced only a minor shift. Addition of excess TTP peptide restored fast diffusion of RhB-CIM1_c_, indicating displacement of CIM1_c_ from the CNOT1 complex.

**Figure 3.**
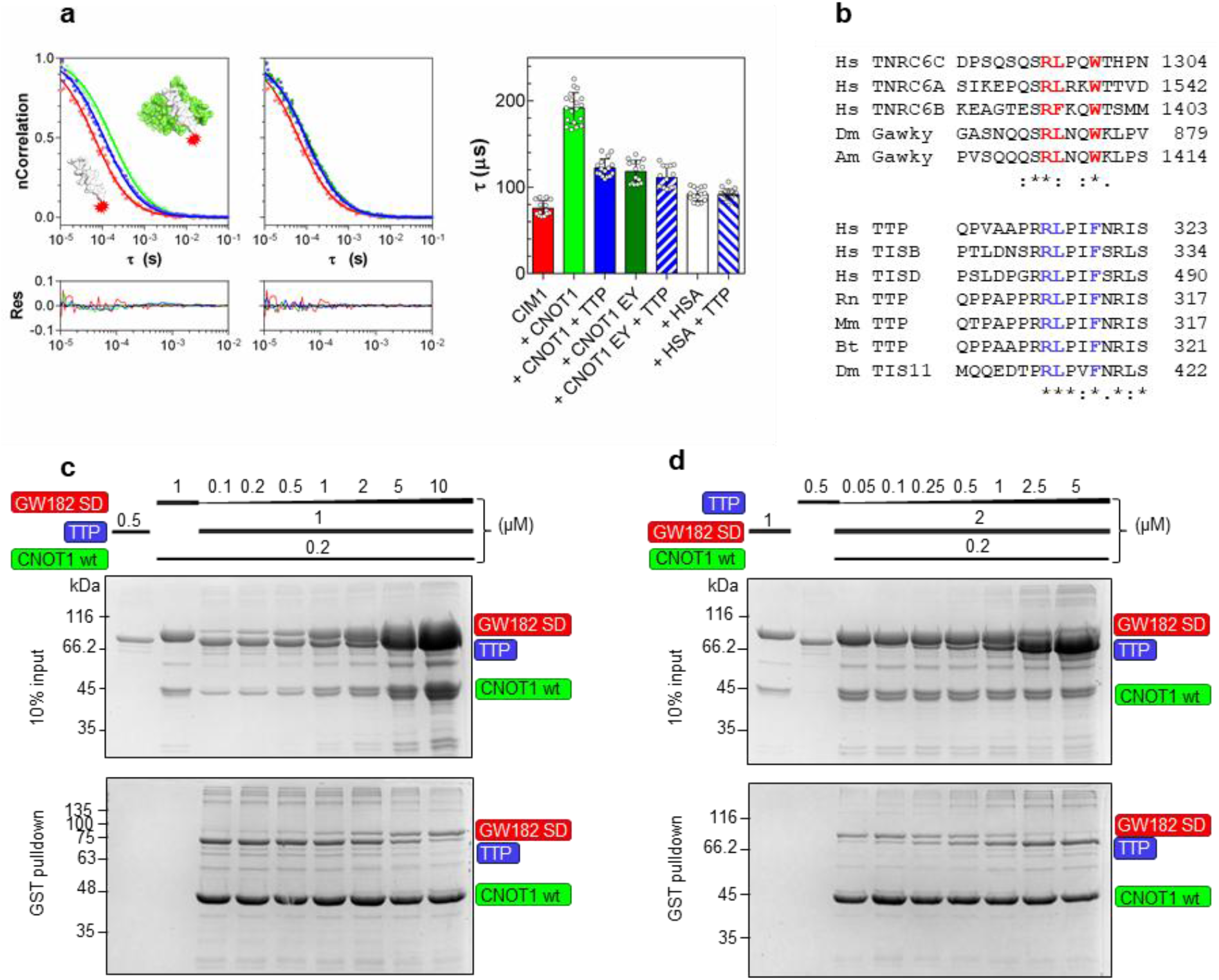
GW182 SD CIM1 directly competes with TTP for the same binding site on CNOT1 through a convergent linear motif. **a**, (Left panel) Normalized FCS autocorrelation curve for diffusion of fluorescently labeled (RhoB) CIM1_c_ peptide alone (at 20 nM, red), in the presence of GST-CNOT1(800-999) (left panel) or GST-CNOT1(800-999) E893A/Y900A (middle panel) at 10 μM (light and dark green, respectively), and after addition of the TTP peptide at a final concentration of 50 μM as the third component (blue). The diffusion time of CIM1_c_ increases significantly upon the complex formation with CNOT1 (from red to green line), and then decreases (from green to blue line) due to the displacement of CIM1_c_ from the complex by TTP. (Middle panel) Influence of CNOT1 EY on CIM1_c_ diffusion is less pronounced and unperturbed by TTP, pointing to nonspecific interactions. (Right panel) Quantitative changes in CIM1_c_ diffusion time in the binary (with CNOT1, CNOT EY, or HSA as a negative control) and ternary (in the presence of TTP) mixtures. **b**, Sequence similarity of GW182 SD CIM1 and CNOT1-binding fragment of TTP within the convergent RL(P/ζ)XΩ motif, where: ζ, hydrophilic; X, any; and Ω, aromatic residue. Hs, human (*Homo sapiens*); Dm, fruit fly (*Drosophila melanogaster*); Am, honeybee (*Apis mellifica*); Rn, rat (*Rattus norvegicus*); Mm, mouse (*Mus musculus*); Bt, cattle (*Bos taurus*). **c**, GST pull-down assay for the displacement of MBP-TTP by MBP-GW182 SD from the complex with GST-CNOT1(800-999). **d**, GST pull-down assay for the displacement of MBP-GW182 SD by MBP-TTP from the complex with GST-CNOT1(800-999).

We next examined whether the GW182 SD and TTP proteins compete for CNOT1 binding. GST-CNOT1 pulldown assays performed with constant MBP-TTP and increasing MBP-GW182 SD concentrations revealed a progressive loss of TTP retention (Fig. 3c). Conversely, increasing MBP-TTP concentration reduced the binding of MBP-GW182 SD to CNOT1 (Fig. 3d), demonstrating reciprocal competition.

These results provide the evidence that TTP directly competes with the CIM1 of GW182 for CNOT1 binding.

### Convergent short linear motifs in GW182 and TTP

Sequence analysis revealed an unexpected convergence between the CNOT1-binding motifs of GW182 and TTP (Fig. 3b). Although both proteins are evolutionarily unrelated, they contain a conserved five-residue motif conforming to RL(P/ζ)XΩ (where ζ is hydrophilic, Ω is aromatic, and X is any residue), represented by RLPQW in TNRC6 C and RPLIF in TTP. The key aromatic residues (Trp1300 in GW182, Phe319 in TTP) occupy equivalent positions and are critical for CNOT1 binding^7,26^.

The regions flanking the motif differ markedly (Fig. 3b). In GW182s, N-terminal extensions are enriched in polar residues and potential phosphorylation sites, whereas TTP flanking sequences are more rigid and hydrophobic. These differences likely underlie the distinct binding modes and regulatory properties of the two interactions.

### GW182 silencing domain undergoes liquid-liquid phase separation

We next examined whether the GW182 SD can undergo liquid-liquid phase separation. Purified GW182 SD formed liquid droplets at room temperature at micromolar concentrations (Fig. 4a). Fusion of GW182 SD to mCherry slightly increased solubility but preserved LLPS behavior, which was reproducibly observed in the presence of low concentrations of PEG 1000 (Fig. 4b-f). Droplets formed at 6 μM GW182 SD were highly dynamic, exhibiting rapid coalescence and fluorescence recovery after photobleaching (FRAP) with half-times of 10-20 s (Fig. 4c-f). The mobile fraction decreased with increasing PEG concentration (Fig. 4d), consistent with crowding effects reported for other phase-separating proteins^71^.

**Figure 4.**
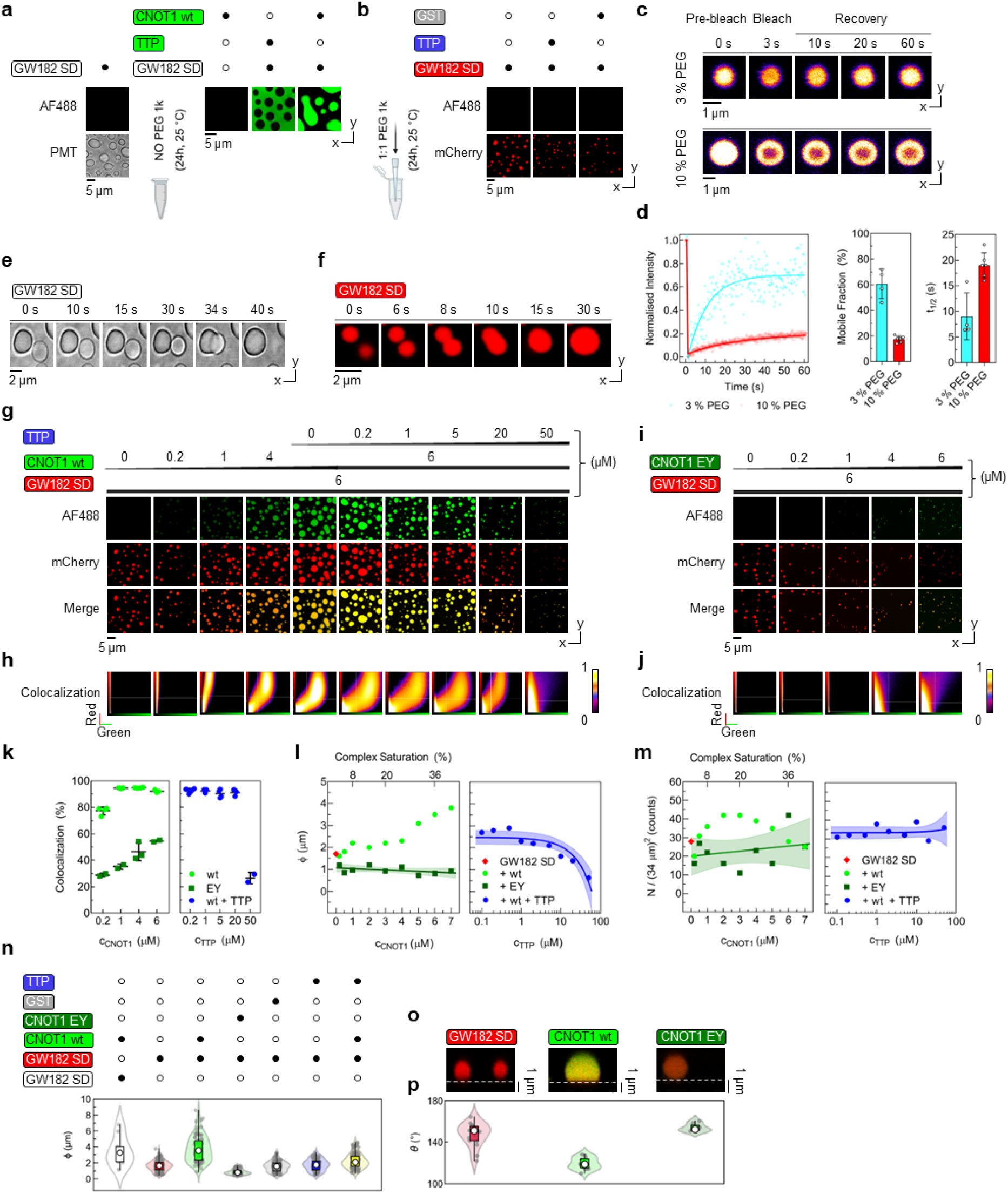
GW182 SD triggers liquid-liquid phase separation and recruits CNOT1(800-999) to multiprotein droplets *in vitro*. **a**, Representative confocal images showing liquid droplets of GW182 SD at 6 µM in transmitted light (left panel) and incorporation of fluorescently labeled (AF488) GST-CNOT1(800-999) into GW182 SD droplets (right panel, equimolar concentrations of 6 µM). CNOT1 alone does not undergo LLPS. **b**, Representative confocal images of LLPS of GW182 SD-mCherry (6 µM) in 10% (w/v) PEG 1k in the presence of GST-AF488 (6 µM) or TTP peptide (50 µM) as controls. **c**, FRAP time-lapse images for GW182 SD-mCherry droplets formed in 3% and 10% PEG 1k. **d**, Quantification of mobile fraction and fluorescence recovery half-time (*t₁_/_₂*) for GW182 SD-mCherry droplets in 3% (cyan) and 10% (red) PEG 1k, n = 2-5. **e**, Time-lapse confocal images showing coalescence of droplets formed by GW182 SD (6 µM) without PEG and, **f**, GW182 SD-mCherry (6 µM) in 3% (w/v) PEG 1k. **g**, Representative images of multiprotein droplets formed by GW182 SD-mCherry incorporating GST-CNOT1(800-999)-AF488 at increasing concentrations from 0.2-6 µM, and further, at increasing concentrations of TTP peptide (0.2-50 µM) in the ternary mixture. **h**, Colocalization diagrams showing fluorescence intensity correlation between mCherry and AF488 channels in droplets formed by GW182 SD-mCherry with CNOT1 or with CNOT1 and TTP peptide. The upward curvature of the correlation is due to FRET (explained in Fig. 5). **i**, Multiprotein droplets of GW182 SD-mCherry with GST-CNOT1(800-999)-AF488 E893A/Y900A at 0.2-6 µM. **j**, Colocalization diagrams showing fluorescence intensity correlation between mCherry and AF488 channels in droplets formed by GW182 SD-mCherry with CNOT1 EY. **k**, Quantitative dependence of colocalization (value calculated for the lower left corner of the diagram) in multiprotein droplets on the concentration of CNOT1 (light green circles) or CNOT1 EY (green squares), and on TTP peptide concentration at 6 µM CNOT1 (blue circles). **l**, Quantitative dependence of the median droplet diameter (φ) and, **m**, number of droplets (N) on CNOT1 (light green circles) or CNOT1 EY (green squares) concentration, and on TTP peptide concentration at 6 µM CNOT1 (blue circles). **n**, Violin plots showing diameter distribution of droplets formed by GW182 SD at 6 µM with CNOT1 at 6 µM (transparent), GW182 SD-mCherry alone (6 µM, red), and GW182 SD-mCherry (6 µM) in the presence of CNOT1 (6 µM, light green), CNOT1 EY (6 µM, dark green), GST (6 µM, grey), TTP peptide (5 µM, blue), and CNOT1 at 6 µM with TTP at 5 µM (yellow); open circles, medians; boxes, interquartile ranges (Q1-Q3); whiskers, range; grey dots, individual droplets. **o**, Confocal Z-stack images showing droplets formed by GW182 SD-mCherry (6 µM) alone (left), with CNOT1 at 5 µM (middle), and with CNOT1 EY at 5 µM (right). **p**, Violin plots showing contact angle distribution of droplets formed by GW182 SD-mCherry alone (red), with CNOT1 (light green), or with CNOT1 EY (dark green). Images acquired after 24 h of incubation; number of images, n = 2-5 at each condition. See Fig. S13-S18 for more data and statistics after 12 h of incubation.

### CNOT1 is recruited as a client into GW182 SD condensates

While GW182 SD undergoes LLPS on its own (Fig. 4a,b), CNOT1(800-999) does not form condensates under the same conditions (Fig. 4a). The TTP peptide was also excluded from GW182 SD condensates (Fig. 4a, S13), indicating that it does not contribute to condensate formation through independent phase separation or partitioning. However, when CNOT1 was mixed with GW182 SD at equimolar concentrations, they formed multiprotein liquid droplets (Fig. 4a), indicating that GW182 SD functions as a scaffold, recruiting CNOT1 as a client into the condensates^52^. This motivated us to use the scaffold-client relationships as a functional assay to examine the molecular basis of GW182/TTP binding to CNOT1.

Titration of AF488-CNOT1 into 6 μM GW182 SD-mCherry revealed concentration-dependent recruitment into the dense phase (Fig. 4g, h, S14, S16), with near-complete colocalization achieved in the lower left quadrants at ≥0.2 μM CNOT1 (Fig. 4k). Incorporation of CNOT1 promoted droplet nucleation, growth, and coalescence (Fig. 4l, m), leading to the increase in the median droplet diameter and in droplet number (Fig. 4l, m) up to a client protein concentration at which the distances between droplets became comparable to their diameter, as quantified by Delaunay triangulation (Fig. S14, S16). At this point, coalescence began to dominate, resulting in a stronger increase in diameter and a decrease in the number of droplets. Notably, CNOT1 recruitment also altered the surface wetting behavior of the droplets, with increased spreading on hydrophobic surfaces, as reflected by a reduced contact angle (Fig. 4o,p). This change in interfacial behavior suggests that incorporation of CNOT1 as a client tunes condensate physicochemical properties. These observations indicate that CNOT1 promotes condensate assembly and growth through specific heterotypic interactions, rather than acting as an additional independent phase-separating scaffold.

In contrast, the CNOT1 EY mutant showed markedly reduced recruitment into GW182 SD condensates (Fig. 4i,j, S18), as quantified by the degree of colocalization (Fig. 4k), and impaired droplet growth without affecting nucleation (Fig. 4l, m). The EY mutant also failed to alter droplet wetting (Fig. 4o, p), indicating a distinct mode of scaffold-client organization within the unspecific condensates.

### TTP inhibits GW182 SD-CNOT1 multiprotein condensation

To test whether TTP competes with GW182 SD for CNOT1 during condensate formation, we performed LLPS competition assays. The TTP peptide alone did not affect GW182 SD-mCherry phase separation (Fig. 4b, n) and did not partition into droplets (Fig. 4a). Addition of increasing concentrations of TTP to equimolar GW182 SD-mCherry and CNOT1 mixtures resulted in a pronounced reduction in multiprotein droplet size (Fig. 4g, l), with a narrowing of the size distribution toward that observed for GW182 SD alone (Fig. 4n). Droplet number remained largely unchanged (Fig. 4m), indicating that TTP primarily inhibits the client recruitment and droplet growth rather than the scaffold nucleation (Fig. S15, S17). At higher TTP concentrations (20-50 μM), LLPS was strongly suppressed, resembling the unspecific effect of the presence of the CNOT1 EY mutant (Fig. 4i).

This uncoupling of nucleation from droplet growth is consistent with TTP selectively limiting CNOT1-mediated network expansion rather than interfering with GW182 SD homotypic interactions, which further supports the possibility of competition between TTP and GW182 SD for CNOT1 binding.

### FRET reveals specific heterotypic interactions within condensates

Confocal imaging revealed enhanced red fluorescence under green excitation in GW182 SD-CNOT1 condensates in a form of an upward curvature in the colocalization diagrams (Fig. 4h), suggesting Förster resonance energy transfer (FRET) between AF488-labeled CNOT1 and GW182 SD-mCherry. Replacement of mCherry with Alexa Fluor 647 (at Cys378, Fig. S19) abolished this nonlinear signal increase (Fig. 5b-d, S20, S21), consistent with reduced spectral overlap and increased donor-acceptor distance (Fig. 5e).

**Figure 5.**
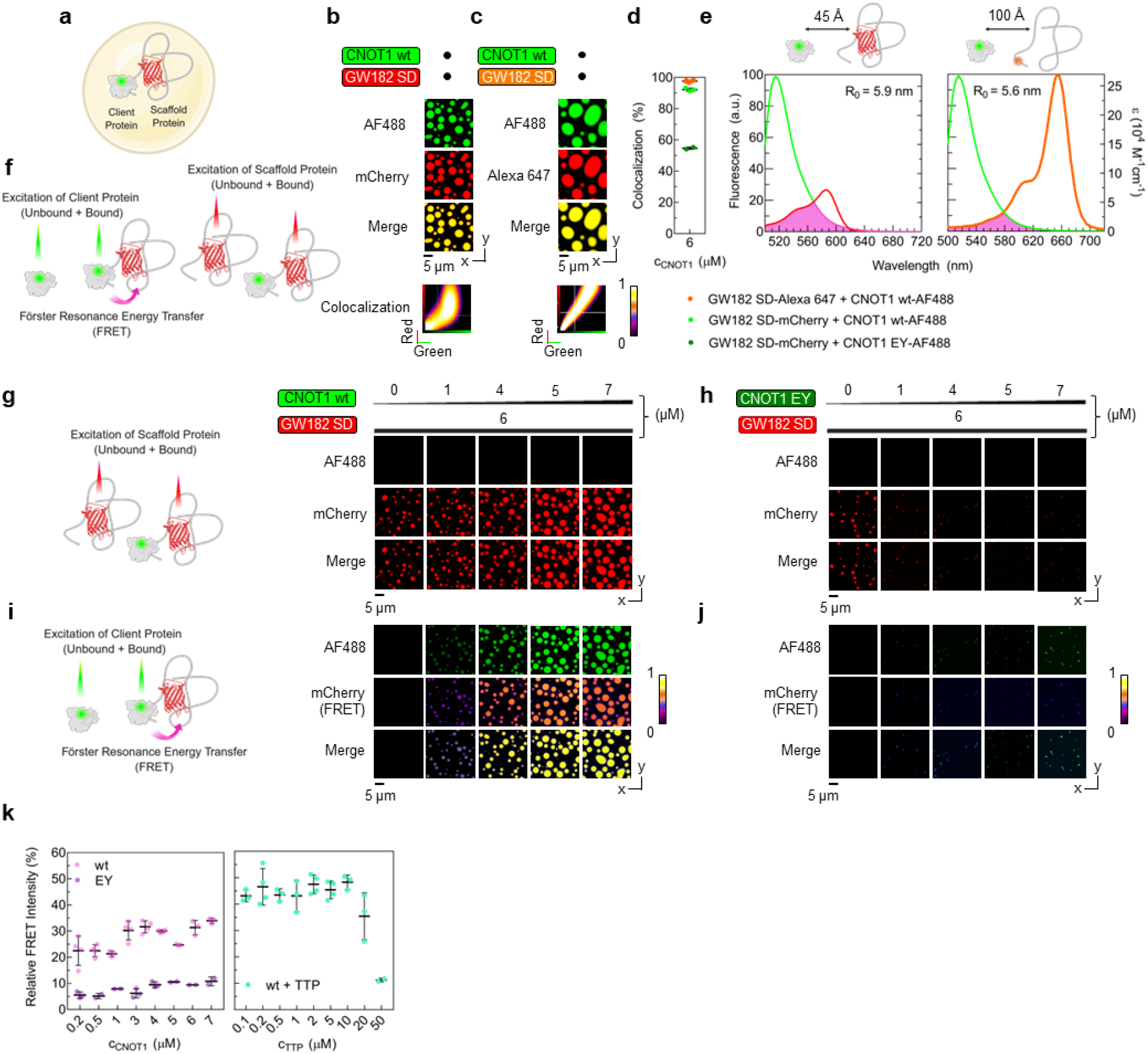
Intact TTP-binding site of CNOT1 is required for specific scaffold-client interactions with GW182 SD in multiprotein droplets. **a**, Schematic illustration of a multiprotein droplet composed of the scaffold GW182 SD-mCherry protein, which drives LLPS, and the client GST-CNOT1(800-999)-AF488 protein. **b**, Confocal images after double excitation (red and green) of droplets formed by GW182 SD-mCherry and CNOT1 (both at 6 µM), with colocalization diagram showing fluorescence intensity distributions of AF488 and mCherry. **c**, Confocal images after double excitation of droplets formed by GW182 SD-Alexa Fluor 647 and CNOT1, with colocalization diagram showing fluorescence intensity distributions of AF488 and Alexa Fluor 647. **d**, Quantification of colocalization levels in droplets formed by equimolar concentrations (6 µM) of GW182 SD-Alexa Fluor 647 with CNOT1 (orange points), and GW182 SD-mCherry with either CNOT1 (light green) or with CNOT1 EY (dark green). **e**, Fluorophores used in **b** and **c** as donor-acceptor FRET pairs, with the spectral overlap marked pink; normalized emission spectrum of AF488 (light green) and absorption spectrum of mCherry (left, red) or Alexa Fluor 647 (right, orange) with corresponding Fӧrster radii. **f**, Schematic illustration of double-excitation confocal imaging experiments, enabling visualization of both client CNOT1 (green emission) and scaffold GW182 SD molecules (red emission). The signal from CNOT1 (donor) is attenuated while the GW182 SD signal (acceptor) is enhanced due to FRET from AF488, which is stronger with mCherry (**b**, colocalization panel, similarly as in Fig. 4h) than with Alexa Fluor 647 (**c**, colocalization panel). **g**, Single-excitation (red) confocal imaging of total population of GW182 SD-mCherry molecules, both unbound and bound, upon LLPS at increasing concentrations (1-7 µM) of CNOT1 or, **h**, CNOT1 EY. **i**, Single-excitation (green) confocal imaging of CNOT1 (AF488 channel) and the FRET-derived GW182 SD-mCherry signal coming selectively from a part of the CNOT1-bound fraction upon LLPS at increasing concentrations (1-7 µM) of CNOT1 or, **j**, CNOT1 EY. **k**, Quantitative dependence of relative FRET intensity from CNOT1-AF488 to GW182 SD-mCherry (at 6 µM) on CNOT1 (pink) or CNOT1 EY (purple) concentration, and on TTP peptide concentration at 6 µM CNOT1 (cyan). Images acquired after 24 h of incubation with 10% PEG; number of images, n = 2-5 at each condition. See Fig. S21-S27 for larger images, more data and statistics after 12 h of incubation.

Quantitative analysis revealed that FRET intensity increased with CNOT1 concentration (Fig. 5g, i, k, S22), whereas the EY mutant produced a markedly weaker response (Fig. 5h, j, k, S23). Addition of TTP reduced FRET in a concentration-dependent manner, with FRET levels approaching those of the EY mutant at higher TTP concentrations (Fig. 5k), demonstrating competitive displacement of CNOT1 from GW182 SD within condensates. Altogether, these findings suggest that CNOT1 recruitment into the dense phase is determined by the specificity and possible inhibition of its heterotypic interactions with GW182 SD.

### FRET-FRAP distinguishes specific from nonspecific client recruitment

To further probe the scaffold-client interaction specificity, we employed a combined FRET-FRAP approach^72^ (Fig. 6a). In this assay, photobleaching of the donor abolished FRET, and the recovery was monitored in both donor and FRET channels. For CNOT1, mobile fractions in the donor (CNOT1) and acceptor (GW182 SD) channels were nearly identical (Fig. 6b, d), indicating stable heterotypic complexes that co-diffused within the dense phase, with the same mobile fraction as GW182 SD-mCherry alone (Fig. 4c, d). In contrast, CNOT1 EY exhibited a higher donor mobile fraction than that observed through FRET for the complex (Fig. 6c, e), consistent with weaker, nonspecific or transient interactions that led to residual binding. These measurements distinguish stable, sequence-specific heterotypic interactions from nonspecific client enrichment and demonstrate that CNOT1 recruitment into GW182 SD condensates is governed by defined molecular recognition rather than generic IDP stickiness.

**Figure 6.**
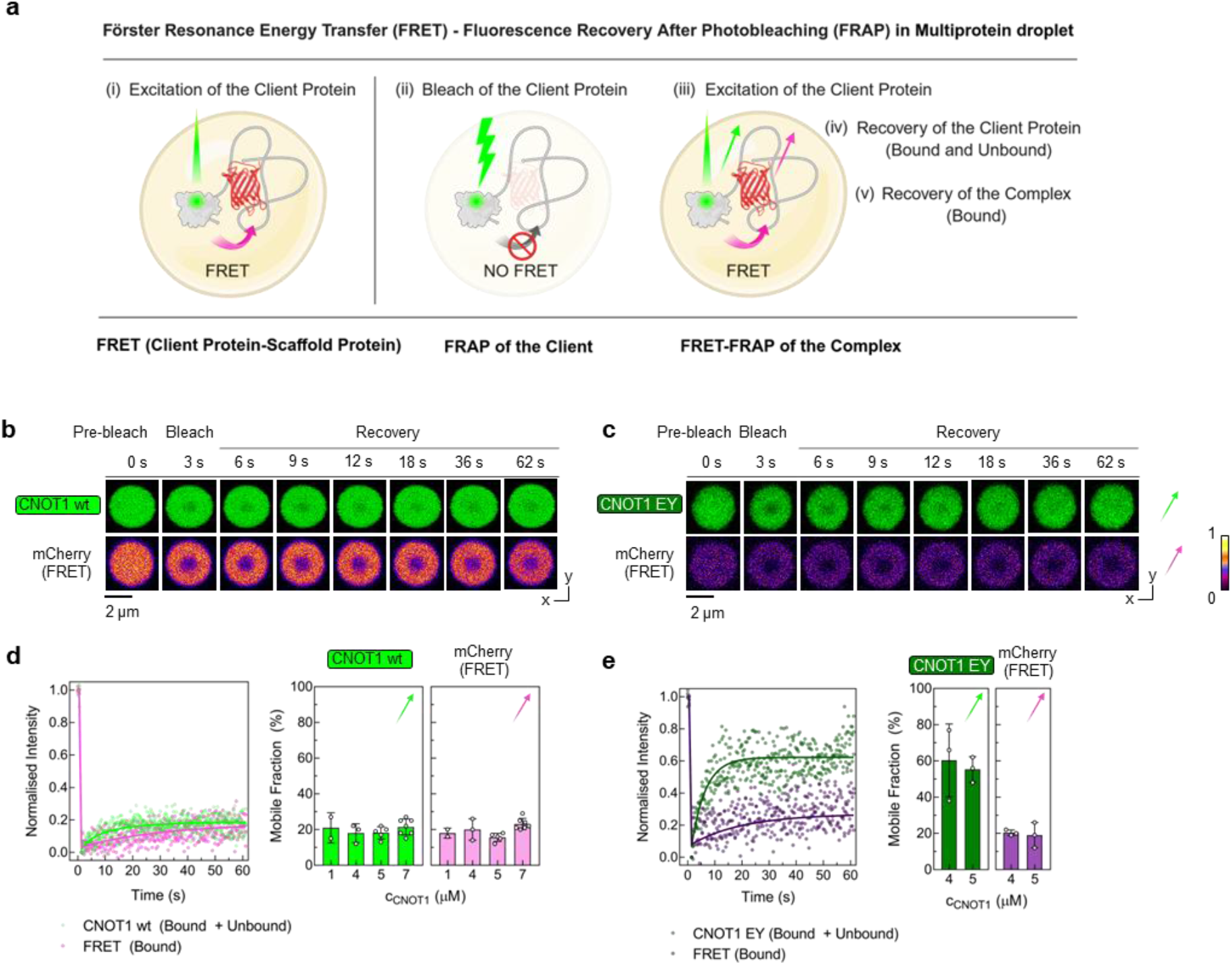
CNOT1 client mobility is governed by heterotypic interactions with GW182 SD scaffold. **a**, Schematic illustration of the combined Förster resonance energy transfer and fluorescence recovery after photobleaching (FRET-FRAP) assay. After selectively bleaching the donor (client CNOT1), FRAP is observed separately for the donor (unbound and a fraction of bound that does not undergo FRET) and the acceptor (scaffold GW182 SD-mCherry, only a fraction of bound that is excited *via* FRET). Green and pink arrows indicate fluorescence recovery. **b**, FRET-FRAP time-lapse images of droplets formed by GW182 SD-mCherry (6 µM) and CNOT1 (5 µM) or, **c**, CNOT1 EY (5 µM), showing fluorescence recovery for CNOT1 directly in the AF488 channel and for GW182 SD-mCherry *via* FRET in the mCherry channel. For FRET-FRAP comparison between CNOT1 wt and EY, droplets of the same size were selected. **d**, (left) FRET-FRAP kinetics in multiprotein droplets of GW182 SD-mCherry (6 µM) and CNOT1 (5 µM), CNOT1 (light green) observed directly, GW182 SD (pink) observed *via* FRET, n = 5; (right) quantification of mean mobile fractions for CNOT1 (light green) and the FRET-excited bound fraction of GW182 SD (pink) at increasing concentrations of CNOT1 (1-7 µM), n = 2-7. **e**, (left) FRET-FRAP kinetics in multiprotein droplets of GW182 SD-mCherry (6 µM) and CNOT1 EY (5 µM), CNOT1 EY (dark green) observed directly, GW182 SD (purple) observed *via* FRET, n = 3; (right) quantification of mean mobile fractions for CNOT1 EY (dark green) and the FRET-excited bound fraction of GW182 SD (purple) at increasing concentrations of CNOT1 EY (4-5 µM), n = 3. Images acquired after 24 h of incubation with 10% PEG. See Fig. S28, S29 for more FRET-FRAP data and negative controls.

## Discussion

We identify a previously unrecognized mode of interaction between GW182 and the central CCR4-NOT subunit CNOT1 that necessitates a redefinition of the CCR4-NOT-interacting motif 1 (CIM1)^26^. Rather than constituting a minimal element, CIM1 functions as part of an extended sequence that engages the 800-999 region of CNOT1 through a low-affinity interaction. The N-terminally extended DPSQSQSRLPQWTHPN sequence, herein termed N-CIM1, strongly enhances the binding, establishing the extended N-CIM1 sequence as the functional unit mediating GW182 binding to CNOT1.

An additional point of convergence in distinct gene silencing pathways was detected, namely that GW182 SD and TTP compete for a partially shared binding site on CNOT1(800-999). This site is located within a hydrophobic groove formed by helices α1, α3, and α4 and encompasses the critical residues E893 and Y900.^7^ The overlap is explained by a local sequence similarity, as both proteins contain a SLiM, RL(P/ζ)XΩ, despite being evolutionarily unrelated. In both cases, the conserved Trp or Phe side chains dock in equivalent positions within the CNOT1 groove (Fig. S1b, 7). The broader RLXXΩ motif is conserved across GW182 and TTP paralogs and species^35,36^ (Fig. 3b), supporting its classification as a SLiM that likely arose through convergent evolution. A comparable mechanism is observed in the regulation of translation initiation, where unrelated proteins employ similar motifs to engage a shared site^38,73–75^.

SLiMs frequently act as recruitment interfaces within large multiprotein assemblies, mediating weak and transient interactions that enable rapid regulatory switching^37^. In this context, the shared RLPXF/W motif of GW182 SD and TTP recognized by CNOT1(800-999) exhibits micromolar affinities, consistent with dynamic regulation rather than stable complex formation. These interactions modulate CCR4-NOT engagement by distinct silencing pathways, establishing CNOT1 as a regulatory hub that coordinates miRNA-and ARE-mediated gene silencing across diverse physiological contexts.

GW182 and TTP engage CCR4-NOT through multivalent contacts. GW182 SD can associate with CCR4-NOT indirectly *via* CNOT9 through the CN9BD region of CNOT1^14^. However, CNOT1 variants incapable of binding CNOT9 retain interaction with GW182^15^, implying the existence of independent binding sites^26^. Direct engagement of GW182 CIM1 with the TTP-binding region of CNOT1 in the absence of CNOT9 demonstrates that both CNOT9-and CNOT1-dependent interactions can contribute to the complex formation. Similarly, TTP engages CCR4-NOT through both CNOT1 and CNOT9^73^, utilizing the same tryptophan-binding pockets involved in GW182 SD recognition, thereby stabilizing CCR4-NOT association and promoting efficient deadenylation of target mRNAs^76^. This multivalent view is consistent with recent reconstitution studies showing that IDRs of RNA-adaptors can use dispersed interaction segments to bind Ccr4–Not and tune deadenylation rates, with related evidence for human TTP^63^. In this context, the CNOT1(800–999) groove defined here represents a molecularly resolved competitive interface within a broader multivalent adaptor– CCR4–NOT interaction network. Together, these observations provide a shared mechanistic basis for potential crosstalk between miRNA-and ARE-dependent silencing pathways.

Beyond direct binding, the GW182 SD functions as a scaffold within multiprotein condensates, linking SLiM-mediated recognition to mesoscale organization. GW182 SD undergoes LLPS, forming dynamic assemblies that recruit the TTP-binding region of CNOT1 (Fig. 7). Incorporation of CNOT1 promotes condensate growth and alters droplet wetting behavior, indicating that heterotypic interactions modulate condensate physicochemical properties. Recent work on natural biomolecular condensates has shown that low interfacial tension facilitates exchange of molecules *via* the dilute phase, highlighting interfacial organization as a key determinant of condensate material properties^77^. More broadly, condensate organization is thought to emerge from the balance between homotypic scaffold interactions and selective heterotypic client contacts^52,79,74^. Additionally, *in silico* models suggest that the energetic barrier imposed by conformational entropy of intrinsically disordered regions, such as GW182 SD, can be overcome only through favorable binding interactions^80^.

**Figure 7.**
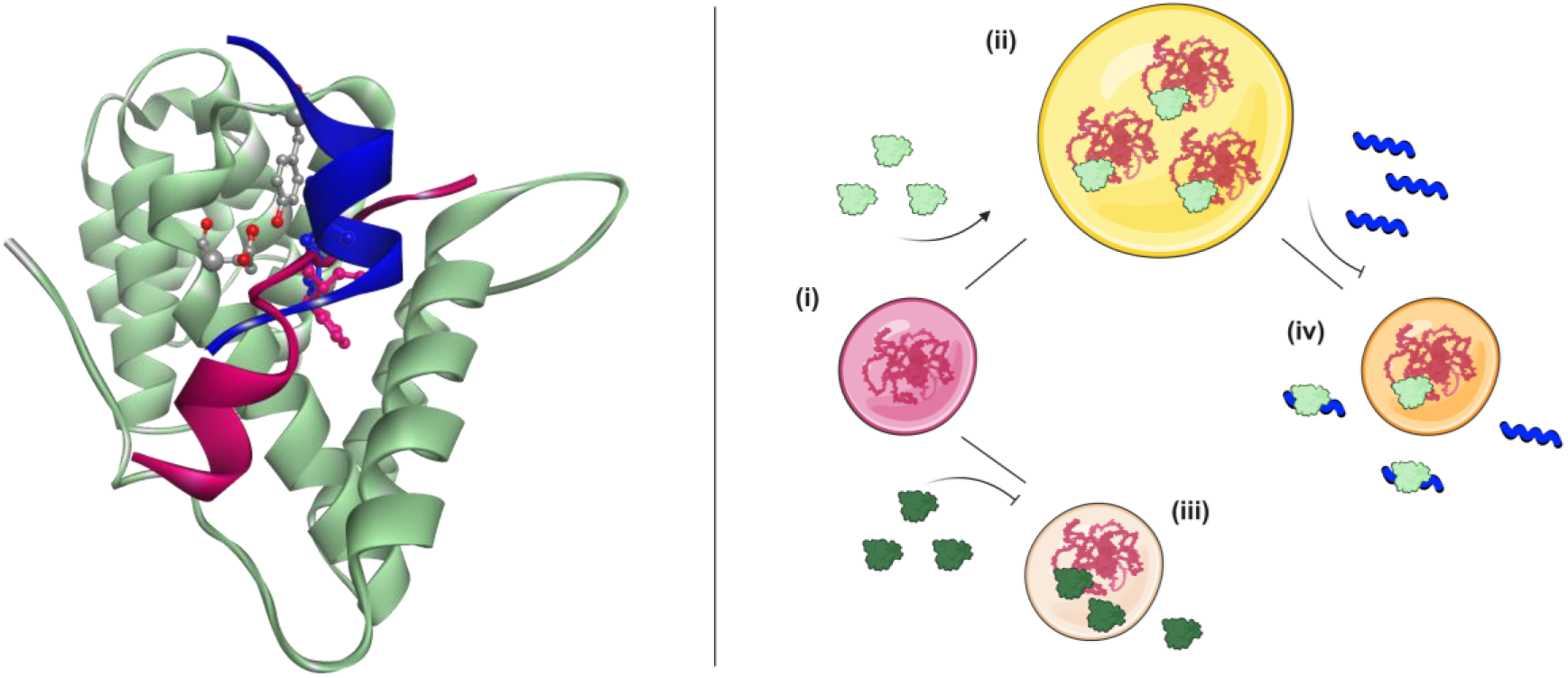
(Left) The N-terminal CNOT1 domain (residues 800-999; light green) accommodates the shared binding site for tristetraprolin peptide (TTP; blue) and CIM1 of GW182 SD (TNRC6C; magenta). The aromatic side chains of GW182 Trp1300 and TTP Phe319^7^, which occupy equivalent positions, are shown in ball-and-stick representation. The key CNOT1 residues Glu893 and Tyr900 are shown in CPK colors. (Right) GW182 SD undergoes liquid-liquid phase separation (i) and recruits CNOT1(800–999) into multiprotein droplets (ii). CNOT1 point mutations that impair CIM1 binding (E893A/900A; iii), or competitive binding of TTP to CNOT1 (iv), disrupt the assembly and growth of GW182 SD– CNOT1 condensates.

Although reconstituted GW182 SD–CNOT1 droplets do not recapitulate cellular P-bodies, studies of the EDC4–XRN1 axis show that altered decay-factor interactions can remodel P-body dynamics and shift miRNA-mediated silencing from mRNA decay toward translational repression^81,82^. In this context, our data provide a defined client–scaffold system in which specific CNOT1 recruitment reshapes condensate growth and interfacial behavior, linking molecular recognition to condensate-level organization *in vitro*.

In this framework, the role of the GW182 silencing domain in multiprotein condensate assembly complements that of the N-terminal GW182 Ago-binding domain, which recruits Argonaute proteins as clients^54^. Together, these domains provide a modular architecture through which GW182 can influence CCR4-NOT-dependent deadenylation kinetics by promoting PB condensation^54,83^. GW182 thus emerges as both an adaptor for targeted CCR4-NOT recruitment^31^ and an active organizer of the dynamic miRISC machinery *via* phase separation.

The importance of specific CNOT1 residues for condensate organization further underscores the role of defined heterotypic contacts in GW182 SD-driven phase separation. Residues E893 and Y900 of CNOT1 are critical for both GW182 SD binding and selective recruitment into condensates. Mutation of these residues impairs the heterotypic interaction network required for condensation and secondarily weakens GW182 SD homotypic interactions, resulting in reduced condensate size and altered internal dynamics. These observations suggest that a client with impaired heterotypic interactions can diminish the inherent phase-separation capacity of a scaffold.

Similar sensitivity to single-residue perturbations has been observed in stress granule regulation in living cells^84^. Limited partitioning of the CNOT1 EY mutant into condensates reflects residual weak interactions, consistent with reports that nonspecific amino-acid “stickiness” can drive partial client enrichment without supporting functional phase separation^85^.

TTP modulates GW182 SD-CNOT1 condensation by limiting the pool of CNOT1 available for engagement. TTP itself remains excluded from GW182 SD droplets and does not act as a client. Instead, formation of the CNOT1-TTP complex interferes with the homo-and heterotypic interaction network required for condensate expansion, selectively suppressing droplet growth. This behavior aligns with principles showing that client recruitment and condensate stability depend on the strength, valency, and network compatibility of heterotypic interactions^86^, and underscores the role of SLiMs as tunable regulators of IDP-driven condensation that encode client-scaffold specificity^87^.

Taken together, the findings support a model in which the CNOT1(800-999) domain acts as a hub for SLiM-mediated interactions with either GW182 SD or TTP, linking miRNA-and ARE-dependent silencing pathways. In this model, GW182 SD can regulate CCR4-NOT engagement through a dual-mode mechanism that integrates CNOT1 binding in the dilute phase with modulation of reconstituted multiprotein condensate assembly and organization *in vitro*.

## Methods

### DNA constructs

Experiments were performed on human CNOT1(800-999) (UniProtKB: A5YKK6, isoform 2), GW182 fragments (TNRC6C paralog, UniProtKB: Q9HCJ0, isoform 1) denoted to as GW182 SD10 (residues 1260-1620)^27^ or GW182 SD (residues1260-1690), and tristetraprolin (UniProtKB: P26651) expressed from a pET28b(+) vector with a N-terminal His_6_-Sumo tag^88^; on tristetraprolin and GW182 SD expressed from a pMAL-c5x vector with an N-terminal MBP-tag and a C-terminal His-tag; and on CNOT1(800-999) cloned into the pGEX 6p1 vector with a GST-tag using *EcoRI* and *NotI* restriction sites. Point mutations in GST-CNOT1(800-999) were introduced using a standard site-directed mutagenesis protocol.

### Protein preparation

His_6_-Sumo proteins were overexpressed and purified as described previously^67^. MBP-GW182 SD/TTP-His were purified on HIS-Select Nickel Affinity Gel (Sigma-Aldrich) in 50 mM Tris/HCl, 150 mM NaCl, 5 mM imidazole, 2 mM DTT, pH 8.0. After purification, MBP-GW182 SD contained a mixture of the full length protein and a degradation product, which was identified by mass spectrometry to be GW182 SD without the MBP-tag. GST-CNOT1(800-999) wt or mutant were purified on Glutathione Sepharose 4B (GE Healthcare) in 50 mM Tris/HCl, 150 mM NaCl, 2 mM DTT, pH 8.0. All proteins were dialyzed into 50 mM Tris, 150 mM NaCl, 2 mM DTT, 1 mM EDTA, pH 7.0, and concentrated, if needed, on the day of the experiment using Amicon Ultra Centrifugal Filters (Merck Millipore) at 4 °C.

Proteins were labeled with either AF488 NHS ester (Lumiprobe) that interacts with primary amines (lysine residues and N-terminus) or Cy5, Alexa Fluor 647 C_2_ maleimide (Thermo Fisher Scientific) that interacts with cysteine residues, following the manufacturers’ protocols. Excess unreacted dye was removed by either multi-step dialysis using Purcysteine residues, following the manufacturers’ protocols. Excess unreacted dye was removed by either multi-Acysteine residues, following the manufacturers’ protocols. Excess unreacted dye was removed by either multi-Lyzers (Sigma-Aldrich) or size-exclusion chromatography (SEC) on a Superdex 200 Increase 10/300 GL column (Cytiva).

### Peptides

The TTP peptide (referred to as TTP; sequence: APRRLPIFNRISVSE), CIM1 (QSRLPQWTHP), N-CIM1 (DPSQSQSRLPQWTHPN), and CIM2 (GSSELLWGGVPQYSSSLWGPPS) were ordered from Genemed Synthesis (98% purity; San Antonio, Texas, USA). The peptide concentration was determined (± 5%) by quantitative amino acid analysis performed by BioCentrum (Kraków, Poland). Aliquots of the peptides were dissolved in 50 mM Tris/HCl pH 7.0, 150 mM NaCl for further studies.

The fluorescently labeled peptides, RhB-CIM1_c_ (rhodamine B-APSQSQSRLPQWTHPNSMD-NH_2_; GW182 TNRC6C sequence), and FITC-TTP (fluorescein-QPVAAPRRLPIFNRISVSE-NH_2_), were synthesized with a β-alanine linker separating the dye from the peptide sequence, HPLC-purified (95-97%), and MS-confirmed by Lipopharm (Gdańsk, Poland). In the TNRC6C peptide sequence, a change from D to A was necessary to avoid the acid hydrolysis of the D-P bond, which caused cleavage of the rhodamine fragment. Aliquots of the peptides were dissolved in 50 mM Tris/HCl pH 7.0, 150 mM NaCl, 0.5 mM EDTA, 1 mM TCEP for further studies. The concentrations of fluorescently labeled peptides were determined from the absorbance of RhB and FITC at their maxima.

### Hydrogen-deuterium exchange mass spectrometry measurements (HDX-MS)

HDX-MS measurements, including controls for in-exchange and back-exchange, for individual pepsin-generated peptides of CNOT1(800-999) upon interactions with N-CIM1, CIM1, TTP peptides, and GW182 SD10 were performed in Ion Mobility Mode and analyzed as previously described^67^ in three or four technical replicates. Individual data points on the HDX-MS kinetic plots are shown with error bars to visualize the MS experimental uncertainty. The fraction of amide hydrogens that were exchanged for deuterium, *ΔFr*, for each CNOT1 peptide in the presence of TTP, CIM1, or N-CIM1 was calculated as:

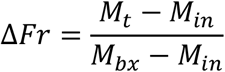

where *M_t_* was the average mass of the peptide after a given exchange time, *t*; *M_in_* and *M_bx_* were the average masses of the peptide in the in-exchange and back-exchange control samples^67^, respectively. The number of deuterons taken up, *ND*, was calculated as:

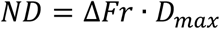

where *D_max_*was the maximum number of amide protons exchangeable for deuterium, calculated as the number of residues in a given peptide after subtracting the N-terminal residue and all proline residues.

### HDX-MS titration of CNOT1(800-999) by the N-CIM1 peptide

CNOT1(800-999) at 2 µM was exchanged for 2 h in deuterated reaction buffer in the presence of N-CIM1 (0, 1.4, 3.5, 14, 70, 140, 280, and 700 µM). The observed decrease in the mass of each CNOT1 peptide at higher N-CIM1 concentrations is due to the complex formation, which prevents deuterium uptake. These mass changes follow the equilibrium binding equation and can be interpreted in terms of local contribution to binding affinity. The apparent local dissociation constants, *K_d_^local^*, determined this way for the subsequent CNOT1 peptides reflect the local thermodynamic stability of the hydrogen bonds between N-CIM1 and individual CNOT1 fragments or allosterically stabilized in the complex, and are thus the upper-limit estimates for the overall *K_d_* of the entire CNOT1(800-999). The observed mass of a given CNOT1 peptic peptide, *M_t_*, after a given H/D exchange time, *t*, is a function of increasing ligand concentration, [*L*]:

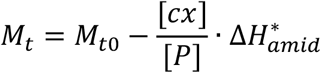

where *M_t0_*is the CNOT1 peptide mass in the absence of N-CIM1, [*P*] is CNOT1 concentration, [*L*] is N-CIM1 concentration, and [*cx*] is their complex concentration:

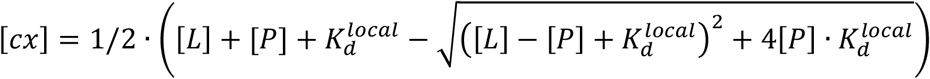

Δ*H*^∗^*_amid_* is the maximum number of amide hydrogens protected against the exchange due to the ligand binding, uncorrected for back-exchange. The ligand binding can alter amide protection against HDX due to direct binding or conformational changes. Non-linear least squares fitting of the above equations allowed the determination of *K_d_^local^* and Δ*H*^∗^*_amid_* for all CNOT1 peptides with their numerical errors. The maximum number of protected amide hydrogens corrected for back-exchange, ΔH_amid_, was calculated for each CNOT1 peptide individually, based on the correction factor, *bx*, determined for a specific CNOT1 peptide^67^, as:

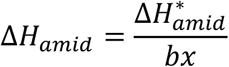

Δ*H_amid_*describes the spatial extent of the interaction as the maximum number of amide hydrogens protected from HDX upon the complex saturation by N-CIM1. The effectiveness of the interaction should thus be analyzed in terms of both parameters (Suppl. Table T6).

The corresponding Gibbs free energy, *ΔG°*, was calculated according to the equation:

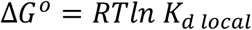

where *R* is the universal gas constant and *T* is the absolute temperature. The *ΔG°* values resulting from the *K_d_^local^* values, however, cannot be summed up since the free energy of an interacting system cannot simply be expressed as a sum of its components^89^.

### *In vitro* pulldown assays

0.2 µM GST-CNOT1(800-999) wt or mutant were incubated at 4°C for 1.5 hours with desired amount of MBP-GW182 SD, and/or MBP-TTP in chilled pulldown buffer (50 mM Tris/HCl, 150 mM NaCl, 1 mM EDTA, 2 mM DTT, 0.05% Triton X-100, pH 7.0). 15 µL of packed Glutathione Sepharose 4B resin (GE Healthcare) was added to each sample, followed by 15 min incubation on rotator at 4°C. Beads were washed three times with 1 ml of assay buffer. Proteins were eluted by adding 40 µL of 2x Laemmli sample buffer (60 mM Tris/HCl, pH 6.8, 2% sodium dodecyl sulfate, 10% glycerol, 5% β-mercaptoethanol, 0.01% bromophenol blue) and boiling them for 10 minutes at 95 °C. Samples were spun down for 1 min at ∼16600 g and 30 µL of each supernatant was separated by 12% SDS-PAGE. Gel was stained using Bio-Safe Coomasie Stain (Bio-Rad).

### Absorption and fluorescence spectroscopy

Absorption spectra were recorded using a Cary 50 spectrophotometer every 0.5 nm, at a rate of 120 nm/min, and baseline-corrected using the corresponding buffer. Emission spectra were measured using a Fluorolog 3.11 spectrofluorimeter in a thermostated (± 0.2 °C) semi-micro quartz cuvette (optical path length for excitation and emission, 4 and 10 mm, respectively) with magnetic stirring. The actual temperature inside the cuvette was measured using a thermocouple.

### Fluorescence correlation spectroscopy (FCS)

FCS measurements were performed at Axio Observer LSM 780 (Zeiss) with ConfoCor 3 and C-Apochromat 40x/1.2 W corr M27 essentially as described previously ^88^. The temperature inside each droplet (20 ± 0.5 °C) was measured after FCS acquisition using a calibrated micro-thermocouple (0.5 mm diameter). The structural parameter (s) was determined for each microscopic slide previously passivated with BSA using the appropriate calibration dye in deionized water: AF488 (D = 435 μm^2^s^−1^), Alexa Fluor 546 (D = 341 μm^2^s^−1^), Cy5 (D = 360 μm^2^s^−1^), rhodamine B (D = 450 μm^2^s^−1^), or fluorescein (D = 425 μm^2^s^−1^)^85–87^. All samples were prepared individually in Eppendorf tubes and incubated at 20 °C for at least 30 minutes before measurement.

Protein-protein FCS titrations with increasing concentrations of His_6_-Sumo-GW182 SD10 (10 nM-16 µM) were performed for Cy5-labeled GST-CNOT1(800-999), GST-CNOT1(800-999) E893A/Y900A, and α-chymotrypsinogen as a negative control, at 1 µM, in 50 mM NaH_2_PO_4_/Na_2_HPO_4_, 150 mM NaCl, 1 mM EDTA, 1 mM TCEP, pH 7.0, at 20 °C, in 15 µL droplets, in 2 technical replicates. Additionally, to account for viscosity changes, control titrations were run for free Cy5. Cy5 was excited at 633 nm using a HeNe laser (1% relative power) with an MBS 488/561/633 nm and an LP 655 nm emission filter. The triplet state population and relaxation time of Cy5 bound to the protein were included in the FCS data analysis as free parameters due to the possibility of Cy5 *cis*-*trans* isomerization. A two-component diffusion model was used to account for any residual free dye^88^.

Since the observed diffusion time, τ, in is proportional to the hydrodynamic radius of the diffusing species, it reflects the complex formation between two proteins. The dissociation constant, *K_d_*, was determined by measuring the increase in the τ value for GST-CNOT1(800-999) in response to the concentration of His_6_-Sumo-GW182 SD10, according to the equation:

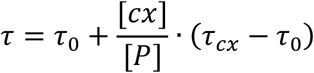

where *τ_0_* is the diffusion time in the absence of His_6_-Sumo-GW182 SD10, *τ_cx_* is the diffusion time of the complex, and [*cx*] and *ΔG°* were calculated using equations analogous to those used for HDX-MS titrations.

FCS displacement assays were performed for RhB-CIM1_c_ and FITC-TTP peptides at 20 nM, with GST-CNOT1(800-999), GST-CNOT1(800-999) E893A/Y900A, or human serum albumin as a negative control, at 10 µM, and by addition of unlabeled TTP peptide to 50 µM. This molar excess was estimated based on *K_d_* for His_6_-Sumo-GW182 SD10 and GST-CNOT1. The experiments were conducted in 50 mM Tris-HCl, 150 mM NaCl, 1 mM EDTA, 1 mM TCEP, pH 7.0, at 20 °C, in 8 µL droplets placed in wells formed by silicone slide gaskets (capacity 3-10 μL, Grace Bio-Labs), in 2 technical replicates. RhB-CIM1_c_ was excited at 561 nm using a DPSS laser (2% relative power), with an MBS 488/561 nm and LP 580 nm emission filter. FITC-TTP was excited at 488 nm using an Argon multiline laser (3% relative power), with an MBS 488 nm and BP 495-555 nm emission filter. The triplet state parameters of RhB and FITC were characterized in independent experiments, using 2-25% laser power, yielding an average triplet-state lifetime of ∼3 µs for both dyes, which was further used as a fixed parameter in FCS data fitting using a single-component model.

### Confocal microscopy

Confocal imaging, FRET, and FRAP experiments were performed on Axio Observer LSM 780 (Zeiss) with a Plan-Apochromat 63×/1.40 oil immersion objective at 25 °C, with scan dimensions of 34 × 34 µm or 24 × 24 µm, using either an MBS 488/561 nm with 495-555 nm emission (excitation at 488 nm, 0.7% Argon multiline laser power) and 690-720 nm emission (excitation at 561 nm, 1.4% DPSS laser power), or an MBS 488/561/633 nm with 505-550 nm emission (excitation at 488 nm, 0.5% Argon multiline laser power) and 684-754 nm emission (excitation at 633 nm, 0.7% HeNe laser power). Glass slides were passivated with a Silanization solution I (Supelco). Samples of 20 µL were placed in wells formed by Press-To-Seal silicone isolators (9 mm diameter, 0.8 mm depth, Grace Bio-Labs) under cover glasses. All samples were prepared individually in Eppendorf tubes, randomized in order, and incubated at 25 °C for at least 30 minutes before being transferred into passivated imaging chambers. Condensates were analysed after 12 and 24 hours.

### Liquid-liquid phase separation assays

LLPS experiments were performed for GW182 SD or its mixtures with GST-CNOT1(800- 999)-AF488 or FITC-TTP in 50 mM Tris-HCl, 150 mM NaCl, 0.5 mM EDTA, 1 mM TCEP, pH 7.2, at 25 °C, and for GW182 SD-mCherry or its mixtures with GST-CNOT1(800-999)- AF488 or GST-CNOT1(800-999)-AF488 E893A/Y900A in the same buffer supplemented by PEG 1000 up to the final concentration of 3% or 10% (w/v), according to the guidelines^93,94^. Control experiments were done for GW182 SD-mCherry (6 µM) with GST-AF488 (6 µM) or TTP (50 µM).

Comparison of droplet size distribution was performed for GW182 SD with GST- CNOT1(800-999)-AF488 (both at 6 µM) without PEG, and for GW182 SD-mCherry alone (6 µM) and in the presence of GST-CNOT1(800-999)-AF488, or GST-CNOT1(800-999)- AF488 E893A/Y900A, or GST-AF488 (each at 6 µM), or in the presence of TTP at 5 µM in the buffer with 10% (w/v) PEG 1000.

Multiprotein LLPS experiments were performed for GW182SD-mCherry (6 µM) titrated with increasing concentrations of GST-CNOT1(800-999)-AF488 (0-7 µM) or GST- CNOT1(800-999)-AF488 E893A/Y900A (0-7 µM), and for the 1:1 mixture of GW182SD-mCherry with GST-CNOT1(800-999)-AF488 (6 µM) with increasing concentrations of TTP (0-50 µM) in the buffer with 10% (w/v) PEG 1000.

### Liquid droplet number and diameter analysis

Image stacks of condensates were analyzed using an automated macro developed for Fiji (1.54p). Further data processing was performed with custom-written Python scripts. Both tools and a detailed usage protocol are freely available under the GNU General Public License v3 (GPLv3) on *GitHub* https://github.com/bialobrezuski.

### Förster Resonance Energy Transfer (FRET)

Based on the client-scaffold model, we designed an assay to test for proximity-dependent energy transfer between interacting proteins in the dense phase (Fig. 5a). The spectral overlap between the emission spectrum of AF488 and the absorption spectrum of Alexa Fluor 647 is approximately 1.5-fold lower than that for mCherry (Fig. 5e), *J* = 1.73 and 2.52 · 10^15^ M^-1^cm^-1^nm^4^, and the Fӧrster radius, *R_0_*, is 55.68 and 59.29 Å, respectively. Additionally, structural predictions suggest an average donor-acceptor distance of approximately 45 Å between CNOT1 and the GW182 SD mCherry fluorophore, whereas the corresponding distance to the Alexa 647 fluorophore attached to C378 is close to 100 Å (Fig. S19). Due to the sixth-power dependence of FRET efficiency on the distance, this difference is expected to reduce energy transfer by two orders of magnitude. Thus, the enhancement of the red signal under green excitation was specific to the AF488-mCherry FRET.

To quantify relative FRET intensity more precisely, we performed confocal imaging of droplets assembled in the presence of either CNOT1, CNOT1 EY, or CNOT1 titrated with increasing concentrations of TTP, using single-wavelength excitation of either the donor or the acceptor fluorophore. Upon direct excitation of the acceptor, red fluorescence could be detected from any GW182 SD-mCherry molecules, irrespective of whether they were engaged in the complex with CNOT1 or remained unbound within the condensates (Fig. 5g, S22). Excitation of the donor led to emission in both the donor (green) and acceptor (red) channels, indicating FRET between AF488-labeled CNOT1 and GW182 SD-mCherry in the dense phase (Fig. 5i).

To prove that the observed increase in the FRET channel comes from the extent of client recruitment into GW182 SD condensates, we quantified the relative FRET intensity (FRET_rel_) by comparing the mCherry fluorescence under donor excitation (FRET signal) to the total mCherry fluorescence originating from both bound and unbound GW182 SD-mCherry.

Each stack of dual-channel (AF488 - donor, mCherry - acceptor) confocal images was analysed independently. Fluorescence intensities were measured both inside and outside the condensates in each channel, with corrections for background and noise^94^. To maintain consistent brightness and contrast settings across channels and prevent the artificial enhancement of fluorescence due to auto-adjustment, the adjustment was first applied to the mCherry channel. Then, the resulting brightness and contrast parameters were then propagated to the AF488 channel, ensuring that the AF488 intensity was defined relative to the mCherry reference. This analysis was fully integrated into an automated pipeline combining Fiji macros with Python-based post-processing freely available under the GNU General Public License v3 (GPLv3) on *GitHub* https://github.com/bialobrezuski.

The relative FRET intensity was determined from dual-channel confocal images. First, condensates were imaged with donor excitation at 488 nm, and emission was collected in two channels: 495-555 nm (green, *I_donor_*) and 690-720 nm or 684-754 nm (red, *I_acceptor_*). The *I_acceptor_* signal represents only the emission from the acceptor excited *via* FRET from the bound donor. Next, imaging was performed with acceptor excitation at 561 nm or 633 nm, and emission was recorded in the respective channels (*I_total_*). The *I_total_* represents the total emission of the directly excited mCherry or Alexa Fluor 647. The fluorescence intensities were corrected for uneven illumination and background-subtracted, using an automated analysis pipeline combining Fiji macros with Python-based post processing.

The relative FRET intensity was calculated as:

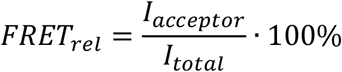

### Colocalization

Artefactual green-to-red bleed-through was excluded in independent experiments. Colocalization was evaluated on condensate image stacks as:

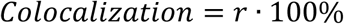

where *r* was the Pearson’s coefficient determined using the JaCoP plugin for Fiji/ImageJ^95^. The workflow was incorporated into the same automated analysis pipeline, combining Fiji macros with Python-based post processing.

### Delaunay triangulation

The Delaunay triangulation method was applied to the image stacks using the threshold model that was identified as the optimal one for droplet analysis^96^, *via* a Fiji/ImageJ plugin. This step was integrated into the existing automated scripts.

### Contact angles

Contact angle measurements were performed on confocal Z-stack images of condensates. Zen 2010 (Zeiss) was used to generate orthogonal (y-z) views. Contact angles were measured using the angle tool in both Zen 2010 and Fiji/ImageJ.

### Fluorescence recovery after photobleaching (FRAP)

A defined region of interest (ROI) within a droplet was photobleached, and fluorescence recovery within the ROI was recorded over time. Recovery curves were normalized and corrected for overall photobleaching using ZEN 2010 software. Curve fitting was performed using the ImageJ FRAP Analyzer plugin (v. 1.8) with a single-exponential model. For multiprotein condensates with GST-CNOT1(800-999) or GST-CNOT1(800-999) E893A/Y900A as clients, FRAP measurements were conducted on droplets of comparable size.

### FRET-based FRAP

In FRET-FRAP experiments, condensates were first imaged with donor excitation at 488 nm, and emission was collected in both the donor (*I_donor_*) and acceptor (*I_acceptor_*) channels. The *I_donor_* signal includes AF488 fluorescence from unbound CNOT1 client proteins and from those CNOT1 molecules bound to GW182 SD scaffold proteins that did not transfer energy to mCherry. The *I_acceptor_* signal originates exclusively from the GW182-mCherry molecules that are excited *via* FRET from bound CNOT1. Photobleaching was performed at 488 nm, and fluorescence recovery was recorded over time in the *I_donor_* and *I_acceptor_* channels. The former was used to determine the mobility of CNOT1, while the latter was used to evaluate the mobility of the CNOT1-GW182 SD complex. Measurements were carried out on droplets of comparable size and repeated across image stacks.

## Data analysis and visualization

Statistical data analysis and non-linear fitting were performed using Prism 7 (GraphPad Software), ZEN 2010 (Zeiss), Frap Analyzer, or custom-written Python scripts based on the Matplotlib library. Errors of reported values (one standard deviation) for HDX-MS and FCS were calculated according to the propagation rules^97^, on the basis of the numerical uncertainty resulting from the fitting and statistical variation across measurements, by using the exact differential method. Automated analysis of condensates was conducted using a pipeline that combines Fiji macros with Python-based post processing. All scripts and detailed documentation are freely available under the GNU General Public License v3 (GPLv3) on *GitHub* https://github.com/bialobrezuski. Data visualization was performed using Prism 7 (GraphPad Software), and the violin plots were generated using custom-written Python scripts. Docking of GW182 N-CIM1 to CNOT1 (PDB 4J8S)^7^ was done by CABS-dock^68^; structure visualization was done by BIOVIA Discovery Studio (Dassault Systèmes). Toy models and schematic figures were created using BioRender.com.

## Data availability

The Source data are provided with this paper.

## Acknowledgements

We thank Dr Marc R. Fabian for performing the pull-down experiment shown in Fig. 1c and for valuable discussions during the early development of the project, Dr Krzysztof Tarnowski for the assistance in HDX-MS experiments, Ms Barbara P. Klepka for excellent laboratory assistance, Dr Beata Kliszcz for helpful discussion, and Prof. E. Darzynkiewicz for access to his laboratory at the Division of Biophysics, Institute of Experimental Physics, Faculty of Physics, University of Warsaw. The work was partially supported by the Foundation for Polish Science International PhD Projects Programme at the Faculty of Physics, University of Warsaw, to M.K.C.-R., and by grants from the National Science Centre of Poland (Sonata-Bis 2016/22/E/NZ1/00656 to A.N., Preludium 2023/49/B/NZ1/04320 to M.K.B.). The research was performed in the NanoFun laboratories co-financed by ERDF within the POIG.02.02.00-00-025/09 Project.

## Contributions

M.K.B. performed the FCS experiments, conceptualized and performed the LLPS experiments, and wrote Python scripts for image analysis. M.K.C.-R. performed investigation, data curation, and formal analysis for the HDX-MS studies. A.M. performed the pulldown assays, with contributions from M.K.C.-R. M.K.C.-R. and A.M. performed mutagenesis. M.K.C.-R., A.M. and Z.S. performed protein expression and purification. N.S. consulted on the design of the experiments. M.D. developed the HDX-MS methodology, provided resources and validated the results. A.N. conceived the project, conceptualized the experiments, contributed to their execution, validated the results, supervised the study and wrote the manuscript with contributions from M.K.B. and M.K.C.-R. All authors contributed to experimental design, data analysis, writing and editing the manuscript.

## Ethics declarations

The authors declare no competing interests.

## References

1. Matoulkova, E., Michalova, E., Vojtesek, B. & Hrstka, R. The role of the 3’ untranslated region in post-transcriptional regulation of protein expression in mammalian cells. RNA Biol. 9, 563–576 (2012).

2. Moore, M. J. From Birth to Death: The Complex Lives of Eukaryotic mRNAs. Science 309, 1514– 1518 (2005).

3. Heck, A. M. & Wilusz, J. The Interplay between the RNA Decay and Translation Machinery in Eukaryotes. Cold Spring Harb. Perspect. Biol. 10, a032839 (2018).

4. Luo, Z. et al. High-resolution mapping of CCR4-NOT recruitment elements reveals transcriptome-wide drivers of mRNA decay. Cell Rep. 45, 117348 (2026).

5. Chen, C.-Y. A. & Shyu, A.-B. AU-rich elements: characterization and importance in mRNA degradation. Trends Biochem. Sci. 20, 465–470 (1995).

6. Brooks, S. A. & Blackshear, P. J. Tristetraprolin (TTP): interactions with mRNA and proteins, and current thoughts on mechanisms of action. Biochim. Biophys. Acta 1829, 666–679 (2013).

7. Fabian, M. R. et al. Structural basis for the recruitment of the human CCR4–NOT deadenylase complex by tristetraprolin. Nat. Struct. Mol. Biol. 20, 735–739 (2013).

8. Fu, M. & Blackshear, P. J. RNA-binding proteins in immune regulation: a focus on CCCH zinc finger proteins. Nat. Rev. Immunol. 17, 130–143 (2017).

9. Collart, M. A., Audebert, L. & Bushell, M. Roles of the CCR4-NOT complex in translation and dynamics of co-translation events. WIREs RNA 15, e1827 (2024).

10. Ukleja, M. et al. The architecture of the Schizosaccharomyces pombe CCR4-NOT complex. Nat. Commun. 7, 10433 (2016).

11. Lau, N.-C. et al. Human Ccr4–Not complexes contain variable deadenylase subunits. Biochem. J. 422, 443–453 (2009).

12. Bai, Y. et al. The CCR4 and CAF1 Proteins of the CCR4-NOT Complex Are Physically and Functionally Separated from NOT2, NOT4, and NOT5. Mol. Cell. Biol. 19, 6642–6651 (1999).

13. Bawankar, P., Loh, B., Wohlbold, L., Schmidt, S. & Izaurralde, E. NOT10 and C2orf29/NOT11 form a conserved module of the CCR4-NOT complex that docks onto the NOT1 N-terminal domain. RNA Biol. 10, 228–244 (2013).

14. Mathys, H. et al. Structural and Biochemical Insights to the Role of the CCR4-NOT Complex and DDX6 ATPase in MicroRNA Repression. Mol. Cell 54, 751–765 (2014).

15. Chen, Y. et al. A DDX6-CNOT1 Complex and W-Binding Pockets in CNOT9 Reveal Direct Links between miRNA Target Recognition and Silencing. Mol. Cell 54, 737–750 (2014).

16. Petit, A.-P. et al. The structural basis for the interaction between the CAF1 nuclease and the NOT1 scaffold of the human CCR4–NOT deadenylase complex. Nucleic Acids Res. 40, 11058–11072 (2012).

17. Basquin, J. et al. Architecture of the Nuclease Module of the Yeast Ccr4-Not Complex: the Not1-Caf1-Ccr4 Interaction. Mol. Cell 48, 207–218 (2012).

18. Boland, A. et al. Structure and assembly of the NOT module of the human CCR4-NOT complex. Nat. Struct. Mol. Biol. 20, 1289–1297 (2013).

19. Behm-Ansmant, I. et al. mRNA degradation by miRNAs and GW182 requires both CCR4:NOT deadenylase and DCP1:DCP2 decapping complexes. Genes Dev. 20, 1885–1898 (2006).

20. Sandler, H., Kreth, J., Timmers, H. Th. M. & Stoecklin, G. Not1 mediates recruitment of the deadenylase Caf1 to mRNAs targeted for degradation by tristetraprolin. Nucleic Acids Res. 39, 4373– 4386 (2011).

21. Chekulaeva, M. et al. miRNA repression involves GW182-mediated recruitment of CCR4–NOT through conserved W-containing motifs. Nat. Struct. Mol. Biol. 18, 1218–1226 (2011).

22. Cooke, A., Prigge, A. & Wickens, M. Translational Repression by Deadenylases. J. Biol. Chem. 285, 28506–28513 (2010).

23. Zekri, L., Kuzuoğlu-Öztürk, D. & Izaurralde, E. GW182 proteins cause PABP dissociation from silenced miRNA targets in the absence of deadenylation. EMBO J. 32, 1052–1065 (2013).

24. Jonas, S. & Izaurralde, E. Towards a molecular understanding of microRNA-mediated gene silencing. Nat. Rev. Genet. 16, 421–433 (2015).

25. Eulalio, A., Tritschler, F. & Izaurralde, E. The GW182 protein family in animal cells: New insights into domains required for miRNA-mediated gene silencing. RNA 15, 1433–1442 (2009).

26. Fabian, M. R. et al. miRNA-mediated deadenylation is orchestrated by GW182 through two conserved motifs that interact with CCR4–NOT. Nat. Struct. Mol. Biol. 18, 1211–1217 (2011).

27. Braun, J. E., Huntzinger, E., Fauser, M. & Izaurralde, E. GW182 Proteins Directly Recruit Cytoplasmic Deadenylase Complexes to miRNA Targets. Mol. Cell 44, 120–133 (2011).

28. Liu, J. et al. A role for the P-body component GW182 in microRNA function. Nat. Cell Biol. 7, 1261–1266 (2005).

29. Sala, L. et al. AGO2 silences mobile transposons in the nucleus of quiescent cells. Nat. Struct. Mol. Biol. 30, 1985–1995 (2023).

30. Liu, J., Liu, Z. & Corey, D. R. The Requirement for GW182 Scaffolding Protein Depends on Whether Argonaute Is Mediating Translation, Transcription, or Splicing. Biochemistry 57, 5247–5256 (2018).

31. Johnson, S. T., Chu, Y., Liu, J. & Corey, D. R. Impact of scaffolding protein TNRC6 paralogs on gene expression and splicing. RNA 27:1004–1016 (2021).

32. Guo, H., Kazadaeva, Y., Ortega, F. E., Manjunath, N. & Desai, T. J. Trinucleotide repeat containing 6c (TNRC6c) is essential for microvascular maturation during distal airspace sacculation in the developing lung. Dev. Biol. 430, 214–223 (2017).

33. Granadillo, J. L. et al. Pathogenic variants in *TNRC6B* cause a genetic disorder characterised by developmental delay/intellectual disability and a spectrum of neurobehavioural phenotypes including autism and ADHD. J. Med. Genet. 57, 717–724 (2020).

34. Cieplak-Rotowska, M. K. et al. Structural Dynamics of the GW182 Silencing Domain Including its RNA Recognition motif (RRM) Revealed by Hydrogen-Deuterium Exchange Mass Spectrometry. J. Am. Soc. Mass Spectrom. 29, 158–173 (2018).

35. Lazzaretti, D., Tournier, I. & Izaurralde, E. The C-terminal domains of human TNRC6A, TNRC6B, and TNRC6C silence bound transcripts independently of Argonaute proteins. RNA 15, 1059–1066 (2009).

36. Sheth, U. & Parker, R. Decapping and Decay of Messenger RNA Occur in Cytoplasmic Processing Bodies. Science 300, 805–808 (2003).

37. Youn, J.-Y. et al. High-Density Proximity Mapping Reveals the Subcellular Organization of mRNA-Associated Granules and Bodies. Mol. Cell 69, 517–532.e11 (2018).

38. Eulalio, A., Behm-Ansmant, I., Schweizer, D. & Izaurralde, E. P-Body Formation Is a Consequence, Not the Cause, of RNA-Mediated Gene Silencing. Mol. Cell. Biol. 27, 3970–3981 (2007).

39. Hubstenberger, A. et al. P-Body Purification Reveals the Condensation of Repressed mRNA Regulons. Mol. Cell 68, 144–157.e5 (2017).

40. Standart, N. & Weil, D. P-Bodies: Cytosolic Droplets for Coordinated mRNA Storage. Trends Genet. 34, 612–626 (2018).

41. Safieddine, A. et al. Cell-cycle-dependent mRNA localization in P-bodies. Mol. Cell 84, 4191–4208.e7 (2024).

42. Kamenska, A. et al. The DDX6–4E-T interaction mediates translational repression and P-body assembly. Nucleic Acids Res. 44, 6318–6334 (2016).

43. Kedersha, N. et al. Stress granules and processing bodies are dynamically linked sites of mRNP remodeling. J. Cell Biol. 169, 871–884 (2005).

44. Majerciak, V., Zhou, T., Kruhlak, M. J. & Zheng, Z.-M. RNA helicase DDX6 and scaffold protein GW182 in P-bodies promote biogenesis of stress granules. Nucleic Acids Res. 51, 9337–9355 (2023).

45. Ripin, N., Macedo De Vasconcelos, L., Ugay, D. A. & Parker, R. DDX6 modulates P-body and stress granule assembly, composition, and docking. J. Cell Biol. 223, e202306022 (2024).

46. Banani, S. F., Lee, H. O., Hyman, A. A. & Rosen, M. K. Biomolecular condensates: organizers of cellular biochemistry. Nat. Rev. Mol. Cell Biol. 18, 285–298 (2017).

47. Brangwynne, C. P., Tompa, P. & Pappu, R. V. Polymer physics of intracellular phase transitions. Nat. Phys. 11, 899–904 (2015).

48. Lin, Y., Protter, D. S. W., Rosen, M. K. & Parker, R. Formation and Maturation of Phase-Separated Liquid Droplets by RNA-Binding Proteins. Mol. Cell 60, 208–219 (2015).

49. Bose, M., Lampe, M., Mahamid, J. & Ephrussi, A. Liquid-to-solid phase transition of oskar ribonucleoprotein granules is essential for their function in Drosophila embryonic development. Cell 185, 1308–1324.e23 (2022).

50. Lazar, T., Connor, A., DeLisle, C. F., Burger, V. & Tompa, P. Targeting protein disorder: the next hurdle in drug discovery. Nat. Rev. Drug Discov. 10.1038/s41573-025-01220-6 (2025).

51. Liu, C. et al. Diffusing protein binders to intrinsically disordered proteins. Nature 10.1038/s41586-025-09248-9 (2025).

52. Ditlev, J. A., Case, L. B. & Rosen, M. K. Who’s In and Who’s Out—Compositional Control of Biomolecular Condensates. J. Mol. Biol. 430, 4666–4684 (2018).

53. Farag, M., Borcherds, W. M., Bremer, A., Mittag, T. & Pappu, R. V. Phase separation of protein mixtures is driven by the interplay of homotypic and heterotypic interactions. Nat. Commun. 14, 5527 (2023).

54. Sheu-Gruttadauria, J. & MacRae, I. J. Phase Transitions in the Assembly and Function of Human miRISC. Cell 173, 946–957.e16 (2018).

55. Currie, S. L. & Rosen, M. K. Using quantitative reconstitution to investigate multicomponent condensates. RNA 28, 27–35 (2022).

56. Bulbrook, D. et al. Tryptophan-Mediated Interactions between Tristetraprolin and the CNOT9 Subunit Are Required for CCR4-NOT Deadenylase Complex Recruitment. J. Mol. Biol. 430, 722–736 (2018).

57. Ruscica, V. et al. Direct role for the Drosophila GIGYF protein in 4EHP-mediated mRNA repression. Nucleic Acids Res. 47, 7035–7048 (2019).

58. Peter, D. et al. Molecular basis for GIGYF–Me31B complex assembly in 4EHP-mediated translational repression. Genes Dev. 33, 1355–1360 (2019).

59. Sobti, M., Mead, B. J., Stewart, A. G., Igreja, C. & Christie, M. Molecular basis for GIGYF– TNRC6 complex assembly. RNA 29, 724–734 (2023).

60. Christie, M. & Igreja, C. eIF4E-homologous protein (4EHP): a multifarious cap-binding protein. FEBS J. 290, 266–285 (2023).

61. Harris Snell, P., et al. Coordinated regulation of mRNA translation and stability by ZC3H7A and ZC3H7B RNA-binding proteins. Cell Rep. 45, 117511 (2026).

62. Welte, T. et al. Convergence of multiple RNA-silencing pathways on GW182/TNRC6. Mol. Cell 83, 2478–2492.e8 (2023).

63. Stowell, J. A. W. et al. Phosphorylation-dependent tuning of mRNA deadenylation rates. Nat. Struct. Mol. Biol. 33, 63–70 (2026).

64. James, E. I., Murphree, T. A., Vorauer, C., Engen, J. R. & Guttman, M. Advances in Hydrogen/Deuterium Exchange Mass Spectrometry and the Pursuit of Challenging Biological Systems. Chem. Rev. 122, 7562–7623 (2022).

65. Masson, G. R. et al. Recommendations for performing, interpreting and reporting hydrogen deuterium exchange mass spectrometry (HDX-MS) experiments. Nat. Methods 16, 595–602 (2019).

66. Chen, W. & Komives, E. A. Combination of single-molecule Förster resonance energy transfer and hydrogen-deuterium exchange mass spectrometry toward dynamic structural biology. Curr. Opin. Struct. Biol. 99, 103300 (2026).

67. Cieplak-Rotowska, M. K., Dadlez, M. & Niedzwiecka, A. Exploring the CNOT1(800–999) HEAT Domain and Its Interactions with Tristetraprolin (TTP) as Revealed by Hydrogen/Deuterium Exchange Mass Spectrometry. Biomolecules 15, 403 (2025).

68. Kurcinski, M., Jamroz, M., Blaszczyk, M., Kolinski, A. & Kmiecik, S. CABS-dock web server for the flexible docking of peptides to proteins without prior knowledge of the binding site. Nucleic Acids Res. 43, W419–W424 (2015).

69. Davey, N. E. SLiMDisc: short, linear motif discovery, correcting for common evolutionary descent. Nucleic Acids Res. 34, 3546–3554 (2006).

70. Davey, N. E., Simonetti, L. & Ivarsson, Y. The next wave of interactomics: Mapping the SLiM-based interactions of the intrinsically disordered proteome. Curr. Opin. Struct. Biol. 80, 102593 (2023).

71. André, A. A. M. & Spruijt, E. Liquid–Liquid Phase Separation in Crowded Environments. Int. J. Mol. Sci. 21, 5908 (2020).

72. Rothenberg, K. E., Scott, D. W., Christoforou, N. & Hoffman, B. D. Vinculin Force-Sensitive Dynamics at Focal Adhesions Enable Effective Directed Cell Migration. Biophys. J. 114, 1680–1694 (2018).

73. Liu, J., Valencia-Sanchez, M. A., Hannon, G. J. & Parker, R. MicroRNA-dependent localization of targeted mRNAs to mammalian P-bodies. Nat. Cell Biol. 7, 719–723 (2005).

74. Kumar, M. et al. ELM—the eukaryotic linear motif resource in 2020. Nucleic Acids Res. gkz1030 (2019).

75. Marcotrigiano, J., Gingras, A.-C., Sonenberg, N. & Burley, S. K. Cap-Dependent Translation Initiation in Eukaryotes Is Regulated by a Molecular Mimic of eIF4G. Mol. Cell 3, 707–716 (1999).

76. Pekovic, F. et al. Multivalent interactions with CCR4–NOT and PABPC1 determine mRNA repression efficiency by tristetraprolin. Nat. Commun. 16, 7528 (2025).

77. Schneider, T. N. et al. Engineering nanocondensate formation through sequence composition and patterning (2026).

78. Mizutani, A., Tan, C., Sugita, Y. & Takada, S. Heterogeneous condensates of transcription factors in embryonic stem cells: Molecular simulations. Biophys. J. 124, 1587–1598 (2025).

79. Espinosa, J. R. et al. Liquid network connectivity regulates the stability and composition of biomolecular condensates with many components. Proc. Natl. Acad. Sci. 117, 13238–13247 (2020).

80. Grigorev, V., Wingreen, N. S. & Zhang, Y. Conformational Entropy of Intrinsically Disordered Proteins Bars Intruders from Biomolecular Condensates. PRX Life 3, 013011 (2025).

81. Brothers, W. R., Ali, F., Kajjo, S. & Fabian, M. R. The EDC4-XRN1 interaction controls P-body dynamics to link mRNA decapping with decay. EMBO J. 42, e113933 (2023).

82. Malsick, L. E. & Wilusz, J. Dynamic “Cap”-abilities of P-bodies and the XRN1-EDC4 axis. EMBO J. 42, EMBJ2023115310 (2023).

83. Smokers, I. B. A., Visser, B. S., Slootbeek, A. D., Huck, W. T. S. & Spruijt, E. How Droplets Can Accelerate Reactions─Coacervate Protocells as Catalytic Microcompartments. Acc. Chem. Res. 57, 1885–1895 (2024).

84. Cheng, S.-J. et al. Regulation of stress granule maturation and dynamics by poly(ADP-ribose) interaction with PARP13. Nat. Commun. 16, 621 (2025).

85. Villegas, J. A. & Levy, E. D. A unified statistical potential reveals that amino acid stickiness governs nonspecific recruitment of client proteins into condensates. Protein Sci. 31, e4361 (2022).

86. Kelley, F. M. et al. Controlled and orthogonal partitioning of large particles into biomolecular condensates (2024).

87. Cermakova, K. & Hodges, H. C. Interaction modules that impart specificity to disordered protein. Trends Biochem. Sci. 48, 477–490 (2023).

88. Waszkiewicz, R. et al. Hydrodynamic Radii of Intrinsically Disordered Proteins: Fast Prediction by Minimum Dissipation Approximation and Experimental Validation. J. Phys. Chem. Lett. 15, 5024– 5033 (2024).

89. Mark, A. E. & Van Gunsteren, W. F. Decomposition of the Free Energy of a System in Terms of Specific Interactions. J. Mol. Biol. 240, 167–176 (1994).

90. Gendron, P.-O., Avaltroni, F. & Wilkinson, K. J. Diffusion Coefficients of Several Rhodamine Derivatives as Determined by Pulsed Field Gradient–Nuclear Magnetic Resonance and Fluorescence Correlation Spectroscopy. J. Fluoresc. 18, 1093–1101 (2008).

91. Culbertson, C. Diffusion coefficient measurements in microfluidic devices. Talanta 56, 365–373 (2002).

92. Petrášek, Z. & Schwille, P. Precise Measurement of Diffusion Coefficients using Scanning Fluorescence Correlation Spectroscopy. Biophys. J. 94, 1437–1448 (2008).

93. Alberti, S., Gladfelter, A. & Mittag, T. Considerations and Challenges in Studying Liquid-Liquid Phase Separation and Biomolecular Condensates. Cell 176, 419–434 (2019).

94. Alberti, S. et al. A User’s Guide for Phase Separation Assays with Purified Proteins. J. Mol. Biol. 430, 4806–4820 (2018).

95. Bolte, S. & Cordelières, F. P. A guided tour into subcellular colocalization analysis in light microscopy. J. Microsc. 224, 213–232 (2006).

96. Nichele, L., Persichetti, V., Lucidi, M. & Cincotti, G. Quantitative evaluation of ImageJ thresholding algorithms for microbial cell counting. OSA Contin. 3, 1417 (2020).

97. Taylor, J. Introduction to Error Analysis, the Study of Uncertainties in Physical Measurements, 2nd Edition. Published by University Science Books (1997).

