## Supplementary information, figures, tables for "GW182 silencing domain engages the tristetraprolin-binding pocket of CNOT1 to recruit CNOT1 into multiprotein condensates"

July 16, 2026

**Supplementary figure S1 to Fig 1e.**

**(a)** HDX differences (ΔFr) for pepsin-generated CNOT1 peptides in the presence of the CIM2 peptide at 90.5 μM *vs* the *apo* state (CNOT1 at 2 μM) after 2h of deuteration. Vertical thin lines show the standard deviation, n=3. Two regions of the highest protection against HDX in the presence of TTP- or GW182-derived peptides are marked in grey (as in Fig 1e). (b) The best-scoring model of the CNOT1(800-999) complex with N-CIM1 (magenta) obtained by coarse-grained docking (RSMD 3.8 Å) superimposed on the crystal structure of CNOT1 with the TTP peptide (blue). The aromatic side chain of Trp1300 of GW182 (magenta balls-and-sticks) occupies the same position as Phe319 of TTP (blue balls-and-sticks), while the N-CIM1 peptide backbone adopts a slightly altered orientation relative to the hydrophobic groove.

**a**

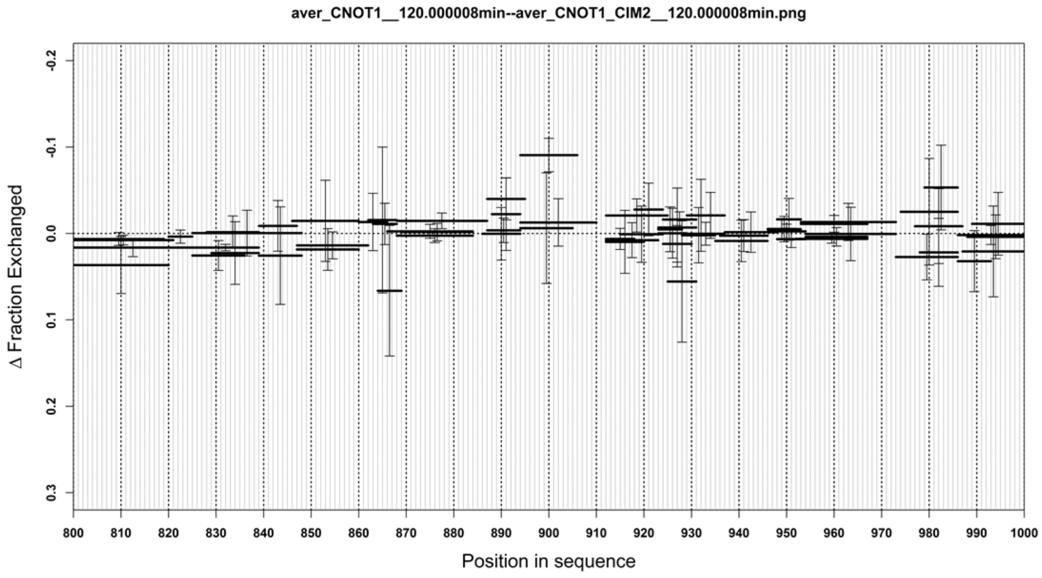

**b**

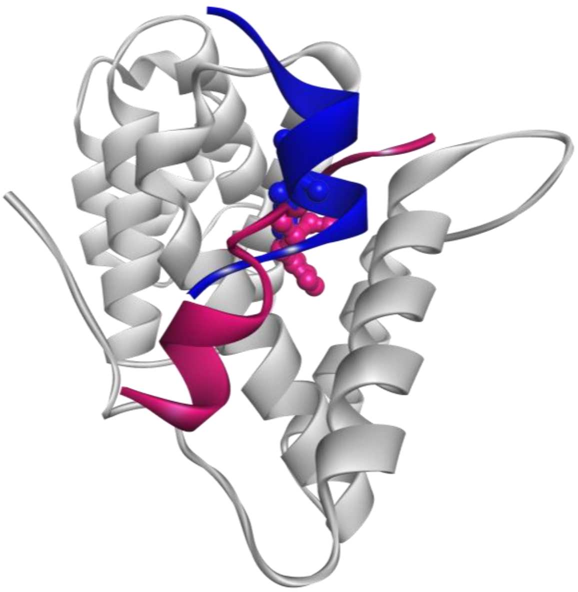

**Supplementary figure S2 to Fig 1f.**

Kinetics of deuterium uptake by CNOT1(800-999) at 2 µM alone (black) and in the presence of N-CIM1 peptide at 140 µM (magenta). *ND* - maximum number of protons exchanged.

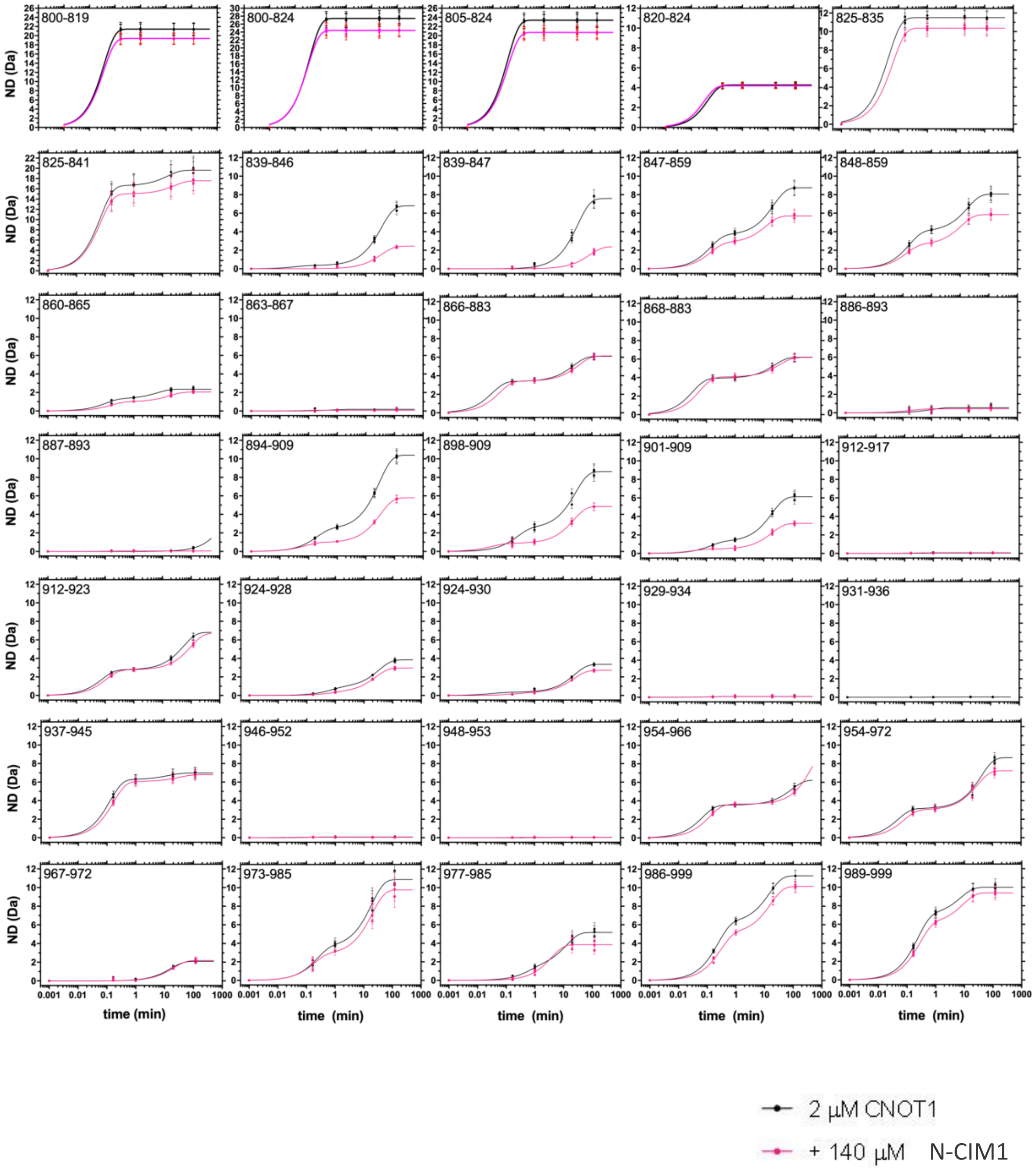

**Supplementary figure S3 to Fig 1f.**
Kinetics of deuterium uptake by CNOT1(800-999) at 2 µM alone (black), in the presence of the CIM1 peptide at 12.6 µM (orange) or 126 µM (red). *ND* - maximum number of protons exchanged. The control data (the *apo* state) is the same as in [6].

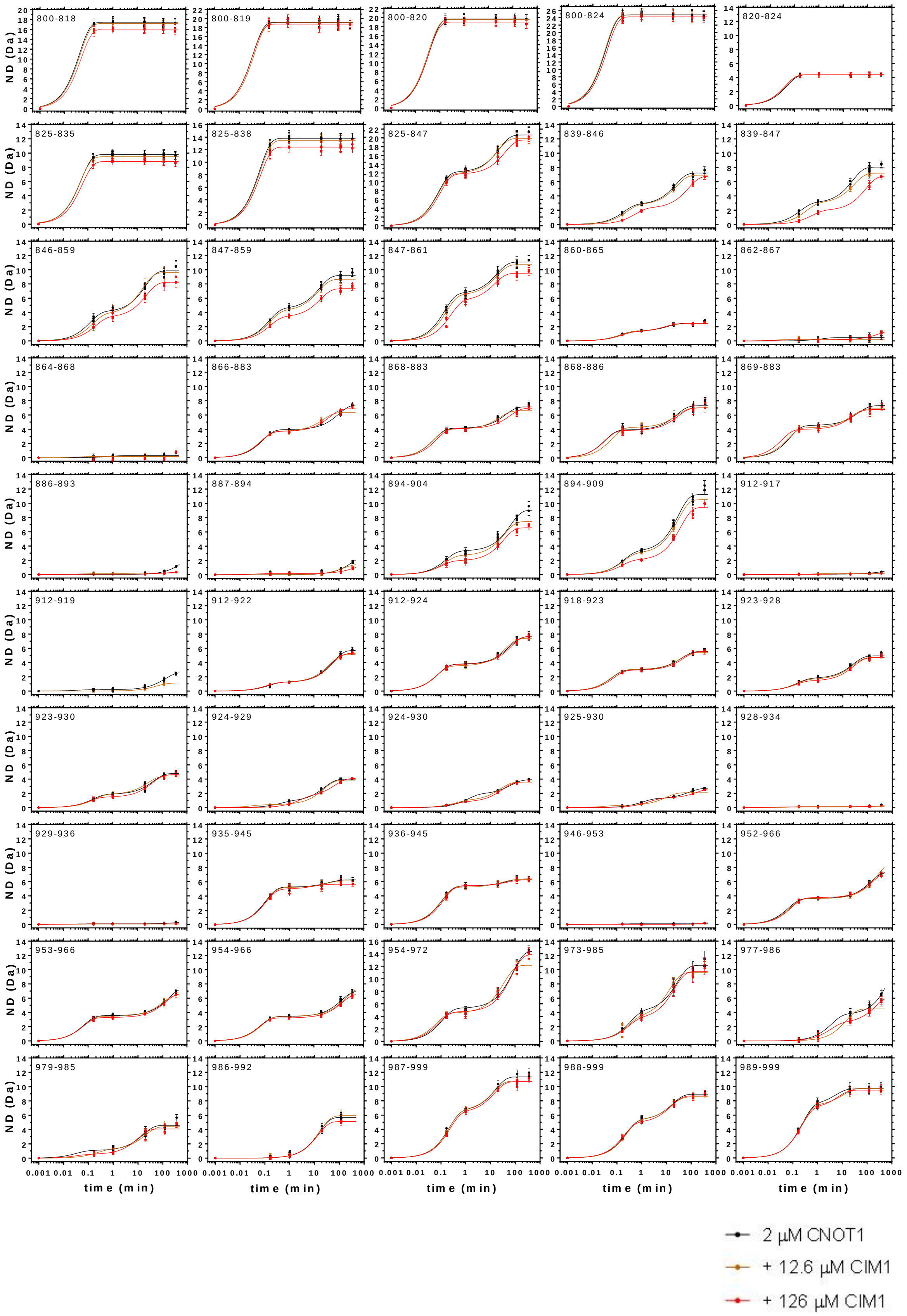

**Supplementary figure S4 to Fig 1f.**

Kinetics of deuterium uptake by CNOT1(800-999) at 2 µM alone or in the presence of SD10 at 20 µM (blue). Please note, colors differ from those used in Fig 1f. *ND* - maximum number of protons exchanged.

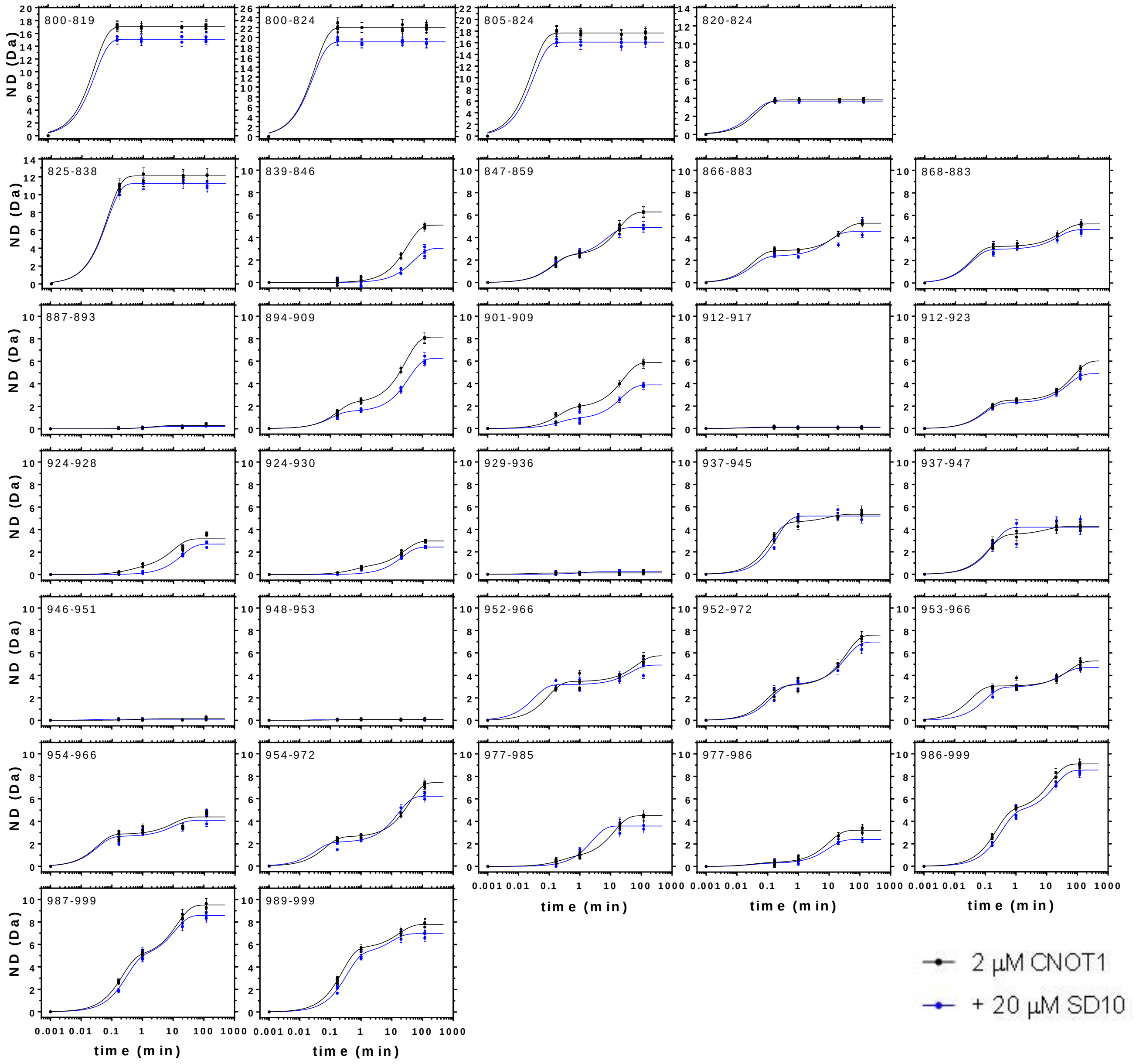

### Supplementary figure S5 to Fig 1f.

Kinetics of deuterium uptake by CNOT1(800-999) at 2 µM alone (black), in the presence of the TTP peptide at 3.3 µM (green) or the CIM1 peptide at 126 µM (red). Please note, colors differ from those used in Fig 1f. *ND* - maximum number of protons exchanged. Independent biological replicate of data in Fig S3: CNOT1 in the presence of the CIM peptide at 126 µM starting from protein expression and purification.

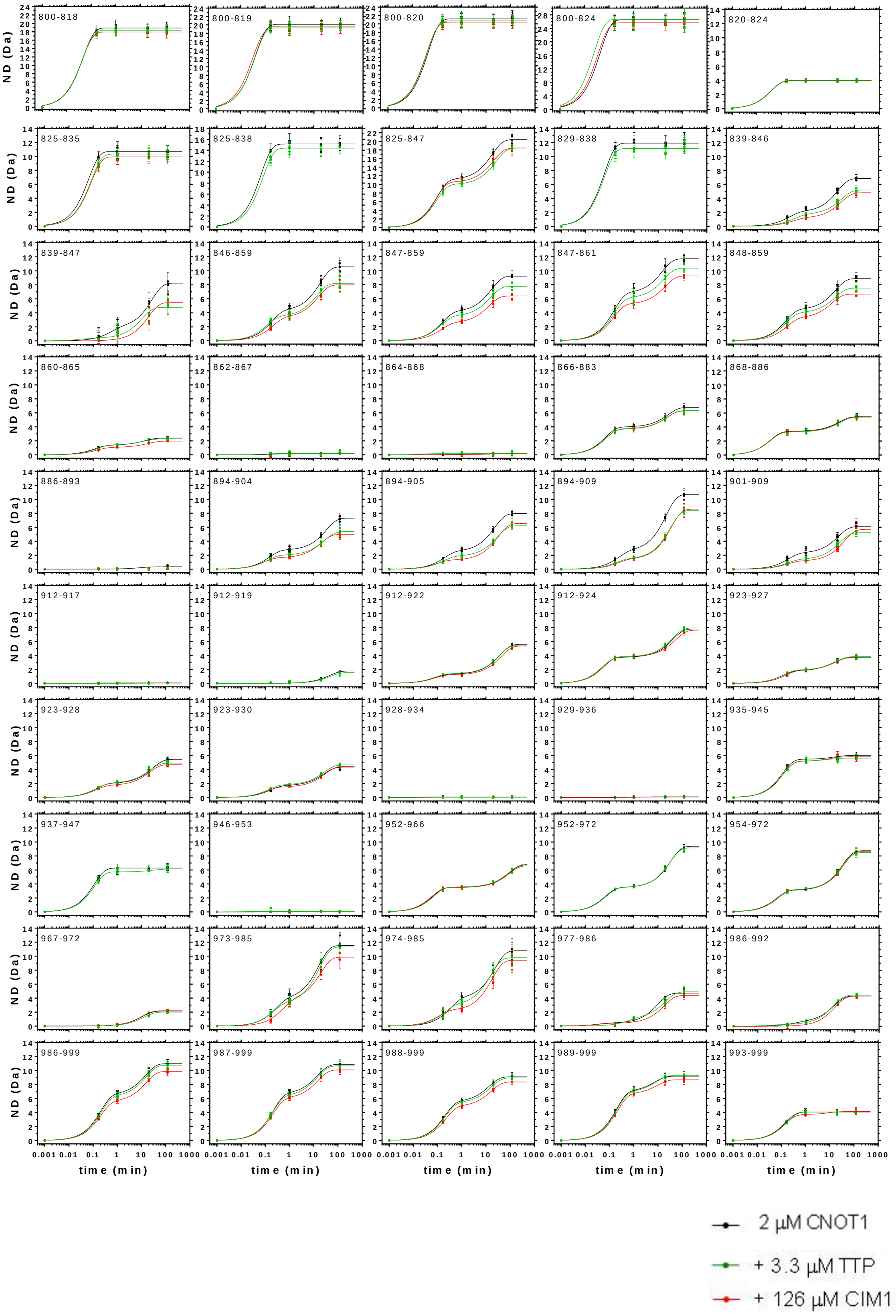

**Supplementary figure S6 to Fig 1f.**
Kinetics of deuterium uptake of chosen peptides of CNOT1(800-999) E893A/Y900A at 2 µM alone (black), in the presence of the TTP peptide at 3.3 µM (green) or the CIM1 peptide at 126 µM (red). Please note, colors differ from those used in Fig 1f. *ND* - maximum number of protons exchanged.

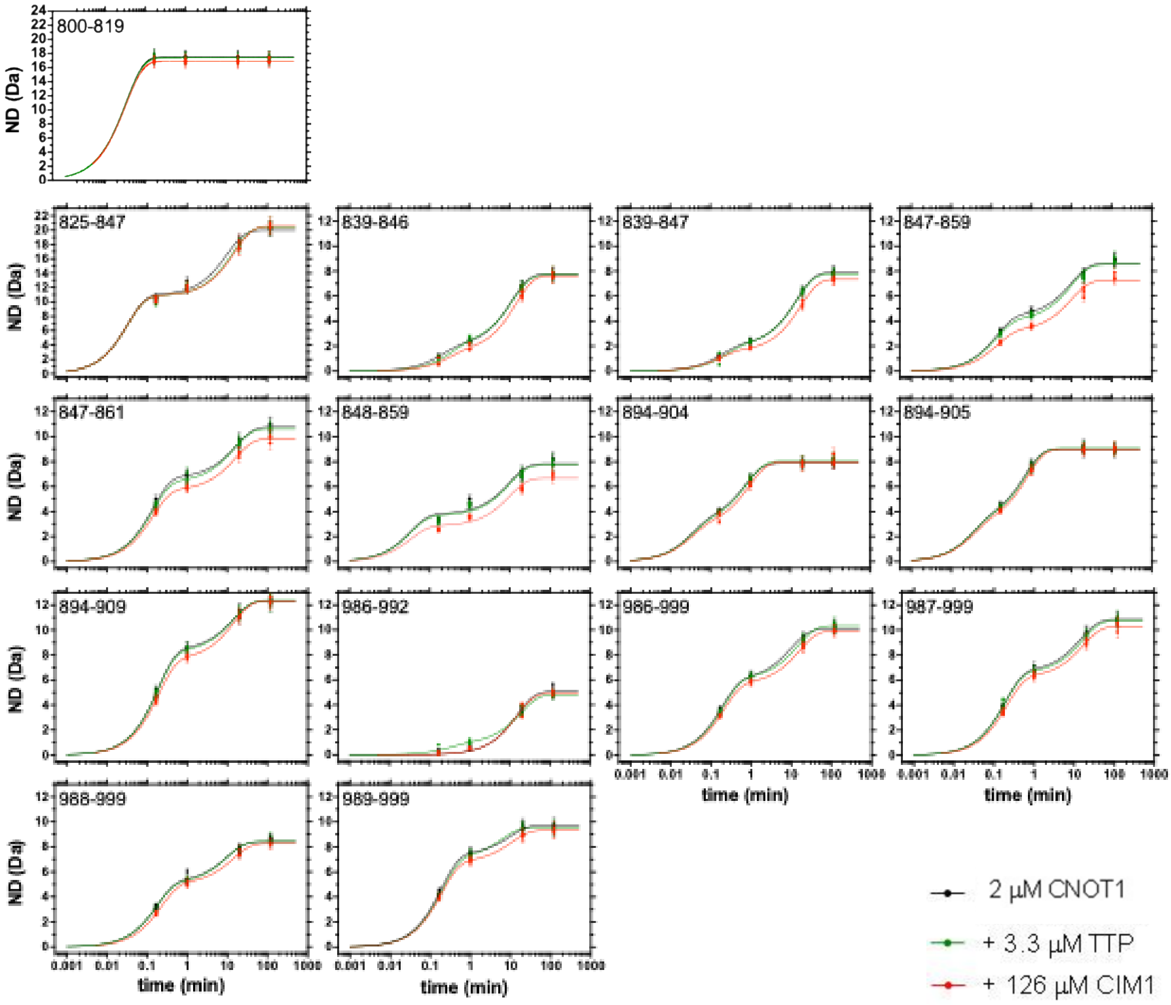

**Supplementary figure S7 to Fig 2a.**

Apparent local N-CIM1 binding isotherms (or titration curves) obtained for subsequent CNOT1(800-999) peptides after 2 hours of H/D exchange in the presence of increasing concentrations of the N-CIM1 peptide. CNOT1 concentration was 2 µM. Vertical axis: mass of peptide in Da. Horizontal axis: N-CIM1 peptide concentration in mM. MS data uncorrected for back-exchange.

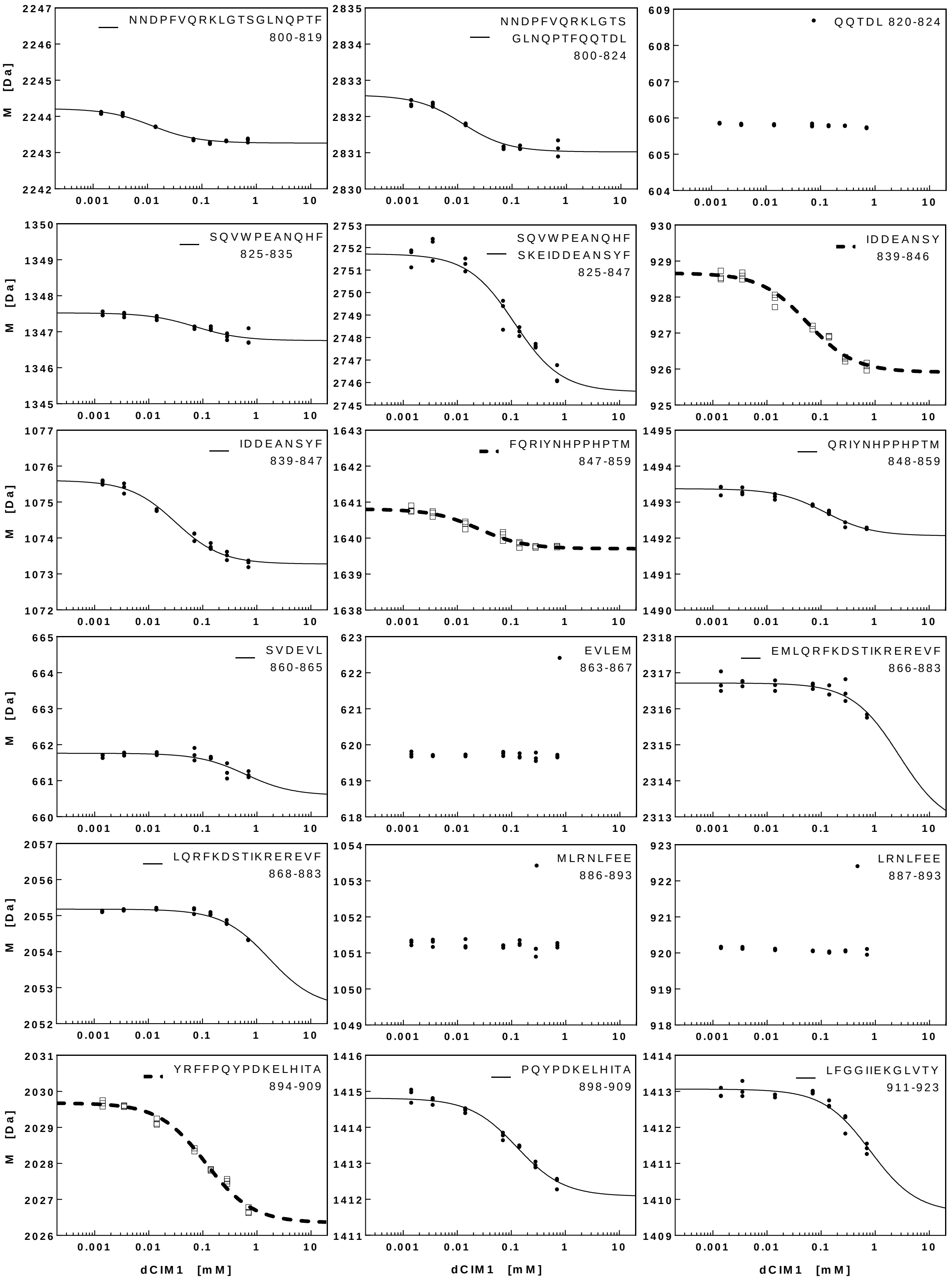

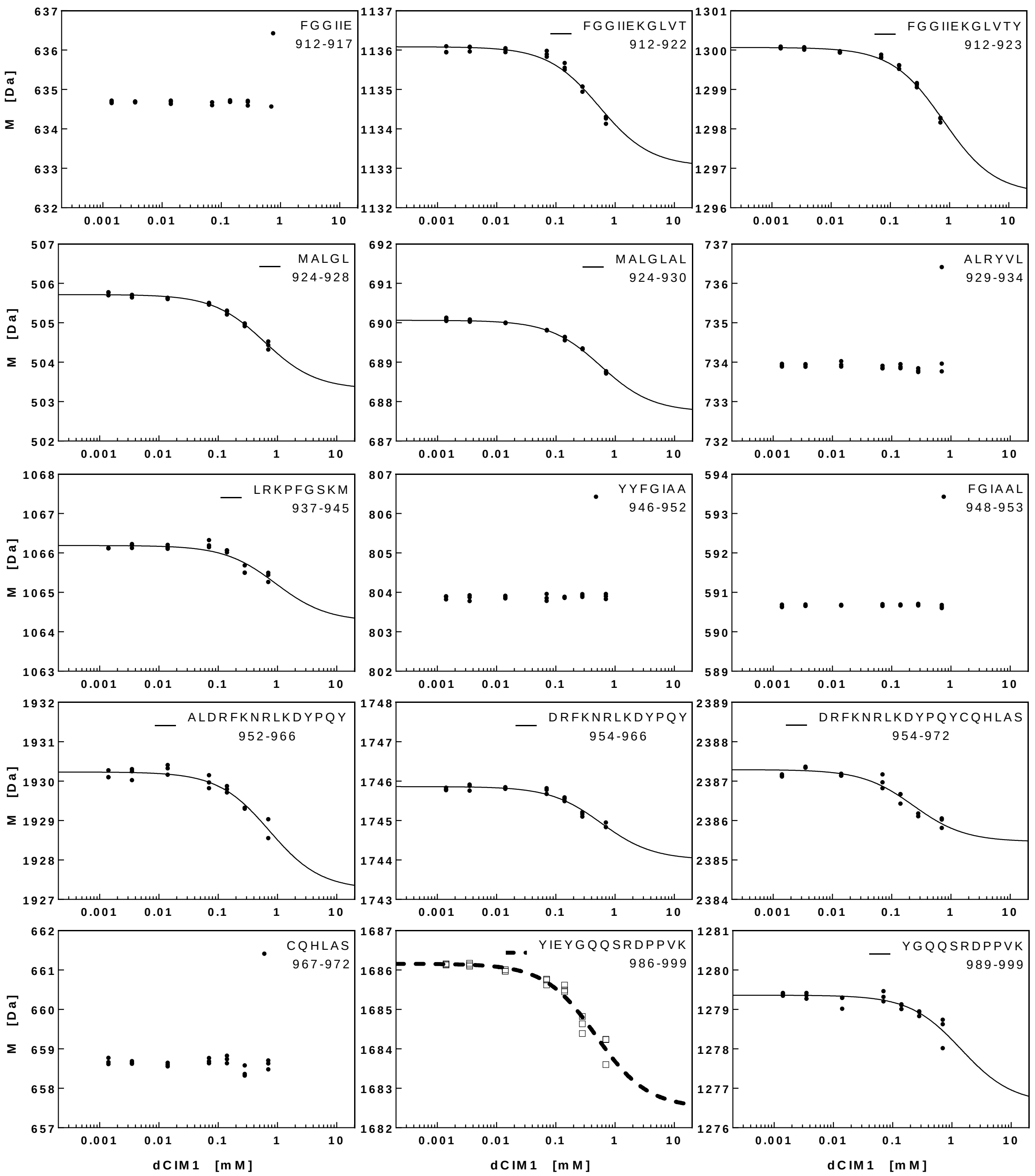
FGGIIE
912-917
FGGIIEKGLVT
912-922
FGGIIEKGLVTY
912-923
MALGL
924-928
MALGLAL
924-930
ALRYVL
929-934
LRKPFGSKM
937-945
YYFGIAA
946-952
FGIAAL
948-953
ALDRFKNRLKDYPQY
952-966
DRFKNRLKDYPQY
954-966
DRFKNRLKDYPQYCQHLAS
954-972
CQHLAS
967-972
YIEYGQQSRDPPVK
986-999
YGQQSRDPPVK
989-999
M [Da]
dCIM1 [mM]

**Supplementary figure S8 to Fig 2e,f.**

GST-pulldown assay for titration of GST-CNOT1(800-999) containing selected point mutations with either MBP-GW182 SD (left column) or MBP-TTP (right column). See also Fig. S9.

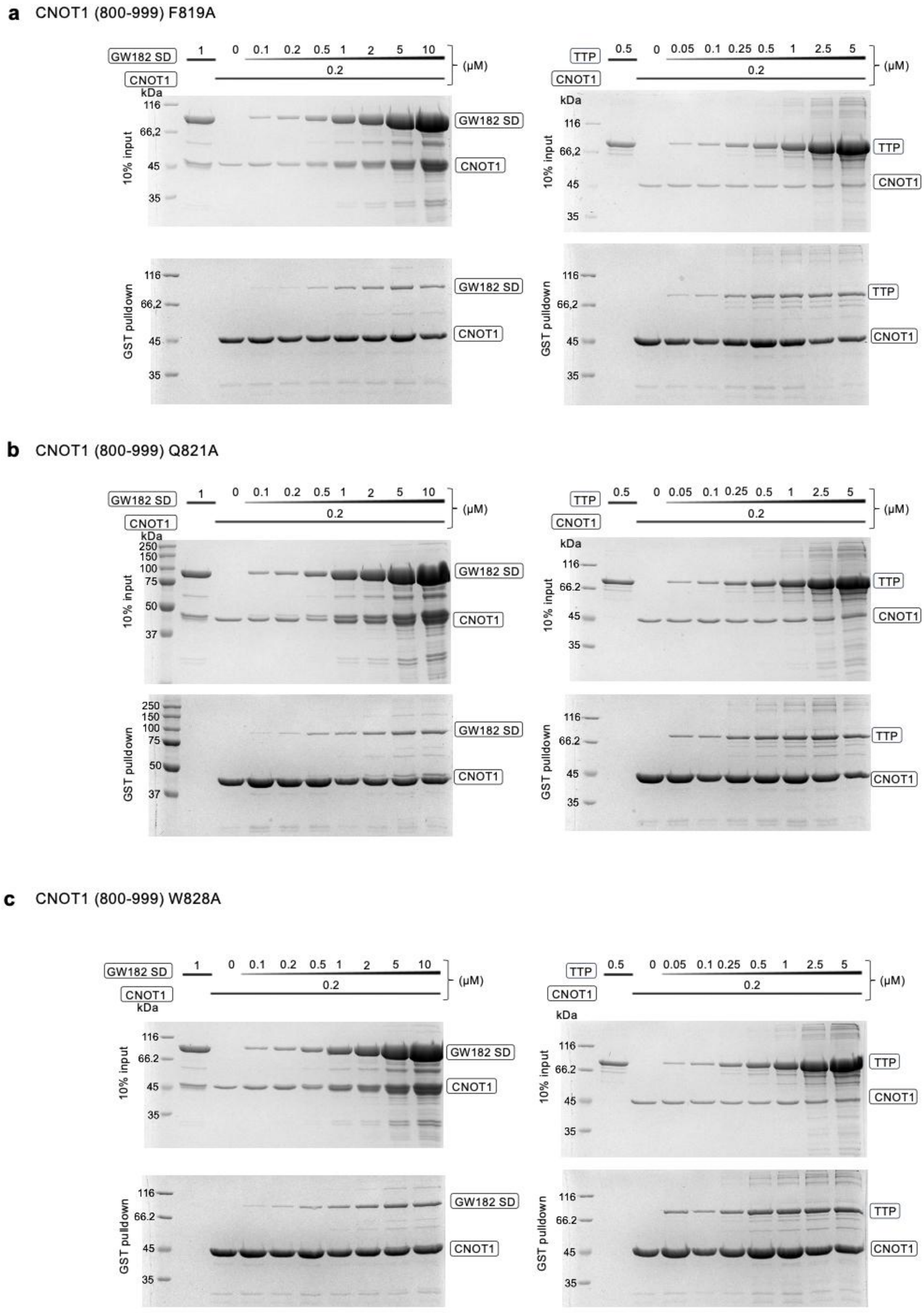

**d** CNOT1 (800-999) D840A

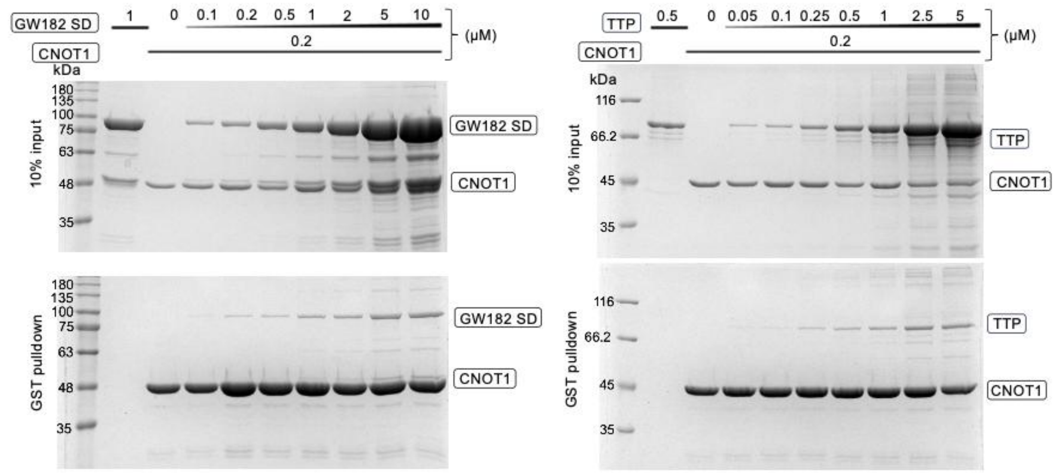

**e** CNOT1 (800-999) E842A

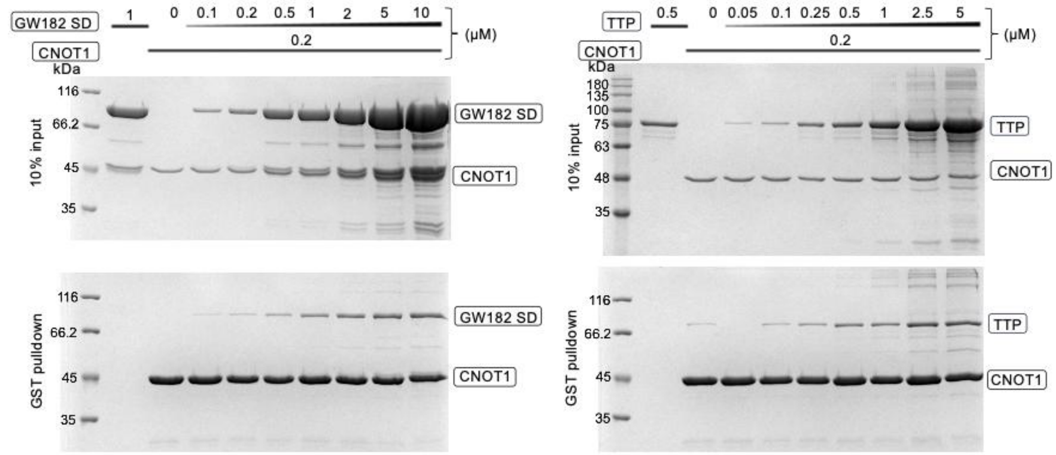

**f** CNOT1 (800-999) W828A

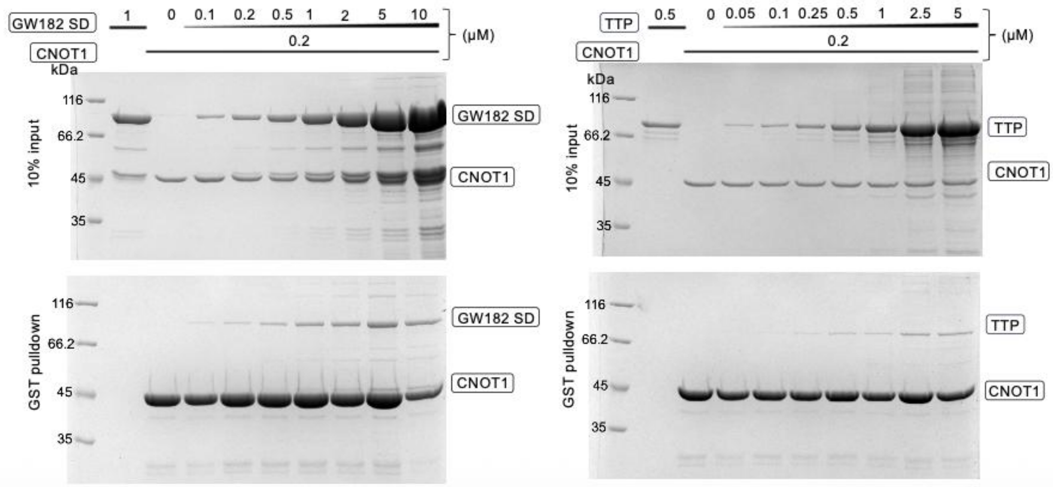

**g** CNOT1 (800-999) Y846A

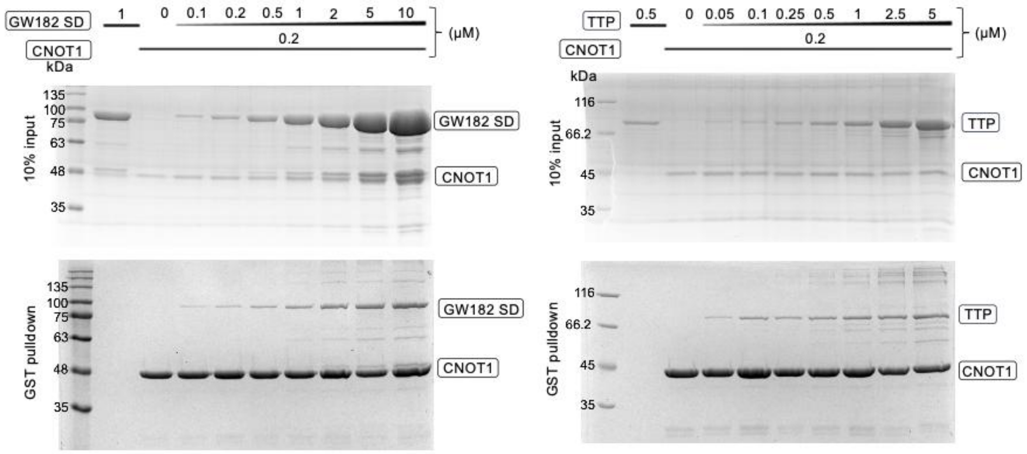

**h** CNOT1 (800-999) F847A

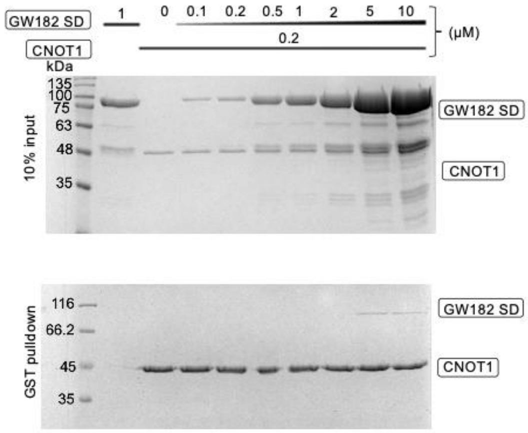

**i** CNOT1 (800-999) F847A/Q848A

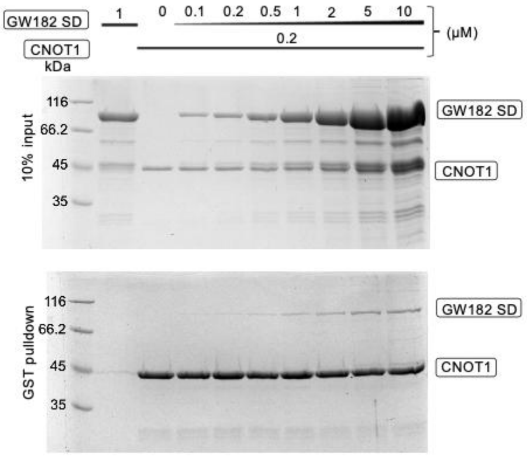

**j** CNOT1 (800-999) Q848A

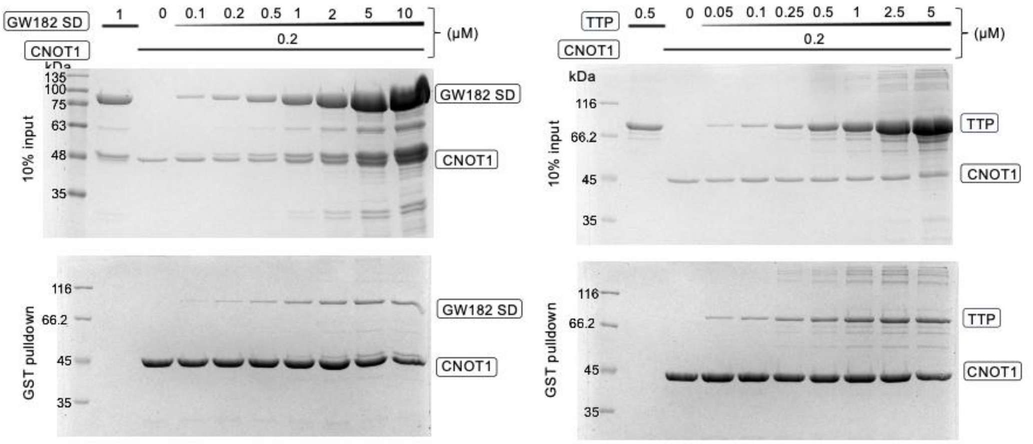

**k** CNOT1 (800-999) F847A/Q848A/P857A/T852A

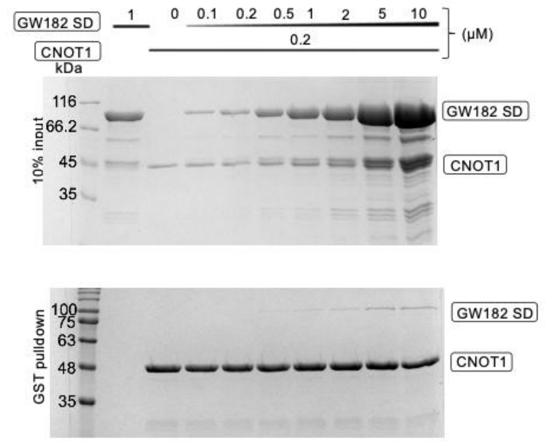

**l** CNOT1 (800-999) Y851A

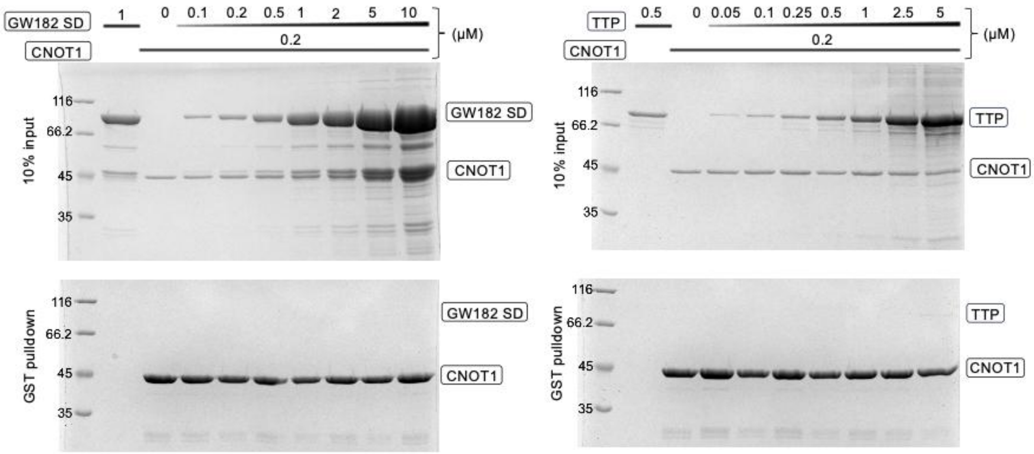

**m** CNOT1 (800-999) N852A

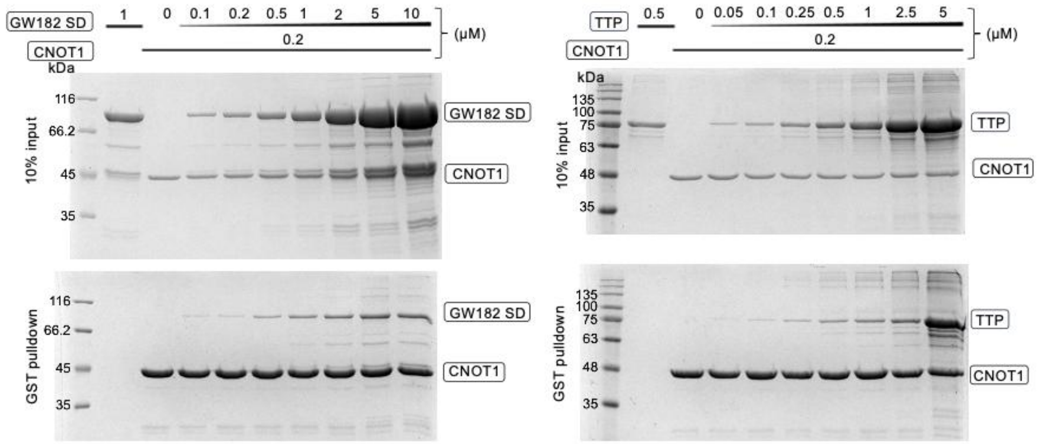

**n** CNOT1 (800-999) N852A/H853A/P854A

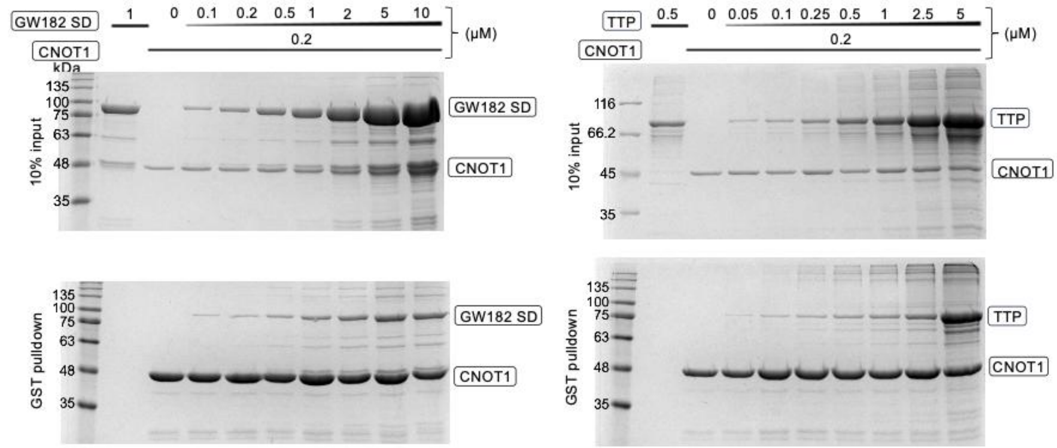

**o** CNOT1 (800-999) P855A/H856A

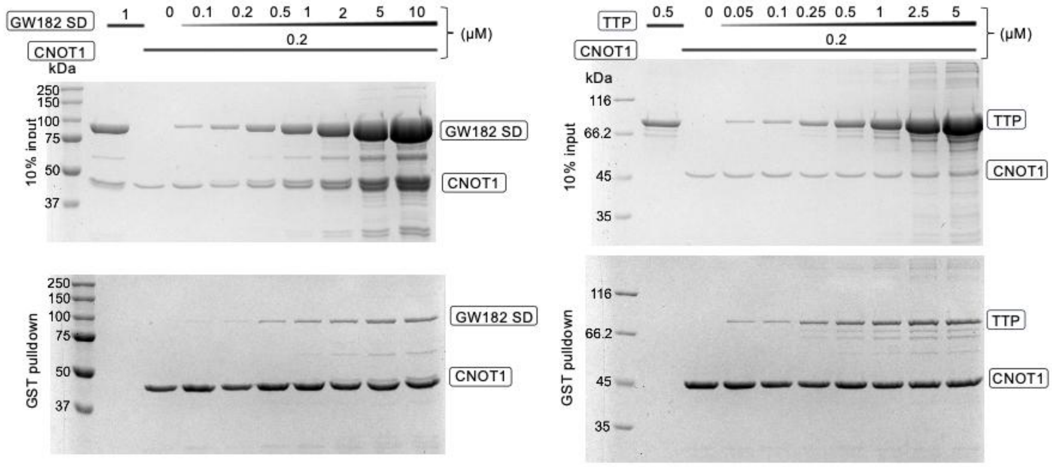

**p** CNOT1 (800-999) P857A/T858A

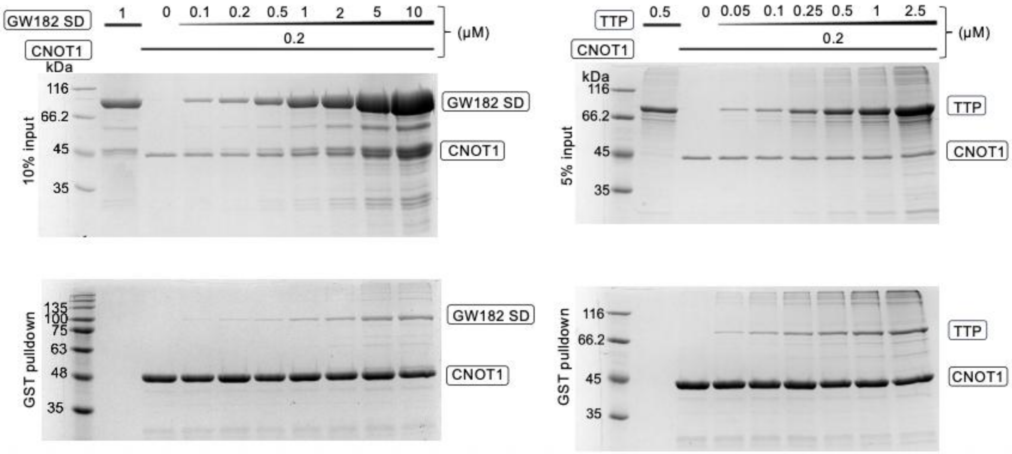

**q** CNOT1 (800-999) P857A/T858A/E893A/Y900A

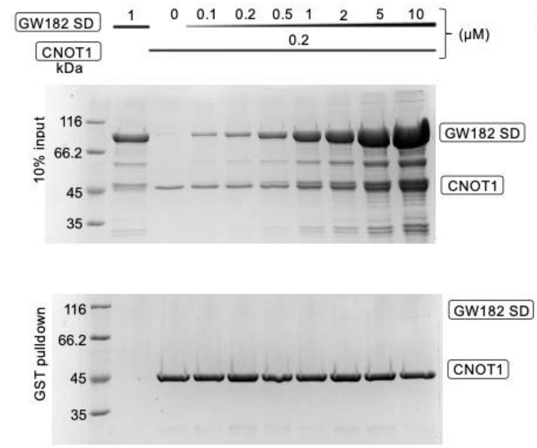

**r** CNOT1 (800-999) E893A/Y900A

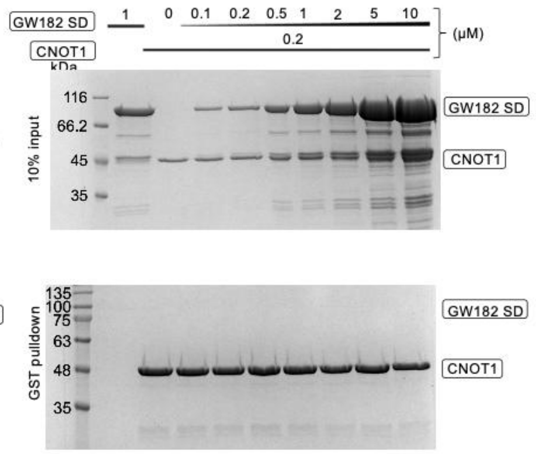

**s** CNOT1 (800-999) E904A

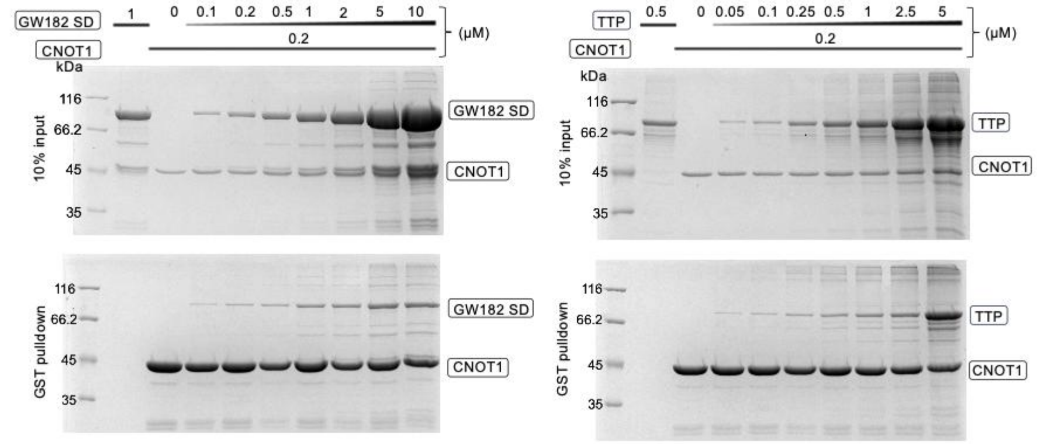

### Supplementary figure S9 to Fig 2b,e and S8.

MBP-SDfl satellite band similar in size to GST-CNOT1 was found to be a degradation product of MBP-SDfl consisting of SDfl (TNRC6C 1260-1690) with additional 6 residues from the MBP-tag:

A) The product was found to have a mass of : 47061.5 ± 5.9 Da, while SDfl without the MBP-tag has a predicted mass of 46316.21 Da, and with 6 additional residues (MHVPNY) 47058.08 Da.

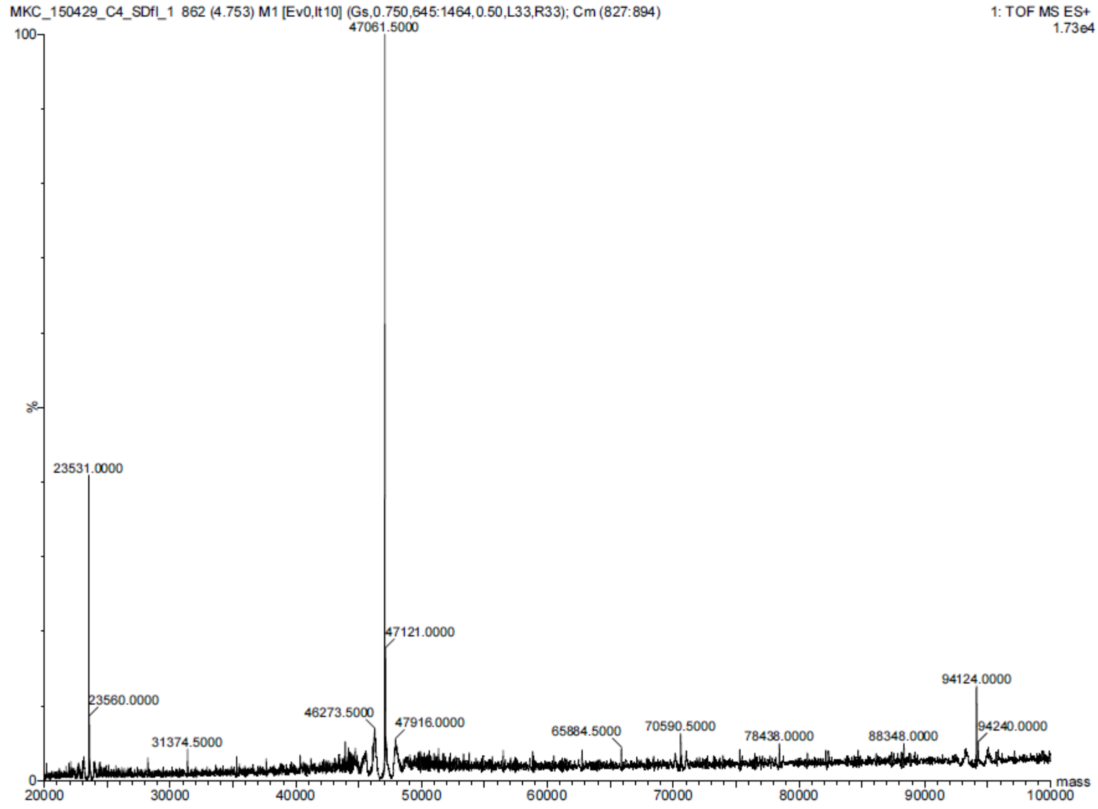

B) The product was identified as TNRC6C:

1. Q9HCJ0 **Mass:** 176427 **Score:** 4811 **Matches:** 111(110) **Sequences:** 74(74) **emPAI:** 4.07

Trinucleotide repeat-containing gene 6C protein OS=Homo sapiens GN=TNRC6C PE=1 SV=3

| Query | Observed | Mr(expt) | Mr(calc) | ppm | Miss | Score | Expect | Rank | Unique | Peptide |
|---|---|---|---|---|---|---|---|---|---|---|
| 14 | 361.2020 | 720.3894 | 720.3919 | -3.45 | 0 | 50 | 0.0043 | 1 | U | K.HGAIPGGL.S |

C) The product contains peptides only from TNRC6C fragment 1260-1690, that is, SDfl:

| | | | | | |
|---|---|---|---|---|---|
| **1101** | LNSSQPSLRA | QVPQFLSPQV | QAQLLQFAAK | NIGLNPALLT | SPINPQHMTM |
| **1151** | LNQLYQLQLA | YQRLQIQQQM | LQAQRNVSGS | MRQQEQQVAR | TITNLQQQIQ |
| **1201** | QHQRQLAQAL | LVKQPPPPPP | PPHLSLHPSA | GKSAMDSFPS | HPQTPGLPDL |
| **1251** | QTKEQQSSPN | TFAPYPLAGL | NPNMNVNSMD | MTGGLSVKDP | SQSQSRLPQW |
| **1301** | THPNSMDNLP | SAASPLEQNP | SKHGAIPGGL | SIGPPGKSSI | DDSYGRYDLI |
| **1351** | QNSESPASPP | VAVPHSWSRA | KSDSDKISNG | SSINWPPEFH | PGVPWKGLQN |
| **1401** | IDPENDPDVT | PGSVPTGPTI | NTTIQDVNRY | LLKSGGKLSD | IKSTWSSGPT |
| **1451** | SHTQASLSHE | LWKVPRNSTA | PTRPPPGLTN | PKPSSTWGAS | PLGWTSSYSS |
| **1501** | GSAWSTDTSG | RTSSWLVLRN | LTPQIDGSTL | RTLCLQHGPL | ITFHLNLTQG |
| **1551** | NAVVRYSSKE | EAAKAQKSLH | MCVLGNTTIL | AEFAGEEEVN | RFLAQGQALP |
| **1601** | PTSSWQSSSA | SSQPRLSAAG | SSHGLVRSDA | GHWNAPCLGG | KGSSELLWGG |
| **1651** | VPQYSSSLWG | PPSADDSRVI | GSPTPLTTLL | PGDLLSGESL | |

**Supplementary figure S10 to Fig 2g. Negative control experiments probing the interaction of GW182 SD with the tristetraprolin-binding site of CNOT1. (a)** Representative normalized FCS autocorrelation curve from one of two independent titrations of fluorescently labeled (Cy5) α-chymotrypsinogen at 1 µM in the absence (black line) and presence of 15.6 µM GW182 SD10 (grey line), shown with fits to the experimental data (translucent points) and the corresponding raw residuals (bottom panel). **(b)** Comparison of diffusion times (τ) obtained in two independent titration experiments. Increasing concentrations of GW182 SD10 did not produce significant changes in the diffusion time of α-chymotrypsinogen, indicating a lack of crowding agent and viscosity related scaffolds.

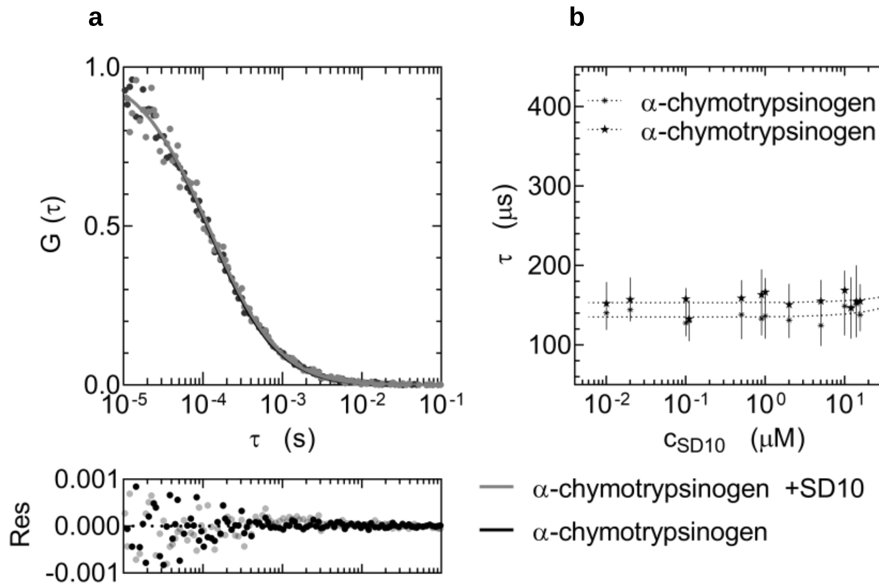

**Supplementary figure S11 to Fig 3a - Fig 5. (a)** Absorbance spectra of human serum albumin (HSA, black), GST-CNOT1 EY mutant (800-999; green), GST-CNOT1 wt (800-999; lime), and **(b)** fluorescein-labelled TTP peptide (purple), and rhodamine-labelled GW182 peptide (red). **(c)** Absorbance spectra of fluorescently labelled proteins: GST-CNOT1 wt-AF488 (lime), GST-CNOT1 EY-AF488 (green), GST-AF488 (grey) and **(d)** GW182 SD-mCherry (red), GW182 SD-Alexa 647 (orange). Stoichiometry of free AF488 and Alexa 647 is shown in Supplementary Fig. S12.

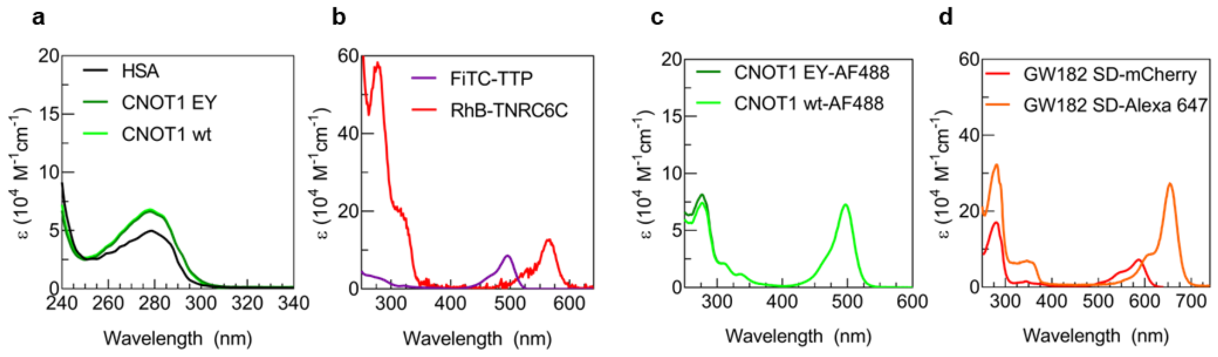

**Supplementary figure S12 to Fig 3a. Tristetraprolin (TTP) and intrinsically disordered CIM1 region of GW182 SD displacement on CNOT1 wt (800-999) of CCR4-NOT. (a)** (*left*) FCS autocorrelation curves of GST-CNOT1 wt-AF488, GST-CNOT1 EY-AF488, and GST-AF488 and *(right)* estimated from autocorrelation curves proportion (%) of AF488-labeled proteins. **(c)** Size-exclusion chromatography purification of GW182 SD-Alexa 647 from unlabelled Alexa 647 dye. **(d)** FCS autocorrelation curves from negative control displacement experiments showing TTP (blue) and GW182 SD CIM1 (red) in the presence of HSA (white, open circles). **(e)** FCS autocorrelation curves showing displacement of fluorescein-labelled TTP (purple) by 50 μM unlabelled TTP (blue) in the presence of CNOT1 wt (lime), **(f)** CNOT1 EY (green), or **(g)** HSA (white). **(h)** Diffusion times (τ) indicating complex formation between increasing concentrations of FITC-TTP (purple) or unlabelled TTP (blue) with CNOT1 wt (lime), CNOT1 EY (green), and HSA (white).

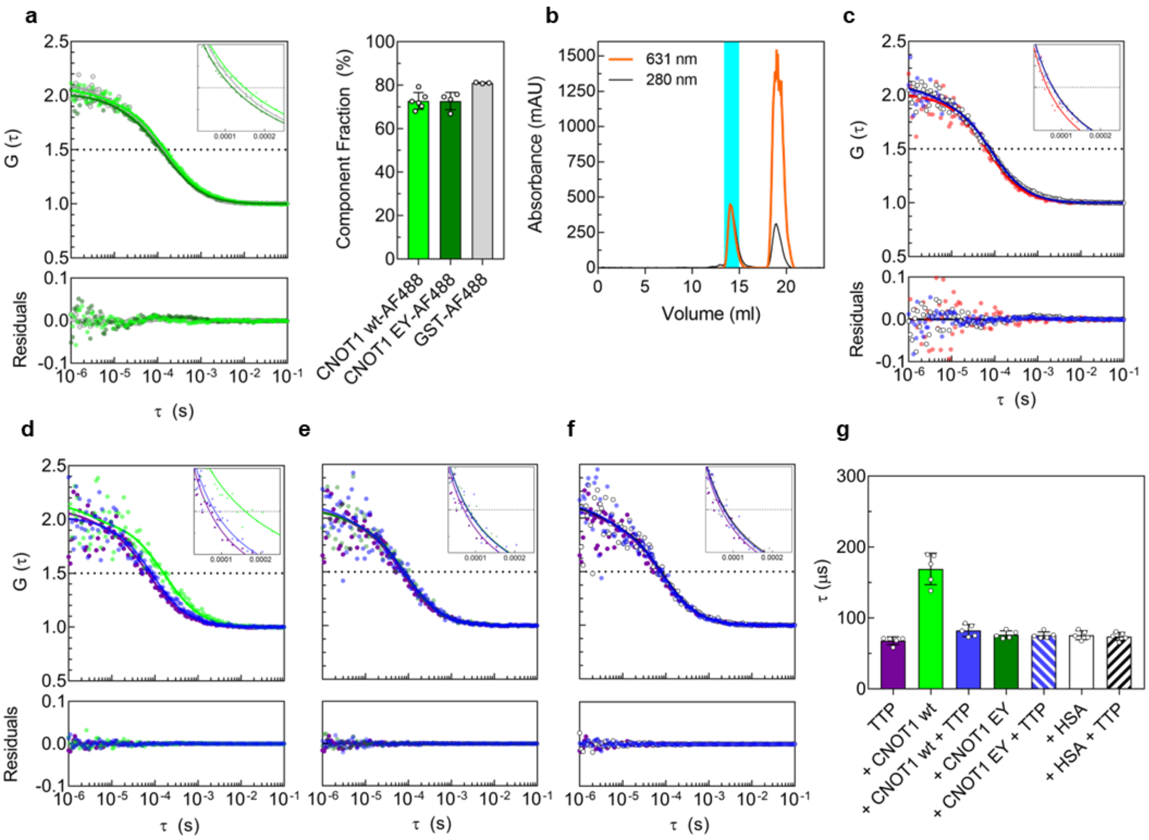

**Supplementary figure S13 to Fig 4. LLPS of GW182 SD and formation of multiprotein droplets. (a)** Confocal images showing lack of propensity for phase separation of mCherry and GST-CNOT1 EY (800-999) mutant in 10% (w/v) PEG 1k. **(b)** Representative confocal images showing LLPS of GW182 SD-mCherry (6 µM) in 10% (w/v) PEG 1k in the presence of equimolar concentrations (6 µM) of GST-CNOT1 wt (800-999)-AF488, GST-CNOT1 EY (800-999)-AF488, GST-AF488, or GST-CNOT1 wt (800-999)-AF488 in the presence of excess TTP peptide (5 µM). Images acquired after 24 h, n = 2-5. Scale bar: 5 µm. **(c)** Colocalization fluorograms showing AF488 and mCherry intensity distributions within the dense phase of GW182 SD-mCherry droplets formed with either, GST-AF488 or TTP peptide alone, or in multi-protein droplets formed with GST-CNOT1 wt, GST-CNOT1 EY or GST-CNOT1 wt with TTP peptide. **(d)** Violin plots showing the distribution of liquid droplet counts (N) formed by (6 µM) GW182 SD in the presence of: GST-CNOT1 wt-AF488 (6 µM, white), GW182 SD-mCherry alone (6 µM, red), and GW182 SD-mCherry (6 µM) in the presence of GST-CNOT1 wt-AF488 (6 µM, lime), GST-CNOT1 EY-AF488 (6 µM, green), GST-AF488 (6 µM, grey), TTP peptide (5 µM, blue), or GST-CNOT1 wt-AF488 (6 µM) and TTP peptide (5 µM, yellow). Medians are shown as open circles; boxes represent interquartile ranges ($Q_1$-$Q_3$) with whiskers; grey dots represent individual droplets. Images were acquired after 24 h, n = 2-5.

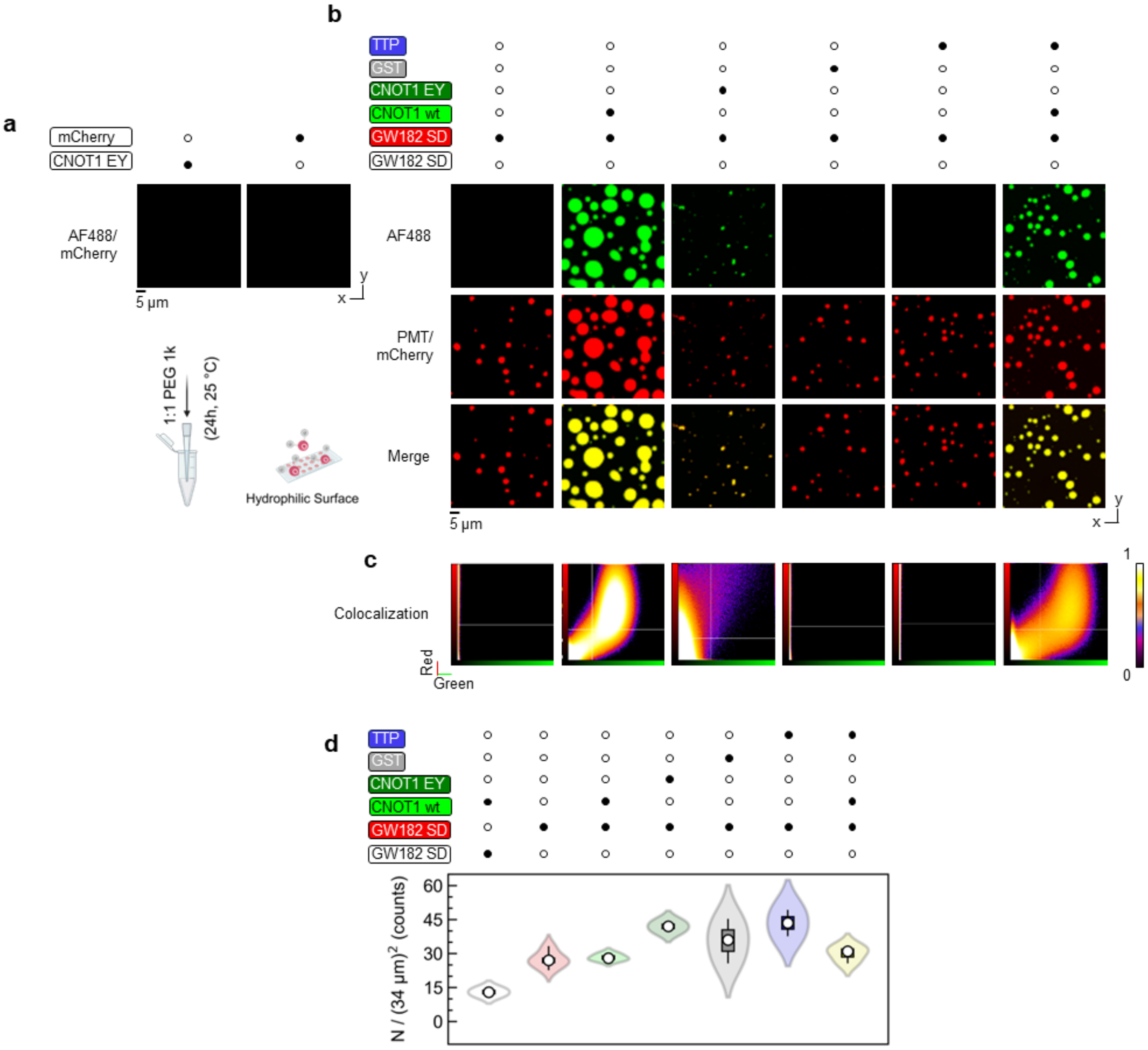

**Supplementary figure S14 to Fig 4g. LLPS of GW182 SD and formation of multiprotein droplets with CNOT1 wt (800-999) after 12h. (a)** Confocal images showing formation of multiprotein liquid droplets by GW182 SD-mCherry (6 µM) in 10% (w/v) PEG 1k in the presence of increasing concentrations (0-7 µM) of GST-CNOT1 wt (800-999)-AF488. Images were acquired after 12 h; n = 2-5. Scale bar: 5 µm. **(b)** Colocalization fluorograms showing AF488 and mCherry intensity distributions within the dense phase of GW182 SD-mCherry droplets formed with GST-CNOT1 wt-AF488. **(c)** Representative confocal images showing Delaunay triangulation (in red) performed between geometric centres of mass of droplets. Triangulation performed on n = 2-5 confocal images. Scale bar: 5 µm. **(d)** Droplet-droplet distances estimated from Delaunay triangulations, showing median distances (red) and minimal distances (blue). **(e-f)** Normalized histograms of integrated fluorescence intensities from AF488 and mCherry in the presence of increasing concentrations of CNOT1 wt (e, lime) and CNOT1 EY (f, green). **(g)** Colocalization (%) between AF488 and mCherry fluorescence intensities across increasing concentrations of CNOT1 wt (lime) and CNOT1 EY (green). **(h-j)** Violin plots showing distributions of droplet parameters in the absence (red) and presence of GST-CNOT1 wt-AF488: **(h)** diameters (φ), **(i)** droplet counts (*N*), **(j)** total droplet area occupied. Medians are indicated by open circles; boxes represent interquartile ranges ($Q_1$-$Q_3$) with whiskers; grey dots represent individual droplets. Images acquired after 12 h; n = 2-5.

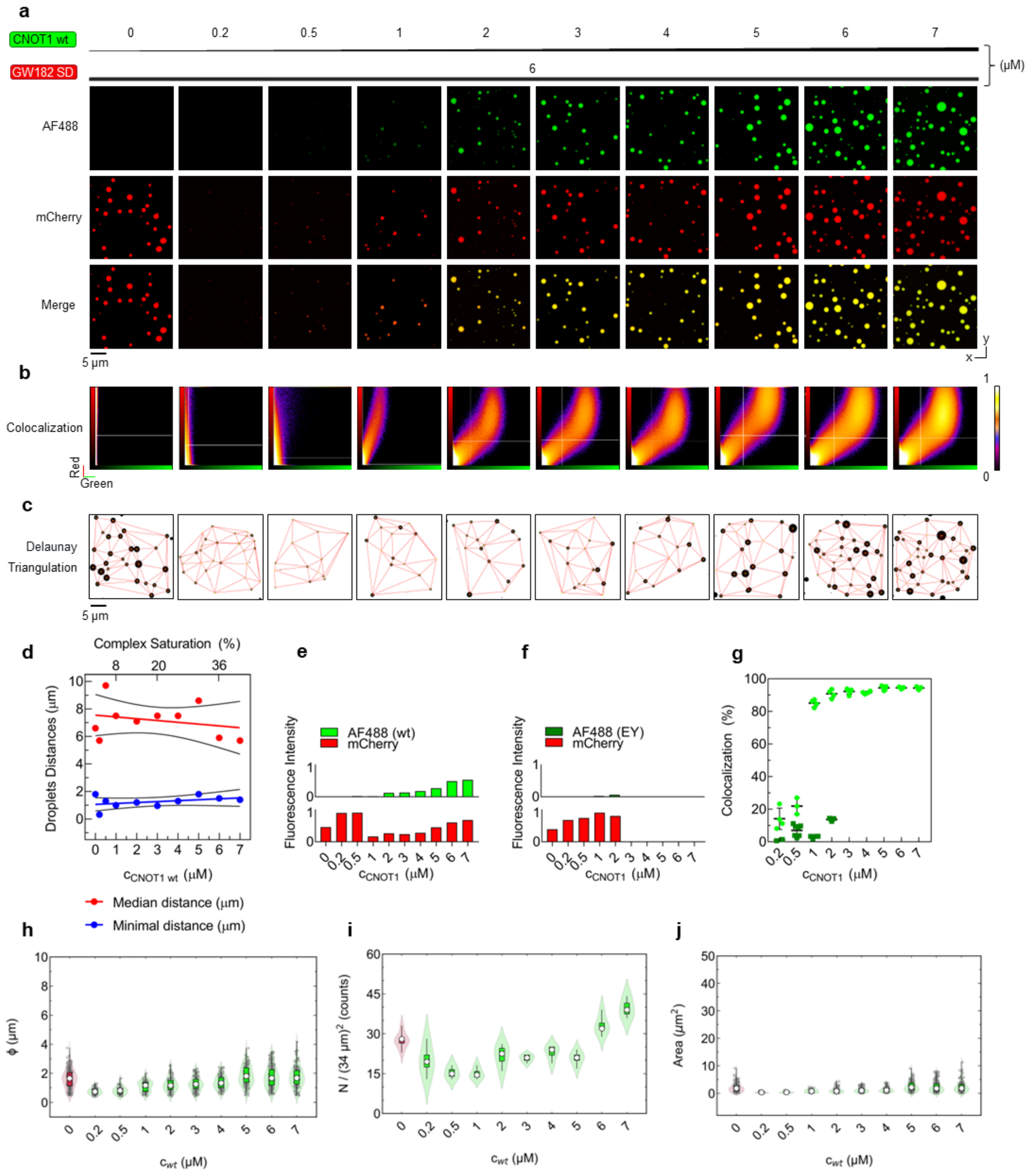

**Supplementary figure S15 to Fig 4g. LLPS of GW182 SD and formation of multiprotein droplets with CNOT1 wt (800-999) inhibited by tristetraproline (TTP) after 12h. (a)** Confocal images showing the formation of multiprotein liquid droplets with equimolar concentrations of GW182 SD-mCherry (6 µM) and GST-CNOT1 wt (800-999)-AF488 (6 µM) in 10% (w/v) PEG 1k, in the presence of increasing concentrations (0-50 µM) of TTP peptide. Images were acquired after 12 h; n = 2-5. Scale bar: 5 µm. **(b)** Colocalization fluorograms showing AF488 and mCherry intensity distributions within the dense phase of GW182 SD-mCherry droplets formed with GST-CNOT1 wt (800-999)-AF488 across increasing TTP peptide concentrations. **(c)** Normalized histograms of integrated fluorescence intensities from AF488 (lime) and mCherry (red) in the presence of increasing TTP peptide concentrations. **(d)** Colocalization (%) between AF488 and mCherry fluorescence intensities across increasing of TTP peptide concentrations (blue). **(e-g)** Violin plots showing distributions of droplet parameters in the absence (lime) and presence of TTP peptide (blue): **(c)** diameters (φ), **(d)** droplet counts (*N*), **(e)** total droplet area occupied. Medians are indicated by open circles; boxes represent interquartile ranges ($Q_1$-$Q_3$) with whiskers; grey dots represent individual droplets. Images acquired after 12 h; n = 2-5.

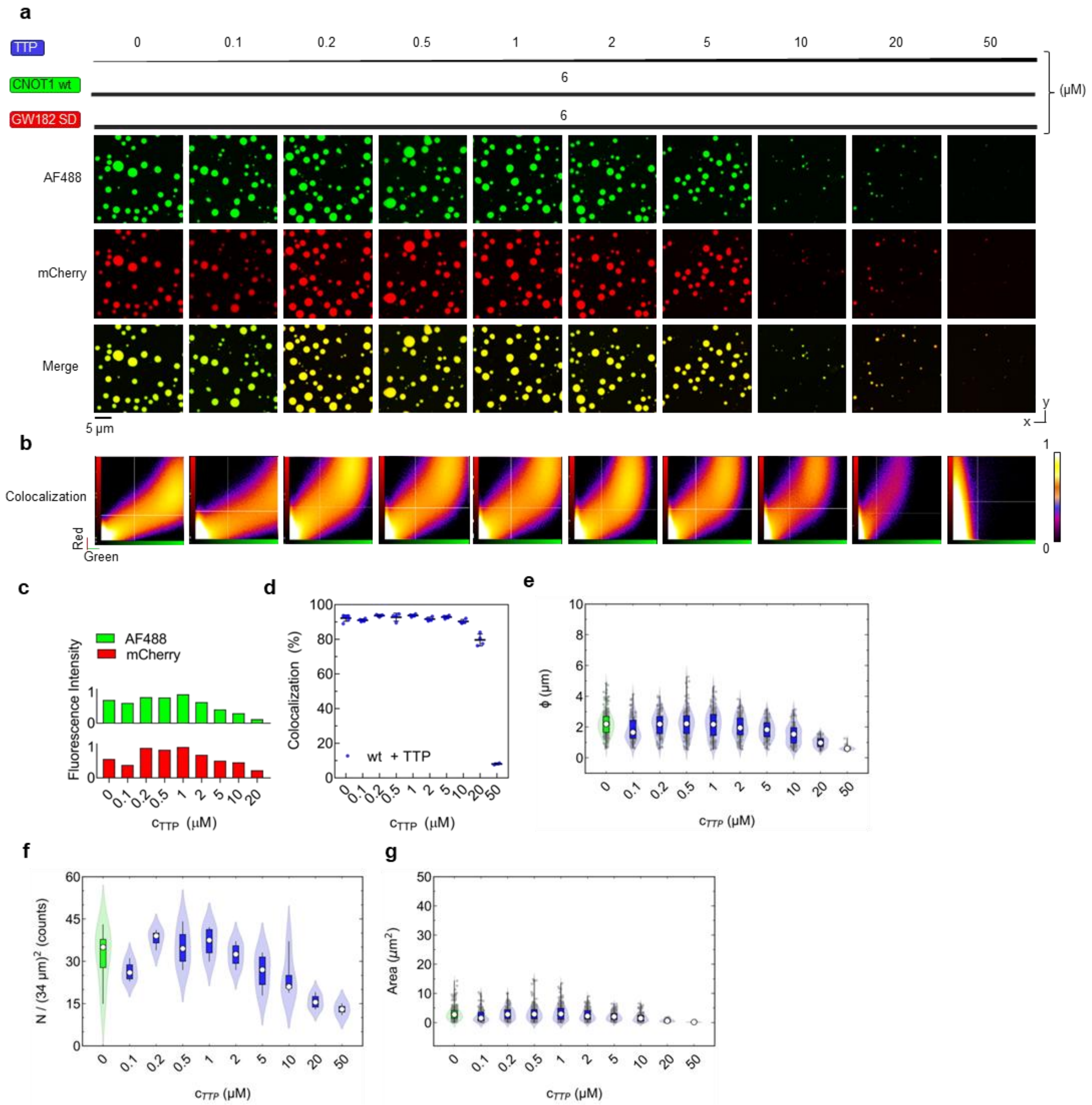

**Supplementary figure S16 to Fig 4g. LLPS of GW182 SD and formation of multiprotein droplets with CNOT1 wt (800-999) after 24h, n = 2-5. (a)** Confocal images showing formation of multiprotein liquid droplets by GW182 SD-mCherry (6 μM) in 10% (w/v) PEG 1k in the presence of increasing concentrations (0-7 μM) of GST-CNOT1 wt (800-999)-AF488. Images were acquired after 24 h; n = 2-5. Scale bar: 5 μm. **(b)** Colocalization fluorograms showing AF488 and mCherry intensity distributions within the dense phase of GW182 SD-mCherry droplets formed with GST-CNOT1 wt-AF488. **(c)** Representative confocal images showing Delaunay triangulation (in red) performed between geometric centres of mass of droplets. Triangulation performed on n = 2-5 confocal images. Scale bar: 5 μm. **(d)** Droplet-droplet distances estimated from Delaunay triangulations, showing median distances (red) and minimal distances (blue). **(e-f)** Normalized histograms of integrated fluorescence intensities from AF488 and mCherry in the presence of increasing concentrations of CNOT1 wt (e, lime) and CNOT1 EY (f, green). **(g)** Colocalization (%) between AF488 and mCherry fluorescence intensities across increasing concentrations of CNOT1 wt (lime) and CNOT1 EY (green). **(h-j)** Violin plots showing distributions of droplet parameters in the absence (red) and presence of GST-CNOT1 wt-AF488: **(h)** diameters (φ), **(i)** droplet counts (*N*), **(j)** total droplet area occupied. Medians are indicated by open circles; boxes represent interquartile ranges ($Q_1$-$Q_3$) with whiskers; grey dots represent individual droplets.

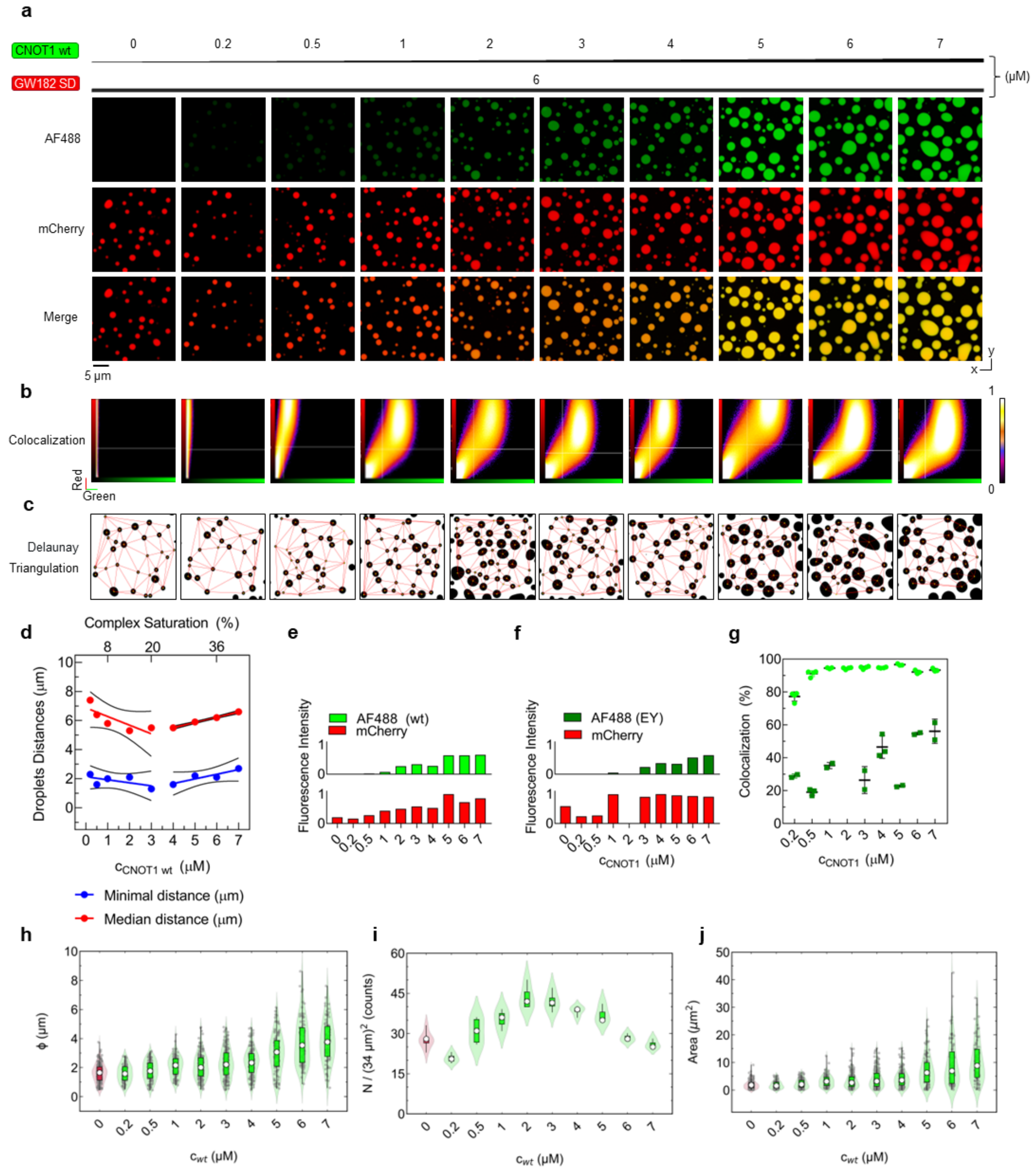

**Supplementary figure S17 to Fig 4g. LLPS of GW182 SD and formation of multiprotein droplets with CNOT1 wt (800-999) inhibited by tristetraproline (TTP) after 24h. (a)** Confocal images showing the formation of multiprotein liquid droplets with equimolar concentrations of GW182 SD-mCherry (6 µM) and GST-CNOT1 wt (800-999)-AF488 (6 µM) in 10% (w/v) PEG 1k, in the presence of increasing concentrations (0-50 µM) of TTP peptide. Images were acquired after 24 h; n = 2-5. Scale bar: 5 µm. **(b)** Colocalization fluorograms showing AF488 and mCherry intensity distributions within the dense phase of GW182 SD-mCherry droplets formed with GST-CNOT1 wt (800-999)-AF488 across increasing TTP peptide concentrations. **(c)** Normalized histograms of integrated fluorescence intensities from AF488 (lime) and mCherry (red) in the presence of increasing TTP peptide concentrations. **(d)** Colocalization (%) between AF488 and mCherry fluorescence intensities across increasing of TTP peptide concentrations (blue). **(e-g)** Violin plots showing distributions of droplet parameters in the absence (lime) and presence of TTP peptide (blue): **(c)** diameters (φ), **(d)** droplet counts (*N*), **(e)** total droplet area occupied. Medians are indicated by open circles; boxes represent interquartile ranges ($Q_1$-$Q_3$) with whiskers; grey dots represent individual droplets. Images acquired after 24 h; n = 2-5.

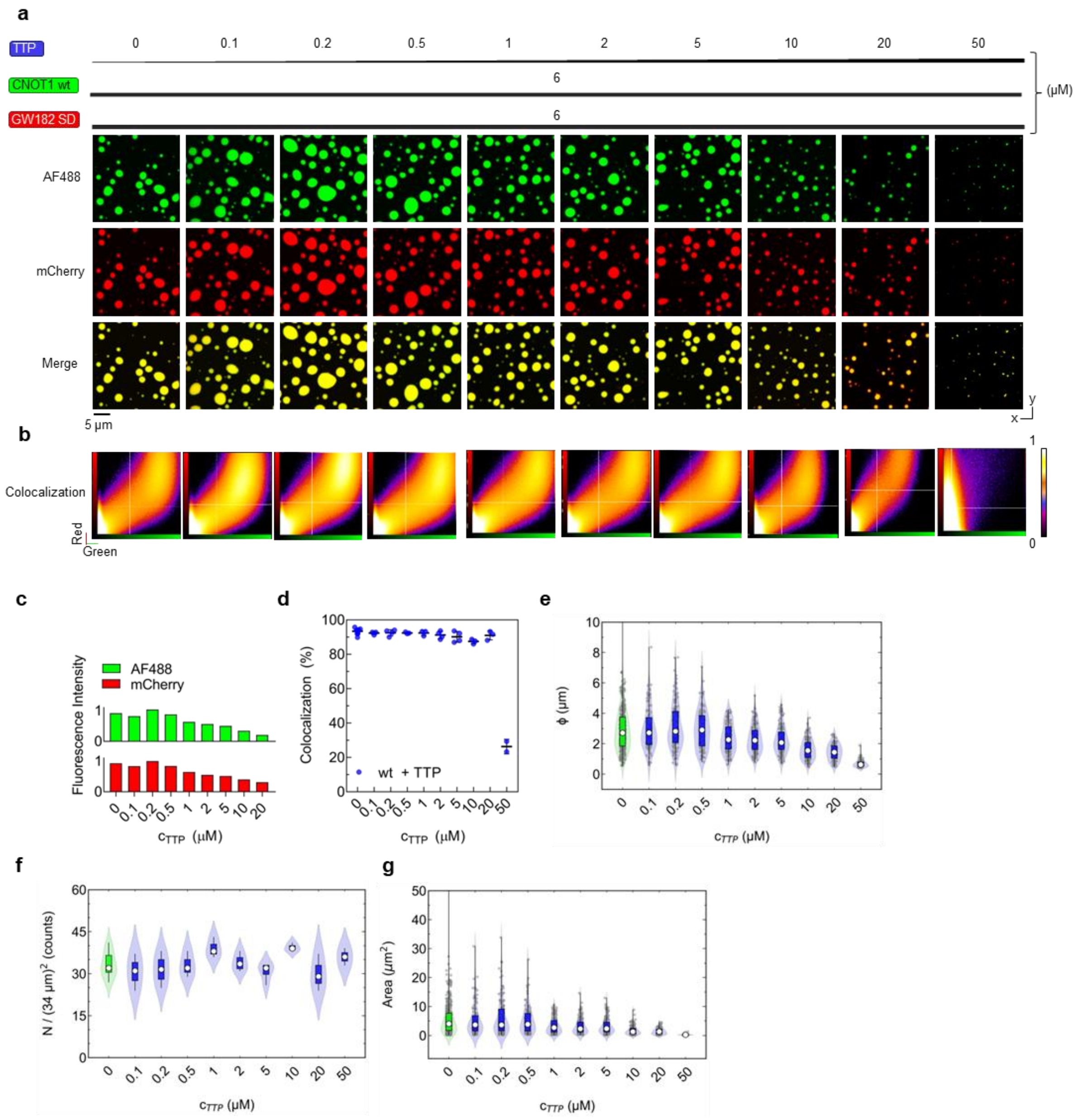

**Supplementary figure S18 to Fig 4i. LLPS of GW182 SD and formation of multiprotein droplets with CNOT1 EY (800-999) after 12-24h.** **(a)** Confocal images showing formation of multiprotein liquid droplets by GW182 SD-mCherry (6 μM) in 10% (w/v) PEG 1k in the presence of increasing concentrations (0-2 μM) of GST-CNOT1 EY (800-999)-AF488. Images were acquired after 12 h; n = 2-5. Scale bar: 5 μm. **(b)** Colocalization fluorograms showing AF488 and mCherry intensity distributions within the dense phase of GW182 SD-mCherry droplets formed with GST-CNOT1 EY-AF488. **(c-e)** Violin plots showing distributions of droplet parameters in the absence (red) and presence of GST-CNOT1 EY-AF488 after 12 h: **(c)** diameters (φ), **(d)** droplet counts (*N*), and **(e)** total droplet area. Medians are indicated by open circles; boxes represent interquartile ranges ($Q_1$-$Q_3$) with whiskers; grey dots represent individual droplets. Images acquired after 12 h; n = 2-5. **(f)** Confocal images showing formation of multiprotein liquid droplets by GW182 SD-mCherry (6 μM) in 10% (w/v) PEG 1k in the presence of increasing concentrations (0-7 μM) of GST-CNOT1 EY (800-999)-AF488. Images were acquired after 24 h; n = 2-5. Scale bar: 5 μm. **(g)** Colocalization fluorograms showing AF488 and mCherry intensity distributions within the dense phase of GW182 SD-mCherry droplets formed with GST-CNOT1 EY-AF488. **(h-j)** Violin plots showing distributions of droplet parameters in the absence (red) and presence of GST-CNOT1 EY-AF488: **(h)** diameters (φ), **(i)** droplet counts (*N*), **(j)** total droplet area occupied. Medians are indicated by open circles; boxes represent interquartile ranges ($Q_1$-$Q_3$) with whiskers; grey dots represent individual droplets. Images acquired after 12 h; n = 2-5.

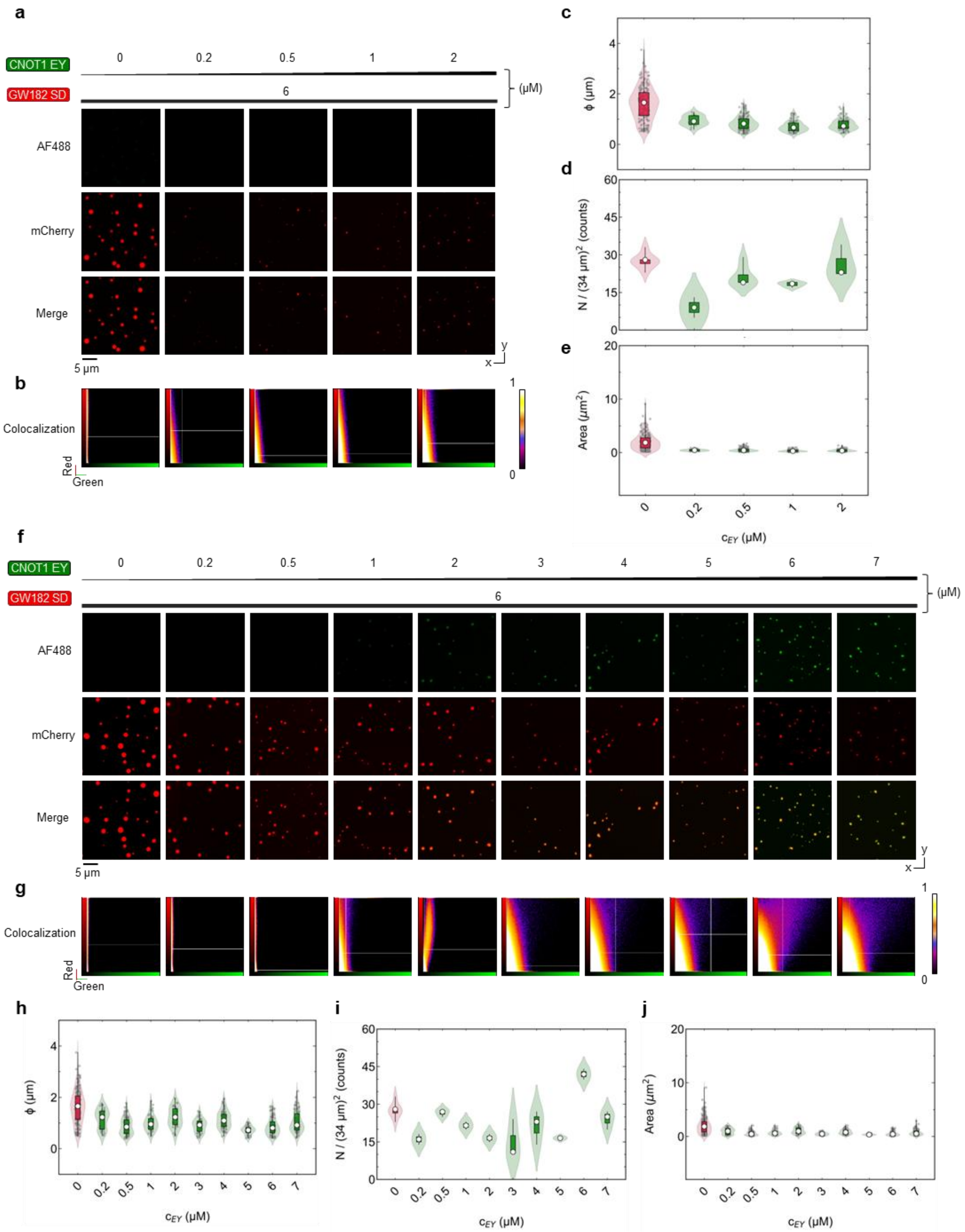
a
CNOT1 EY
GW182 SD
0
0.2
0.5
1
2
6
(µM)
AF488
mCherry
Merge
5 µm
y
x
b
Colocalization
Red
Green
1
0
c
ϕ (µm)
d
N / (34 µm)² (counts)
e
Area (µm²)
$c_{EY}$ (µM)
f
CNOT1 EY
GW182 SD
0
0.2
0.5
1
2
3
4
5
6
7
(µM)
AF488
mCherry
Merge
5 µm
g
Colocalization
Red
Green
h
ϕ (µm)
$c_{EY}$ (µM)
i
N / (34 µm)² (counts)
$c_{EY}$ (µM)
j
Area (µm²)
$c_{EY}$ (µM)

**Supplementary figure S19 to Fig 5b-c. Crystal structure of CNOT1 (820-999), Alpha Fold 2 prediction of GW182 SD (a)** Putative conformation of GW182 SD predicted by AlphaFold2. Cysteine residues are highlighted in orange, with solvent accessible Cys378 marked by an orange star. RRM domain is shown as a ribbon with the solvent-accessible surface displayed. **(b)** Crystal structure of CNOT1 (820-999) (PDB: 4J8S) with a docked CIM1 peptide of GW182 SD (red). Lysine residues available for labelling with AF488 are marked with lime stars.

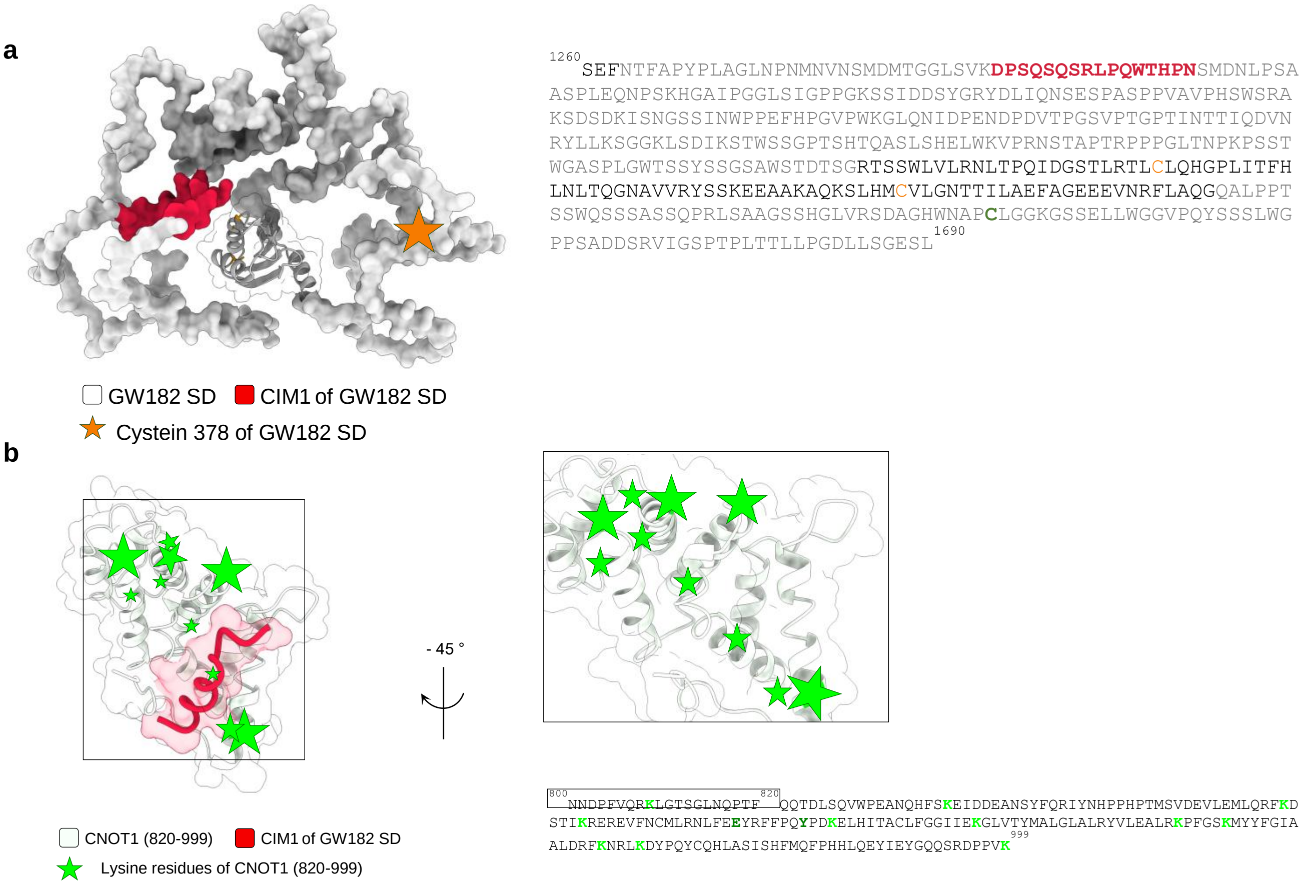

**Supplementary figure S20 to Fig 5c. Comparison of LLPS of GW182 SD, GW182 SD-Alexa 647 and GW182 SD-mCherry and formation of multiprotein droplets with GST-CNOT1 wt (800-999) after 12-24h. (a)** Confocal images showing formation of multiprotein liquid droplets by GW182 SD (6 µM) in the presence of GST-CNOT1 wt (800-999)-AF488 (6 µM). Images acquired after 12 h; n = 2-3. Scale bar: 5 µm. **(b)** Confocal images showing LLPS of GW182 SD-Alexa 647 (6 µM) in 10% (w/v) PEG 1k and formation of multiprotein liquid droplets in the presence of GST-CNOT1 wt (800-999)-AF488 (6 µM). Images acquired after 12 h; n = 2-5. Scale bar: 5 µm. **(c)** Colocalization fluorograms showing AF488 and Alexa 647 intensity distributions within the dense phase of GW182 SD-Alexa 647 droplets formed with GST-CNOT1 wt-AF488**.** **(d-f)** Violin plots showing distributions of droplet parameters after 12 h for: GW182 SD with GST-CNOT1 wt-AF488 (white), GW182 SD-mCherry alone (red), GW182 SD-mCherry with GST-CNOT1 wt-AF488 (lime), GW182 SD-Alexa 647 alone (orange), and GW182 SD-Alexa 647 with GST-CNOT1 wt-AF488 (yellow): **(d)** diameters (φ), **(e)** droplet counts (*N*), **(f)** total droplet area occupied. Medians are indicated by open circles; boxes represent interquartile ranges ($Q_1$-$Q_3$) with whiskers; grey dots represent individual droplets. Images acquired after 12 h; n = 2-5. **(g-h)** Normalized histograms of integrated fluorescence intensities from: **(g)** AF488 and mCherry or **(h)** AF488 and Alexa 647 in the presence of CNOT1 wt. **(i)** Colocalization (%) between AF488 and mCherry (lime) or Alexa 647 (orange) fluorescence intensities, in the presence of CNOT1 wt (lime) after 12h.**(j)** Confocal images showing multiprotein liquid droplet formation by GW182 SD (6 µM) in the presence of GST-CNOT1 wt (800-999)-AF488 (6 µM). Images acquired after 21 h; n = 2-3. Scale bar: 5 µm. **(k)** Confocal images showing LLPS of GW182 SD-Alexa 647 (6 µM) in 10% (w/v) PEG 1k and formation of multiprotein liquid droplets in the presence of GST-CNOT1 wt (800-999)-AF488 (6 µM). Images acquired after 24 h; n = 2-5. Scale bar: 5 µm.**(l)** Colocalization fluorograms showing AF488 and Alexa 647 intensity distributions within the dense phase of GW182 SD-Alexa 647 droplets formed with GST-CNOT1 wt-AF488 after 24 h. **(m-o)** Violin plots showing distributions of droplet parameters after 24 h for: GW182 SD with GST-CNOT1 wt-AF488 (white), GW182 SD-mCherry (red), GW182 SD-mCherry with GST-CNOT1 wt-AF488 (lime), GW182 SD-Alexa 647 (orange), and GW182 SD-Alexa 647 with GST-CNOT1 wt-AF488 **(d)** diameters (φ), **(e)** droplet counts (*N*), **(f)** total droplet area occupied. Medians are indicated by open circles; boxes represent interquartile ranges ($Q_1$-$Q_3$) with whiskers; grey dots represent individual droplets. Images acquired after 24 h; n = 2-5. **(p-r)** Normalized histograms of integrated fluorescence intensities from: **(p)** AF488 and mCherry or **(r)** AF488 and Alexa 647 in the presence of CNOT1 wt. **(s)** Colocalization (%) between AF488 and mCherry (lime), or Alexa 647 (orange), in the presence of CNOT1 wt after 24 h.

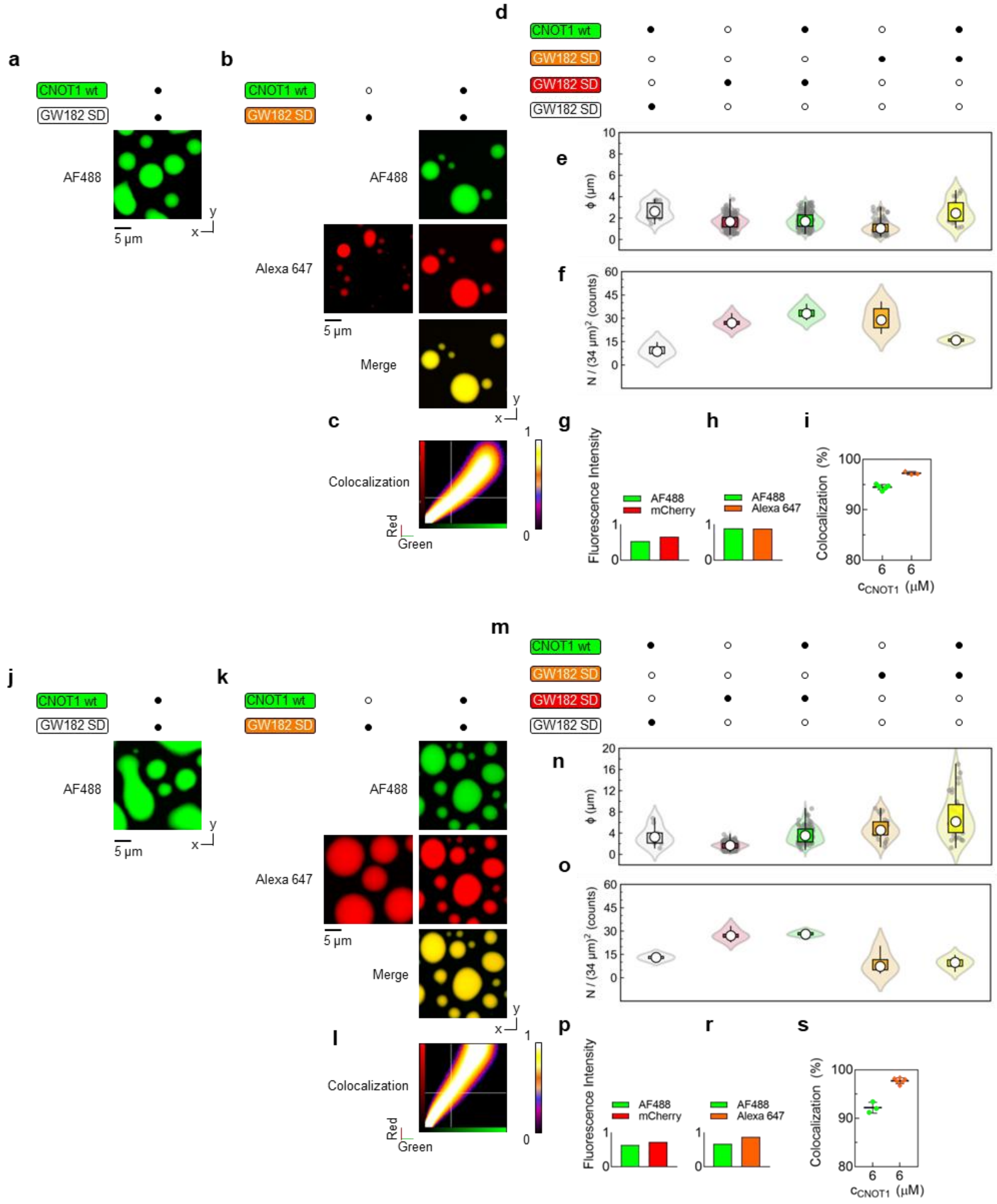

**Supplementary figure S21 to Fig 5c. FRET observed for multiprotein droplets composed of GW182 SD-Alexa 647 and GST-CNOT1 wt (800-999)-AF488 after 12-24 h. (a)** Confocal images from single-wavelength excitation (Alexa 647 channel) showing GW182 SD-Alexa 647 (6 µM) fluorescence in the dense phase upon interaction with GST-CNOT1 wt (800-999)-AF488 (6 µM). Images acquired after 12 h; n = 2-5. Scale bar: 5 µm. **(b)** Confocal images acquired using single-laser excitation (AF488 channel), showing fluorescence from GST-CNOT1 wt-AF488 in unbound and bound states, and FRET-derived Alexa 647 emission indicating complex formation with GW182 SD-Alexa 647 (6 µM) in droplets formed with CNOT1 wt (6 µM). Images acquired after 12 h; n = 2-5. Scale bar: 5 µm. **(c)** Confocal images from single-wavelength excitation showing GW182 SD-Alexa 647 (6 µM) fluorescence in the dense phase and upon interaction with GST-CNOT1 wt-AF488 (6 µM). Images acquired after 24 h; n = 2-5. Scale bar: 5 µm. **(d)** Confocal images from single-laser excitation (AF488 channel) showing GST-CNOT1 wt-AF488 fluorescence and corresponding FRET-derived Alexa 647 signal, confirming complex formation with GW182 SD-Alexa 647 (6 µM) in droplets formed with CNOT1 wt (6 µM). Images acquired after 24 h; n = 2-5. Scale bar: 5 µm. **(e)** Relative FRET intensity comparison after 12 h between equimolar (6 µM) GST-CNOT1 wt-AF488 and either GW182 SD-mCherry (pink) or GW182 SD-Alexa 647 (orange). **(f)** Relative FRET intensity comparison after 24 h between equimolar (6 µM) GST-CNOT1 wt-AF488 and either GW182 SD-mCherry (pink) or GW182 SD-Alexa 647 (orange).

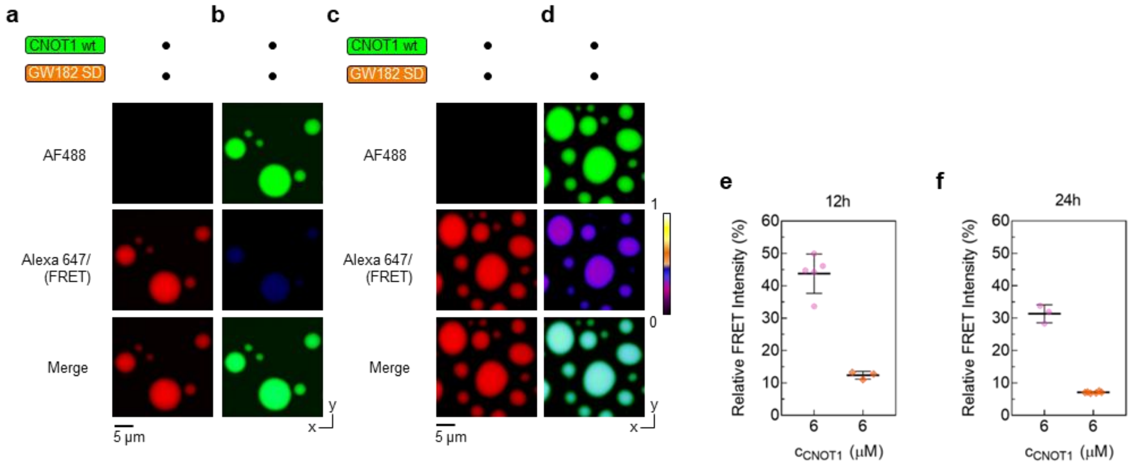

**Supplementary figure S22 to Fig 5g. FRET observed for multiprotein droplets composed of GW182 SD-mCherry and GST-CNOT1 wt (800-999)-AF488 after 12 h. (a)** Confocal images acquired using single-wavelength excitation (mCherry channel), showing GW182 SD-mCherry (6 µM) fluorescence in the dense phase upon interaction with increasing concentrations (0-7 µM) of GST-CNOT1 wt (800-999)-AF488. Images acquired after 12 h; n = 2-5. Scale bar: 5 µm. **(b)** Confocal images acquired using single-wavelength excitation (AF488 channel), showing fluorescence from GST-CNOT1 wt-AF488 in both unbound and bound states, and FRET-derived mCherry emission, indicating limited complex formation with GW182 SD-mCherry (6 µM) in droplets formed with increasing concentrations (0-7 µM) of CNOT1 wt. Images acquired after 12 h; n = 2-5. Scale bar: 5 µm. **(c)** Normalized histograms of integrated fluorescence intensities from single-wavelength excitation of mCherry (FRET, pink), and mCherry (red) in the presence of increasing concentrations of CNOT1 wt. **(d)** Normalized histograms of integrated fluorescence intensities from single-wavelength excitation of mCherry (FRET, purple), and mCherry (red) in the presence of increasing concentrations of CNOT1 EY. **(e)** Relative FRET intensity comparison after 12 h between increasing concentrations of either GST-CNOT1 wt-AF488 (pink) or GST-CNOT1 EY-AF488 (purple) and GW182 SD-mCherry (6 µM).

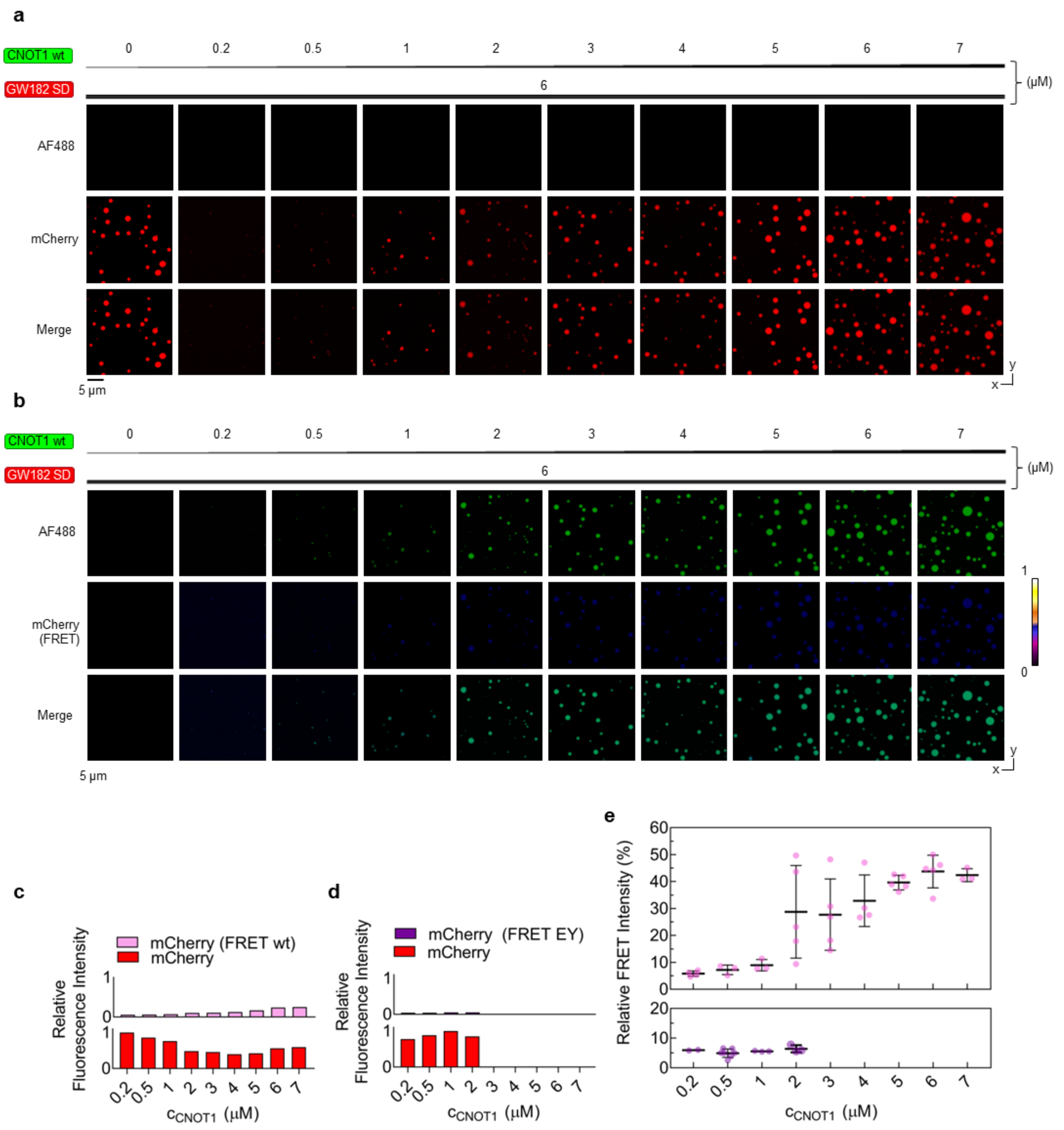

**Supplementary figure S23 to Fig 5h. FRET observed for multiprotein droplets composed of GW182 SD-mCherry and GST-CNOT1 EY (800-999)-AF488 after 12 h. (a)** Confocal images acquired using single-wavelength excitation (mCherry channel), showing GW182 SD-mCherry (6 µM) fluorescence in the dense phase upon interaction with increasing concentrations (0-2 µM) of GST-CNOT1 EY (800-999)-AF488. Images acquired after 12 h; n = 2-5. Scale bar: 5 µm. **(b)** Confocal images acquired using single-laser excitation (AF488 channel), showing fluorescence from GST-CNOT1 EY-AF488 in both unbound and bound states, and weak FRET-derived mCherry emission, indicating limited complex formation with GW182 SD-mCherry (6 µM) in droplets formed with increasing concentrations (0-2 µM) of CNOT1 EY. Images acquired after 12 h; n = 2-5. Scale bar: 5 µm.

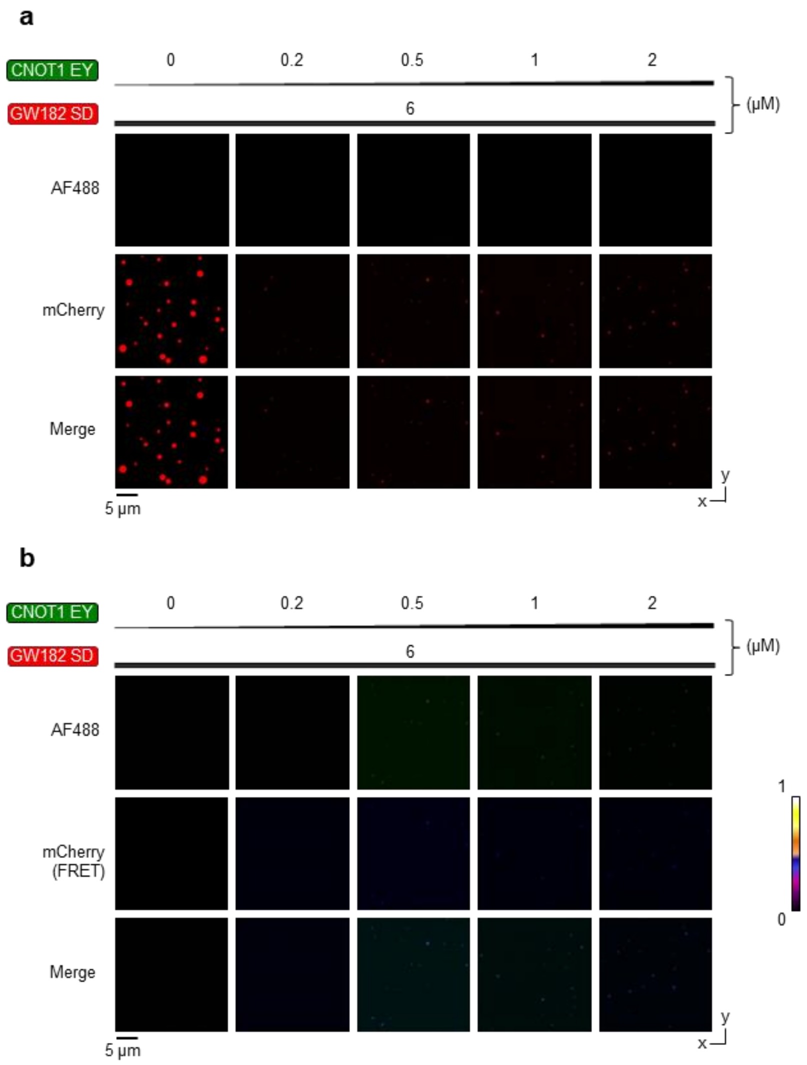

**Supplementary figure S24 to Fig 5i. FRET observed for multiprotein droplets composed of GW182 SD-mCherry and GST-CNOT1 wt (800-999)-AF488 after 24 h. (a)** Confocal images acquired using single-wavelength excitation (mCherry channel), showing GW182 SD-mCherry (6 µM) fluorescence in the dense phase upon interaction with increasing concentrations (0-7 µM) of GST-CNOT1 wt (800-999)-AF488. Images acquired after 24 h; n = 2-5. Scale bar: 5 µm. **(b)** Confocal images acquired using single-wavelength excitation (AF488 channel), showing fluorescence from GST-CNOT1 wt-AF488 in both unbound and bound states, and FRET-derived mCherry emission, indicating limited complex formation with GW182 SD-mCherry (6 µM) in droplets formed with increasing concentrations (0-7 µM) of CNOT1 wt. Images acquired after 24 h; n = 2-5. Scale bar: 5 µm. **(c)** Normalized histograms of integrated fluorescence intensities from single-wavelength excitation of mCherry (FRET, pink), and mCherry (red) in the presence of increasing concentrations of CNOT1 wt. **(d)** Normalized histograms of integrated fluorescence intensities from single-wavelength excitation of mCherry (FRET, purple), and mCherry (red) in the presence of increasing concentrations of CNOT1 EY**.** **(e)** Relative FRET intensity comparison after 24 h between increasing concentrations of either GST-CNOT1 wt-AF488 (pink) or GST-CNOT1 EY-AF488 (purple) and GW182 SD-mCherry (6 µM).

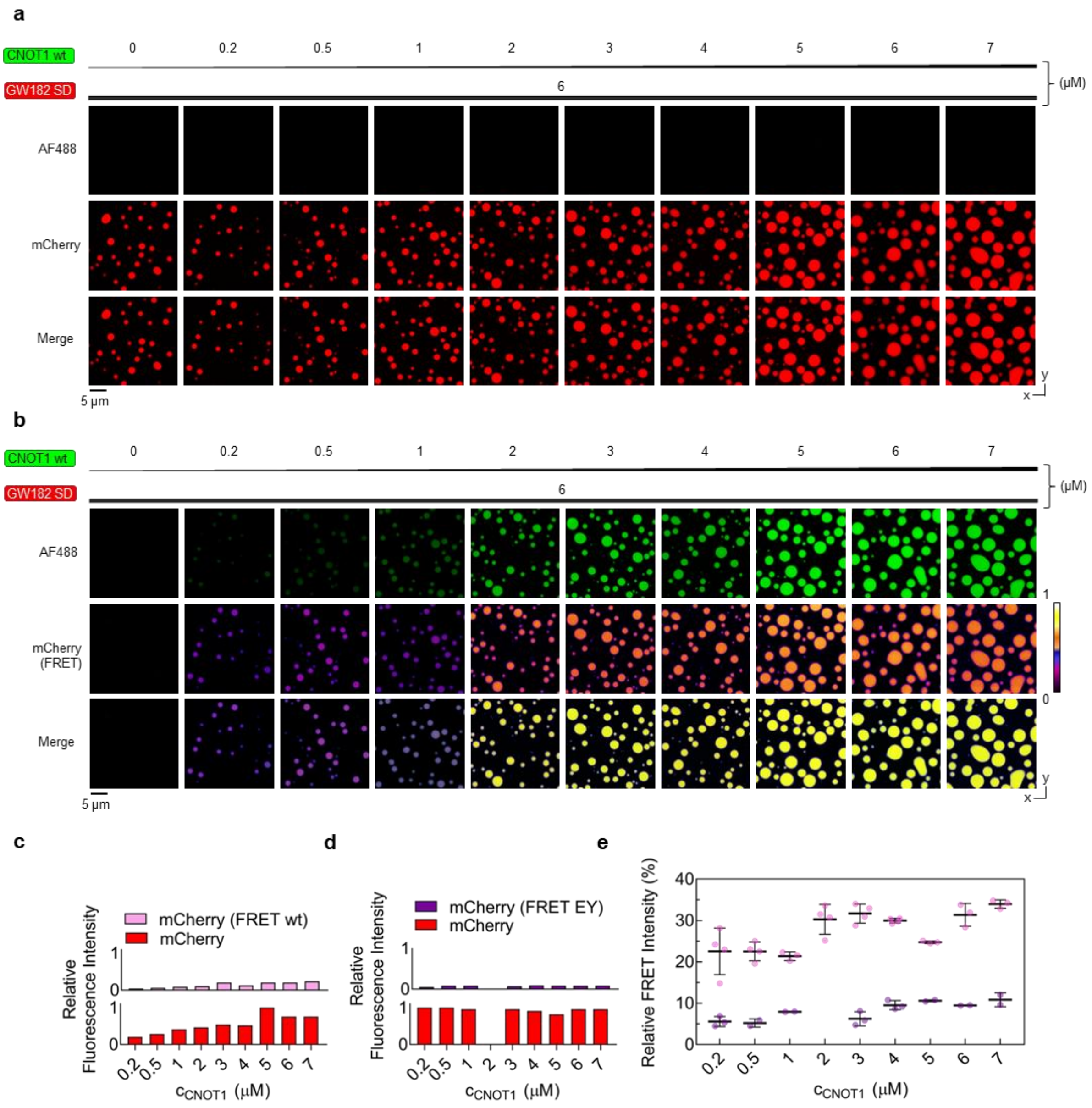

**Supplementary figure S25 to Fig 5j. FRET observed for multiprotein droplets composed of GW182 SD-mCherry and GST-CNOT1 EY (800-999)-AF488 after 24 h. (a)** Confocal images acquired using single-wavelength excitation (mCherry channel), showing GW182 SD-mCherry (6 µM) fluorescence in the dense phase upon interaction with increasing concentrations (0-7 µM) of GST-CNOT1 EY (800-999)-AF488. Images acquired after 24 h; n = 2-5. Scale bar: 5 µm. **(b)** Confocal images acquired using single-laser excitation (AF488 channel), showing fluorescence from GST-CNOT1 EY-AF488 in both unbound and bound states, and weak FRET-derived mCherry emission, indicating limited complex formation with GW182 SD-mCherry (6 µM) in droplets formed with increasing concentrations (0-7 µM) of CNOT1 EY. Images acquired after 24 h; n = 2-5. Scale bar: 5 µm

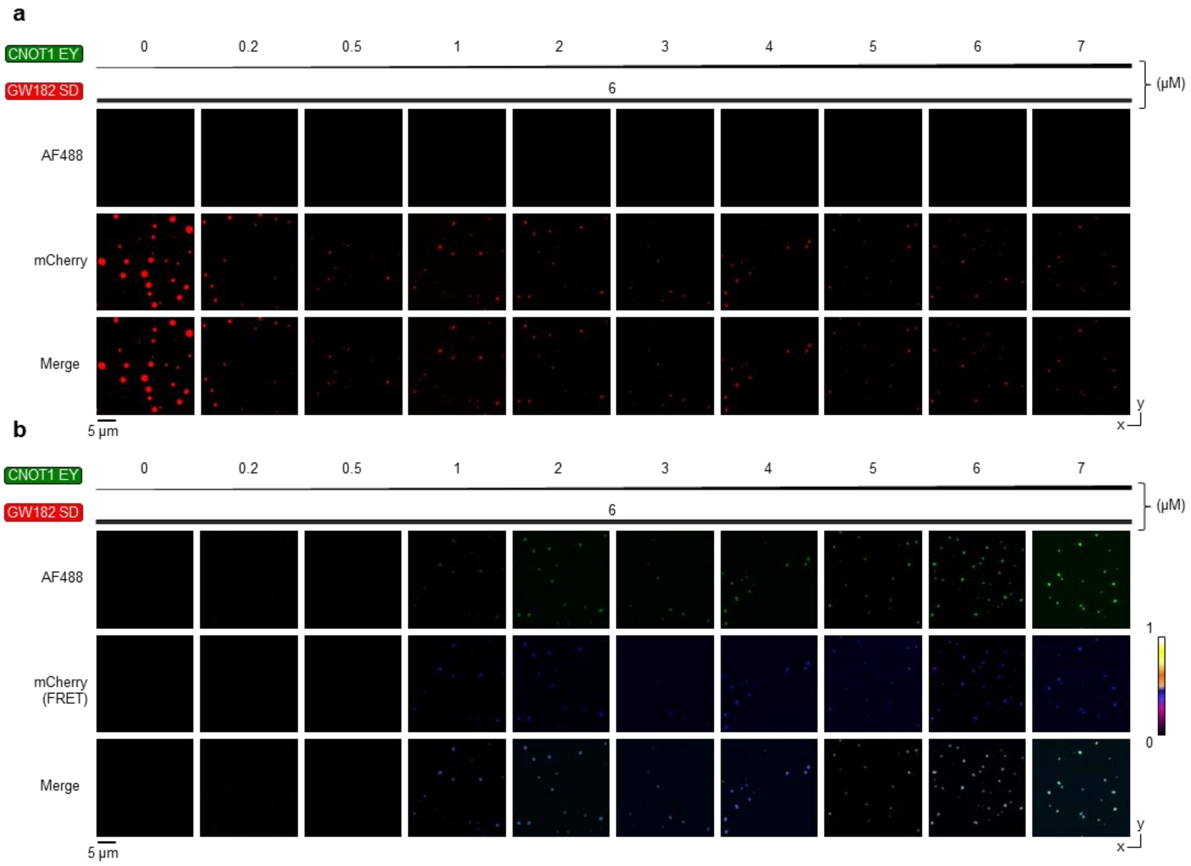

**Supplementary figure S26 to Fig 5k. FRET observed for the inhibition of multiprotein droplets composed of GW182 SD-mCherry and GST-CNOT1 wt (800-999)-AF488 by tristetraprolin (TTP) after 12 h. (a)** Confocal images acquired using single-wavelength excitation (mCherry channel), showing fluorescence of GW182 SD-mCherry (6 µM) in the dense phase upon formation of multiprotein liquid droplets with equimolar concentrations of GST-CNOT1 wt (800-999)-AF488 (6 µM) in 10% (w/v) PEG 1k, in the presence of increasing concentrations (0-50 µM) of TTP peptide. Images acquired after 12 h; n = 2-5. Scale bar: 5 µm. **(b)** Confocal images acquired using single-wavelength excitation (AF488 channel), showing fluorescence from GST-CNOT1 wt-AF488 (6 µM) in both unbound and bound states, and FRET-derived mCherry emission indicating complex formation with GW182 SD-mCherry (6 µM) in droplets formed in the presence of increasing concentrations (0-50 µM) of TTP peptide. Images acquired after 12 h; n = 2-5. Scale bar: 5 µm. **(c)** Normalized histograms of integrated fluorescence intensities from single-wavelength excitation of mCherry (FRET, teal) and mCherry (red) in the presence of increasing concentrations of TTP peptide. **(d)** Relative FRET intensity change after 12 h in liquid droplets formed by equimolar concentrations of GST-CNOT1 wt-AF488 and GW182 SD-mCherry in the presence of increasing concentrations of TTP peptide (teal).

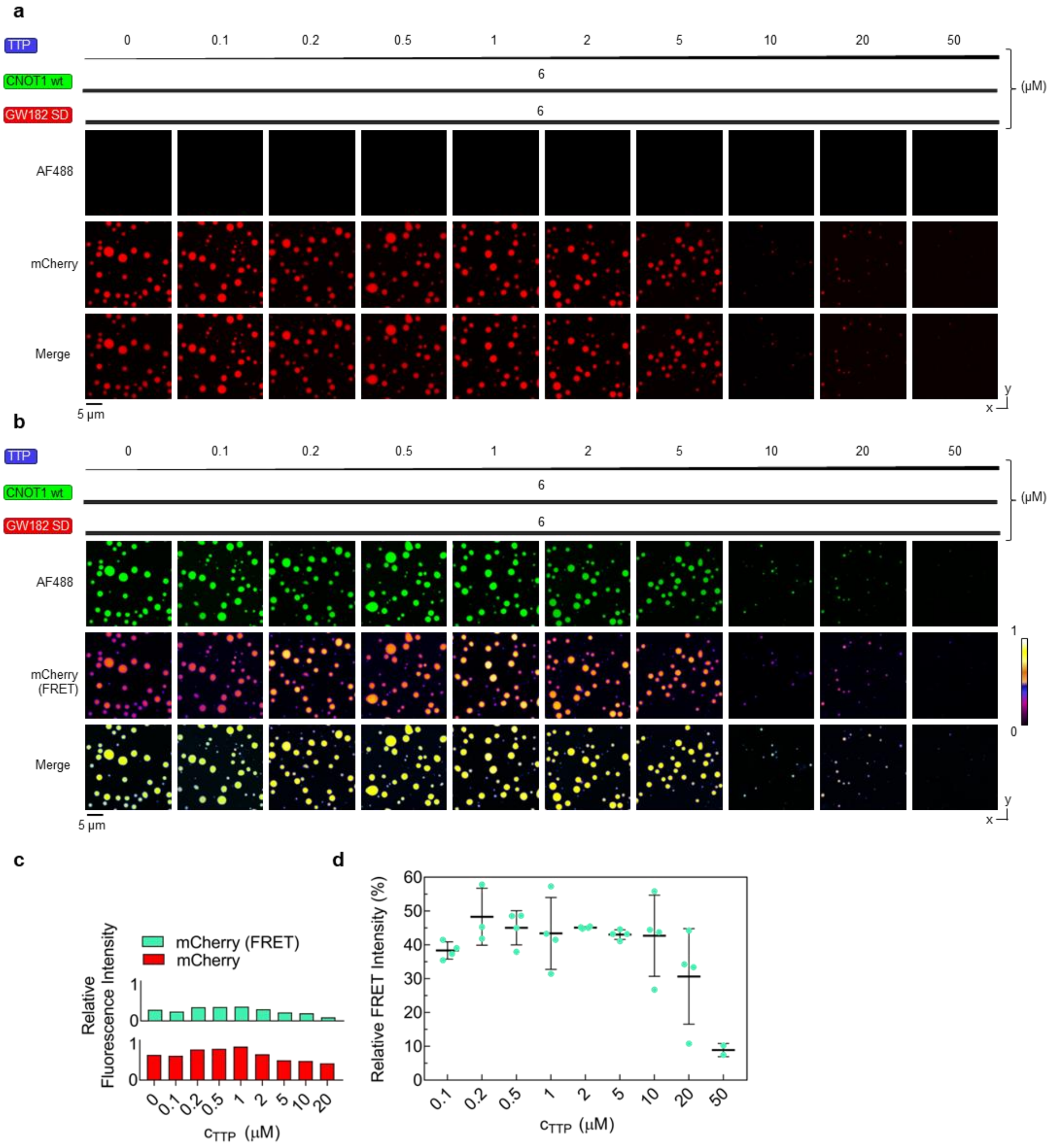

**Supplementary figure S27 to Fig 5k. FRET observed for the inhibition of multiprotein droplets composed of GW182 SD-mCherry and GST-CNOT1 wt (800-999)-AF488 by tristetraprolin (TTP) after 24 h. (a)** Confocal images acquired using single-wavelength excitation (mCherry channel), showing fluorescence of GW182 SD-mCherry (6 µM) in the dense phase upon formation of multiprotein liquid droplets with equimolar concentrations of GST-CNOT1 wt (800-999)-AF488 (6 µM) in 10% (w/v) PEG 1k, in the presence of increasing concentrations (0-50 µM) of TTP peptide. Images acquired after 24 h; n = 2-5. Scale bar: 5 µm. **(b)** Confocal images acquired using single-wavelength excitation (AF488 channel), showing fluorescence from GST-CNOT1 wt-AF488 (6 µM) in both unbound and bound states, and FRET-derived mCherry emission indicating complex formation with GW182 SD-mCherry (6 µM) in droplets formed in the presence of increasing concentrations (0-50 µM) of TTP peptide. Images acquired after 24 h; n = 2-5. Scale bar: 5 µm. **(c)** Normalized histograms of integrated fluorescence intensities from single-wavelength excitation of mCherry (FRET, teal) and mCherry (red) in the presence of increasing concentrations of TTP peptide. **(d)** Relative FRET intensity change after 24 h in liquid droplets formed by equimolar concentrations of GST-CNOT1 wt-AF488 and GW182 SD-mCherry in the presence of increasing concentrations of TTP peptide (teal).

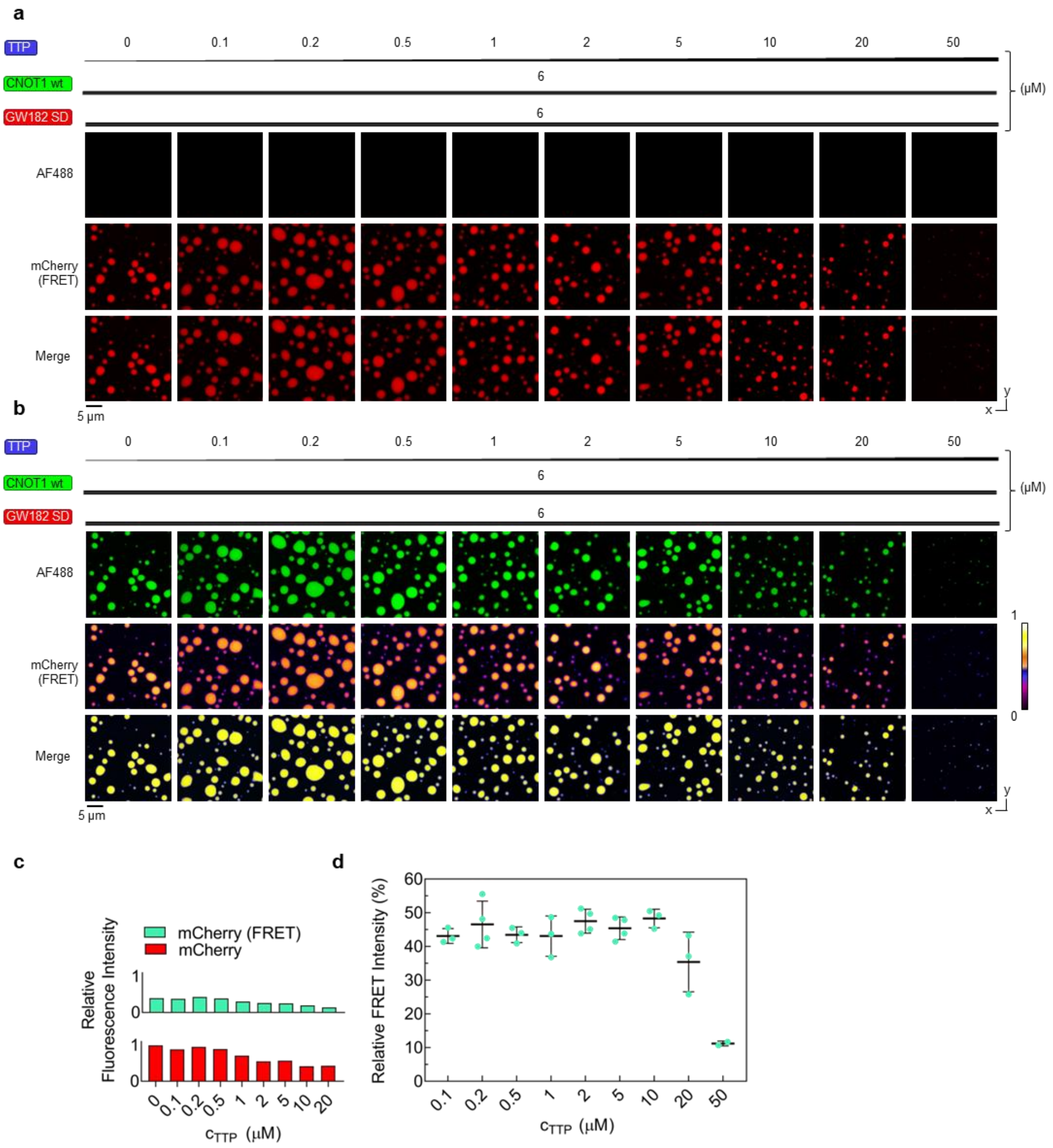

**Supplementary figure S28 to Fig 6a. Negative controls for the FRET-FRAP experiments. (a-c)** Comparisons of FRAP time-lapse images of liquid droplets formed by GW182 SD-mCherry (6 µM) in 10% (w/v) PEG 1k, acquired using single-wavelength excitation (mCherry channel) and photobleaching with: **(a)** 20% relative power of the Argon multiline laser, **(b)** 20% relative power of the DPSS laser. **(c-d)** Comparisons of FRAP time-lapse images of liquid droplets formed by GW182 SD-mCherry (6 µM) in 10% (w/v) PEG 1k, acquired using single-wavelength excitation (mCherry channel) and photobleaching with: **(c)** 100% relative power of the Argon multiline laser, **(d)** 100% relative power of the DPSS laser.

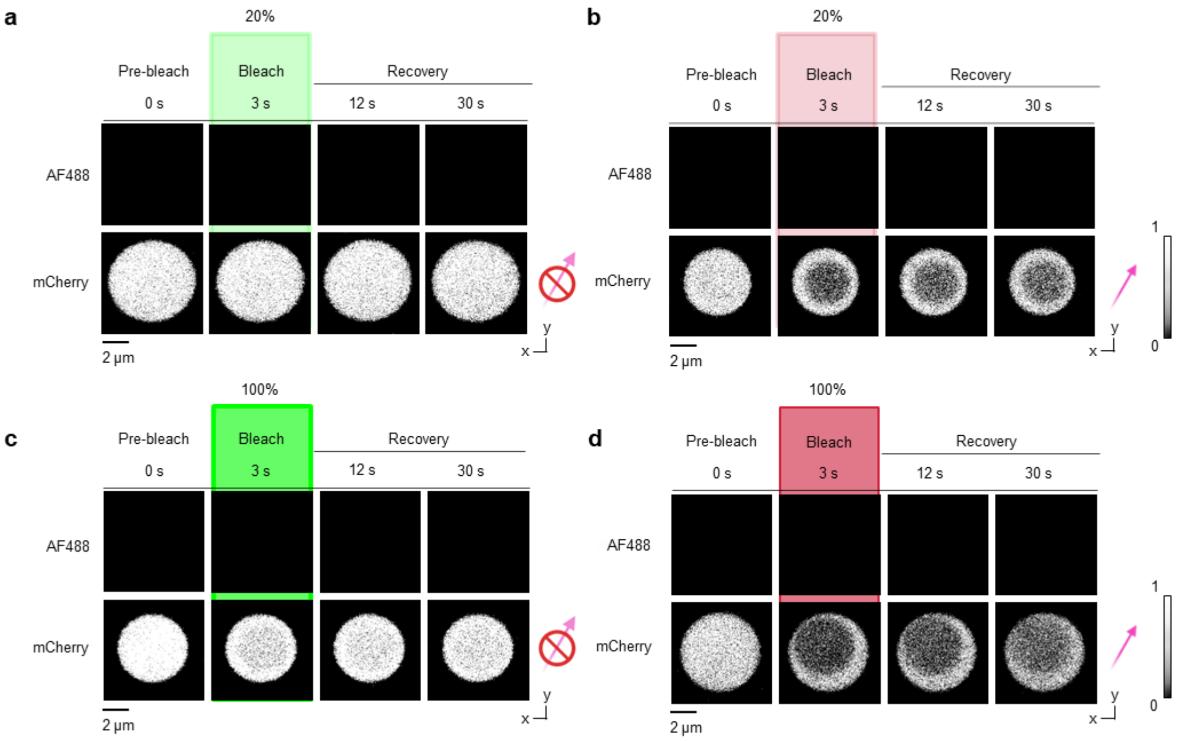

**Supplementary figure S29 to Fig 6b-e. FRET-FRAP for multiprotein droplets composed of GW182 SD-mCherry and either GST-CNOT1 wt (800-999)-AF488 or GST-CNOT1 EY (800-999)-AF488 after 24 h. (a-d)** FRET-FRAP time-lapse images of multiprotein droplets formed by GW182 SD-mCherry (6 µM) with increasing concentrations of GST-CNOT1 wt-AF488: **(a)** 1 µM, **(b)** 4 µM, **(c)** 7 µM, and **(d)** 4 µM GST-CNOT1 EY-AF488. Fluorescence recovery is shown in both the AF488 channel (unbound and bound states) and the mCherry channel (FRET, bound states). Images were acquired after 24 h; n = 2-5. Recovery was measured over 60 s. Scale bar: 1 µm. **(e-h)** Quantification of mean mobile fractions (%) in the bound and unbound state (AF488 channel, lime or green) and the FRET-only bound state (mCherry channel) for: **(e)** 1 µM CNOT1 wt (pink circles), **(f)** 4 µM CNOT1 wt (pink circles), **(g)** 7 µM CNOT1 wt (pink circles) and **(h)** 4 µM CNOT1 EY (purple circles)

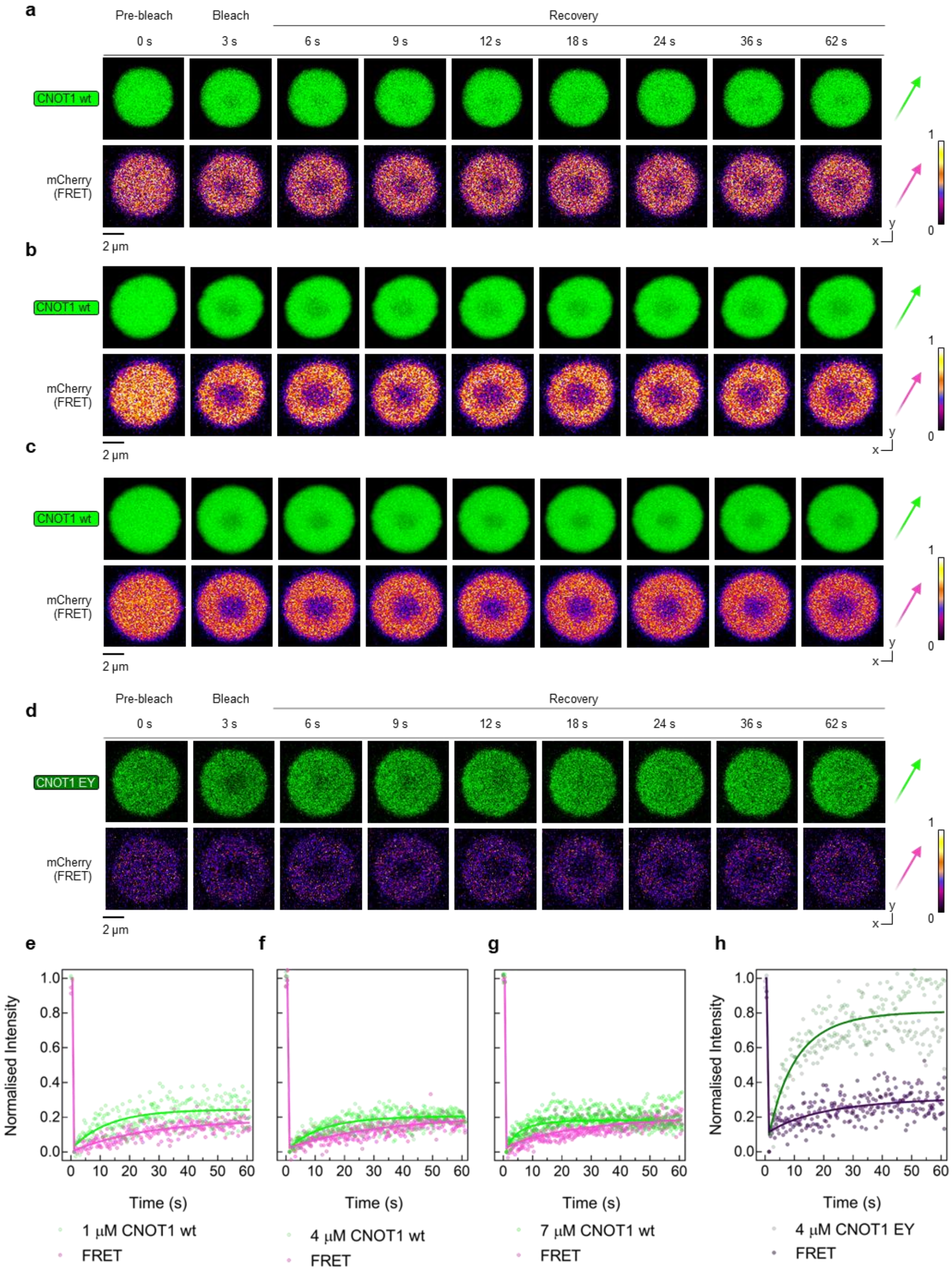

**Supplementary figure S30 to Fig 1f.**

In **Figure 1f**, the CNOT1 peptide IDDEANSY(839-846) displays a fraction of fast-exchanging protons in experiments with CIM1 and TTP, while only slow-exchanging protons are present in experiments with N-CIM1 and GW182 SD10. This difference comes from a bimodal **(a)** or monomodal **(b)** distribution of isotopic envelopes of the IDDEANSY peptide. The bimodal envelope **(a)** can be attributed to postranslational modifications of a fraction of CNOT1 used in the experiments with CIM1 and TTP. *E.g.*, triple oxidation increases the mass by 48 Da **(c, d)**. These modifications could make some protons in the IDDEANSY peptide more accessible to solvent, which is reflected in the m/z spectra as a bimodal isotopic envolope, best visible for 1 min of HDX as shown below **(a)**. Nevertheless, all tested ligands protected this peptide from HDX highlighting its' involvement in the formation of all the complexes.

**(a)**

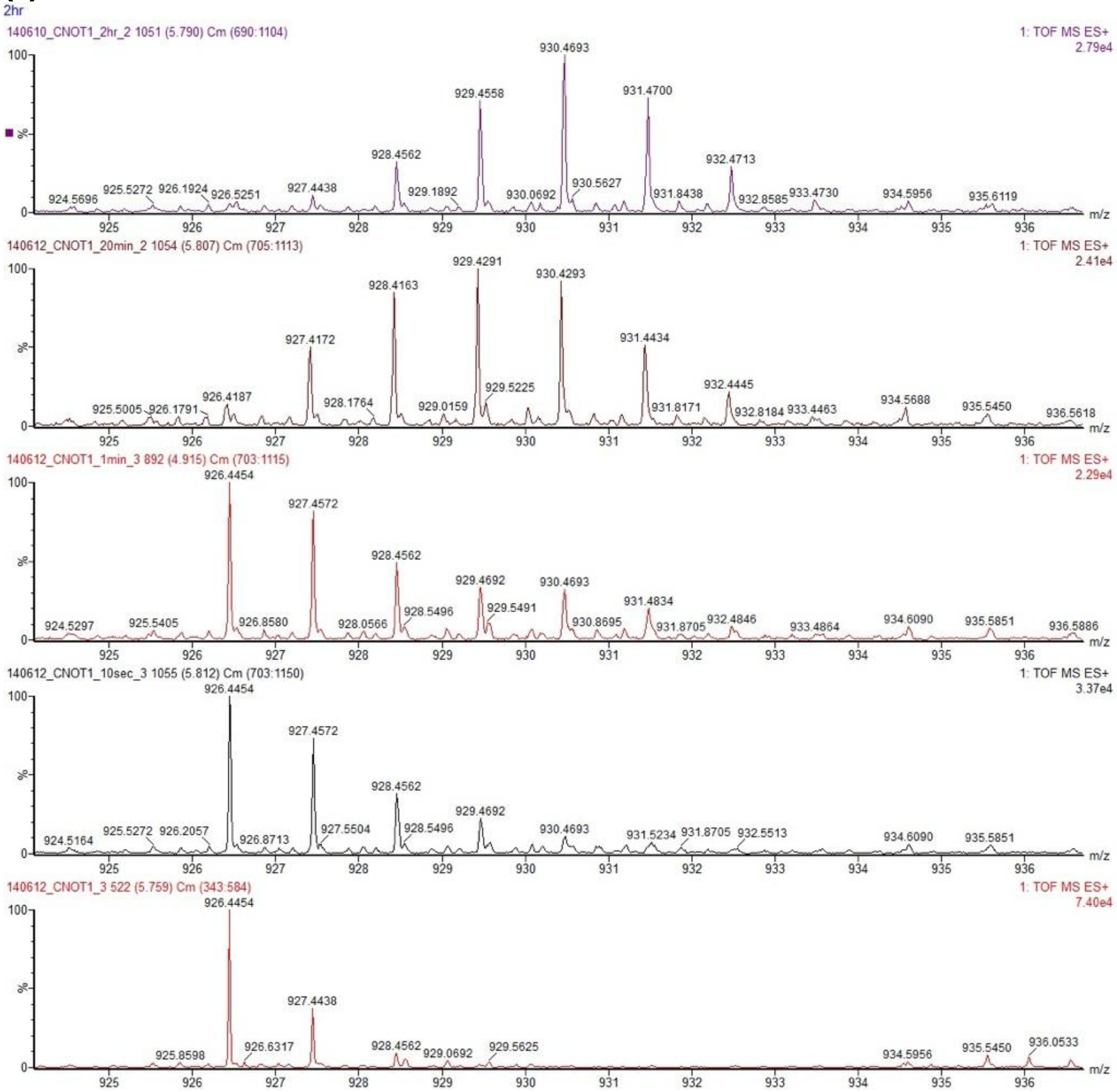

**(b)**

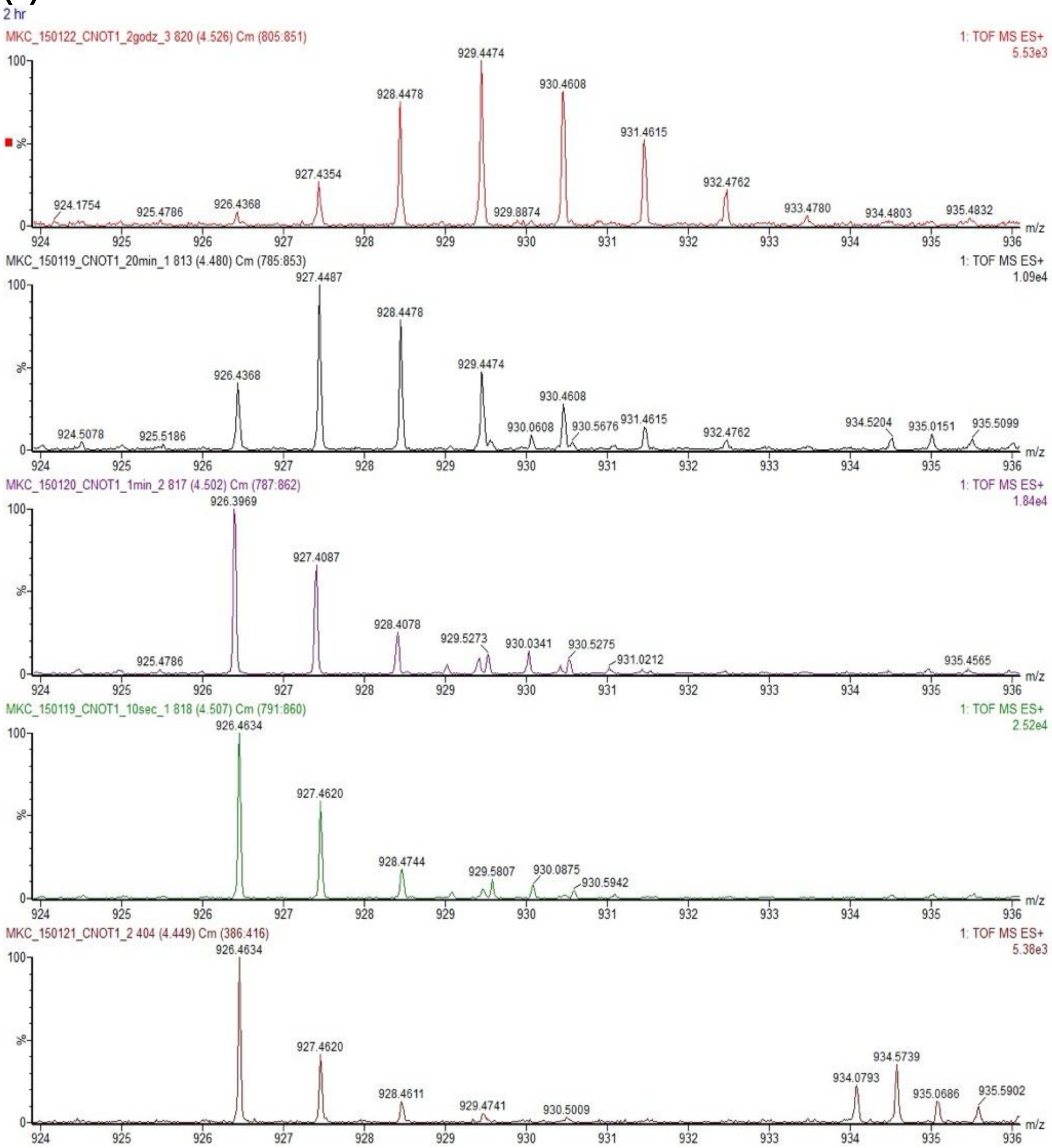
2 hr
MKC_150122_CNOT1_2godz_3 820 (4.526) Cm (805:851)
1: TOF MS ES+
5.53e3
929.4474
930.4608
928.4478
931.4615
927.4354
932.4762
924.1754
925.4786
926.4368
929.8874
933.4780
934.4803
935.4832
MKC_150119_CNOT1_20min_1 813 (4.480) Cm (785:853)
1: TOF MS ES+
1.09e4
927.4487
928.4478
929.4474
926.4368
930.4608
924.5078
925.5186
930.0608
930.5676
931.4615
932.4762
934.5204
935.0151
935.5099
MKC_150120_CNOT1_1min_2 817 (4.502) Cm (787:862)
1: TOF MS ES+
1.84e4
926.3969
927.4087
928.4078
929.5273
930.0341
930.5275
931.0212
925.4786
935.4565
MKC_150119_CNOT1_10sec_1 818 (4.507) Cm (791:860)
1: TOF MS ES+
2.52e4
926.4634
927.4620
928.4744
929.5807
930.0875
930.5942
MKC_150121_CNOT1_2 404 (4.449) Cm (386:416)
1: TOF MS ES+
5.38e3
926.4634
927.4620
928.4611
929.4741
930.5009
934.0793
934.5739
935.0686
935.5902
100
%
0
m/z

**(c)** Mass of CNOT1 used in the experiment with CIM1 and TTP:

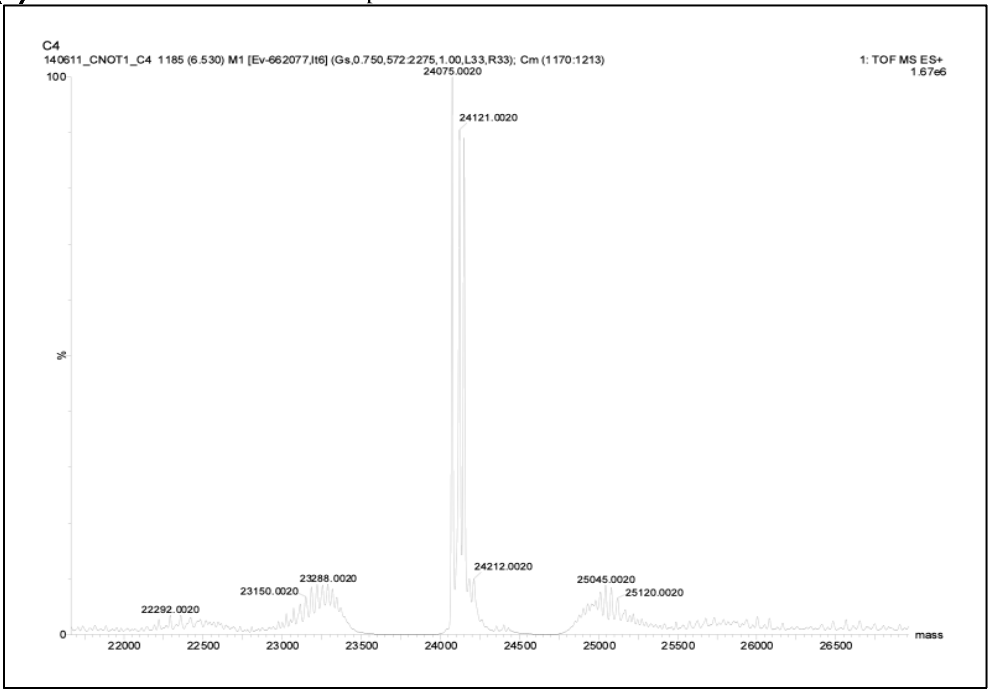

**(d)** Mass of CNOT1 used in the experiment with N-CIM1 and SD10:

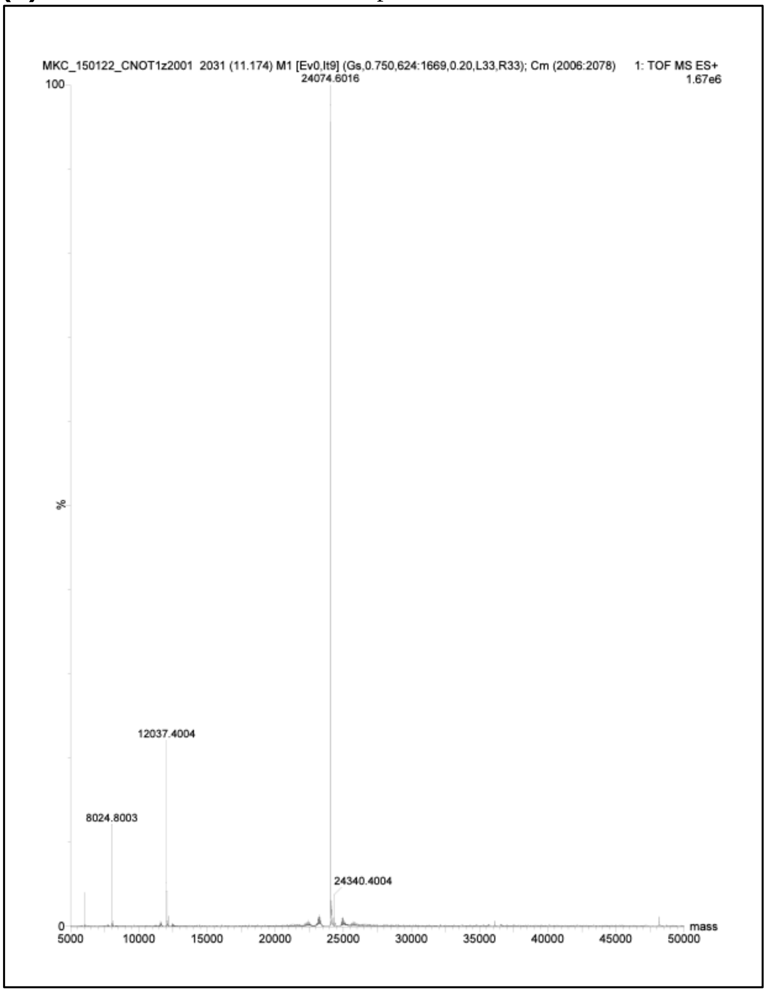

### Supplementary Table T1 to Fig 2f.

Kinetic parameters of hydrogen-deuterium exchange for CNOT1 at 2 µM in the presence of the N-CIM1 peptide at 144 µM. *state* denotes that CNOT1(800-999) was measured at 2 µM alone, *apo*, or in the presence of different binding partners, as indicated; $ND_i$, maximal number of protons exchanging at the corresponding HDX rate constant, $k_i$; *ND*, maximum number of protons exchanged; $D_{\%}$, percent of exchangeable amide hydrogens that undergoe exchange to deuterium; *P-value*, possibility that the improvement of the fit with use of the selected model could be random, according to the F-Snedecor's test; *%AIC*, chance (in %) that the selected model is correct, from the Akaike's Information Criteria; $\delta$, one standard deviation resulting from numerical fit.

| | | Best-fit values | | | | | | | | | | | | | | | | | |
|---|---|---|---|---|---|---|---|---|---|---|---|---|---|---|---|---|---|---|---|
| Amino acids/sequence | state | $ND_1$ [Da] | $\delta ND_1$ | $k_1$ [$min^{-1}$] | $\delta k_1$ | $ND_2$ [Da] | $\delta ND_2$ | $k_2$ [$min^{-1}$] | $\delta k_2$ | $ND_3$ [Da] | $\delta ND_3$ | $k_3$ [$min^{-1}$] | $\delta k_3$ | ND [Da] | $\delta$ND | $D_{\%}$ | $\delta D_{\%}$ | P-value | % AIC |
| 800-819 | apo | | | | | | | | | 21.44 | 0.08 | 27 | 4 | 21.44 | 0.08 | 126.1 | 0.5 | - | 90 |
| NNDPFVQRKLGTSGLNQPTF | + 70xN-CIM1 | | | | | | | | | 19.41 | 0.07 | 28 | 4 | 19.41 | 0.07 | 114.2 | 0.4 | - | 90 |
| 800-824 | apo | | | | | | | | | 27.44 | 0.13 | 27 | 5 | 27.44 | 0.13 | 124.7 | 0.6 | 0.3348 | 82 |
| NNDPFVQRKLGTSGLNQPTFQQTDL | + 70xN-CIM1 | | | | | | | | | 24.41 | 0.15 | 31 | 11 | 24.41 | 0.15 | 111.0 | 0.7 | - | 90 |
| 805-824 | apo | | | | | | | | | 23.39 | 0.10 | 26 | 4 | 23.39 | 0.10 | 129.9 | 0.6 | - | 90 |
| VQRKLGTSGLNQPTFQQTDL | + 70xN-CIM1 | | | | | | | | | 20.73 | 0.11 | 26 | 5 | 20.73 | 0.11 | 115.2 | 0.6 | - | 90 |
| 820-824 | apo | | | | | | | | | 4.271 | 0.015 | 22.9 | 1.8 | 4.271 | 0.015 | 106.8 | 0.4 | - | 90 |
| QQTDL | + 70xN-CIM1 | | | | | | | | | 4.195 | 0.018 | 31 | 8 | 4.195 | 0.018 | 104.9 | 0.5 | - | 90 |
| 825-835 | apo | | | | | | | | | 11.50 | 0.06 | 22 | 3 | 11.50 | 0.06 | 127.8 | 0.7 | - | 80 |
| SQVWPEANQHF | + 70xN-CIM1 | | | | | | | | | 10.37 | 0.05 | 16.1 | 0.9 | 10.37 | 0.05 | 115.2 | 0.6 | 0.9526 | > 99 |
| 825-838 | apo | | | | | | | | | 15.69 | 0.06 | 13.3 | 0.4 | 15.69 | 0.06 | 130.8 | 0.5 | - | > 99 |
| SQVWPEANQHFSKE | + 70xN-CIM1 | peptide n.d. | | | | | | | | | | | | | | | | | |
| 825-841 | apo | 3.1 | 0.3 | 0.064 | 0.016 | | | | | 16.6 | 0.2 | 14.4 | 1.2 | 19.6 | 0.2 | 130.9 | 1.4 | < 0.0001 | > 99 |
| SQVWPEANQHFSKEIDD | + 70xN-CIM1 | 2.6 | 0.3 | 0.038 | 0.010 | | | | | 14.97 | 0.20 | 14.3 | 1.1 | 17.59 | 0.20 | 117.3 | 1.3 | < 0.0001 | > 99 |
| 828-838 | apo | | | | | | | | | 11.78 | 0.09 | 15.8 | 1.2 | 11.78 | 0.09 | 130.9 | 0.9 | - | 96 |
| WPEANQHFSKE | + 70xN-CIM1 | peptide n.d. | | | | | | | | | | | | | | | | | |
| 839-846 | apo | 6.47 | 0.14 | 0.0287 | 0.0020 | | | | | 0.33 | 0.08 | 30[a] | | 6.80 | 0.13 | 97.2 | 1.9 | 0.6434 | 93 |
| IDDEANSY | + 70xN-CIM1 | 2.44 | 0.11 | 0.028 | 0.004 | | | | | | | | | 2.44 | 0.11 | 34.8 | 1.6 | 0.0638 | 73 |
| 839-847 | apo | 7.6 | 0.2 | 0.031 | 0.003 | | | | | | | | | 7.6 | 0.2 | 95 | 3 | 0.1038 | 59 |
| IDDEANSYF | + 70xN-CIM1 | 2.4 | 0.5 | 0.012 | 0.005 | | | | | | | | | 2.4 | 0.5 | 30 | 7 | 0.3315 | 97 |
| 847-859 | apo | 5.15 | 0.12 | 0.044 | 0.002 | 3.60 | 0.08 | 7.1 | 0.5 | | | | | 8.75 | 0.09 | 97.2 | 1.0 | < 0.0001 | > 99 |
| FQRIYNHPPHPTM | + 70xN-CIM1 | 2.96 | 0.13 | 0.080 | 0.010 | 2.76 | 0.11 | 6.9 | 0.8 | | | | | 5.72 | 0.09 | 63.5 | 1.0 | < 0.0001 | > 99 |
| 848-859 | apo | 4.06 | 0.14 | 0.055 | 0.004 | 4.01 | 0.10 | 6.1 | 0.5 | | | | | 8.07 | 0.09 | 100.8 | 1.1 | < 0.0001 | > 99 |
| QRIYNHPPHPTM | + 70xN-CIM1 | 3.29 | 0.11 | 0.090 | 0.009 | 2.56 | 0.09 | 7.5 | 0.8 | | | | | 5.85 | 0.07 | 73.1 | 0.9 | < 0.0001 | > 99 |
| 860-865 | apo | 1.05 | 0.14 | 0.15 | 0.12 | | | | | 1.30 | 0.16 | 10 | 3 | 2.35 | 0.08 | 46.9 | 1.7 | 0.0001 | > 99 |
| SVDEVL | + 70xN-CIM1 | 1.07 | 0.06 | 0.054 | 0.007 | 0.98 | 0.05 | 6.2 | 0.9 | | | | | 2.05 | 0.04 | 41.0 | 0.8 | < 0.0001 | > 99 |
| 863-867 | apo | HDX n.d. | | | | | | | | | | | | | | 0 | | | |
| EVLEM | + 70xN-CIM1 | HDX n.d. | | | | | | | | | | | | | | 0 | | | |
| 866-883 | apo | 2.73 | 0.07 | 0.043 | 0.003 | | | | | 3.38 | 0.04 | 30[a] | | 6.10 | 0.06 | 35.9 | 0.3 | 0.4134 | 91 |
| EMLQRFKDSTIKREREVF | + 70xN-CIM1 | 2.73 | 0.08 | 0.029 | 0.003 | | | | | 3.41 | 0.06 | 17 | 2 | 6.15 | 0.07 | 36.1 | 0.4 | < 0.0001 | > 99 |
| 868-883 | apo | 2.29 | 0.07 | 0.042 | 0.003 | | | | | 3.91 | 0.04 | 30[a] | | 6.20 | 0.06 | 41.3 | 0.4 | 0.5823 | 93 |
| LQRFKDSTIKREREVF | + 70xN-CIM1 | 2.13 | 0.11 | 0.028 | 0.005 | | | | | 4.05 | 0.08 | 17 | 3 | 6.18 | 0.10 | 41.2 | 0.6 | < 0.0001 | > 99 |
| 886-893 | apo | HDX n.d. | | | | | | | | | | | | | | 0 | | | |
| MLRNLFEE | + 70xN-CIM1 | HDX n.d. | | | | | | | | | | | | | | 0 | | | |
| 887-893 | apo | HDX n.d. | | | | | | | | | | | | | | 0 | | | |
| LRNLFEE | + 70xN-CIM1 | HDX n.d. | | | | | | | | | | | | | | 0 | | | |
| 894-909 | apo | 8.02 | 0.08 | 0.0338 | 0.0010 | 2.37 | 0.06 | 5.2 | 0.4 | | | | | 10.39 | 0.06 | 79.9 | 0.5 | < 0.0001 | > 99 |
| YRFFPQYPDKELHITA | + 70xN-CIM1 | 4.85 | 0.07 | 0.0310 | 0.0014 | | | | | 0.93 | 0.05 | 13 | 4 | 5.79 | 0.06 | 44.5 | 0.5 | < 0.0001 | > 99 |
| 898-909 | apo | 6.2 | 0.4 | 0.037 | 0.006 | 2.5 | 0.3 | 4.0 | 1.3 | | | | | 8.6 | 0.2 | 86 | 2 | < 0.0001 | > 99 |
| PQYPDKELHITA | + 70xN-CIM1 | 4.00 | 0.14 | 0.043 | 0.004 | | | | | 0.86 | 0.09 | 30[a] | | 4.86 | 0.12 | 48.6 | 1.2 | 0.8249 | 94 |
| 901-909 | apo | 4.89 | 0.16 | 0.051 | 0.004 | 1.24 | 0.12 | 7 | 2 | | | | | 6.13 | 0.11 | 76.6 | 1.4 | < 0.0001 | > 99 |
| PDKELHITA | + 70xN-CIM1 | 2.79 | 0.11 | 0.051 | 0.005 | | | | | 0.46 | 0.06 | 30[a] | | 3.25 | 0.09 | 40.6 | 1.1 | 0.8642 | 94 |
| 912-917 | apo | HDX n.d. | | | | | | | | | | | | | | 0 | | | |
| FGGIIE | + 70xN-CIM1 | HDX n.d. | | | | | | | | | | | | | | 0 | | | |
| 912-923 | apo | 4.06 | 0.06 | 0.0183 | 0.0008 | | | | | 2.75 | 0.03 | 12.9 | 0.6 | 6.82 | 0.06 | 62.0 | 0.6 | < 0.0001 | > 99 |
| FGGIIEKGLVTY | + 70xN-CIM1 | 4.0 | 0.3 | 0.0101 | 0.0018 | 2.76 | 0.04 | 9.1 | 0.5 | | | | | 6.7 | 0.4 | 61 | 3 | < 0.0001 | > 99 |
| 924-928 | apo | 3.0 | 0.2 | 0.030 | 0.005 | 0.8 | 0.2 | 1.4 | 0.7 | | | | | 3.85 | 0.06 | 96.2 | 1.5 | < 0.0001 | > 99 |
| MALGL | + 70xN-CIM1 | 2.6 | 0.2 | 0.039 | 0.005 | 0.3 | 0.2 | 1.4 | 1.7 | | | | | 2.97 | 0.04 | 74.3 | 0.9 | 0.0066 | 94 |
| 924-930 | apo | 3.03 | 0.13 | 0.040 | 0.005 | | | | | 0.34 | 0.08 | 30[a] | | 3.36 | 0.11 | 56.0 | 1.9 | 0.0015 | 99 |
| MALGLAL | + 70xN-CIM1 | 2.45 | 0.06 | 0.042 | 0.003 | 0.27 | 0.05 | 4 | 2 | | | | | 2.72 | 0.04 | 45.3 | 0.7 | 0.0002 | > 99 |
| 925-930 | apo | 2.45 | 0.18 | 0.051 | 0.014 | | | | | | | | | 2.45 | 0.18 | 49 | 6 | 0.0804 | 85 |
| ALGLAL | + 70xN-CIM1 | peptide n.d. | | | | | | | | | | | | | | | | | |
| 929-934 | apo | HDX n.d. | | | | | | | | | | | | | | 0 | | | |
| ALRYVL | + 70xN-CIM1 | HDX n.d. | | | | | | | | | | | | | | 0 | | | |
| 929-936 | apo | peptide n.d. | | | | | | | | | | | | | | | | | |
| ALRYVLEA | + 70xN-CIM1 | peptide n.d. | | | | | | | | | | | | | | | | | |
| 931-936 | apo | HDX n.d. | | | | | | | | | | | | | | 0 | | | |
| RYVLEA | + 70xN-CIM1 | peptide n.d. | | | | | | | | | | | | | | | | | |
| 937-945 | apo | 0.71 | 0.10 | 0.07 | 0.03 | 6.27 | 0.08 | 7.8 | 0.3 | | | | | 6.98 | 0.07 | 99.7 | 1.0 | 0.0002 | > 99 |
| LRKPFGSKM | + 70xN-CIM1 | 0.74 | 0.07 | 0.030 | 0.009 | 6.05 | 0.05 | 5.92 | 0.15 | | | | | 6.79 | 0.06 | 97.1 | 0.8 | < 0.0001 | > 99 |
| 946-951 | apo | HDX n.d. | | | | | | | | | | | | | | 0 | | | |

| | | Best-fit values | | | | | | | | | | | | | | | | | |
|---|---|---|---|---|---|---|---|---|---|---|---|---|---|---|---|---|---|---|---|
| Amino acids/sequence | state | $ND_1$ [Da] | $\delta ND_1$ [Da] | $k_1$ [$min^{-1}$] | $\delta k_1$ [$min^{-1}$] | $ND_2$ [Da] | $\delta ND_2$ [Da] | $k_2$ [$min^{-1}$] | $\delta k_2$ [$min^{-1}$] | $ND_3$ [Da] | $\delta ND_3$ [Da] | $k_3$ [$min^{-1}$] | $\delta k_3$ [$min^{-1}$] | ND [Da] | δND [Da] | $D_{\%}$ | $\delta D_{\%}$ | P-value | % AIC |
| YYFGIA | + 70xN-CIM1 | HDX n.d. | | | | | | | | | | | | | | 0 | | | |
| 946-952 | apo | HDX n.d. | | | | | | | | | | | | | | 0 | | | |
| YYFGIAA | + 70xN-CIM1 | HDX n.d. | | | | | | | | | | | | | | 0 | | | |
| 948-953 | apo | HDX n.d. | | | | | | | | | | | | | | 0 | | | |
| FGIAAL | + 70xN-CIM1 | HDX n.d. | | | | | | | | | | | | | | 0 | | | |
| 952-966 | apo | 3.3 | 1.0 | 0.009 | 0.005 | | | | | 3.48 | 0.06 | 30[a] | | 6.7 | 1.0 | 52 | 8 | < 0.0001 | > 99 |
| ALDRFKNRLKDYPQY | + 70xN-CIM1 | 1.9 | 0.6 | 0.014 | 0.010 | | | | | 3.34 | 0.13 | 10.2 | 1.6 | 5.3 | 0.6 | 41 | 5 | < 0.0001 | > 99 |
| 954-966 | apo | 2.7 | 0.5 | 0.010 | 0.004 | | | | | 3.54 | 0.06 | 13.5 | 1.3 | 7.6 | 1.6 | 69 | 14 | < 0.0001 | > 99 |
| DRFKNRLKDYPQY | + 70xN-CIM1 | 7 | 27 | 0.002 | 0.008 | 3.62 | 0.07 | 7.7 | 0.5 | | | | | 11 | 27 | 100 | 200 | < 0.0001 | > 99 |
| 954-972 | apo | 5.5 | 0.3 | 0.025 | 0.005 | | | | | 3.1 | 0.2 | 20 | 14 | 8.6 | 0.3 | 50.9 | 1.7 | < 0.0001 | > 99 |
| DRFKNRLKDYPQYCQHLAS | + 70xN-CIM1 | 4.2 | 0.3 | 0.034 | 0.007 | | | | | 3.0 | 0.2 | 12 | 5 | 7.2 | 0.2 | 42.5 | 1.4 | < 0.0001 | > 99 |
| 967-972 | apo | 2.09 | 0.08 | 0.061 | 0.008 | | | | | | | | | 2.09 | 0.08 | 41.9 | 1.5 | 0.3513 | 97 |
| CQHLAS | + 70xN-CIM1 | 2.19 | 0.08 | 0.049 | 0.006 | | | | | | | | | 2.19 | 0.08 | 43.8 | 1.7 | - | > 99 |
| 973-985 | apo | 7.4 | 0.5 | 0.053 | 0.008 | 3.5 | 0.4 | 4.3 | 1.4 | | | | | 10.9 | 0.3 | 99 | 3 | 0.0003 | > 99 |
| ISHFMQFPHHLQE | + 70xN-CIM1 | 7.0 | 0.9 | 0.050 | 0.012 | 2.8 | 0.8 | 5 | 3 | | | | | 9.8 | 0.4 | 89 | 4 | 0.0044 | 90 |
| 977-985 | apo | 3.8 | 0.9 | 0.080 | 0.019 | 1.4 | 0.9 | 1.6 | 2.0 | | | | | 5.18 | 0.16 | 74 | 2 | 0.0257 | 58 |
| MQFPHHLQE | + 70xN-CIM1 | 3.86 | 0.15 | 0.27 | 0.07 | | | | | | | | | 3.86 | 0.15 | 55 | 2 | 0.8631 | 90 |
| 977-986 | apo | 3.9 | 0.3 | 0.074 | 0.011 | 1.1 | 0.2 | 2.9 | 1.7 | | | | | 5.03 | 0.14 | 62.9 | 1.7 | 0.0028 | 97 |
| MQFPHHLQEY | + 70xN-CIM1 | peptide n.d. | | | | | | | | | | | | | | | | | |
| 986-999 | apo | 5.12 | 0.05 | 0.0668 | 0.0015 | 6.16 | 0.04 | 4.20 | 0.07 | | | | | 11.28 | 0.03 | 102.5 | 0.3 | < 0.0001 | > 99 |
| YIEYGQQSRDPPVK | + 70xN-CIM1 | 5.12 | 0.16 | 0.060 | 0.004 | 5.02 | 0.14 | 3.4 | 0.3 | | | | | 10.13 | 0.09 | 92.1 | 0.8 | < 0.0001 | > 99 |
| 989-999 | apo | 3.06 | 0.15 | 0.13 | 0.03 | 6.94 | 0.16 | 4.3 | 0.2 | | | | | 10.00 | 0.09 | 125.0 | 1.2 | < 0.0001 | > 99 |
| YGQQSRDPPVK | + 70xN-CIM1 | 3.38 | 0.13 | 0.110 | 0.014 | 6.02 | 0.12 | 3.68 | 0.19 | | | | | 9.40 | 0.08 | 117.5 | 1.0 | < 0.0001 | > 99 |

[a] Fixed values, according to (Gemmecker et al., 1993; Hoofnagle et al., 2001; Mandell et al., 2001; Rutkowska-Wlodarczyk et al., 2008).

[b] exceptions to the classification that $k_1 \leq 0.1$, $0.1 < k_2 < 10$ and $10 < k_3 \leq 30$, due to lack of space in appropriate column.

**Supplementary Table T2 to Fig 2f.**

Kinetic parameters of hydrogen-deuterium exchange for CNOT1 at 2 µM in the presence of the CIM1 peptide at 12.6 µM or 126 µM. The control data (the *apo* state) is the same as in (Cieplak-Rotowska et al., 2025).

| | | Best-fit values | | | | | | | | | | | | | | | | | |
|---|---|---|---|---|---|---|---|---|---|---|---|---|---|---|---|---|---|---|---|
| Amino acids/sequence | state | $ND_1$ [Da] | $\delta ND_1$ | $k_1$ [min$^{-1}$] | $\delta k_1$ | $ND_2$ [Da] | $\delta ND_2$ | $k_2$ [min$^{-1}$] | $\delta k_2$ | $ND_3$ [Da] | $\delta ND_3$ | $k_3$ [min$^{-1}$] | $\delta k_3$ | ND [Da] | $\delta$ND | $D_{\%}$ | $\delta D_{\%}$ | P-value | % AIC |
| 800-818 | apo | | | | | | | | | 17.42 | 0.05 | 24 | 2 | 17.42 | 0.05 | 108.9 | 0.3 | - | 86 |
| NNDPFVQRKLGTSGLNQPT | + 6.3 x CIM1 | | | | | | | | | 17.20 | 0.06 | 22.5 | 1.8 | 17.20 | 0.06 | 107.5 | 0.4 | - | 90 |
| | + 63 x CIM1 | | | | | | | | | 16.03 | 0.06 | 20.2 | 1.6 | 16.03 | 0.06 | 100.2 | 0.4 | >0.9999 | 87 |
| 800-819 | apo | | | | | | | | | 19.18 | 0.12 | 27 | 7 | 19.18 | 0.12 | 112.8 | 0.7 | - | 86 |
| NNDPFVQRKLGTSGLNQPTF | + 6.3 x CIM1 | | | | | | | | | 19.10 | 0.16 | 24 | 5 | 19.10 | 0.16 | 112.4 | 0.9 | - | 90 |
| | + 63 x CIM1 | | | | | | | | | 18.76 | 0.11 | 28 | 8 | 18.76 | 0.11 | 110.4 | 0.6 | - | 87 |
| 800-820 | apo | | | | | | | | | 19.71 | 0.07 | 27 | 4 | 19.71 | 0.07 | 109.5 | 0.4 | - | 86 |
| NNDPFVQRKLGTSGLNQPTF | + 6.3 x CIM1 | | | | | | | | | 19.56 | 0.10 | 29 | 8 | 19.56 | 0.10 | 108.7 | 0.6 | - | 90 |
| Q | + 63 x CIM1 | | | | | | | | | 19.01 | 0.09 | 28 | 6 | 19.01 | 0.09 | 105.6 | 0.5 | - | 87 |
| 800-824 | apo | | | | | | | | | 24.76 | 0.09 | 29 | 5 | 24.76 | 0.09 | 112.5 | 0.4 | - | 86 |
| NNDPFVQRKLGTSGLNQPTF | + 6.3 x CIM1 | | | | | | | | | 24.75 | 0.12 | 30 | 8 | 24.75 | 0.12 | 112.5 | 0.5 | - | 90 |
| QQTDL | + 63 x CIM1 | | | | | | | | | 24.24 | 0.10 | 26 | 4 | 24.24 | 0.10 | 110.2 | 0.5 | - | 87 |
| 820-824 | apo | | | | | | | | | 4.380 | 0.012 | 20.8 | 1.2 | 4.380 | 0.012 | 109.5 | 0.3 | 0.6800 | 85 |
| QQTDL | + 6.3 x CIM1 | | | | | | | | | 4.388 | 0.017 | 24 | 3 | 4.388 | 0.017 | 109.7 | 0.4 | - | 90 |
| | + 63 x CIM1 | | | | | | | | | 4.376 | 0.013 | 23.6 | 1.9 | 4.376 | 0.013 | 109.4 | 0.3 | - | 87 |
| 825-835 | apo | | | | | | | | | 9.79 | 0.04 | 18.7 | 1.1 | 9.79 | 0.04 | 108.9 | 0.4 | - | > 99 |
| SQVWPEANQHF | + 6.3 x CIM1 | | | | | | | | | 9.49 | 0.05 | 19.9 | 1.8 | 9.49 | 0.05 | 105.4 | 0.6 | - | 96 |
| | + 63 x CIM1 | | | | | | | | | 8.81 | 0.04 | 16.9 | 1.0 | 8.81 | 0.04 | 97.8 | 0.5 | - | > 99 |
| 825-838 | apo | | | | | | | | | 13.82 | 0.04 | 15.5 | 0.5 | 13.82 | 0.04 | 115.2 | 0.4 | - | > 99 |
| SQVWPEANQHFSKE | + 6.3 x CIM1 | | | | | | | | | 13.48 | 0.05 | 15.0 | 0.6 | 13.48 | 0.05 | 112.3 | 0.4 | - | > 99 |
| | + 63 x CIM1 | | | | | | | | | 12.40 | 0.08 | 14.7 | 0.9 | 12.40 | 0.08 | 103.3 | 0.6 | 0.4862 | 96 |
| 825-847 | apo | 8.5 | 0.4 | 0.043 | 0.005 | | | | | 12.1 | 0.3 | 12.4 | 1.5 | 20.7 | 0.2 | 98.3 | 1.0 | < 0.0001 | > 99 |
| SQVWPEANQHFSKEIDDEA | + 6.3 x CIM1 | 8.1 | 0.2 | 0.052 | 0.004 | | | | | 11.78 | 0.18 | 12.4 | 0.9 | 19.87 | 0.16 | 94.6 | 0.8 | < 0.0001 | > 99 |
| NSYF | + 63 x CIM1 | 7.8 | 0.4 | 0.026 | 0.005 | | | | | 11.7 | 0.3 | 10.8 | 1.3 | 19.5 | 0.3 | 93.0 | 1.4 | < 0.0001 | > 99 |
| 828-838 | apo | | | | | | | | | 10.56 | 0.09 | 18 | 2 | 10.56 | 0.09 | 117.3 | 1.0 | 0.9701 | 98 |
| WPEANQHFSKE | + 6.3 x CIM1 | | | | | | | | | 10.37 | 0.09 | 17.3 | 1.9 | 10.37 | 0.09 | 115.2 | 1.1 | - | 96 |
| | + 63 x CIM1 | peptide n.d. | | | | | | | | | | | | | | | | | |
| 829-838 | apo | | | | | | | | | 10.56 | 0.09 | 16.5 | 1.7 | 10.56 | 0.09 | 117.3 | 1.0 | - | 99 |
| PEANQHFSKE | + 6.3 x CIM1 | peptide n.d. | | | | | | | | | | | | | | | | | |
| | + 63 x CIM1 | peptide n.d. | | | | | | | | | | | | | | | | | |
| 839-846 | apo | 4.5 | 0.2 | 0.041 | 0.005 | 2.74 | 0.18 | 5.2 | 1.0 | | | | | 7.22 | 0.11 | 103.1 | 1.6 | < 0.0001 | > 99 |
| IDDEANSY | + 6.3 x CIM1 | 4.19 | 0.14 | 0.038 | 0.003 | 2.71 | 0.11 | 4.1 | 0.5 | | | | | 6.90 | 0.10 | 98.6 | 1.4 | < 0.0001 | > 99 |
| | + 63 x CIM1 | 4.50 | 0.15 | 0.0125 | 0.0006 | 2.24 | 0.15 | 1.8 | 0.3 | | | | | 6.74 | 0.06 | 96.2 | 0.8 | < 0.0001 | > 99 |
| 839-847 | apo | 5.1 | 0.3 | 0.040 | 0.005 | 3.0 | 0.2 | 5.2 | 1.1 | | | | | 8.04 | 0.14 | 100.5 | 1.8 | < 0.0001 | > 99 |
| IDDEANSYF | + 6.3 x CIM1 | 4.26 | 0.16 | 0.035 | 0.003 | 2.93 | 0.14 | 3.0 | 0.4 | | | | | 7.18 | 0.10 | 89.8 | 1.3 | < 0.0001 | > 99 |
| | + 63 x CIM1 | 4.9 | 0.5 | 0.0118 | 0.0016 | 1.9 | 0.4 | 1.8 | 0.9 | | | | | 6.79 | 0.18 | 85 | 2 | 0.0011 | 99 |
| 846-859 | apo | 5.8 | 0.3 | 0.047 | 0.007 | 4.0 | 0.3 | 8.3 | 2.0 | | | | | 9.84 | 0.19 | 98.4 | 1.9 | < 0.0001 | > 99 |
| YFQRIYNHPPHPTM | + 6.3 x CIM1 | 5.9 | 0.2 | 0.060 | 0.006 | 3.71 | 0.19 | 6.5 | 1.0 | | | | | 9.58 | 0.17 | 95.8 | 1.7 | < 0.0001 | > 99 |
| | + 63 x CIM1 | 5.0 | 0.3 | 0.044 | 0.006 | 3.3 | 0.2 | 5.0 | 1.1 | | | | | 8.23 | 0.16 | 82.3 | 1.6 | < 0.0001 | > 99 |
| 847-859 | apo | 4.75 | 0.19 | 0.056 | 0.006 | 4.44 | 0.17 | 6.4 | 0.7 | | | | | 9.19 | 0.10 | 102.1 | 1.2 | < 0.0001 | > 99 |
| FQRIYNHPPHPTM | + 6.3 x CIM1 | 4.51 | 0.09 | 0.064 | 0.003 | 4.15 | 0.07 | 6.1 | 0.3 | | | | | 8.65 | 0.06 | 96.1 | 0.7 | < 0.0001 | > 99 |
| | + 63 x CIM1 | 4.02 | 0.19 | 0.054 | 0.006 | 3.32 | 0.16 | 5.9 | 0.8 | | | | | 7.34 | 0.11 | 81.5 | 1.2 | < 0.0001 | > 99 |
| 847-861 | apo | 4.5 | 0.2 | 0.053 | 0.007 | 6.59 | 0.19 | 7.1 | 0.6 | | | | | 11.07 | 0.12 | 100.6 | 1.1 | < 0.0001 | > 99 |
| FQRIYNHPPHPTMSV | + 6.3 x CIM1 | 4.3 | 0.2 | 0.054 | 0.007 | 6.37 | 0.19 | 6.1 | 0.5 | | | | | 10.70 | 0.17 | 97.3 | 1.5 | < 0.0001 | > 99 |
| | + 63 x CIM1 | 3.9 | 0.4 | 0.065 | 0.017 | 5.6 | 0.4 | 4.1 | 0.8 | | | | | 9.5 | 0.2 | 86. | 2 | 0.001 | 99 |
| 860-865 | apo | 1.16 | 0.17 | 0.08 | 0.03 | 1.34 | 0.15 | 7 | 2 | | | | | 2.51 | 0.09 | 50.1 | 1.8 | 0.0018 | 99 |
| SVDEVL | + 6.3 x CIM1 | 1.07 | 0.05 | 0.106 | 0.017 | 1.33 | 0.04 | 7.0 | 0.6 | | | | | 2.39 | 0.03 | 47.9 | 0.6 | < 0.0001 | > 99 |
| | + 63 x CIM1 | 1.19 | 0.07 | 0.11 | 0.03 | 1.25 | 0.07 | 6.3 | 0.9 | | | | | 2.44 | 0.04 | 48.9 | 0.8 | < 0.0001 | > 99 |
| 862-867 | apo | HDX n.d. | | | | | | | | | | | | | | 0 | | | |
| DEVLEM | + 6.3 x CIM1 | HDX n.d. | | | | | | | | | | | | | | 0 | | | |
| | + 63 x CIM1 | HDX n.d. | | | | | | | | | | | | | | 0 | | | |
| 863-867 | apo | HDX n.d. | | | | | | | | | | | | | | 0 | | | |
| EVLEM | + 6.3 x CIM1 | HDX n.d. | | | | | | | | | | | | | | 0 | | | |
| | + 63 x CIM1 | HDX n.d. | | | | | | | | | | | | | | 0 | | | |
| 864-868 | apo | HDX n.d. | | | | | | | | | | | | | | 0 | | | |
| VLEML | + 6.3 x CIM1 | HDX n.d. | | | | | | | | | | | | | | 0 | | | |
| | + 63 x CIM1 | HDX n.d. | | | | | | | | | | | | | | 0 | | | |
| 866-883 | apo | 3.5 | 0.2 | 0.0098 | 0.0018 | | | | | 3.93 | 0.13 | 11.4 | 1.9 | 7.4 | 0.2 | 43.7 | 1.4 | < 0.0001 | > 99 |
| EMLQRFKDSTIKREREVF | + 6.3 x CIM1 | 2.67 | 0.14 | 0.041 | 0.006 | | | | | 3.68 | 0.11 | 14 | 2 | 6.36 | 0.10 | 37.4 | 0.6 | < 0.0001 | > 99 |
| | + 63 x CIM1 | 3.2 | 0.2 | 0.023 | 0.005 | | | | | 3.68 | 0.17 | 13 | 3 | 6.88 | 0.16 | 40.4 | 0.9 | < 0.0001 | > 99 |
| 868-883 | apo | 3.02 | 0.20 | 0.024 | 0.006 | | | | | 4.12 | 0.17 | 18 | 7 | 7.14 | 0.15 | 47.6 | 1.0 | < 0.0001 | > 99 |
| LQRFKDSTIKREREVF | + 6.3 x CIM1 | 2.61 | 0.09 | 0.042 | 0.003 | | | | | 3.99 | 0.06 | 19 | 3 | 6.60 | 0.06 | 44.0 | 0.4 | < 0.0001 | > 99 |
| | + 63 x CIM1 | 2.9 | 0.2 | 0.016 | 0.004 | | | | | 4.07 | 0.16 | 15 | 4 | 6.95 | 0.20 | 46.3 | 1.3 | < 0.0001 | > 99 |
| 868-886 | apo | 3.4 | 0.3 | 0.038 | 0.010 | | | | | 3.91 | 0.20 | 30[a] | | 7.31 | 0.20 | 40.6 | 1.1 | 0.8842 | 90 |
| LQRFKDSTIKREREVFNCM | + 6.3 x CIM1 | 2.78 | 0.09 | 0.037 | 0.003 | | | | | 4.25 | 0.07 | 16.0 | 1.7 | 7.03 | 0.07 | 39.0 | 0.4 | < 0.0001 | > 99 |
| | + 63 x CIM1 | 3.17 | 0.17 | 0.035 | 0.006 | | | | | 3.83 | 0.12 | 30[a] | | 7.01 | 0.13 | 38.9 | 0.7 | 0.6838 | 90 |
| 869-883 | apo | 2.79 | 0.15 | 0.025 | 0.005 | | | | | 4.53 | 0.13 | 14 | 2 | 7.32 | 0.11 | 52.3 | 0.8 | < 0.0001 | > 99 |
| QRFKDSTIKREREVF | + 6.3 x CIM1 | 2.69 | 0.20 | 0.043 | 0.008 | | | | | 4.20 | 0.15 | 17 | 5 | 6.89 | 0.14 | 49.2 | 1.0 | < 0.0001 | > 99 |
| | + 63 x CIM1 | 2.79 | 0.19 | 0.044 | 0.009 | | | | | 3.99 | 0.13 | 30[a] | | 6.78 | 0.14 | 48.4 | 1.0 | 0.9945 | 91 |
| 886-893 | apo | 3 | 3 | 0.0012 | 0.0015 | | | | | | | | | 3 | 3 | 50 | 50 | - | > 99 |

| | | Best-fit values | | | | | | | | | | | | | | | | | |
|---|---|---|---|---|---|---|---|---|---|---|---|---|---|---|---|---|---|---|---|
| Amino acids/sequence | state | $ND_1$ [Da] | $\delta ND_1$ | $k_1$ [min$^{-1}$] | $\delta k_1$ | $ND_2$ [Da] | $\delta ND_2$ | $k_2$ [min$^{-1}$] | $\delta k_2$ | $ND_3$ [Da] | $\delta ND_3$ | $k_3$ [min$^{-1}$] | $\delta k_3$ | ND [Da] | δND | $D_{\%}$ | $\delta D_{\%}$ | P-value | % AIC |
| MLRNLFEE | + 6.3 x CIM1 | 0.11 | 0.08 | 0.1[a] | | | | | | 0.11 | 0.05 | 30[a] | | 0.22 | 0.05 | 3.2 | 0.8 | - | 74 |
| | + 63 x CIM1 | 0.33 | 0.09 | 0.005 | 0.003 | | | | | 0.048 | 0.020 | 30[a] | | 0.38 | 0.09 | 5.4 | 1.4 | 0.3906 | 86 |
| 887-893 | apo | 1.7 | 0.2 | 0.0034 | 0.0007 | | | | | | | | | 1.7 | 0.2 | 28.8 | 3.8 | - | > 99 |
| LRNLFEE | + 6.3 x CIM1 | 0.55 | 0.09 | 0.016 | 0.007 | | | | | | | | | 0.55 | 0.09 | 9.2 | 1.5 | - | > 99 |
| | + 63 x CIM1 | 0.43 | 0.06 | 0.007 | 0.002 | | | | | | | | | 0.43 | 0.06 | 7.2 | 1.0 | 0.2636 | 92 |
| 887-894 | apo | 2.8 | 1.0 | 0.0029 | 0.0016 | | | | | | | | | 2.8 | 1.0 | 40 | 15 | - | > 99 |
| LRNLFEEY | + 6.3 x CIM1 | 1.5 | 1.9 | 0.007 | 0.013 | | | | | | | | | 1.5 | 1.9 | 20 | 30 | - | > 99 |
| | + 63 x CIM1 | 1.4 | 0.7 | 0.0021 | 0.0016 | | | | | 0.15 | 0.03 | 30[a] | | 1.5 | 0.8 | 22 | 11 | 0.2789 | 82 |
| 888-893 | apo | 0.32 | 0.16 | 0.1[a] | | | | | | 0.19 | 0.12 | 30[a] | | 0.51 | 0.10 | 10.1 | 1.9 | 0.9623 | 86 |
| RNLFEE | + 6.3 x CIM1 | | | | | 0.33 | 0.06 | 1.2 | 0.8 | | | | | 0.33 | 0.06 | 6.7 | 1.1 | 0.3977 | 84 |
| | + 63 x CIM1 | | | | | 0.25 | 0.04 | 7 | 7 | | | | | 0.25 | 0.04 | 5.0 | 0.9 | 0.7118 | 86 |
| 894-904 | apo | 5.8 | 0.3 | 0.015 | 0.003 | 3.3 | 0.2 | 6.1 | 1.4 | | | | | 9.0 | 0.3 | 100 | 3 | < 0.0001 | > 99 |
| YRFFPQYPDKE | + 6.3 x CIM1 | 4.8 | 0.3 | 0.027 | 0.005 | 2.62 | 0.19 | 6.6 | 1.5 | | | | | 7.4 | 0.2 | 83 | 3 | < 0.0001 | > 99 |
| | + 63 x CIM1 | 4.7 | 0.2 | 0.029 | 0.005 | | | | | 1.89 | 0.20 | 10 | 4 | 6.56 | 0.16 | 72.9 | 1.8 | < 0.0001 | > 99 |
| 894-905 | apo | 5.4 | 0.6 | 0.008 | 0.003 | 4.8 | 0.5 | 1.1 | 0.3 | | | | | 10.2 | 0.6 | 102 | 6 | < 0.0001 | > 99 |
| YRFFPQYPDKEL | + 6.3 x CIM1 | 5.9 | 0.3 | 0.038 | 0.005 | 2.4 | 0.2 | 7 | 2 | | | | | 8.4 | 0.2 | 84 | 2 | < 0.0001 | > 99 |
| | + 63 x CIM1 | 6.0 | 0.3 | 0.023 | 0.004 | | | | | 1.9 | 0.2 | 11 | 6 | 7.9 | 0.2 | 79 | 2 | < 0.0001 | > 99 |
| 894-909 | apo | 8.1 | 0.4 | 0.032 | 0.005 | 3.2 | 0.4 | 5.1 | 1.8 | | | | | 11.2 | 0.3 | 80.1 | 1.8 | < 0.0001 | > 99 |
| YRFFPQYPDKELHITA | + 6.3 x CIM1 | 7.61 | 0.12 | 0.0326 | 0.0015 | 2.90 | 0.09 | 5.4 | 0.5 | | | | | 10.50 | 0.09 | 75.0 | 0.7 | < 0.0001 | > 99 |
| | + 63 x CIM1 | 7.4 | 0.3 | 0.026 | 0.003 | 2.0 | 0.2 | 6 | 2 | | | | | 9.4 | 0.2 | 67.2 | 1.5 | < 0.0001 | > 99 |
| 912-917 | apo | HDX n.d. | | | | | | | | | | | | | | 0 | | | |
| FGGIIE | + 6.3 x CIM1 | HDX n.d. | | | | | | | | | | | | | | 0 | | | |
| | + 63 x CIM1 | HDX n.d. | | | | | | | | | | | | | | 0 | | | |
| 912-919 | apo | 2.44 | 0.17 | 0.0075 | 0.0012 | | | | | 0.18 | 0.06 | 30[a] | | 2.63 | 0.17 | 38 | 2 | 0.7 | 89 |
| FGGIIEKG | + 6.3 x CIM1 | 1.13 | 0.10 | 0.016 | 0.004 | | | | | | | | | 1.13 | 0.10 | 16.1 | 1.5 | 0.4384 | 98 |
| | + 63 x CIM1 | peptide n.d. | | | | | | | | | | | | | | | | | |
| 912-922 | apo | 4.47 | 0.13 | 0.0170 | 0.0016 | 1.23 | 0.10 | 6.4 | 1.6 | | | | | 5.70 | 0.11 | 57.0 | 1.1 | < 0.0001 | > 99 |
| FGGIIEKGLVT | + 6.3 x CIM1 | 3.93 | 0.11 | 0.023 | 0.002 | 1.21 | 0.07 | 7.3 | 1.4 | | | | | 5.14 | 0.11 | 51.4 | 1.1 | < 0.0001 | > 99 |
| | + 63 x CIM1 | 4.10 | 0.09 | 0.0186 | 0.0014 | 1.20 | 0.07 | 7.9 | 1.5 | | | | | 5.30 | 0.07 | 53.0 | 0.7 | < 0.0001 | > 99 |
| 912-924 | apo | 3.87 | 0.15 | 0.017 | 0.002 | | | | | 3.76 | 0.11 | 12.7 | 2.0 | 7.63 | 0.13 | 63.6 | 1.1 | < 0.0001 | > 99 |
| FGGIIEKGLVTYM | + 6.3 x CIM1 | 3.90 | 0.12 | 0.024 | 0.003 | | | | | 3.51 | 0.08 | 14.5 | 2.0 | 7.41 | 0.12 | 61.7 | 1.0 | < 0.0001 | > 99 |
| | + 63 x CIM1 | 3.99 | 0.12 | 0.0145 | 0.0014 | | | | | 3.72 | 0.08 | 12.9 | 1.5 | 7.71 | 0.11 | 64.3 | 0.9 | < 0.0001 | > 99 |
| 915-923 | apo | 4.06 | 0.13 | 0.0161 | 0.0017 | | | | | 2.65 | 0.09 | 11.5 | 2.0 | 6.71 | 0.12 | 83.8 | 1.5 | < 0.0001 | > 99 |
| IIEKGLVTY | + 6.3 x CIM1 | 3.31 | 0.12 | 0.024 | 0.003 | | | | | 2.66 | 0.08 | 15 | 3 | 5.97 | 0.12 | 74.6 | 1.5 | < 0.0001 | > 99 |
| | + 63 x CIM1 | peptide n.d. | | | | | | | | | | | | | | | | | |
| 918-923 | apo | 2.59 | 0.06 | 0.024 | 0.002 | | | | | 2.97 | 0.05 | 15.7 | 1.9 | 5.55 | 0.05 | 111.1 | 0.9 | < 0.0001 | > 99 |
| KGLVTY | + 6.3 x CIM1 | 2.43 | 0.06 | 0.026 | 0.002 | | | | | 3.01 | 0.04 | 15.9 | 1.6 | 5.43 | 0.06 | 108.7 | 1.2 | < 0.0001 | > 99 |
| | + 63 x CIM1 | 2.64 | 0.10 | 0.020 | 0.003 | | | | | 2.91 | 0.08 | 13.3 | 1.9 | 5.56 | 0.08 | 111.1 | 1.6 | < 0.0001 | > 99 |
| 923-927 | apo | 2.24 | 0.11 | 0.049 | 0.006 | 1.73 | 0.09 | 8.0 | 1.4 | | | | | 3.96 | 0.06 | 99.1 | 1.5 | < 0.0001 | > 99 |
| YMALG | + 6.3 x CIM1 | 2.00 | 0.10 | 0.050 | 0.006 | 1.91 | 0.07 | 6.7 | 0.8 | | | | | 3.91 | 0.07 | 97.7 | 1.7 | < 0.0001 | > 99 |
| | + 63 x CIM1 | peptide n.d. | | | | | | | | | | | | | | | | | |
| 923-928 | apo | 3.19 | 0.18 | 0.038 | 0.006 | 1.76 | 0.15 | 6.7 | 1.8 | | | | | 4.94 | 0.10 | 99 | 2 | < 0.0001 | > 99 |
| YMALGL | + 6.3 x CIM1 | 3.09 | 0.11 | 0.040 | 0.004 | 1.62 | 0.09 | 7.2 | 1.2 | | | | | 4.71 | 0.08 | 94.2 | 1.7 | < 0.0001 | > 99 |
| | + 63 x CIM1 | 3.29 | 0.13 | 0.037 | 0.004 | 1.40 | 0.11 | 8 | 2 | | | | | 4.69 | 0.08 | 93.8 | 1.7 | < 0.0001 | > 99 |
| 923-930 | apo | 3.0 | 0.2 | 0.022 | 0.006 | 1.85 | 0.19 | 6 | 2 | | | | | 4.80 | 0.18 | 69 | 3 | < 0.0001 | > 99 |
| YMALGLAL | + 6.3 x CIM1 | 2.60 | 0.16 | 0.038 | 0.006 | 1.84 | 0.12 | 5.4 | 1.0 | | | | | 4.43 | 0.12 | 63.3 | 1.7 | < 0.0001 | > 99 |
| | + 63 x CIM1 | 3.17 | 0.13 | 0.025 | 0.003 | 1.45 | 0.11 | 9 | 2 | | | | | 4.62 | 0.10 | 66.0 | 1.4 | < 0.0001 | > 99 |
| 924-929 | apo | 3.12 | 0.20 | 0.037 | 0.005 | 0.86 | 0.20 | 2.3 | 1.3 | | | | | 3.98 | 0.06 | 79.6 | 1.2 | < 0.0001 | > 99 |
| MALGLA | + 6.3 x CIM1 | 3.46 | 0.20 | 0.040 | 0.006 | | | | | 0.40 | 0.12 | 30[a] | | 3.86 | 0.17 | 77 | 3 | 0.0005 | > 99 |
| | + 63 x CIM1 | 2.61 | 0.10 | 0.0140 | 0.0015 | 1.49 | 0.10 | 0.38[b] | 0.05 | | | | | 4.10 | 0.07 | 81.9 | 1.3 | < 0.0001 | > 99 |
| 924-930 | apo | 2.07 | 0.09 | 0.0145 | 0.0019 | 1.84 | 0.10 | 0.81[b] | 0.08 | | | | | 3.91 | 0.06 | 65.2 | 1.1 | < 0.0001 | > 99 |
| MALGLAL | + 6.3 x CIM1 | 2.63 | 0.06 | 0.0388 | 0.0020 | 0.96 | 0.06 | 2.5 | 0.4 | | | | | 3.59 | 0.03 | 59.8 | 0.5 | < 0.0001 | > 99 |
| | + 63 x CIM1 | 2.82 | 0.11 | 0.032 | 0.003 | 0.77 | 0.11 | 2.9 | 1.1 | | | | | 3.59 | 0.06 | 59.8 | 0.9 | < 0.0001 | > 99 |
| 925-930 | apo | 1.51 | 0.11 | 0.012 | 0.002 | 1.22 | 0.10 | 0.90[b] | 0.15 | | | | | 2.72 | 0.08 | 54.4 | 1.7 | < 0.0001 | > 99 |
| ALGLAL | + 6.3 x CIM1 | 1.81 | 0.20 | 0.1[a] | | | | | | 0.29 | 0.13 | 30[a] | | 2.10 | 0.13 | 42 | 3 | 0.1974 | 74 |
| | + 63 x CIM1 | 1.45 | 0.13 | 0.0051 | 0.0012 | 1.29 | 0.05 | 0.48[c] | 0.05 | | | | | 2.74 | 0.14 | 55 | 3 | < 0.0001 | > 99 |
| 928-934 | apo | HDX n.d. | | | | | | | | | | | | | | 0 | | | |
| LALRYVL | + 6.3 x CIM1 | HDX n.d. | | | | | | | | | | | | | | 0 | | | |
| | + 63 x CIM1 | HDX n.d. | | | | | | | | | | | | | | 0 | | | |
| 929-934 | apo | HDX n.d. | | | | | | | | | | | | | | 0 | | | |
| ALRYVL | + 6.3 x CIM1 | HDX n.d. | | | | | | | | | | | | | | 0 | | | |
| | + 63 x CIM1 | HDX n.d. | | | | | | | | | | | | | | 0 | | | |
| 929-936 | apo | HDX n.d. | | | | | | | | | | | | | | 0 | | | |
| ALRYVLEA | + 6.3 x CIM1 | HDX n.d. | | | | | | | | | | | | | | 0 | | | |
| | + 63 x CIM1 | HDX n.d. | | | | | | | | | | | | | | 0 | | | |
| 931-936 | apo | HDX n.d. | | | | | | | | | | | | | | 0 | | | |
| RYVLEA | + 6.3 x CIM1 | HDX n.d. | | | | | | | | | | | | | | 0 | | | |
| | + 63 x CIM1 | peptide n.d. | | | | | | | | | | | | | | | | | |
| 935-945 | apo | 1.01 | 0.13 | 0.026 | 0.012 | 5.25 | 0.11 | 9.0 | 0.7 | | | | | 6.26 | 0.10 | 69.5 | 1.1 | < 0.0001 | > 99 |
| EALRKPFGSKM | + 6.3 x CIM1 | 0.96 | 0.10 | 0.031 | 0.009 | 5.18 | 0.07 | 8.6 | 0.4 | | | | | 6.13 | 0.08 | 68.1 | 0.9 | < 0.0001 | > 99 |
| | + 63 x CIM1 | 0.7 | 0.3 | 0.1[a] | | 5.0 | 0.2 | 9.0 | 1.5 | | | | | 5.65 | 0.15 | 62.8 | 1.6 | 0.0368 | 71 |
| 936-945 | apo | 1.01 | 0.13 | 0.021 | 0.010 | 5.37 | 0.11 | 8.7 | 0.6 | | | | | 6.38 | 0.11 | 79.8 | 1.4 | < 0.0001 | > 99 |
| ALRKPFGSKM | + 6.3 x CIM1 | 1.10 | 0.10 | 0.032 | 0.008 | 5.21 | 0.07 | 9.4 | 0.5 | | | | | 6.32 | 0.08 | 79.0 | 1.0 | < 0.0001 | > 99 |
| | + 63 x CIM1 | 0.82 | 0.10 | 0.019 | 0.008 | 5.42 | 0.08 | 7.6 | 0.4 | | | | | 6.24 | 0.09 | 78.1 | 1.1 | < 0.0001 | > 99 |
| 937-947 | apo | 0.92 | 0.08 | 0.027 | 0.008 | 5.02 | 0.07 | 8.7 | 0.4 | | | | | 5.94 | 0.05 | 65.9 | 0.6 | < 0.0001 | > 99 |
| LRKPFGSKMYY | + 6.3 x CIM1 | 0.90 | 0.14 | 0.039 | 0.016 | 4.87 | 0.10 | 9.2 | 0.7 | | | | | 5.76 | 0.10 | 64.0 | 1.1 | 0.0004 | > 99 |
| | + 63 x CIM1 | peptide n.d. | | | | | | | | | | | | | | | | | |
| 946-951 | apo | HDX n.d. | | | | | | | | | | | | | | 0 | | | |

| Amino acids/sequence | state | Best-fit values |  |  |  |  |  |  |  |  |  |  |  |  |  |  |  |  |  |
|---|---|---|---|---|---|---|---|---|---|---|---|---|---|---|---|---|---|---|---|
| | | $ND_1$ [Da] | $\delta ND_1$ | $k_1$ [min$^{-1}$] | $\delta k_1$ | $ND_2$ [Da] | $\delta ND_2$ | $k_2$ [min$^{-1}$] | $\delta k_2$ | $ND_3$ [Da] | $\delta ND_3$ | $k_3$ [min$^{-1}$] | $\delta k_3$ | ND [Da] | δND | $D_\%$ | $\delta D_\%$ | P-value | % AIC |
| YYFGIA | + 6.3 x CIM1 | HDX n.d. |  |  |  |  |  |  |  |  |  |  |  |  |  | 0 |  |  |  |
|  | + 63 x CIM1 | HDX n.d. |  |  |  |  |  |  |  |  |  |  |  |  |  | 0 |  |  |  |
| 946-952 | apo | HDX n.d. |  |  |  |  |  |  |  |  |  |  |  |  |  | 0 |  |  |  |
| YYFGIAA | + 6.3 x CIM1 | HDX n.d. |  |  |  |  |  |  |  |  |  |  |  |  |  | 0 |  |  |  |
|  | + 63 x CIM1 | HDX n.d. |  |  |  |  |  |  |  |  |  |  |  |  |  | 0 |  |  |  |
| 946-953 | apo | HDX n.d. |  |  |  |  |  |  |  |  |  |  |  |  |  | 0 |  |  |  |
| YYFGIAAL | + 6.3 x CIM1 | HDX n.d. |  |  |  |  |  |  |  |  |  |  |  |  |  | 0 |  |  |  |
|  | + 63 x CIM1 | HDX n.d. |  |  |  |  |  |  |  |  |  |  |  |  |  | 0 |  |  |  |
| 948-952 | apo | HDX n.d. |  |  |  |  |  |  |  |  |  |  |  |  |  | 0 |  |  |  |
| FGIAA | + 6.3 x CIM1 | HDX n.d. |  |  |  |  |  |  |  |  |  |  |  |  |  | 0 |  |  |  |
|  | + 63 x CIM1 | HDX n.d. |  |  |  |  |  |  |  |  |  |  |  |  |  | 0 |  |  |  |
| 948-953 | apo | HDX n.d. |  |  |  |  |  |  |  |  |  |  |  |  |  | 0 |  |  |  |
| FGIAAL | + 6.3 x CIM1 | HDX n.d. |  |  |  |  |  |  |  |  |  |  |  |  |  | 0 |  |  |  |
|  | + 63 x CIM1 | HDX n.d. |  |  |  |  |  |  |  |  |  |  |  |  |  | 0 |  |  |  |
| 952-966 | apo | 3.75 | 0.10 | 0.0073 | 0.0005 |  |  |  |  | 3.62 | 0.04 | 15.9 | 1.5 | 7.38 | 0.10 | 56.7 | 0.8 | < 0.0001 | > 99 |
| ALDRFKNRLKDYPQY | + 6.3 x CIM1 | 5 | 5 | 0.004 | 0.005 |  |  |  |  | 3.62 | 0.08 | 14.0 | 1.8 | 9 | 5 | 70 | 40 | 0.0133 | 87 |
|  | + 63 x CIM1 | 3.7 | 0.3 | 0.0060 | 0.0011 |  |  |  |  | 3.74 | 0.09 | 12.1 | 1.5 | 7.4 | 0.3 | 57 | 2 | < 0.0001 | > 99 |
| 953-966 | apo | 3.90 | 0.20 | 0.0055 | 0.0006 |  |  |  |  | 3.54 | 0.06 | 14.6 | 1.7 | 7.4 | 0.2 | 62.0 | 1.7 | < 0.0001 | > 99 |
| LDRFKNRLKDYPQY | + 6.3 x CIM1 | 3.1 | 0.5 | 0.009 | 0.003 |  |  |  |  | 3.44 | 0.05 | 14.4 | 1.2 | 6.5 | 0.5 | 54 | 4 | < 0.0001 | > 99 |
|  | + 63 x CIM1 | 3.41 | 0.14 | 0.0074 | 0.0008 |  |  |  |  | 3.28 | 0.06 | 15 | 2 | 6.69 | 0.14 | 55.7 | 1.2 | < 0.0001 | > 99 |
| 953-972 | apo | 8.8 | 0.4 | 0.0115 | 0.0014 |  |  |  |  | 4.7 | 0.2 | 12 | 3 | 13.5 | 0.4 | 75 | 2 | < 0.0001 | > 99 |
| LDRFKNRLKDYPQYCQHLAS | + 6.3 x CIM1 | 6.9 | 0.3 | 0.026 | 0.003 |  |  |  |  | 4.25 | 0.19 | 15 | 4 | 11.2 | 0.3 | 62.1 | 1.4 | < 0.0001 | > 99 |
|  | + 63 x CIM1 | peptide n.d. |  |  |  |  |  |  |  |  |  |  |  |  |  |  |  |  |  |
| 954-966 | apo | 3.61 | 0.17 | 0.0076 | 0.0008 |  |  |  |  | 3.42 | 0.07 | 14.2 | 2.0 | 7.03 | 0.17 | 63.9 | 1.5 | < 0.0001 | > 99 |
| DRFKNRLKDYPQY | + 6.3 x CIM1 | 4.0 | 1.7 | 0.005 | 0.003 |  |  |  |  | 3.47 | 0.04 | 12.8 | 0.8 | 7.475 | 1.68 | 66 | 15 | < 0.0001 | > 99 |
|  | + 63 x CIM1 | 3.38 | 0.11 | 0.0073 | 0.0006 |  |  |  |  | 3.22 | 0.05 | 15.5 | 1.7 | 6.60 | 0.11 | 60.0 | 1.0 | < 0.0001 | > 99 |
| 954-972 | apo | 9.1 | 0.4 | 0.0121 | 0.0016 | 5.3 | 0.3 | 8.8 | 1.7 |  |  |  |  | 14.4 | 0.4 | 85 | 2 | < 0.0001 | > 99 |
| DRFKNRLKDYPQYCQHLAS | + 6.3 x CIM1 | 7.64 | 0.16 | 0.0279 | 0.0018 |  |  |  |  | 4.50 | 0.11 | 15 | 2 | 12.14 | 0.14 | 71.4 | 0.8 | < 0.0001 | > 99 |
|  | + 63 x CIM1 | 9.2 | 0.5 | 0.014 | 0.002 |  |  |  |  | 4.7 | 0.3 | 12 | 4 | 13.9 | 0.4 | 82 | 3 | < 0.0001 | > 99 |
| 973-985 | apo | 6.5 | 0.6 | 0.036 | 0.007 | 4.1 | 0.6 | 2.8 | 1.1 |  |  |  |  | 10.6 | 0.2 | 96 | 2 | < 0.0001 | > 99 |
| ISHFMQFPHHLQE | + 6.3 x CIM1 | 6.4 | 0.5 | 0.071 | 0.012 | 3.2 | 0.4 | 3.5 | 1.2 |  |  |  |  | 9.6 | 0.3 | 88 | 3 | 0.0038 | 96 |
|  | + 63 x CIM1 | 6.7 | 0.5 | 0.047 | 0.007 | 3.1 | 0.4 | 3.5 | 1.3 |  |  |  |  | 9.7 | 0.2 | 89 | 2 | < 0.0001 | > 99 |
| 974-985 | apo | 7.4 | 0.5 | 0.1[a] |  |  |  |  |  | 1.9 | 0.4 | 30[a] |  | 9.3 | 0.3 | 93 | 3 | 0.308 | 76 |
| SHFMQFPHHLQE | + 6.3 x CIM1 | 5.8 | 0.2 | 0.076 | 0.008 | 2.90 | 0.19 | 4.4 | 0.8 |  |  |  |  | 8.75 | 0.15 | 87.5 | 1.5 | < 0.0001 | > 99 |
|  | + 63 x CIM1 | 6.2 | 0.4 | 0.049 | 0.008 | 2.5 | 0.4 | 4.3 | 1.8 |  |  |  |  | 8.7 | 0.2 | 87 | 2 | 0.0002 | > 99 |
| 977-986 | apo | 10 | 12 | 0.0010 | 0.0015 | 3.68 | 0.17 | 0.29[b] | 0.05 |  |  |  |  | 13 | 12 | 170 | 150 | < 0.0001 | > 99 |
| MQFPHHLQEY | + 6.3 x CIM1 | 4.2 | 0.2 | 0.067 | 0.009 |  |  |  |  | 0.28 | 0.13 | 30[a] |  | 4.53 | 0.17 | 57 | 2 | 0.0524 | 60 |
|  | + 63 x CIM1 | 3.8 | 0.6 | 0.0046 | 0.0018 | 2.5 | 0.2 | 0.31[b] | 0.09 |  |  |  |  | 6.3 | 0.7 | 78 | 9 | < 0.0001 | > 99 |
| 978-985 | apo | 4.3 | 0.3 | 0.057 | 0.012 |  |  |  |  | 0.9 | 0.2 | 30[a] |  | 5.2 | 0.2 | 87 | 3 | 0.3399 | 82 |
| QFPHHLQE | + 6.3 x CIM1 | 3.77 | 0.11 | 0.061 | 0.004 | 0.94 | 0.09 | 3.5 | 1.0 |  |  |  |  | 4.71 | 0.06 | 78.5 | 1.1 | < 0.0001 | > 99 |
|  | + 63 x CIM1 | peptide n.d. |  |  |  |  |  |  |  |  |  |  |  |  |  |  |  |  |  |
| 979-985 | apo | 3.5 | 0.3 | 0.061 | 0.015 |  |  |  |  | 1.1 | 0.2 | 30[a] |  | 4.60 | 0.19 | 92 | 4 | 0.3162 | 82 |
| FPHHLQE | + 6.3 x CIM1 | 3.2 | 0.3 | 0.051 | 0.011 | 1.2 | 0.2 | 4 | 3 |  |  |  |  | 4.4 | 0.2 | 88 | 4 | 0.0045 | 95 |
|  | + 63 x CIM1 | 3.6 | 0.3 | 0.1[a] |  |  |  |  |  | 0.5 | 0.2 | 30[a] |  | 4.08 | 0.18 | 82 | 4 | 0.2211 | 72 |
| 986-992 | apo | 5.70 | 0.11 | 0.064 | 0.005 |  |  |  |  |  |  |  |  | 5.70 | 0.11 | 94.9 | 1.9 | 0.142 | 64 |
| YIEYGQQ | + 6.3 x CIM1 | 5.93 | 0.19 | 0.060 | 0.006 |  |  |  |  |  |  |  |  | 5.93 | 0.19 | 99 | 3 | 0.1742 | 92 |
|  | + 63 x CIM1 | 5.12 | 0.06 | 0.071 | 0.003 |  |  |  |  |  |  |  |  | 5.12 | 0.06 | 85.3 | 1.1 | 0.7593 | 89 |
| 986-999 | apo | 4.7 | 0.2 | 0.061 | 0.007 | 6.16 | 0.19 | 4.6 | 0.4 |  |  |  |  | 10.85 | 0.11 | 98.6 | 1.0 | < 0.0001 | > 99 |
| YIEYGQQSRDPPVK | + 6.3 x CIM1 | 4.29 | 0.12 | 0.060 | 0.004 | 6.17 | 0.10 | 4.52 | 0.20 |  |  |  |  | 10.46 | 0.08 | 95.1 | 0.7 | < 0.0001 | > 99 |
|  | + 63 x CIM1 | 4.6 | 0.2 | 0.056 | 0.006 | 5.93 | 0.18 | 4.5 | 0.4 |  |  |  |  | 10.52 | 0.11 | 95.6 | 1.0 | < 0.0001 | > 99 |
| 987-999 | apo | 4.8 | 0.4 | 0.062 | 0.011 | 6.5 | 0.3 | 4.7 | 0.6 |  |  |  |  | 11.36 | 0.19 | 113.6 | 1.9 | < 0.0001 | > 99 |
| IEYGQQSRDPPVK | + 6.3 x CIM1 | 4.23 | 0.14 | 0.076 | 0.007 | 6.54 | 0.12 | 4.6 | 0.2 |  |  |  |  | 10.77 | 0.09 | 107.7 | 0.9 | < 0.0001 | > 99 |
|  | + 63 x CIM1 | 4.4 | 0.3 | 0.068 | 0.012 | 6.3 | 0.3 | 4.4 | 0.5 |  |  |  |  | 10.69 | 0.16 | 106.9 | 1.6 | < 0.0001 | > 99 |
| 988-999 | apo | 3.7 | 0.3 | 0.056 | 0.010 | 5.2 | 0.2 | 4.4 | 0.6 |  |  |  |  | 8.92 | 0.13 | 99.2 | 1.5 | < 0.0001 | > 99 |
| EYGQQSRDPPVK | + 6.3 x CIM1 | 3.32 | 0.13 | 0.062 | 0.006 | 5.25 | 0.11 | 4.1 | 0.2 |  |  |  |  | 8.57 | 0.08 | 95.2 | 0.9 | < 0.0001 | > 99 |
|  | + 63 x CIM1 | 3.82 | 0.18 | 0.063 | 0.007 | 4.87 | 0.15 | 5.2 | 0.5 |  |  |  |  | 8.69 | 0.10 | 96.6 | 1.1 | < 0.0001 | > 99 |
| 989-999 | apo | 2.2 | 0.2 | 0.1[a] |  | 7.6 | 0.2 | 4.1 | 0.3 |  |  |  |  | 9.76 | 0.10 | 122.1 | 1.3 | 0.2355 | 77 |
| YGQQSRDPPVK | + 6.3 x CIM1 | 2.47 | 0.13 | 0.092 | 0.015 | 7.28 | 0.12 | 4.29 | 0.18 |  |  |  |  | 9.75 | 0.08 | 121.8 | 1.1 | < 0.0001 | > 99 |
|  | + 63 x CIM1 | 2.63 | 0.14 | 0.15 | 0.04 | 6.88 | 0.16 | 4.5 | 0.2 |  |  |  |  | 9.50 | 0.06 | 118.8 | 0.8 | < 0.0001 | > 99 |

[a] Fixed values, according to (Gemmecker et al., 1993; Hoofnagle et al., 2001; Mandell et al., 2001; Rutkowska-Wlodarczyk et al., 2008).

[b] exceptions to the classification that $k_1 \leq 0.1$, $0.1 < k_2 < 10$ and $10 < k_3 \leq 30$, due to lack of space in appropriate column.

### Supplementary Table T3 to Fig 2f.

Kinetic parameters of hydrogen-deuterium exchange for CNOT1 at 2 µM in the presence of SD10 at 20 µM.

| | | Best-fit values | | | | | | | | | | | | | | | | | |
|---|---|---|---|---|---|---|---|---|---|---|---|---|---|---|---|---|---|---|---|
| Amino acids/sequence | state | $ND_1$ [Da] | $\delta ND_1$ | $k_1$ [min$^{-1}$] | $\delta k_1$ | $ND_2$ [Da] | $\delta ND_2$ | $k_2$ [min$^{-1}$] | $\delta k_2$ | $ND_3$ [Da] | $\delta ND_3$ | $k_3$ [min$^{-1}$] | $\delta k_3$ | ND [Da] | δND | $D_{\%}$ | $\delta D_{\%}$ | P-value | % AIC |
| 800-819 | apo | | | | | | | | | 17.06 | 0.07 | 32 | 8 | 17.06 | 0.07 | 100.4 | 0.4 | - | 85 |
| NNDPFVQRKLGTSGLNQPTF | + 10 x SD10 | | | | | | | | | 15.08 | 0.12 | 31 | 14 | 15.08 | 0.12 | 88.7 | 0.7 | 0.7693 | 89 |
| 800-824 | apo | | | | | | | | | 22.03 | 0.13 | 32 | 12 | 22.03 | 0.13 | 100.1 | 0.6 | - | 85 |
| NNDPFVQRKLGTSGLNQPTFQQTDL | + 10 x SD10 | | | | | | | | | 19.12 | 0.15 | 36 | 23 | 19.12 | 0.15 | 86.9 | 0.7 | - | 90 |
| 805-824 | apo | | | | | | | | | 17.67 | 0.14 | 37 | 23 | 17.67 | 0.14 | 98.2 | 0.8 | - | 88 |
| VQRKLGTSGLNQPTFQQTDL | + 10 x SD10 | | | | | | | | | 16.10 | 0.14 | 32 | 18 | 16.10 | 0.14 | 89.4 | 0.8 | - | 90 |
| 820-824 | apo | | | | | | | | | 3.81 | 0.02 | 23 | 3 | 3.81 | 0.02 | 95.3 | 0.5 | - | 87 |
| QQTDL | + 10 x SD10 | | | | | | | | | 3.69 | 0.03 | 31 | 16 | 3.69 | 0.03 | 92.3 | 0.8 | - | 91 |
| 825-838 | apo | | | | | | | | | 12.11 | 0.08 | 14.0 | 0.7 | 12.11 | 0.08 | 100.9 | 0.6 | - | > 99 |
| SQVWPEANQHFSKE | + 10 x SD10 | | | | | | | | | 11.26 | 0.08 | 14.7 | 1.0 | 11.26 | 0.08 | 93.8 | 0.7 | | > 99 |
| 839-846 | apo | 5.11 | 0.14 | 0.031 | 0.003 | | | | | | | | | 5.11 | 0.14 | 73 | 2 | 0.3348 | 98 |
| IDDEANSY | + 10 x SD10 | 3.0 | 0.4 | 0.020 | 0.008 | | | | | | | | | 3.0 | 0.4 | 44 | 6 | | 93 |
| 847-859 | apo | 3.9 | 0.2 | 0.051 | 0.006 | 2.37 | 0.15 | 7.1 | 1.3 | | | | | 6.28 | 0.14 | 69.7 | 1.5 | < 0.0001 | > 99 |
| FQRIYNHPPHPTM | + 10 x SD10 | 2.6 | 0.2 | 0.13 | 0.06 | 2.3 | 0.2 | 8 | 2 | | | | | 4.89 | 0.16 | 54.3 | 1.7 | 0.0003 | > 99 |
| 848-859 | apo | 3.5 | 0.3 | 0.1[a] | | | | | | 2.09 | 0.18 | 30[a] | | 5.63 | 0.18 | 70 | 2 | 0.2576 | 78 |
| QRIYNHPPHPTM | + 10 x SD10 | | | | | | | | | 5.8 | 0.2 | 12 | 4 | 5.8 | 0.2 | 72 | 3 | 0.1642 | 91 |
| 866-883 | apo | 2.46 | 0.11 | 0.045 | 0.006 | | | | | 2.84 | 0.07 | 30[a] | | 5.29 | 0.09 | 31.1 | 0.5 | 0.835 | > 99 |
| EMLQRFKDSTIKREREVF | + 10 x SD10 | 2.2 | 0.4 | 0.1[a] | | | | | | 2.4 | 0.3 | 30[a] | | 4.5 | 0.3 | 26.8 | 1.7 | 0.1758 | 89 |
| 868-883 | apo | 2.01 | 0.12 | 0.037 | 0.007 | | | | | 3.23 | 0.07 | 30[a] | | 5.24 | 0.10 | 35.0 | 0.7 | 0.9064 | 96 |
| LQRFKDSTIKREREVF | + 10 x SD10 | 1.8 | 0.2 | 0.046 | 0.015 | | | | | 2.99 | 0.13 | 30[a] | | 4.75 | 0.18 | 31.6 | 1.2 | 0.5197 | 92 |
| 887-893 | apo | HDX n.d. | | | | | | | | | | | | | | 0 | | | |
| LRNLFEE | + 10 x SD10 | HDX n.d. | | | | | | | | | | | | | | 0 | | | |
| 888-893 | apo | HDX n.d. | | | | | | | | | | | | | | 0 | | | |
| RNLFEE | + 10 x SD10 | peptide n.d. | | | | | | | | | | | | | | | | | |
| 894-909 | apo | 5.88 | 0.12 | 0.035 | 0.002 | 2.27 | 0.09 | 5.9 | 0.6 | | | | | 8.14 | 0.09 | 62.6 | 0.7 | < 0.0001 | > 99 |
| YRFFPQYPDKELHITA | + 10 x SD10 | 4.75 | 0.20 | 0.027 | 0.004 | 1.50 | 0.14 | 9 | 3 | | | | | 6.25 | 0.18 | 48.1 | 1.4 | < 0.0001 | > 99 |
| 901-909 | apo | 4.0 | 0.3 | 0.038 | 0.006 | 1.85 | 0.19 | 4.7 | 1.2 | | | | | 5.89 | 0.18 | 74 | 2 | 0.0003 | > 99 |
| PDKELHITA | + 10 x SD10 | 3.1 | 0.3 | 0.042 | 0.011 | 0.8 | 0.2 | 5 | 4 | | | | | 3.9 | 0.2 | 49 | 3 | 0.0103 | 53 |
| 912-917 | apo | HDX n.d. | | | | | | | | | | | | | | 0 | | | |
| FGGIIE | + 10 x SD10 | HDX n.d. | | | | | | | | | | | | | | 0 | | | |
| 912-923 | apo | 3.5 | 0.2 | 0.014 | 0.002 | | | | | 2.49 | 0.04 | 10.1 | 0.6 | 6.0 | 0.2 | 54.8 | 2.0 | < 0.0001 | > 99 |
| FGGIIEKGLVTY | + 10 x SD10 | 2.62 | 0.16 | 0.020 | 0.005 | | | | | 2.27 | 0.10 | 10.4 | 1.7 | 4.89 | 0.18 | 44.4 | 1.6 | < 0.0001 | > 99 |
| 924-928 | apo | 2.6 | 0.5 | 0.1[a] | | 0.5 | 0.4 | 3 | 5 | | | | | 3.18 | 0.16 | 79.4 | 4.01 | 0.0151 | 87 |
| MALGL | + 10 x SD10 | 2.71 | 0.08 | 0.053 | 0.005 | | | | | | | | | 2.71 | 0.08 | 67.7 | 2.0 | 0.4926 | 86 |
| 924-930 | apo | 2.3 | 0.2 | 0.039 | 0.007 | 0.6 | 0.3 | 1.7 | 1.3 | | | | | 2.99 | 0.07 | 49.8 | 1.1 | < 0.0001 | > 99 |
| MALGLAL | + 10 x SD10 | 2.31 | 0.11 | 0.044 | 0.005 | | | | | 0.14 | 0.07 | 30[a] | | 2.46 | 0.09 | 40.9 | 1.4 | 0.0614 | 53 |
| 929-934 | apo | HDX n.d. | | | | | | | | | | | | | | 0 | | | |
| ALRYVL | + 10 x SD10 | HDX n.d. | | | | | | | | | | | | | | 0 | | | |
| 931-936 | apo | HDX n.d. | | | | | | | | | | | | | | 0 | | | |
| RYVLEA | + 10 x SD10 | HDX n.d. | | | | | | | | | | | | | | 0 | | | |
| 937-945 | apo | 0.7 | 0.2 | 0.1[a] | | 4.62 | 0.18 | 7.6 | 0.9 | | | | | 5.34 | 0.12 | 76.3 | 1.7 | 0.0102 | 91 |
| LRKPFGSKM | + 10 x SD10 | | | | | 5.18 | 0.11 | 4.8 | 0.5 | | | | | 5.18 | 0.11 | 74.1 | 1.6 | 0.3277 | 84 |
| 937-947 | apo | 0.8 | 0.2 | 0.1[a] | | 3.52 | 0.20 | 7.9 | 1.3 | | | | | 4.27 | 0.11 | 47.4 | 1.2 | 0.0132 | 88 |
| LRKPFGSKMYY | + 10 x SD10 | | | | | 4.19 | 0.20 | 5.8 | 1.4 | | | | | 4.19 | 0.20 | 47 | 2 | 0.0785 | 51 |
| 946-951 | apo | HDX n.d. | | | | | | | | | | | | | | 0 | | | |
| YYFGIA | + 10 x SD10 | HDX n.d. | | | | | | | | | | | | | | 0 | | | |
| 948-953 | apo | HDX n.d. | | | | | | | | | | | | | | 0 | | | |
| FGIAAL | + 10 x SD10 | HDX n.d. | | | | | | | | | | | | | | 0 | | | |
| 952-966 | apo | 2.3 | 1.0 | 0.014 | 0.017 | | | | | 3.4 | 0.2 | 10 | 3 | 5.7 | 1.1 | 44 | 8 | 0.0009 | 99 |
| ALDRFKNRLKDYPQY | + 10 x SD10 | 1.7 | 0.5 | 0.021 | 0.020 | | | | | 3.2 | 0.2 | 30[a] | | 4.9 | 0.5 | 38 | 4 | 0.6421 | 93 |
| 952-972 | apo | 4.5 | 0.4 | 0.026 | 0.008 | | | | | 3.1 | 0.3 | 10 | 3 | 7.6 | 0.3 | 39.9 | 1.8 | < 0.0001 | > 99 |
| ALDRFKNRLKDYPQYCQHLAS | + 10 x SD10 | 3.8 | 0.4 | 0.027 | 0.009 | 3.2 | 0.3 | 7 | 2 | | | | | 7.0 | 0.4 | 36.7 | 1.9 | < 0.0001 | > 99 |
| 953-966 | apo | 2.2 | 0.5 | 0.019 | 0.012 | | | | | 3.05 | 0.14 | 30[a] | | 5.3 | 0.5 | 44 | 4 | 0.4455 | 94 |
| LDRFKNRLKDYPQY | + 10 x SD10 | 1.76 | 0.19 | 0.033 | 0.012 | | | | | 2.94 | 0.14 | 10 | 2 | 4.69 | 0.15 | 39.1 | 1.3 | < 0.0001 | > 99 |
| 954-966 | apo | 1.5 | 0.3 | 0.1[a] | | | | | | 2.84 | 0.18 | 30[a] | | 4.4 | 0.2 | 39.9 | 2.0 | 0.3486 | 83 |
| DRFKNRLKDYPQY | + 10 x SD10 | 1.4 | 0.3 | 0.1[a] | | | | | | 2.7 | 0.2 | 30[a] | | 4.1 | 0.2 | 37 | 2 | 0.0944 | 60 |
| 954-972 | apo | 4.87 | 0.15 | 0.026 | 0.003 | | | | | 2.58 | 0.10 | 15 | 4 | 7.45 | 0.14 | 43.8 | 0.8 | < 0.0001 | > 99 |
| DRFKNRLKDYPQYCQHLAS | + 10 x SD10 | 4.1 | 0.3 | 0.061 | 0.013 | | | | | 2.10 | 0.16 | 30[a] | | 6.2 | 0.3 | 36.5 | 1.5 | 0.4141 | 96 |
| 973-985 | apo | 5.8 | 0.3 | 0.054 | 0.007 | 2.57 | 0.19 | 6.1 | 1.5 | | | | | 8.35 | 0.17 | 75.9 | 1.6 | 0.0001 | > 99 |
| ISHFMQFPHHLQE | + 10 x SD10 | 6.2 | 0.4 | 0.45 | 0.13 | | | | | | | | | 6.2 | 0.4 | 56 | 3 | 0.235 | 99 |
| 977-985 | apo | 3.75 | 0.15 | 0.075 | 0.009 | 0.76 | 0.13 | 4.2 | 1.7 | | | | | 4.51 | 0.09 | 64.4 | 1.3 | 0.0005 | > 99 |
| MQFPHHLQE | + 10 x SD10 | 3.59 | 0.16 | 0.40 | 0.09 | | | | | | | | | 2.88 | 0.14 | 41.2 | 2.0 | 0.4846 | 98 |
| 977-986 | apo | 2.88 | 0.14 | 0.1[a] | | | | | | 0.33 | 0.09 | 30[a] | | 3.21 | 0.10 | 40.1 | 1.2 | 0.0943 | 60 |
| MQFPHHLQEY | + 10 x SD10 | 2.11 | 0.10 | 0.10 | 0.02 | | | | | 0.27 | 0.06 | 30[a] | | 2.39 | 0.09 | 29.8 | 1.1 | 0.0033 | 97 |
| 986-999 | apo | 4.17 | 0.14 | 0.070 | 0.006 | 4.93 | 0.11 | 4.4 | 0.3 | | | | | 9.10 | 0.09 | 82.7 | 0.8 | < 0.0001 | > 99 |
| YIEYGQQSRDPPVK | + 10 x SD10 | 3.8 | 0.4 | 0.053 | 0.011 | 4.8 | 0.3 | 3.0 | 0.6 | | | | | 8.55 | 0.20 | 77.7 | 1.8 | 0.0006 | > 99 |
| 987-999 | apo | 4.62 | 0.17 | 0.073 | 0.006 | 4.88 | 0.15 | 4.6 | 0.3 | | | | | 9.51 | 0.10 | 95.1 | 1.0 | < 0.0001 | > 99 |
| IEYGQQSRDPPVK | + 10 x SD10 | 3.8 | 0.4 | 0.09 | 0.02 | 4.8 | 0.3 | 3.3 | 0.6 | | | | | 8.6 | 0.2 | 86 | 2 | 0.0033 | 96 |
| 988-999 | apo | 3.3 | 0.2 | 0.056 | 0.009 | 4.34 | 0.18 | 3.8 | 0.4 | | | | | 7.64 | 0.13 | 84.9 | 1.4 | < 0.0001 | > 99 |
| EYGQQSRDPPVK | + 10 x SD10 | peptide n.d. | | | | | | | | | | | | | | | | | |
| 989-999 | apo | 2.15 | 0.17 | 0.056 | 0.010 | 5.64 | 0.14 | 4.1 | 0.3 | | | | | 7.79 | 0.11 | 97.3 | 1.3 | < 0.0001 | > 99 |
| YGQQSRDPPVK | + 10 x SD10 | 1.8 | 0.3 | 0.1[a] | | 5.1 | 0.3 | 2.9 | 0.5 | | | | | 6.97 | 0.13 | 87.1 | 1.7 | 0.3339 | 89 |

[a] Fixed values, according to (Gemmecker et al., 1993; Hoofnagle et al., 2001; Mandell et al., 2001; Rutkowska-Wlodarczyk et al., 2008).

**Supplementary Table T4 to Fig 2f.**

Kinetic parameters of hydrogen-deuterium exchange for CNOT1 at 2 µM in the presence of the TTP peptide at 3.3 µM or the CIM1 peptide at 126 µM.

| | | Best-fit values | | | | | | | | | | | | | | | | | |
|---|---|---|---|---|---|---|---|---|---|---|---|---|---|---|---|---|---|---|---|
| Amino acids/sequence | state | $ND_1$ [Da] | $\delta ND_1$ | $k_1$ [min$^{-1}$] | $\delta k_1$ | $ND_2$ [Da] | $\delta ND_2$ | $k_2$ [min$^{-1}$] | $\delta k_2$ | $ND_3$ [Da] | $\delta ND_3$ | $k_3$ [min$^{-1}$] | $\delta k_3$ | ND [Da] | δND | $D_{\%}$ | $\delta D_{\%}$ | P-value | % AIC |
| 800-818 | apo | | | | | | | | | 18.91 | 0.12 | 21 | 3 | 18.91 | 0.12 | 118.2 | 0.8 | - | 90 |
| NNDPFVQRKLGTSGLNQPT | + 63 x CIM1 | | | | | | | | | 17.96 | 0.09 | 23 | 3 | 17.96 | 0.09 | 112.3 | 0.6 | - | 90 |
| | + 1.7 x TTP | | | | | | | | | 18.36 | 0.10 | 22 | 3 | 18.36 | 0.10 | 114.8 | 0.6 | - | 90 |
| 800-819 | apo | | | | | | | | | 20.13 | 0.11 | 24 | 4 | 20.13 | 0.11 | 118.4 | 0.6 | - | 90 |
| NNDPFVQRKLGTSGLNQPTF | + 63 x CIM1 | | | | | | | | | 19.29 | 0.12 | 31 | 12 | 19.29 | 0.12 | 113.5 | 0.7 | - | 90 |
| | + 1.7 x TTP | | | | | | | | | 19.67 | 0.10 | 25 | 4 | 19.67 | 0.10 | 115.7 | 0.6 | - | 90 |
| 800-820 | apo | | | | | | | | | 21.20 | 0.14 | 23 | 3 | 21.20 | 0.14 | 117.8 | 0.8 | 0.9391 | 90 |
| NNDPFVQRKLGTSGLNQPTFQ | + 63 x CIM1 | | | | | | | | | 20.41 | 0.11 | 27 | 5 | 20.41 | 0.11 | 113.4 | 0.6 | - | 90 |
| | + 1.7 x TTP | | | | | | | | | 20.72 | 0.14 | 25 | 5 | 20.72 | 0.14 | 115.1 | 0.8 | 0.9765 | 90 |
| 800-824 | apo | | | | | | | | | 26.57 | 0.17 | 23 | 3 | 26.57 | 0.17 | 120.8 | 0.8 | - | 90 |
| NNDPFVQRKLGTSGLNQPTFQQTDL | + 63 x CIM1 | | | | | | | | | 25.63 | 0.16 | 28 | 8 | 25.63 | 0.16 | 116.5 | 0.7 | - | 90 |
| | + 1.7 x TTP | | | | | | | | | 26.8 | 0.7 | 30 | | 26.8 | 0.7 | 122 | 3 | 0.9338 | 98 |
| 820-824 | apo | | | | | | | | | 3.974 | 0.017 | 25 | 4 | 3.974 | 0.017 | 99.4 | 0.4 | 0.9338 | 99 |
| QQTDL | + 63 x CIM1 | | | | | | | | | 3.933 | 0.016 | 26 | 4 | 3.933 | 0.016 | 98.3 | 0.4 | - | 90 |
| | + 1.7 x TTP | | | | | | | | | 3.978 | 0.016 | 26 | 4 | 3.978 | 0.016 | 99.5 | 0.4 | - | 94 |
| 825-835 | apo | | | | | | | | | 10.69 | 0.08 | 15.3 | 1.1 | 10.69 | 0.08 | 118.8 | 0.9 | - | > 99 |
| SQVWPEANQHF | + 63 x CIM1 | | | | | | | | | 9.93 | 0.07 | 12.1 | 0.6 | 9.93 | 0.07 | 110.3 | 0.8 | - | > 99 |
| | + 1.7 x TTP | | | | | | | | | 10.32 | 0.11 | 12.0 | 0.9 | 10.32 | 0.11 | 114.7 | 1.2 | - | > 99 |
| 825-838 | apo | | | | | | | | | 15.13 | 0.10 | 15.1 | 1.0 | 15.13 | 0.10 | 126.1 | 0.8 | - | > 99 |
| SQVWPEANQHFSKE | + 63 x CIM1 | peptide n.d. | | | | | | | | | | | | | | | | | |
| | + 1.7 x TTP | | | | | | | | | 14.39 | 0.10 | 12.5 | 0.7 | 14.39 | 0.10 | 119.9 | 0.9 | - | > 99 |
| 825-847 | apo | 9.3 | 0.4 | 0.055 | 0.005 | | | | | 11.0 | 0.3 | 10.5 | 1.0 | 20.4 | 0.3 | 97.1 | 1.2 | < 0.0001 | > 99 |
| SQVWPEANQHFSKEIDDEANSYF | + 63 x CIM1 | 8.0 | 0.4 | 0.048 | 0.006 | | | | | 10.4 | 0.3 | 11.9 | 1.6 | 18.4 | 0.3 | 87.8 | 1.3 | < 0.0001 | > 99 |
| | + 1.7 x TTP | 8.5 | 0.2 | 0.038 | 0.003 | | | | | 9.91 | 0.18 | 10.9 | 0.8 | 18.43 | 0.18 | 87.8 | 0.9 | < 0.0001 | > 99 |
| 828-838 | apo | | | | | | | | | 11.43 | 0.12 | 14.1 | 1.2 | 11.43 | 0.12 | 127.0 | 1.3 | 0.8886 | 90 |
| WPEANQHFSKE | + 63 x CIM1 | peptide n.d. | | | | | | | | | | | | | | | | | |
| | + 1.7 x TTP | | | | | | | | | 10.75 | 0.08 | 14.5 | 0.9 | 10.75 | 0.08 | 119.4 | 0.9 | - | > 99 |
| 829-838 | apo | | | | | | | | | 11.89 | 0.11 | 16.9 | 1.9 | 11.89 | 0.11 | 132.1 | 1.3 | - | 88 |
| PEANQHFSKE | + 63 x CIM1 | peptide n.d. | | | | | | | | | | | | | | | | | |
| | + 1.7 x TTP | | | | | | | | | 11.14 | 0.13 | 17 | 2 | 11.14 | 0.13 | 123.8 | 1.5 | 0.1087 | 60 |
| 839-846 | apo | 4.8 | 0.3 | 0.049 | 0.007 | 2.0 | 0.2 | 4.9 | 1.5 | | | | | 6.82 | 0.18 | 97 | 3 | < 0.0001 | > 99 |
| IDDEANSY | + 63 x CIM1 | 3.6 | 0.2 | 0.035 | 0.006 | 1.1 | 0.2 | 3.3 | 1.7 | | | | | 4.78 | 0.16 | 68 | 2 | < 0.0001 | > 99 |
| | + 1.7 x TTP | 3.5 | 0.3 | 0.034 | 0.008 | 1.6 | 0.3 | 3.0 | 1.6 | | | | | 5.2 | 0.2 | 74 | 3 | 0.0003 | > 99 |
| 839- 847 | apo | 6.4 | 0.7 | 0.038 | 0.007 | 1.8 | 0.7 | 1.9 | 1.6 | | | | | 8.2 | 0.2 | 103 | 3 | < 0.0001 | > 99 |
| IDDEANSYF | + 63 x CIM1 | 5.5 | 0.3 | 0.052 | 0.009 | | | | | | | | | 5.5 | 0.3 | 68 | 4 | 0.1044 | 59 |
| | + 1.7 x TTP | 4.4 | 0.5 | 0.1[a] | | | | | | 0.4 | 0.3 | 30[a] | | 4.8 | 0.3 | 60 | 4 | - | 89 |
| 846-859 | apo | 6.3 | 0.3 | 0.054 | 0.006 | 4.2 | 0.2 | 5.3 | 0.9 | | | | | 10.6 | 0.2 | 106 | 2 | < 0.0001 | > 99 |
| YFQRIYNHPPHPTM | + 63 x CIM1 | 4.7 | 0.4 | 0.065 | 0.014 | 3.2 | 0.3 | 4.9 | 1.4 | | | | | 8.0 | 0.3 | 80 | 3 | 0.0021 | 98 |
| | + 1.7 x TTP | 4.8 | 0.5 | 0.073 | 0.019 | 3.4 | 0.4 | 9 | 3 | | | | | 8.2 | 0.3 | 82 | 3 | 0.0007 | > 99 |
| 847-859 | apo | 5.16 | 0.20 | 0.058 | 0.006 | 4.07 | 0.15 | 6.3 | 0.7 | | | | | 9.23 | 0.13 | 102.5 | 1.5 | < 0.0001 | > 99 |
| FQRIYNHPPHPTM | + 63 x CIM1 | 3.9 | 0.2 | 0.075 | 0.011 | 2.49 | 0.17 | 6.7 | 1.3 | | | | | 6.39 | 0.14 | 71.0 | 1.6 | < 0.0001 | > 99 |
| | + 1.7 x TTP | 4.3 | 0.2 | 0.065 | 0.009 | 3.45 | 0.17 | 7.1 | 1.1 | | | | | 7.75 | 0.15 | 86.1 | 1.7 | < 0.0001 | > 99 |
| 847-861 | apo | 5.0 | 0.3 | 0.066 | 0.011 | 6.7 | 0.2 | 6.5 | 0.7 | | | | | 11.7 | 0.2 | 106.4 | 2.0 | < 0.0001 | > 99 |
| FQRIYNHPPHPTMSV | + 63 x CIM1 | 4.10 | 0.13 | 0.050 | 0.004 | 5.16 | 0.10 | 6.9 | 0.4 | | | | | 9.26 | 0.09 | 84.2 | 0.8 | < 0.0001 | > 99 |
| | + 1.7 x TTP | 4.3 | 0.4 | 0.056 | 0.012 | 6.0 | 0.3 | 6.5 | 0.9 | | | | | 10.4 | 0.3 | 94 | 2 | < 0.0001 | > 99 |
| 848-859 | apo | 4.5 | 0.3 | 0.059 | 0.008 | 4.4 | 0.2 | 6.9 | 1.0 | | | | | 8.88 | 0.18 | 111 | 2 | < 0.0001 | > 99 |
| QRIYNHPPHPTM | + 63 x CIM1 | 3.5 | 0.2 | 0.085 | 0.014 | 3.17 | 0.16 | 6.0 | 0.9 | | | | | 6.66 | 0.13 | 83.2 | 1.7 | < 0.0001 | > 99 |
| | + 1.7 x TTP | 3.6 | 0.2 | 0.061 | 0.010 | 3.92 | 0.18 | 6.7 | 0.9 | | | | | 7.52 | 0.16 | 94.0 | 1.9 | < 0.0001 | > 99 |
| 860-865 | apo | 1.03 | 0.06 | 0.074 | 0.012 | 1.35 | 0.05 | 8.2 | 1.0 | | | | | 2.38 | 0.04 | 47.5 | 0.8 | < 0.0001 | > 99 |
| SVDEVL | + 63 x CIM1 | 0.92 | 0.05 | 0.052 | 0.006 | 1.04 | 0.04 | 7.0 | 0.7 | | | | | 1.96 | 0.03 | 39.2 | 0.6 | < 0.0001 | > 99 |
| | + 1.7 x TTP | 0.89 | 0.05 | 0.077 | 0.013 | 1.39 | 0.04 | 6.6 | 0.6 | | | | | 2.28 | 0.04 | 45.5 | 0.7 | < 0.0001 | > 99 |
| 862-867 | apo | HDX n.d. | | | | | | | | | | | | | | 0 | | | |
| DEVLEM | + 63 x CIM1 | HDX n.d. | | | | | | | | | | | | | | 0 | | | |
| | + 1.7 x TTP | HDX n.d. | | | | | | | | | | | | | | 0 | | | |
| 863-867 | apo | HDX n.d. | | | | | | | | | | | | | | 0 | | | |
| EVLEM | + 63 x CIM1 | HDX n.d. | | | | | | | | | | | | | | 0 | | | |
| | + 1.7 x TTP | HDX n.d. | | | | | | | | | | | | | | 0 | | | |
| 864-868 | apo | HDX n.d. | | | | | | | | | | | | | | 0 | | | |
| VLEML | + 63 x CIM1 | HDX n.d. | | | | | | | | | | | | | | 0 | | | |
| | + 1.7 x TTP | HDX n.d. | | | | | | | | | | | | | | 0 | | | |
| 866-883 | apo | 2.83 | 0.12 | 0.038 | 0.004 | | | | | 3.95 | 0.09 | 14.6 | 1.9 | 6.78 | 0.09 | 40.0 | 0.5 | < 0.0001 | > 99 |
| EMLQRFKDSTIKREREVF | + 63 x CIM1 | 2.65 | 0.19 | 0.043 | 0.008 | | | | | 3.65 | 0.14 | 15 | 4 | 6.30 | 0.14 | 37.1 | 0.8 | < 0.0001 | > 99 |
| | + 1.7 x TTP | 2.63 | 0.10 | 0.042 | 0.004 | | | | | 3.70 | 0.08 | 14.3 | 1.7 | 6.34 | 0.08 | 37.3 | 0.4 | < 0.0001 | > 99 |
| 868-883 | apo | 2.72 | 0.09 | 0.046 | 0.004 | | | | | 3.83 | 0.05 | 30[a] | | 6.54 | 0.07 | 43.6 | 0.5 | 0.4761 | 92 |
| LQRFKDSTIKREREVF | + 63 x CIM1 | 2.20 | 0.07 | 0.052 | 0.005 | | | | | 3.72 | 0.05 | 30[a] | | 5.91 | 0.06 | 39.4 | 0.4 | 0.8785 | 94 |
| | + 1.7 x TTP | 2.56 | 0.07 | 0.042 | 0.003 | | | | | 3.76 | 0.04 | 30[a] | | 6.33 | 0.06 | 42.2 | 0.4 | 0.7625 | 94 |
| 868-886 | apo | 2.11 | 0.09 | 0.044 | 0.005 | | | | | 3.37 | 0.05 | 30[a] | | 5.48 | 0.07 | 30.4 | 0.4 | 0.3539 | 89 |
| LQRFKDSTIKREREVFNCM | + 63 x CIM1 | 2.08 | 0.11 | 0.041 | 0.006 | | | | | 3.33 | 0.07 | 30[a] | | 5.41 | 0.09 | 30.1 | 0.5 | 0.6794 | 93 |
| | + 1.7 x TTP | 2.18 | 0.12 | 0.040 | 0.006 | | | | | 3.28 | 0.07 | 30[a] | | 5.46 | 0.10 | 30.3 | 0.5 | 0.9208 | 94 |
| 869-883 | apo | 2.42 | 0.18 | 0.054 | 0.011 | | | | | 3.83 | 0.11 | 30[a] | | 6.25 | 0.15 | 44.7 | 1.1 | 0.0295 | 74 |
| QRFKDSTIKREREVF | + 63 x CIM1 | 2.14 | 0.13 | 0.051 | 0.009 | | | | | 3.66 | 0.08 | 30[a] | | 5.80 | 0.11 | 41.5 | 0.8 | 0.2056 | 83 |
| | + 1.7 x TTP | 2.37 | 0.14 | 0.049 | 0.008 | | | | | 3.73 | 0.09 | 30[a] | | 6.10 | 0.12 | 43.6 | 0.9 | 0.757 | 94 |

| Amino acids/sequence | state | Best-fit values | | | | | | | | | | | | | | | | | |
|---|---|---|---|---|---|---|---|---|---|---|---|---|---|---|---|---|---|---|---|
| | | $ND_1$ [Da] | $\delta ND_1$ | $k_1$ [min$^{-1}$] | $\delta k_1$ | $ND_2$ [Da] | $\delta ND_2$ | $k_2$ [min$^{-1}$] | $\delta k_2$ | $ND_3$ [Da] | $\delta ND_3$ | $k_3$ [min$^{-1}$] | $\delta k_3$ | ND [Da] | δND | $D_{\%}$ | $\delta D_{\%}$ | P-value | % AIC |
| 886-893 | apo | 0.33 | 0.11 | 0.1[a] | | | | | | 0.01 | 0.07 | 30[a] | | 0.35 | 0.07 | 4.9 | 1.0 | 0.9867 | 90 |
| MLRNLFEE | + 63 x CIM1 | HDX n.d. | | | | | | | | | | | | | | 0 | | | |
| | + 1.7 x TTP | HDX n.d. | | | | | | | | | | | | | | 0 | | | |
| 887-893 | apo | 1.1 | 0.7 | 0.006 | 0.006 | | | | | | | | | 1.1 | 0.7 | 19 | 11 | - | > 99 |
| LRNLFEE | + 63 x CIM1 | HDX n.d. | | | | | | | | | | | | | | 0 | | | |
| | + 1.7 x TTP | HDX n.d. | | | | | | | | | | | | | | 0 | | | |
| 887-894 | apo | 4 | 18 | 0.002 | 0.011 | | | | | | | | | 4 | 18 | 60 | 300 | - | > 99 |
| LRNLFEEY | + 63 x CIM1 | HDX n.d. | | | | | | | | | | | | | | 0 | | | |
| | + 1.7 x TTP | HDX n.d. | | | | | | | | | | | | | | 0 | | | |
| 888-893 | apo | 0.07 | 0.07 | 0.1[a] | | | | | | 0.14 | 0.05 | 30[a] | | 0.21 | 0.05 | 4.2 | 0.9 | - | 64 |
| RNLFEE | + 63 x CIM1 | HDX n.d. | | | | | | | | | | | | | | 0 | | | |
| | + 1.7 x TTP | HDX n.d. | | | | | | | | | | | | | | 0 | | | |
| 894-904 | apo | 4.6 | 0.2 | 0.030 | 0.005 | 2.69 | 0.18 | 6.7 | 1.4 | | | | | 7.3 | 0.2 | 91 | 3 | < 0.0001 | > 99 |
| YRFFPQYPDKE | + 63 x CIM1 | 3.4 | 0.2 | 0.049 | 0.007 | | | | | 1.60 | 0.15 | 12 | 5 | 5.00 | 0.14 | 62.5 | 1.8 | < 0.0001 | > 99 |
| | + 1.7 x TTP | 3.4 | 0.2 | 0.035 | 0.006 | 1.98 | 0.17 | 8 | 2 | | | | | 5.38 | 0.18 | 67 | 2 | < 0.0001 | > 99 |
| 894-905 | apo | 5.48 | 0.15 | 0.044 | 0.003 | 2.48 | 0.12 | 5.4 | 0.7 | | | | | 7.95 | 0.11 | 88.4 | 1.2 | < 0.0001 | > 99 |
| YRFFPQYPDKEL | + 63 x CIM1 | 5.27 | 0.15 | 0.032 | 0.003 | | | | | 1.27 | 0.11 | 10 | 3 | 6.54 | 0.12 | 72.6 | 1.3 | < 0.0001 | > 99 |
| | + 1.7 x TTP | 4.41 | 0.18 | 0.034 | 0.004 | 1.81 | 0.14 | 5.9 | 1.3 | | | | | 6.22 | 0.14 | 69.1 | 1.6 | < 0.0001 | > 99 |
| 894-909 | apo | 8.16 | 0.10 | 0.0455 | 0.0014 | 2.54 | 0.08 | 4.2 | 0.4 | | | | | 10.70 | 0.07 | 82.3 | 0.5 | < 0.0001 | > 99 |
| YRFFPQYPDKELHITA | + 63 x CIM1 | 7.22 | 0.18 | 0.029 | 0.002 | 1.36 | 0.14 | 4.4 | 1.3 | | | | | 8.58 | 0.15 | 66.0 | 1.1 | < 0.0001 | > 99 |
| | + 1.7 x TTP | 6.99 | 0.16 | 0.031 | 0.002 | 1.42 | 0.12 | 5.4 | 1.4 | | | | | 8.41 | 0.13 | 64.7 | 1.0 | < 0.0001 | > 99 |
| 901-909 | apo | 3.8 | 0.4 | 0.046 | 0.011 | 2.3 | 0.3 | 5 | 2 | | | | | 6.1 | 0.3 | 76 | 3 | 0.002 | 98 |
| PDKELHITA | + 63 x CIM1 | 4.6 | 0.2 | 0.026 | 0.004 | 1.13 | 0.14 | 6 | 2 | | | | | 5.69 | 0.19 | 71 | 2 | < 0.0001 | > 99 |
| | + 1.7 x TTP | 3.9 | 0.2 | 0.040 | 0.005 | 1.36 | 0.15 | 5.8 | 1.9 | | | | | 5.23 | 0.14 | 65.4 | 1.8 | < 0.0001 | > 99 |
| 912-917 | apo | HDX n.d. | | | | | | | | | | | | | | 0 | | | |
| FGGIIE | + 63 x CIM1 | HDX n.d. | | | | | | | | | | | | | | 0 | | | |
| | + 1.7 x TTP | HDX n.d. | | | | | | | | | | | | | | 0 | | | |
| 912-919 | apo | 1.77 | 0.08 | 0.022 | 0.003 | | | | | | | | | 1.77 | 0.08 | 25.3 | 1.1 | 0.4239 | 98 |
| FGGIIEKG | + 63 x CIM1 | peptide n.d. | | | | | | | | | | | | | | | | | |
| | + 1.7 x TTP | 1.55 | 0.19 | 0.021 | 0.007 | | | | | | | | | 1.55 | 0.19 | 22 | 3 | 0.2522 | 95 |
| 912-922 | apo | 4.28 | 0.08 | 0.0270 | 0.0017 | | | | | 1.29 | 0.06 | 12 | 2 | 5.57 | 0.07 | 55.7 | 0.7 | < 0.0001 | > 99 |
| FGGIIEKGLVT | + 63 x CIM1 | 4.14 | 0.11 | 0.024 | 0.002 | | | | | 1.20 | 0.07 | 13 | 4 | 5.34 | 0.11 | 53.4 | 1.1 | < 0.0001 | > 99 |
| | + 1.7 x TTP | 4.12 | 0.05 | 0.0283 | 0.0010 | | | | | 1.35 | 0.03 | 13.3 | 1.8 | 5.48 | 0.04 | 54.8 | 0.4 | < 0.0001 | > 99 |
| 912-924 | apo | 4.07 | 0.08 | 0.0242 | 0.0016 | | | | | 3.71 | 0.05 | 20 | 3 | 7.77 | 0.08 | 64.8 | 0.6 | < 0.0001 | > 99 |
| FGGIIEKGLVTYM | + 63 x CIM1 | 3.80 | 0.13 | 0.019 | 0.002 | | | | | 3.81 | 0.06 | 17 | 2 | 7.61 | 0.14 | 63.4 | 1.2 | < 0.0001 | > 99 |
| | + 1.7 x TTP | 4.14 | 0.12 | 0.026 | 0.003 | | | | | 3.78 | 0.08 | 19 | 4 | 7.92 | 0.12 | 66.0 | 1.0 | < 0.0001 | > 99 |
| 915-923 | apo | 3.49 | 0.10 | 0.027 | 0.003 | | | | | 2.76 | 0.07 | 13.7 | 2.0 | 6.25 | 0.09 | 78.1 | 1.2 | < 0.0001 | > 99 |
| IIEKGLVTY | + 63 x CIM1 | peptide n.d. | | | | | | | | | | | | | | | | | |
| | + 1.7 x TTP | 3.73 | 0.11 | 0.026 | 0.002 | | | | | 2.67 | 0.05 | 30[a] | | 6.40 | 0.10 | 80.0 | 1.3 | 0.4981 | 92 |
| 918-923 | apo | 2.25 | 0.13 | 0.031 | 0.005 | | | | | 2.87 | 0.09 | 12.1 | 1.8 | 5.13 | 0.11 | 103 | 2 | < 0.0001 | > 99 |
| KGLVTY | + 63 x CIM1 | 2.4 | 0.4 | 0.014 | 0.007 | 3.04 | 0.11 | 9.5 | 1.3 | | | | | 5.5 | 0.5 | 109 | 10 | < 0.0001 | > 99 |
| | + 1.7 x TTP | 2.30 | 0.08 | 0.040 | 0.003 | | | | | 2.61 | 0.06 | 15 | 2 | 4.90 | 0.06 | 98.1 | 1.1 | < 0.0001 | > 99 |
| 923-927 | apo | 1.90 | 0.10 | 0.070 | 0.010 | 1.78 | 0.08 | 8.9 | 1.3 | | | | | 3.68 | 0.07 | 92.0 | 1.7 | < 0.0001 | > 99 |
| YMALG | + 63 x CIM1 | 1.9 | 0.2 | 0.056 | 0.014 | 1.84 | 0.18 | 6.7 | 1.7 | | | | | 3.77 | 0.13 | 94 | 3 | 0.0005 | > 99 |
| | + 1.7 x TTP | 1.96 | 0.07 | 0.056 | 0.005 | 1.88 | 0.05 | 8.5 | 0.8 | | | | | 3.84 | 0.05 | 96.0 | 1.1 | < 0.0001 | > 99 |
| 923-928 | apo | 3.46 | 0.09 | 0.035 | 0.003 | 1.99 | 0.07 | 7.3 | 0.8 | | | | | 5.45 | 0.07 | 109.0 | 1.4 | < 0.0001 | > 99 |
| YMALGL | + 63 x CIM1 | 3.00 | 0.14 | 0.047 | 0.005 | 1.70 | 0.10 | 7.6 | 1.5 | | | | | 4.70 | 0.09 | 94.0 | 1.9 | < 0.0001 | > 99 |
| | + 1.7 x TTP | 2.9 | 0.2 | 0.047 | 0.009 | 2.04 | 0.18 | 6.8 | 1.8 | | | | | 4.91 | 0.17 | 98 | 3 | 0.0001 | > 99 |
| 923-930 | apo | 2.7 | 0.2 | 0.039 | 0.009 | 1.67 | 0.18 | 6.2 | 1.7 | | | | | 4.35 | 0.17 | 62 | 2 | 0.0009 | 99 |
| YMALGLAL | + 63 x CIM1 | 2.99 | 0.12 | 0.034 | 0.004 | 1.47 | 0.08 | 8.4 | 1.8 | | | | | 4.46 | 0.10 | 63.7 | 1.5 | < 0.0001 | > 99 |
| | + 1.7 x TTP | 2.88 | 0.10 | 0.038 | 0.003 | 1.78 | 0.09 | 7.2 | 1.0 | | | | | 4.67 | 0.07 | 66.7 | 1.0 | < 0.0001 | > 99 |
| 924-930 | apo | 2.16 | 0.13 | 0.02 | 0.02 | 1.6 | 0.7 | 0.7[b] | 0.4 | | | | | 3.74 | 0.796 | 62 | 13 | < 0.0001 | > 99 |
| MALGLAL | + 63 x CIM1 | 2.4 | 0.5 | 0.02 | 0.02 | 0.9 | 0.9 | 0.7[b] | 0.8 | | | | | 3.3 | 0.3 | 55 | 5 | < 0.0001 | > 99 |
| | + 1.7 x TTP | 2.28 | 0.16 | 0.021 | 0.010 | 1.4 | 0.4 | 1.0 | 0.4 | | | | | 3.7 | 0.2 | 61 | 3 | < 0.0001 | > 99 |
| 928-934 | apo | HDX n.d. | | | | | | | | | | | | | | 0 | | | |
| LALRYVL | + 63 x CIM1 | HDX n.d. | | | | | | | | | | | | | | 0 | | | |
| | + 1.7 x TTP | HDX n.d. | | | | | | | | | | | | | | 0 | | | |
| 929-934 | apo | HDX n.d. | | | | | | | | | | | | | | 0 | | | |
| ALRYVL | + 63 x CIM1 | HDX n.d. | | | | | | | | | | | | | | 0 | | | |
| | + 1.7 x TTP | HDX n.d. | | | | | | | | | | | | | | 0 | | | |
| 929-936 | apo | HDX n.d. | | | | | | | | | | | | | | 0 | | | |
| ALRYVLEA | + 63 x CIM1 | HDX n.d. | | | | | | | | | | | | | | 0 | | | |
| | + 1.7 x TTP | HDX n.d. | | | | | | | | | | | | | | 0 | | | |
| 931-936 | apo | HDX n.d. | | | | | | | | | | | | | | 0 | | | |
| RYVLEA | + 63 x CIM1 | peptide n.d. | | | | | | | | | | | | | | | | | |
| | + 1.7 x TTP | HDX n.d. | | | | | | | | | | | | | | 0 | | | |
| 935-945 | apo | 0.57 | 0.09 | 0.06 | 0.02 | 5.45 | 0.07 | 9.4 | 0.4 | | | | | 6.02 | 0.06 | 66.8 | 0.7 | 0.0004 | > 99 |
| EALRKPFGSKM | + 63 x CIM1 | 0.69 | 0.20 | 0.1[a] | | 5.15 | 0.14 | 9.2 | 0.9 | | | | | 5.84 | 0.12 | 64.8 | 1.3 | 0.0080 | 92 |
| | + 1.7 x TTP | 0.44 | 0.17 | 0.1[a] | | 5.19 | 0.13 | 9.1 | 0.8 | | | | | 5.64 | 0.09 | 62.6 | 1.0 | 0.0283 | 75 |
| 936-945 | apo | | | | | 6.59 | 0.07 | 9.4 | 0.6 | | | | | 6.59 | 0.07 | 82.4 | 0.9 | 0.8563 | 91 |
| ALRKPFGSKM | + 63 x CIM1 | | | | | 6.23 | 0.07 | 8.4 | 0.5 | | | | | 6.23 | 0.07 | 77.8 | 0.9 | 0.1038 | 89 |
| | + 1.7 x TTP | 0.8 | 0.2 | 0.1[a] | | 6.18 | 0.18 | 9.2 | 1.0 | | | | | 7.01 | 0.13 | 87.6 | 1.6 | 0.0067 | 95 |
| 937-944 | apo | | | | | 5.95 | 0.13 | 8.9 | 1.1 | | | | | 5.95 | 0.13 | 99.1 | 2 | 0.0775 | 78 |
| LRKPFGSK | + 63 x CIM1 | | | | | 5.54 | 0.12 | 9.1 | 1.3 | | | | | 5.54 | 0.12 | 92.4 | 1.9 | 0.082 | 56 |
| | + 1.7 x TTP | | | | | 5.78 | 0.11 | 8.3 | 0.9 | | | | | 5.78 | 0.11 | 96.4 | 1.9 | 0.0873 | 81 |
| 937-945 | apo | 0.80 | 0.09 | 0.028 | 0.010 | 6.53 | 0.06 | 9.8 | 0.4 | | | | | 7.33 | 0.08 | 104.7 | 1.1 | < 0.0001 | > 99 |
| LRKPFGSKM | + 63 x CIM1 | 0.63 | 0.13 | 0.06 | 0.03 | 6.10 | 0.10 | 9.2 | 0.6 | | | | | 6.72 | 0.09 | 96.0 | 1.3 | 0.0038 | 94 |
| | + 1.7 x TTP | 0.74 | 0.05 | 0.033 | 0.006 | 6.35 | 0.04 | 8.89 | 0.18 | | | | | 7.09 | 0.04 | 101.3 | 0.6 | < 0.0001 | > 99 |

| Amino acids/sequence | state | Best-fit values | | | | | | | | | | | | | | | | | |
|---|---|---|---|---|---|---|---|---|---|---|---|---|---|---|---|---|---|---|---|
| | | $ND_1$ [Da] | $\delta ND_1$ | $k_1$ [min$^{-1}$] | $\delta k_1$ | $ND_2$ [Da] | $\delta ND_2$ | $k_2$ [min$^{-1}$] | $\delta k_2$ | $ND_3$ [Da] | $\delta ND_3$ | $k_3$ [min$^{-1}$] | $\delta k_3$ | ND [Da] | δND | $D_{\%}$ | $\delta D_{\%}$ | P-value | % AIC |
| 937-947 | apo | | | | | 6.26 | 0.06 | 9.0 | 0.5 | | | | | 6.26 | 0.06 | 69.6 | 0.7 | 0.0537 | 67 |
| LRKPFGSKMYY | + 63 x CIM1 | peptide n.d. | | | | | | | | | | | | | | | | | |
| | + 1.7 x TTP | 0.43 | 0.11 | 0.03 | 0.02 | 5.72 | 0.07 | 9.0 | 0.4 | | | | | 6.16 | 0.10 | 68.4 | 1.1 | 0.0075 | 89 |
| 946-951 | apo | HDX n.d. | | | | | | | | | | | | | | 0 | | | |
| YYFGIA | + 63 x CIM1 | HDX n.d. | | | | | | | | | | | | | | 0 | | | |
| | + 1.7 x TTP | HDX n.d. | | | | | | | | | | | | | | 0 | | | |
| 946-952 | apo | HDX n.d. | | | | | | | | | | | | | | 0 | | | |
| YYFGIAA | + 63 x CIM1 | HDX n.d. | | | | | | | | | | | | | | 0 | | | |
| | + 1.7 x TTP | HDX n.d. | | | | | | | | | | | | | | 0 | | | |
| 946-953 | apo | HDX n.d. | | | | | | | | | | | | | | 0 | | | |
| YYFGIAAL | + 63 x CIM1 | HDX n.d. | | | | | | | | | | | | | | 0 | | | |
| | + 1.7 x TTP | HDX n.d. | | | | | | | | | | | | | | 0 | | | |
| 948-952 | apo | HDX n.d. | | | | | | | | | | | | | | 0 | | | |
| FGIAA | + 63 x CIM1 | HDX n.d. | | | | | | | | | | | | | | 0 | | | |
| | + 1.7 x TTP | HDX n.d. | | | | | | | | | | | | | | 0 | | | |
| 948-953 | apo | HDX n.d. | | | | | | | | | | | | | | 0 | | | |
| FGIAAL | + 63 x CIM1 | HDX n.d. | | | | | | | | | | | | | | 0 | | | |
| | + 1.7 x TTP | HDX n.d. | | | | | | | | | | | | | | 0 | | | |
| 952-966 | apo | 3.25 | 0.17 | 0.0110 | 0.0013 | | | | | 3.53 | 0.02 | 15.5 | 0.7 | 6.77 | 0.18 | 52.1 | 1.4 | < 0.0001 | > 99 |
| ALDRFKNRLKDYPQY | + 63 x CIM1 | 3.1 | 0.6 | 0.011 | 0.005 | | | | | 3.48 | 0.08 | 17 | 3 | 6.6 | 0.6 | 51 | 5 | < 0.0001 | > 99 |
| | + 1.7 x TTP | 3.3 | 0.3 | 0.011 | 0.002 | | | | | 3.54 | 0.04 | 14.2 | 1.0 | 6.9 | 0.3 | 52.8 | 2.5 | < 0.0001 | > 99 |
| 952-972 | apo | 5.92 | 0.06 | 0.0300 | 0.0008 | | | | | 3.47 | 0.04 | 15.2 | 1.1 | 9.38 | 0.05 | 49.4 | 0.2 | < 0.0001 | > 99 |
| ALDRFKNRLKDYPQYCQHLAS | + 63 x CIM1 | peptide n.d. | | | | | | | | | | | | | | | | | |
| | + 1.7 x TTP | 5.70 | 0.08 | 0.0327 | 0.0013 | | | | | 3.45 | 0.06 | 14.2 | 1.4 | 9.15 | 0.06 | 48.2 | 0.3 | < 0.0001 | > 99 |
| 953-966 | apo | 3.0 | 0.2 | 0.016 | 0.003 | | | | | 3.41 | 0.05 | 30[a] | | 6.4 | 0.2 | 53.1 | 1.9 | 0.165 | 79 |
| LDRFKNRLKDYPQY | + 63 x CIM1 | 2.9 | 0.5 | 0.010 | 0.003 | | | | | 3.56 | 0.06 | 14.3 | 1.3 | 6.5 | 0.5 | 54 | 4 | < 0.0001 | > 99 |
| | + 1.7 x TTP | 3.1 | 0.4 | 0.012 | 0.003 | | | | | 3.52 | 0.06 | 15.5 | 1.8 | 6.6 | 0.4 | 55 | 3 | < 0.0001 | > 99 |
| 953-972 | apo | 6.06 | 0.06 | 0.0330 | 0.0010 | | | | | 3.03 | 0.03 | 30[a] | | 9.08 | 0.05 | 50.5 | 0.3 | 0.5934 | 93 |
| LDRFKNRLKDYPQYCQHLAS | + 63 x CIM1 | peptide n.d. | | | | | | | | | | | | | | | | | |
| | + 1.7 x TTP | 5.81 | 0.12 | 0.0317 | 0.0019 | | | | | 3.11 | 0.09 | 14 | 2 | 8.92 | 0.09 | 49.5 | 0.5 | < 0.0001 | > 99 |
| 954-966 | apo | 2.94 | 0.19 | 0.0117 | 0.0018 | | | | | 3.37 | 0.03 | 15.9 | 1.0 | 6.3 | 0.2 | 57.3 | 1.9 | < 0.0001 | > 99 |
| DRFKNRLKDYPQY | + 63 x CIM1 | 4.3 | 1.7 | 0.006 | 0.003 | | | | | 3.25 | 0.05 | 12.8 | 1.0 | 7.6 | 1.7 | 69 | 16 | < 0.0001 | > 99 |
| | + 1.7 x TTP | 3.0 | 0.3 | 0.011 | 0.003 | | | | | 3.32 | 0.04 | 15.3 | 1.3 | 6.3 | 0.3 | 58 | 3 | < 0.0001 | > 99 |
| 954-972 | apo | 5.68 | 0.08 | 0.0314 | 0.0012 | | | | | 3.07 | 0.06 | 16 | 2 | 8.75 | 0.06 | 51.5 | 0.4 | < 0.0001 | > 99 |
| DRFKNRLKDYPQYCQHLAS | + 63 x CIM1 | 5.54 | 0.11 | 0.0283 | 0.0018 | | | | | 3.14 | 0.08 | 16 | 3 | 8.68 | 0.10 | 51.0 | 0.6 | < 0.0001 | > 99 |
| | + 1.7 x TTP | 5.36 | 0.09 | 0.0327 | 0.0015 | | | | | 3.13 | 0.06 | 13.8 | 1.6 | 8.49 | 0.07 | 50.0 | 0.4 | < 0.0001 | > 99 |
| 967-972 | apo | 2.14 | 0.03 | 0.094 | 0.005 | | | | | | | | | 2.14 | 0.03 | 42.7 | 0.5 | 0.0773 | 78 |
| CQHLAS | + 63 x CIM1 | 2.20 | 0.04 | 0.095 | 0.007 | | | | | | | | | 2.20 | 0.04 | 44.0 | 0.7 | 0.4001 | 97 |
| | + 1.7 x TTP | 2.00 | 0.09 | 0.079 | 0.013 | | | | | | | | | 2.00 | 0.09 | 40.0 | 1.7 | - | 90 |
| 973-985 | apo | 7.8 | 0.3 | 0.061 | 0.005 | 3.8 | 0.2 | 3.7 | 0.6 | | | | | 11.51 | 0.17 | 104.6 | 1.5 | < 0.0001 | > 99 |
| ISHFMQFPHHLQE | + 63 x CIM1 | 6.3 | 0.8 | 0.052 | 0.009 | 3.6 | 0.8 | 1.9 | 0.9 | | | | | 9.8 | 0.2 | 90 | 2 | 0.0002 | > 99 |
| | + 1.7 x TTP | 8.2 | 0.3 | 0.058 | 0.005 | 3.1 | 0.2 | 4.7 | 1.0 | | | | | 11.31 | 0.19 | 102.8 | 1.7 | < 0.0001 | > 99 |
| 974-985 | apo | 6.9 | 0.4 | 0.042 | 0.005 | 3.9 | 0.3 | 2.9 | 0.7 | | | | | 10.8 | 0.2 | 108 | 2 | < 0.0001 | > 99 |
| SHFMQFPHHLQE | + 63 x CIM1 | 7.2 | 0.5 | 0.055 | 0.009 | 2.2 | 0.4 | 9 | 5 | | | | | 9.4 | 0.3 | 94 | 3 | 0.0003 | > 99 |
| | + 1.7 x TTP | 6.8 | 0.4 | 0.064 | 0.008 | 2.9 | 0.3 | 5.2 | 1.4 | | | | | 9.8 | 0.2 | 98 | 2 | 0.0001 | > 99 |
| 977-986 | apo | 4.3 | 0.2 | 0.1[a] | | | | | | 0.41 | 0.16 | 30[a] | | 4.70 | 0.16 | 58.7 | 2.0 | 0.1005 | 58 |
| MQFPHHLQEY | + 63 x CIM1 | 4.0 | 0.2 | 0.055 | 0.008 | | | | | 0.38 | 0.13 | 30[a] | | 4.38 | 0.17 | 55 | 2 | 0.0127 | 89 |
| | + 1.7 x TTP | 3.8 | 0.6 | 0.049 | 0.009 | 1.1 | 0.6 | 1.6 | 1.7 | | | | | 4.89 | 0.12 | 61.2 | 1.5 | < 0.0001 | > 99 |
| 978-985 | apo | 3.4 | 0.4 | 0.062 | 0.009 | 1.5 | 0.4 | 1.7 | 1.0 | | | | | 4.91 | 0.09 | 81.8 | 1.5 | < 0.0001 | > 99 |
| QFPHHLQE | + 63 x CIM1 | peptide n.d. | | | | | | | | | | | | | | | | | |
| | + 1.7 x TTP | 3 | 2 | 0.05 | 0.04 | 2 | 2 | 0.8[b] | 1.1 | | | | | 4.60 | 0.09 | 76.7 | 1.6 | < 0.0001 | > 99 |
| 986-992 | apo | 3.76 | 0.09 | 0.072 | 0.004 | 0.56 | 0.08 | 3.2 | 1.2 | | | | | 4.32 | 0.05 | 72.1 | 0.8 | < 0.0001 | > 99 |
| YIEYGQQ | + 63 x CIM1 | 4.31 | 0.10 | 0.065 | 0.005 | | | | | | | | | 4.31 | 0.10 | 71.9 | 1.6 | 0.6323 | 88 |
| | + 1.7 x TTP | 4.21 | 0.10 | 0.079 | 0.006 | | | | | 0.23 | 0.06 | 30[a] | | 4.44 | 0.08 | 74.1 | 1.4 | 0.0946 | 66 |
| 986-999 | apo | 4.52 | 0.05 | 0.0669 | 0.0020 | 6.47 | 0.04 | 4.91 | 0.09 | | | | | 10.99 | 0.04 | 99.9 | 0.3 | < 0.0001 | > 99 |
| YIEYGQQSRDPPVK | + 63 x CIM1 | 4.41 | 0.12 | 0.064 | 0.004 | 5.47 | 0.10 | 4.9 | 0.2 | | | | | 9.89 | 0.08 | 89.9 | 0.7 | < 0.0001 | > 99 |
| | + 1.7 x TTP | 4.57 | 0.10 | 0.064 | 0.004 | 6.21 | 0.08 | 4.77 | 0.18 | | | | | 10.77 | 0.07 | 97.9 | 0.6 | < 0.0001 | > 99 |
| 987-999 | apo | 4.27 | 0.06 | 0.068 | 0.003 | 6.61 | 0.05 | 4.73 | 0.10 | | | | | 10.88 | 0.04 | 108.8 | 0.4 | < 0.0001 | > 99 |
| IEYGQQSRDPPVK | + 63 x CIM1 | 4.16 | 0.12 | 0.067 | 0.005 | 5.93 | 0.09 | 4.8 | 0.2 | | | | | 10.09 | 0.08 | 100.9 | 0.8 | < 0.0001 | > 99 |
| | + 1.7 x TTP | 4.40 | 0.14 | 0.070 | 0.006 | 6.27 | 0.11 | 5.1 | 0.2 | | | | | 10.67 | 0.09 | 106.7 | 0.9 | < 0.0001 | > 99 |
| 988-999 | apo | 3.64 | 0.11 | 0.073 | 0.006 | 5.47 | 0.09 | 5.1 | 0.2 | | | | | 9.10 | 0.07 | 101.2 | 0.8 | < 0.0001 | > 99 |
| EYGQQSRDPPVK | + 63 x CIM1 | 3.57 | 0.09 | 0.060 | 0.003 | 4.80 | 0.07 | 4.62 | 0.19 | | | | | 8.37 | 0.06 | 93.0 | 0.6 | < 0.0001 | > 99 |
| | + 1.7 x TTP | 3.67 | 0.05 | 0.063 | 0.002 | 5.29 | 0.04 | 4.90 | 0.11 | | | | | 8.95 | 0.04 | 99.5 | 0.4 | < 0.0001 | > 99 |
| 989-999 | apo | 2.27 | 0.09 | 0.13 | 0.03 | 6.92 | 0.09 | 5.26 | 0.16 | | | | | 9.19 | 0.06 | 114.9 | 0.7 | < 0.0001 | > 99 |
| YGQQSRDPPVK | + 63 x CIM1 | 2.12 | 0.09 | 0.103 | 0.016 | 6.56 | 0.08 | 4.85 | 0.16 | | | | | 8.68 | 0.06 | 108.5 | 0.8 | < 0.0001 | > 99 |
| | + 1.7 x TTP | 2.50 | 0.10 | 0.117 | 0.020 | 6.81 | 0.10 | 5.09 | 0.18 | | | | | 9.31 | 0.07 | 116.4 | 0.9 | < 0.0001 | > 99 |
| 993-999 | apo | | | | | 4.06 | 0.04 | 6.6 | 0.3 | | | | | 4.06 | 0.04 | 101.5 | 1.0 | - | > 99 |
| SRDPPVK | + 63 x CIM1 | 0.48 | 0.13 | 0.1[a] | | 3.67 | 0.10 | 7.1 | 0.6 | | | | | 4.14 | 0.07 | 103.6 | 1.7 | 0.9249 | 94 |
| | + 1.7 x TTP | | | | | 4.07 | 0.04 | 5.9 | 0.3 | | | | | 4.07 | 0.04 | 101.8 | 0.9 | - | > 99 |

[a] Fixed values, according to (Gemmecker et al., 1993; Hoofnagle et al., 2001; Mandell et al., 2001; Rutkowska-Wlodarczyk et al., 2008).

[b] exceptions to the classification that $k_1 \leq 0.1$, $0.1 < k_2 < 10$ and $10 < k_3 \leq 30$, due to lack of space in appropriate column.

### Supplementary Table T5 to Fig 2f.

Kinetic parameters of hydrogen-deuterium exchange for chosen peptides of CNOT1(800-999) E893A/Y900A at 2 µM in the presence of the TTP peptide at 3.3 µM or the CIM1 peptide at 126 µM.

| | | Best-fit values | | | | | | | | | | | | | | | | | |
|---|---|---|---|---|---|---|---|---|---|---|---|---|---|---|---|---|---|---|---|
| **Amino acids/sequence** | **state** | **$ND_1$** | **$\delta ND_1$** | **$k_1$** | **$\delta k_1$** | **$ND_2$** | **$\delta ND_2$** | **$k_2$** | **$\delta k_2$** | **$ND_3$** | **$\delta ND_3$** | **$k_3$** | **$\delta k_3$** | **ND** | **δND** | **$D_{\%}$** | **$\delta D_{\%}$** | **P-value** | **% AIC** |
| | | [Da] | | [$min^{-1}$] | | [Da] | | [$min^{-1}$] | | [Da] | | [$min^{-1}$] | | [Da] | | | | | |
| 800-819 | CNOT1 EY[b] | | | | | | | | | 17.51 | 0.06 | 30[a] | | | | | | - | 86 |
| NNDPFVQRKLGTSGLN | + 63 x CIM1 | | | | | | | | | 16.91 | 0.06 | 30[a] | | | | | | - | 86 |
| QPTF | + 1.7 x TTP | | | | | | | | | 17.38 | 0.05 | 30[a] | | | | | | > 0.9999 | 86 |
| 825-847 | CNOT1 EY[b] | 9.1 | 0.4 | 0.1[a] | | | | | | 10.9 | 0.2 | 30[a] | | 20.0 | 0.2 | 95.0 | 1.2 | 0.3040 | 83 |
| SQVWPEANQHFSKEID | + 63 x CIM1 | 9.7 | 0.4 | 0.064 | 0.008 | | | | | 10.8 | 0.3 | 30[a] | | 20.5 | 0.3 | 97.8 | 1.6 | < 0.0001 | > 99 |
| DEANSYF | + 1.7 x TTP | 9.7 | 0.4 | 0.070 | 0.010 | | | | | 10.8 | 0.3 | 30[a] | | 20.4 | 0.4 | 97.3 | 1.7 | < 0.0001 | > 99 |
| 839-846 | CNOT1 EY[b] | 5.84 | 0.08 | 0.089 | 0.004 | 1.93 | 0.07 | 5.0 | 0.5 | | | | | 7.76 | 0.05 | 110.9 | 0.8 | < 0.0001 | > 99 |
| IDDEANSY | + 63 x CIM1 | 6.1 | 0.3 | 0.070 | 0.007 | 1.5 | 0.2 | 3.3 | 1.4 | | | | | 7.63 | 0.15 | 109 | 2 | 0.0003 | > 99 |
| | + 1.7 x TTP | 5.6 | 0.3 | 0.085 | 0.009 | 2.1 | 0.3 | 2.9 | 0.9 | | | | | 7.70 | 0.13 | 110.0 | 1.9 | 0.0009 | 99 |
| 839-847 | CNOT1 EY[b] | 5.90 | 0.12 | 0.069 | 0.003 | 1.99 | 0.09 | 5.1 | 0.7 | | | | | 7.89 | 0.08 | 98.6 | 1.0 | < 0.0001 | > 99 |
| IDDEANSYF | + 63 x CIM1 | 5.78 | 0.16 | 0.055 | 0.004 | 1.56 | 0.12 | 5.9 | 1.3 | | | | | 7.34 | 0.11 | 91.7 | 1.3 | < 0.0001 | > 99 |
| | + 1.7 x TTP | 5.8 | 0.2 | 0.070 | 0.007 | 2.0 | 0.2 | 3.8 | 1.1 | | | | | 7.77 | 0.15 | 97.2 | 1.8 | 0.0002 | > 99 |
| 847-859 | CNOT1 EY[b] | 4.3 | 0.2 | 0.1[a] | | 4.35 | 0.18 | 7.5 | 1.0 | | | | | 8.63 | 0.12 | 95.9 | 1.4 | - | > 99 |
| FQRIYNHPPHPTM | + 63 x CIM1 | 4.1 | 0.3 | 0.1[a] | | 3.2 | 0.2 | 7.4 | 1.8 | | | | | 7.26 | 0.16 | 80.6 | 1.8 | 0.0183 | 83 |
| | + 1.7 x TTP | 4.5 | 0.3 | 0.1[a] | | 4.01 | 0.20 | 7.8 | 1.2 | | | | | 8.54 | 0.14 | 94.9 | 1.5 | 0.0027 | 98 |
| 847-861 | CNOT1 EY[b] | 4.1 | 0.2 | 0.059 | 0.008 | 6.68 | 0.17 | 7.1 | 0.6 | | | | | 10.78 | 0.15 | 98.0 | 1.4 | 0.0003 | > 99 |
| FQRIYNHPPHPTMSV | + 63 x CIM1 | 4.17 | 0.17 | 0.061 | 0.006 | 5.66 | 0.13 | 7.1 | 0.5 | | | | | 9.83 | 0.11 | 89.4 | 1.0 | < 0.0001 | > 99 |
| | + 1.7 x TTP | 4.3 | 0.2 | 0.068 | 0.010 | 6.32 | 0.19 | 7.2 | 0.7 | | | | | 10.60 | 0.17 | 96.4 | 1.5 | 0.0016 | 99 |
| 848-859 | CNOT1 EY[b] | 4.1 | 0.3 | 0.1[a] | | | | | | 3.8 | 0.2 | 30[a] | | 7.8 | 0.2 | 98 | 3 | 0.2341 | 80 |
| QRIYNHPPHPTM | + 63 x CIM1 | 3.9 | 0.2 | 0.1[a] | | | | | | 2.82 | 0.16 | 30[a] | | 6.67 | 0.16 | 83 | 2 | - | 98 |
| | + 1.7 x TTP | 4.1 | 0.3 | 0.1[a] | | | | | | 3.65 | 0.19 | 30[a] | | 7.73 | 0.20 | 97 | 2 | 0.2287 | 79 |
| 894-904 | CNOT1 EY[b] | | | | | 4.78 | 0.07 | 1.34 | 0.04 | 3.11 | 0.06 | 30[a] | | 7.88 | 0.03 | 98.6 | 0.3 | < 0.0001 | > 99 |
| YRFFPQAPDKE | + 63 x CIM1 | | | | | 5.2 | 0.3 | 1.05 | 0.15 | 2.8 | 0.3 | 30[a] | | 7.97 | 0.13 | 99.6 | 1.7 | < 0.0001 | > 99 |
| | + 1.7 x TTP | | | | | 5.04 | 0.16 | 1.24 | 0.09 | 3.00 | 0.15 | 30[a] | | 8.04 | 0.06 | 100.5 | 0.8 | < 0.0001 | > 99 |
| 894-905 | CNOT1 EY[b] | | | | | 5.8 | 0.2 | 1.48 | 0.11 | 3.17 | 0.20 | 30[a] | | 9.02 | 0.08 | 100.2 | 0.9 | < 0.0001 | > 99 |
| YRFFPQAPDKEL | + 63 x CIM1 | | | | | 6.09 | 0.10 | 1.29 | 0.04 | 2.92 | 0.09 | 30[a] | | 9.00 | 0.04 | 100.0 | 0.4 | < 0.0001 | > 99 |
| | + 1.7 x TTP | | | | | 5.89 | 0.12 | 1.36 | 0.06 | 3.17 | 0.12 | 30[a] | | 9.06 | 0.05 | 100.6 | 0.5 | < 0.0001 | > 99 |
| 894-909 | CNOT1 EY[b] | 4.0 | 0.2 | 0.070 | 0.010 | 8.36 | 0.17 | 5.3 | 0.3 | | | | | 12.34 | 0.14 | 94.9 | 1.1 | 0.0012 | 99 |
| YRFFPQAPDKELHITA | + 63 x CIM1 | 4.75 | 0.17 | 0.068 | 0.006 | 7.60 | 0.14 | 5.1 | 0.3 | | | | | 12.35 | 0.11 | 95.0 | 0.9 | < 0.0001 | > 99 |
| | + 1.7 x TTP | 4.2 | 0.2 | 0.069 | 0.009 | 8.20 | 0.17 | 5.3 | 0.3 | | | | | 12.39 | 0.14 | 95.3 | 1.1 | 0.0006 | > 99 |
| 986-992 | CNOT1 EY[b] | 5.08 | 0.18 | 0.065 | 0.008 | | | | | | | | | 5.08 | 0.18 | 85 | 3 | 0.1580 | 94 |
| YIEYGQQ | + 63 x CIM1 | 4.9 | 0.2 | 0.066 | 0.010 | | | | | | | | | 4.9 | 0.2 | 82 | 4 | 0.4024 | > 99 |
| | + 1.7 x TTP | 4.0 | 0.3 | 0.052 | 0.009 | 0.9 | 0.3 | 3 | 3 | | | | | 4.82 | 0.18 | 80 | 3 | 0.0019 | 98 |
| 986-999 | CNOT1 EY[b] | 4.11 | 0.19 | 0.1[a] | | 5.97 | 0.15 | 5.3 | 0.4 | | | | | 10.08 | 0.10 | 91.6 | 0.9 | - | > 99 |
| YIEYGQQSRDPPVK | + 63 x CIM1 | 4.28 | 0.10 | 0.062 | 0.003 | 5.65 | 0.08 | 4.77 | 0.19 | | | | | 9.93 | 0.06 | 90.3 | 0.6 | < 0.0001 | > 99 |
| | + 1.7 x TTP | 4.29 | 0.12 | 0.068 | 0.005 | 6.05 | 0.10 | 4.9 | 0.2 | | | | | 10.34 | 0.08 | 94.0 | 0.7 | < 0.0001 | > 99 |
| 987-999 | CNOT1 EY[b] | 4.22 | 0.14 | 0.068 | 0.006 | 6.64 | 0.11 | 5.3 | 0.3 | | | | | 10.85 | 0.09 | 108.5 | 0.9 | < 0.0001 | > 99 |
| IEYGQQSRDPPVK | + 63 x CIM1 | 4.1 | 0.2 | 0.060 | 0.008 | 6.18 | 0.19 | 4.8 | 0.4 | | | | | 10.29 | 0.16 | 102.9 | 1.6 | < 0.0001 | > 99 |
| | + 1.7 x TTP | 4.3 | 0.3 | 0.061 | 0.009 | 6.52 | 0.20 | 5.4 | 0.5 | | | | | 10.78 | 0.17 | 107.8 | 1.7 | 0.0007 | > 99 |
| 988-999 | CNOT1 EY[b] | 3.3 | 0.3 | 0.1[a] | | 5.1 | 0.2 | 5.6 | 0.7 | | | | | 8.41 | 0.13 | 93.4 | 1.5 | < 0.0001 | > 99 |
| EYGQQSRDPPVK | + 63 x CIM1 | 3.29 | 0.18 | 0.070 | 0.010 | 4.98 | 0.15 | 4.5 | 0.4 | | | | | 8.27 | 0.11 | 91.9 | 1.3 | < 0.0001 | > 99 |
| | + 1.7 x TTP | 3.40 | 0.20 | 0.1[a] | | 5.03 | 0.15 | 5.8 | 0.5 | | | | | 8.43 | 0.10 | 93.7 | 1.1 | < 0.0001 | > 99 |
| 989-999 | CNOT1 EY[b] | 2.41 | 0.18 | 0.1[a] | | 7.33 | 0.14 | 5.3 | 0.3 | | | | | 9.74 | 0.09 | 121.8 | 1.2 | < 0.0001 | > 99 |
| YGQQSRDPPVK | + 63 x CIM1 | 2.59 | 0.14 | 0.095 | 0.016 | 6.73 | 0.12 | 5.2 | 0.2 | | | | | 9.32 | 0.09 | 116.6 | 1.1 | < 0.0001 | > 99 |
| | + 1.7 x TTP | 2.54 | 0.14 | 0.168 | 0.056 | 7.00 | 0.16 | 5.2 | 0.2 | | | | | 9.54 | 0.07 | 119.3 | 0.8 | < 0.0001 | > 99 |

[a] Fixed values, according to (Gemmecker et al., 1993; Hoofnagle et al., 2001; Mandell et al., 2001; Rutkowska-Wlodarczyk et al., 2008).

[b] CNOT1 EY is CNOT1(800-999) E893A/Y900A.

**Supplementary Table T6 to Supplementary Fig 2a and S7.**

Apparent local dissociation constants ($K_d^{local}$) and maximum number of protons protected against HDX ($\Delta H_{amid}$) observed for subsequent pepsin-generated CNOT1(800-999) peptides due to N-CIM1 binding. The $\Delta H_{amid}$ values are corrected for individual back-exchange of different CNOT1 peptides, determined in the independent experiment. The non-redundant CNOT1 peptides for which both the $K_d^{local}$ is low (≤ *ca*. 100 µM) and $\Delta H_{amid}$ is high (≥ *ca*. 40%) are marked bold.

| Position in sequence | Sequence | $K_d^{local}$ | δ $K_d^{local}$ | $\Delta H_{amid}$ | δ $\Delta H_{amid}$ | $\Delta H_{amid}/D_{max}$ | δ $\Delta H_{amid}/D_{max}$ |
|---|---|---|---|---|---|---|---|
| | | [µM] | | [Da] | | % | |
| 800-819 | NNDPFVQRKLGTSGLNQPTF | 11.1 | 1.7 | 1.98 | 0.13 | 12% | 1% |
| 800-824 | NNDPFVQRKLGTSGLNQPTFQQTDL | 11 | 2 | 3.4 | 0.3 | 15% | 1% |
| 825-835 | SQVWPEANQHF | 70 | 30 | 1.8 | 0.2 | 20% | 2% |
| 825-847 | SQVWPEANQHFSKEIDDEANSYF | 110 | 20 | 13.2 | 1.3 | 63% | 6% |
| 839-846 | IDDEANSY | 56 | 9 | 6.4 | 0.5 | 92% | 7% |
| **839-847** | **IDDEANSYF** | **30** | **6** | **6.6** | **0.8** | **82%** | **10%** |
| **847-859** | **FQRIYNHPPHPTM** | **25** | **6** | **3.3** | **0.2** | **36%** | **3%** |
| 848-859 | QRIYNHPPHPTM | 120 | 30 | 3.9 | 0.4 | 49% | 5% |
| 860-865 | SVDEVL | 600 | 500 | 2.7 | 1.1 | 54% | 23% |
| 866-883 | EMLQRFKDSTIKREREVF | 2500 | 6000 | nd[a] | nd[a] | | |
| 868-883 | LQRFKDSTIKREREVF | 1700 | 1000 | nd[a] | nd[a] | | |
| **894-909** | **YRFFPQYPDKELHITA** | **111** | **13** | **7.8** | **0.5** | **60%** | **4%** |
| 898-909 | PQYPDKELHITA | 130 | 20 | 6.9 | 0.5 | 69% | 5% |
| 911-923 | LFGGIIEKGLVTY | 800 | 400 | nd[a] | nd[a] | | |
| 912-922 | FGGIIEKGLVT | 530 | 140 | nd[a] | nd[a] | | |
| 912-923 | FGGIIEKGLVTY | 770 | 120 | nd[a] | nd[a] | | |
| 924-928 | MALGL | 600 | 100 | 4.3 | 0.4 | 106% | 11% |
| 924-930 | MALGLAL | 570 | 60 | nd[a] | nd[a] | | |
| 937-945 | LRKPFGSKM | 900 | 700 | 5 | 2 | 69% | 33% |
| 952-966 | ALDRFKNRLKDYPQY | 700 | 300 | 5.7 | 1.6 | 44% | 12% |
| 954-966 | DRFKNRLKDYPQY | 600 | 200 | 3.6 | 0.7 | 33% | 6% |
| 954-972 | DRFKNRLKDYPQYCQHLAS | 233 | 6 | 3.4 | 0.4 | 20% | 2% |
| 986-999 | YIEYGQQSRDPPVK | 480 | 140 | 6.7 | 1.0 | 61% | 9% |

[a] For peptides where saturation was not obtained due to a very weak conformational effect, the $K_d^{local}$ value is an estimate, and $\Delta H_{amid}$ was not determined (nd).

**Supplementary Table T7 to Fig 2.**
Apparent equilibrium dissociation constants, $K_d$, for interactions of CNOT1 with GW182 SD or TTP protein fragments.

| CNOT1 protein form | Binding partner | $K_d$ [μM] | Experiment type |
|---|---|---|---|
| GST-CNOT1(800-999) | GW182 SD10 | 6.6 ± 2.1 | FCS |
| CNOT1(800-999) | TTP peptide | 2 | ITC[a] |

[a] determined by (Fabian *et al*., 2013) using isothermal titration calorimetry, shown for comparison

**Supplementary Table T8.**
Primers used to generate CNOT1 mutants.

| Short name | Name | Template | Forward primer | Reverse primer |
|---|---|---|---|---|
| F | F847A | wt | TGAAGCAAACAGCTAGCTCCAGCGAA TATATAA | TTATATATTCGCTGGAGCTAGCTGTTTG CTTCA |
| FQ | F847A/Q 848A | F | GCAAACAGCTATGCCGCGCGAATATA TAATCATC | GATGATTATATATTCGCGCGGCATAGC TGTTTGC |
| PT | P857A/T 858A | wt | TAATCATCCACCACATGCAGCCATGT CTGTTGATGAG | CCTCATCAACAGACATGGCTGCATGTG GTGGATGATTA |
| E893A | E893A | wt | AGGAACTTGTTTGAAGCATATCGTTT TTTTCCC | GGGAAAAAAACGATATGCTTCAAACA AGTTCCT |
| EY | E893A/Y 900A | E893A | CGTTTTTTTCCCCAGGCTCCTGATAAA GAGTTA | TAACTCTTTATCAGGAGCCTGGGGAAA AAAACG |
| FQEY | F847A/Q 848A/ E893A/Y 900A | EY | GCAAACAGCTATGCCGCGCGAATATA TAATCATC | GATGATTATATATTCGCGCGGCATAGC TGTTTGC |
| FQPT | F847A/Q 848A /P857A/ T858A | PT | GCAAACAGCTATGCCGCGCGAATATA TAATCATC | GATGATTATATATTCGCGCGGCATAGC TGTTTGC |
| PTEY | P857A/T 858A/ E893A/Y 900A | EY | TAATCATCCACCACATGCAGCCATGT CTGTTGATGAG | CCTCATCAACAGACATGGCTGCATGTG GTGGATGATTA |
| Y851A | Y851A | wt | CTATTTCCAGCGAATAGCTAATCATCC ACCACATC | GATGTGGTGGATGATTAGCTATTCGCT GGAAATAG |
